# Development of Potent and Cell Active 5-Azaindole-Based Tau Tubulin Kinase Inhibitors

**DOI:** 10.64898/2026.04.27.721186

**Authors:** Raymond Flax, Andrea Lacigová, Stefanie Howell, Haoxi Li, Frances M. Bashore, Lukáš Čajánek, Alison D. Axtman

**Author notes:** **Corresponding Author Alison D. Axtman -** Structural Genomics Consortium, UNC Eshelman School of Pharmacy, University of North Carolina at Chapel Hill, Chapel Hill, North Carolina 27599, United States.

## Abstract

We have developed and characterized a potent and cell active tau tubulin kinase 1 and 2 (TTBK1 and TTBK2) inhibitor, **13**. Compound **13** demonstrates in-cell, kinome-wide selectivity, and potently inhibits both TTBK1 and TTBK2. As part of our medicinal chemistry campaign, we also identified a structurally similar negative control, compound **5**, which lacks in-cell affinity for TTBK1 and TTBK2. Based on their substrates, which include TDP-43, tau, and tubulin, TTBK1 and TTBK2 inhibition has been pursued as a therapeutic approach for Alzheimer’s disease, frontotemporal lobe dementia, and amyotrophic lateral sclerosis. TTBK2 is also an effector of ciliogenesis, acting in concert with CEP164, CP110, and CEP83 to initiate the biogenesis of primary cilia. The development of selective chemical tools for these kinases facilitates investigation into TTBK1/2-mediated pathways and potential disease-altering ramifications linked to their pharmacological perturbation.

## Introduction

Tau tubulin kinases (TTBKs) are a small family of serine/threonine/tyrosine kinases within the larger CMGC group, which consists of Cyclin dependent, Mitogen-activated protein, Glycogen synthase, and CDC-like kinases. TTBKs contain two isoforms: tau tubulin kinase 1 (TTBK1) and tau tubulin kinase 2 (TTBK2). The kinase domains of TTBK1 and TTBK2 are homologous, with 88% identity and 96% similarity.^1, 2^ TTBK1 and TTBK2 also have similar catalytic residues, which are K63/K50 and D164/D141, respectively.^1, 2, 3^ Outside of their kinase domains, whole sequence identity falls to 35%. This sequence variation can help to rationalize the differential roles of TTBK1 and TTBK2 in biological signaling pathways.^1^ For example, the TTBK2 C-terminus contains a CEP164-binding domain, which localizes TTBK2 to the mother centriole where it facilitates the initiation of ciliogenesis. TTBK1 lacks this CEP164 binding domain. Accordingly, TTBK1 can only minimally rescue ciliogenesis in the absence of TTBK2 and cannot replicate the activity of wild-type TTBK2. This is proposed to result, in part, from an inability of TTBK1 to localize to the same regions and make the same protein–protein interactions as TTBK2.^1, 4^

The expression profiles of TTBK1 and TTBK2 also differ. TTBK1 is confined to the central nervous system (CNS), which correlates with its reported phosphorylation of TAR DNA-binding protein 43 (TDP-43), tau, and tubulin at physiologically relevant sites in the brain. This TDP-43 post-translational modification is a hallmark of neurodegenerative diseases such as Alzheimer’s disease, frontotemporal lobe dementia, and amyotrophic lateral sclerosis.^5, 6, 7^ TTBK2 expression is more ubiquitous in the human body, and it serves as a primary regulator of ciliogenesis, facilitating the initiation of this critical process.^8^ TTBK2 is significantly expressed within cerebellum Purkinje cells, hippocampal neurons, midbrain and substantia nigra regions, and the granular cell layer of the CNS. To initiate ciliogenesis, TTBK2 functions at the distal end of the mother centriole where it aids in the removal of capping protein CP110 and recruits the IFT proteins necessary for the ensuing assembly of the ciliary axonemal microtubules. In the absence of TTBK2, ciliogenesis is impaired and, correspondingly, TTBK2 truncation mutations are linked to a rare genetic form of ataxia known as spinocerebellar ataxia type 11 (SCA11).^3, 9^

Although they play important functions in biological and neurodegenerative disease mechanisms, knowledge around TTBK1 and TTBK2 remains limited. To stimulate characterization of these kinases, TTBK1 and TTBK2 were added as understudied kinases to the NIH Illuminating the Druggable Genome (IDG) initiative.^10^ Kinases on this understudied list, which was updated most recently in 2019, represent protein targets that lack and would benefit from high-quality small molecule inhibitors that we term chemical probes. Chemical probes have been defined by the Structural Genomics Consortium (SGC) as compounds with potent target engagement, as reflected by a biochemical IC_50_ <100 nM and cellular IC_50_ <1 µM. Chemical probes must also be highly selective when broadly profiled and are required to demonstrate >30-fold selectivity versus proteins in the target class, excluding highly homologous family members.^11,12^ To increase the robustness of experiments executed with chemical probes, a fully characterized, structurally related negative control is simultaneously released to be used in tandem. Finally, the chemical probe and negative control are distributed without restrictions to the scientific community to facilitate research around historically understudied and/or poorly characterized protein targets. These highly characterized probes are valuable research reagents used for elucidating biological functions of a target protein in a cellular and/or *in vivo* context.^11-13^ These tools also have utility in target validation as part of drug discovery campaigns, as they complement classic genetics-based approaches, and can help elucidate the pathological mechanism(s) of idiopathic and known disease-causing proteins.^11-13^

To date, only a few small molecule inhibitors targeting TTBK1/2 have been published. These compounds are not fully characterized to meet chemical probe criteria and lack screening data versus TTBK1, TTBK2, and/or the broader kinome. Furthermore, confirmation of direct target engagement with TTBK1 and TTBK2 in cells has not been established for most due to the absence of a NanoBRET target engagement assay. However, functional in-cell activity measurements have been provided for these compounds. TTBK1-IN-1 (Figure 1), published by Biogen, is the best available TTBK1/2 inhibitor based on its biochemical IC_50_ values of 2.7 nM and 1.6 nM for TTBK1 and TTBK2, respectively, in a TR-FRET-based assay.^6^ Cellular characterization of TTBK1 inhibition was assessed via phosphorylation of tau at Ser422 using an HTRF-based assay. TTBK1-IN-1 demonstrated an IC_50_ of 315 nM in this cell-based tau phosphorylation assay, but no information regarding TTBK2 inhibition was provided. To assess its kinome-wide selectivity, TTBK1-IN-1 was profiled at 1 µM in the DiscoverX panel of 468 cell-free binding kinases, however, TTBK1 and TTBK2 are not in this panel. Only two kinases were found to bind with high affinity and be potently inhibited by TTBK1-IN-1: CK1δ and PRKD1. Orthogonal enzymatic assays run in dose–response format confirmed that TTBK1-IN-1 inhibits CK1δ and PRKD1 with IC_50_ values of 40 nM and 3 nM, respectively. When used *in vivo*, TTBK1-IN-1 inhibited tau phosphorylation at Ser422 in the brains of hypothermia-induced mice and developing rats, highlighting its suitability for *in vivo* indications as well as favorable CNS penetrance. While TTBK1-IN-1 is the most advanced inhibitor, cellular target engagement data are needed to support its nomination as a TTBK1/2 chemical probe. Nozal et al published TTBK1-IN-2 (Figure 1), which demonstrates cell-free IC_50_ values of 0.24 µM and 4.22 µM for TTBK1 and TTBK2, respectively, making it a weaker inhibitor when compared to TTBK1-IN-1.^5^ Finally, we previously exemplified TTBK1/2-IN-3 (Figure 1), which also has weaker inhibition of TTBK1 and TTBK2 when compared to TTBK1-IN-1. TTBK1 and TTBK2 cell-free enzymatic IC_50_ values of TTBK1/2-IN-3 were 579 nM and 258 nM, respectively. To complement these cell-free measurements, we developed and executed a NanoBRET assay for TTBK1 and TTBK2 to demonstrate that TTBK1/2-IN-3 had weak TTBK affinity in cells (IC_50_ = 11 and 6.4 μM, respectively).^3^ Based on these molecules, TTBK1 and TTBK2 remain without a fully characterized inhibitor that meets chemical probe criteria.

**Figure 1.**
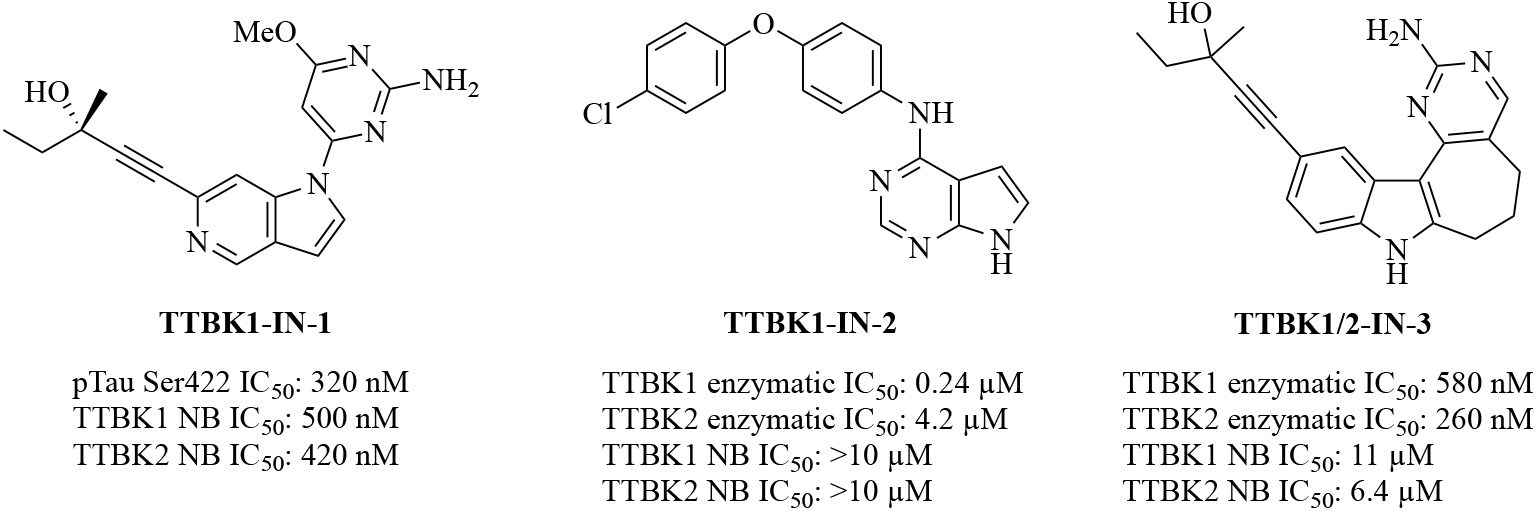
Structures of commercially available TTBK inhibitors with associated cell-free and in-cell potency data. NB = NanoBRET.

Based on the cell-free potency of TTBK1-IN-1 and its promising *in vivo* activity,^6^ we developed analogs of this compound. Building from the 5-azaindole core of TTBK1-IN-1, we synthesized and fully characterized a potent and selective small molecule TTBK1/2 inhibitor, compound **13**. Utilizing a medicinal chemistry optimization strategy guided by direct cellular target engagement assay data for TTBK1 and TTBK2, we modified the 2-aminopyrimidine hinge-binding moiety appended to the 5-azaindole core to impart high affinity for TTBK1/2 in cells. Our structural modifications produced a selective cellular profile, as assessed in panels of 192–300 NanoBRET cellular target engagement assays using full length human kinases. A structurally related negative control compound (**5**) was developed that bears a homologated propargylic alcohol on the alkyne terminus and lacks affinity for TTBK1/2. Compound **13** was further profiled for its ability to disrupt ciliogenesis and found to ablate autophosphorylation of TTBK2, inhibit the recruitment of IFT88 complexes, and impair cilia formation.

**Scheme 1.**
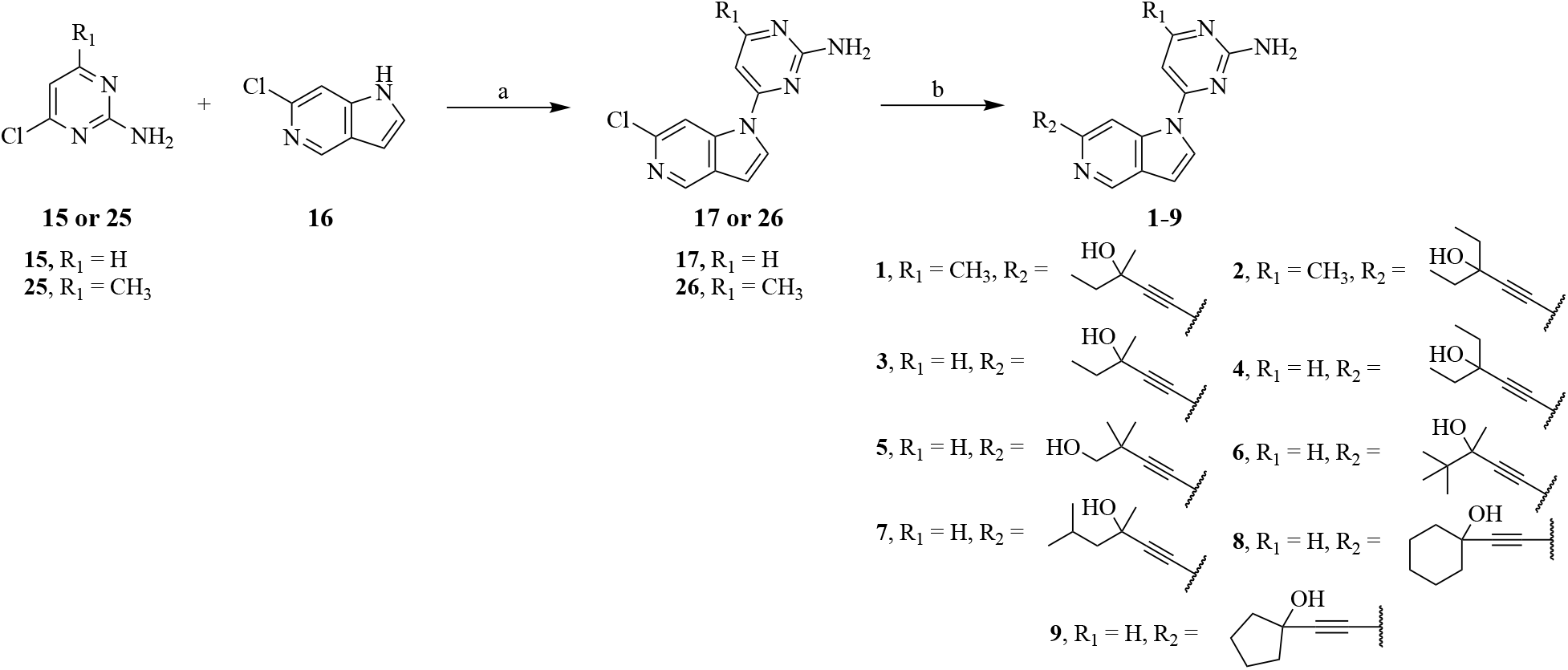
Synthesis of analogs 1–9^a^. ^*a*^ Reagents and conditions: (a) NaH, DMF, 60°C, 16 h, 60*–*80%; (b) alkyne R_2_, K_2_CO_3_, Pd(OAc)_2_, DPPP, DMF, 85°C, 16 h, 1*–*60%.

### Chemistry

Compounds **1**–**9** were assembled via a common synthetic route (Scheme 1). Briefly, the core of this chemical series was synthesized using nucleophilic aromatic substitution conditions to combine commercially available pyrimidine **15** or **25** and 5-azaindole **16**. The final compounds were assembled using intermediate **17** or **26** and Sonogashira coupling conditions with their respective alkynes.

**Scheme 2.**
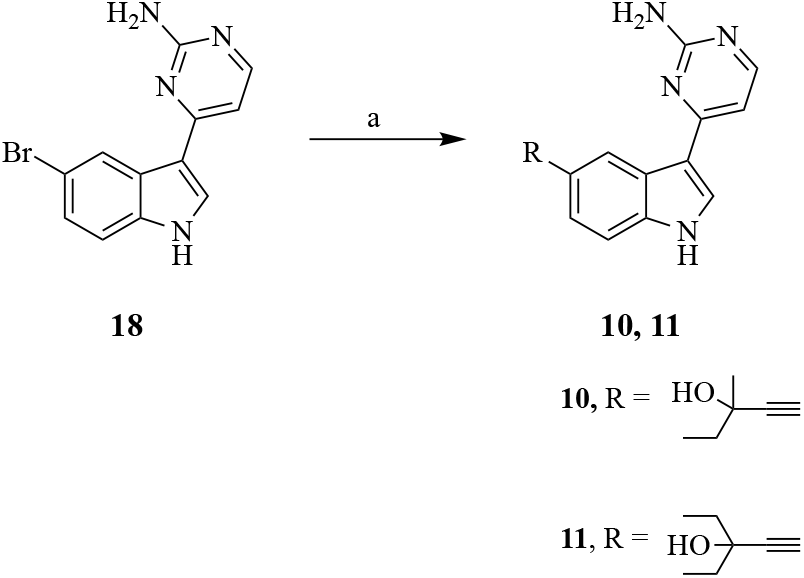
Synthesis of analogs 10 and 11^a^. ^*a*^ Reagents and conditions: (a) alkyne R, DIPA, PdCl_2_(PPh_3_)_2_, CuI, 1-propanol, 85°C, 16 h, 2*–*9%

Compounds **10** and **11** were synthesized from commercially available 2-aminopyrimidine functionalized 6-bromo indole, **18**, via Sonogashira coupling conditions with their respective alkynes to generate the final products (Scheme 2).

Compound **14** was prepared utilizing nucleophilic aromatic substitution conditions with commercially available functionalized pyrimidine **19** and 6-chloro-5-azaindole **16** (Scheme 3). The final compound, **14**, was synthesized via Sonogashira coupling conditions with 3-methylpent-1-yn-3-ol.

**Scheme 3.**
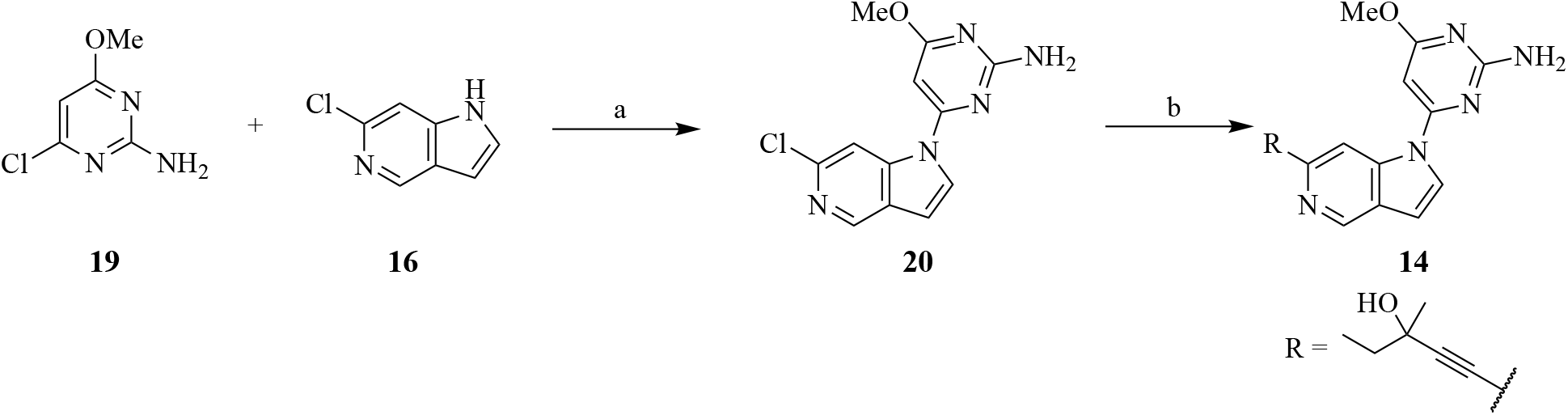
Synthesis of analog 14^a^. ^*a*^ Reagents and conditions: (a) NaH, DMF, 60°C, 16 h, 70%; (b) 3-methylpent-1-yn-3-ol, K_2_CO_3_, Pd(OAc)_2_, DPPP, DMF, 85°C, 16 h, 4%.

Compounds **23** and **24** were synthesized from a commercially available functionalized 6-bromo indole, **18** (Scheme 4). Intermediate **18** was treated with sodium hydride and 3-((*tert*-butoxycarbonyl)amino)propyl 4-methylbenzenesulfonate, followed by acidic cleavage of the Boc-protecting group to yield intermediate **21**. Compound **21** was converted to intermediate **22** via Sonogashira coupling conditions with 3-ethylpent-1-yn-3-ol. The final NanoBRET tracer candidates (**23** and **24**) were generated via an amide coupling between **22** and the commercially available dye, NanoBRET590 SE.

**Scheme 4.**
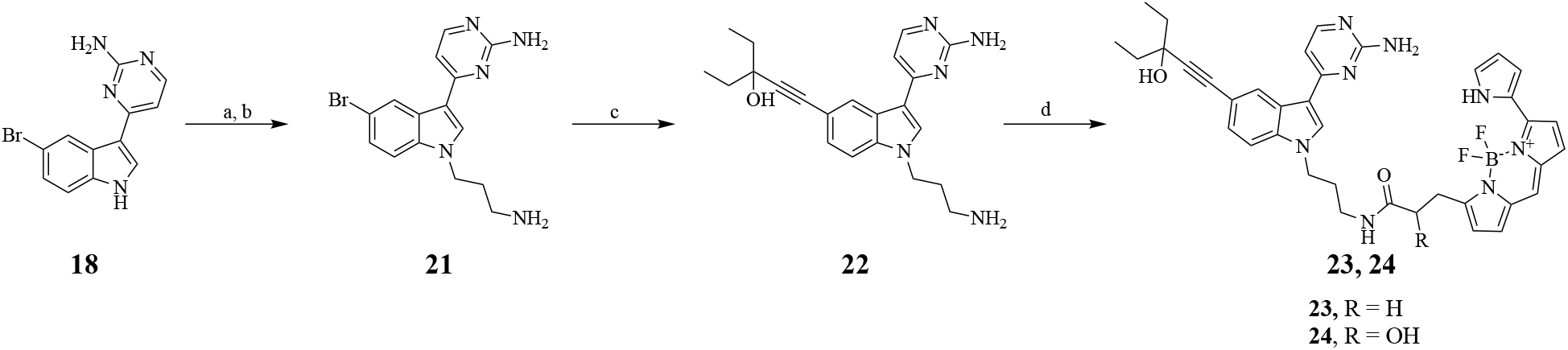
Synthesis of candidate tracers 23 and 24^a^. ^*a*^ Reagents and conditions: (a) 3-((*tert*-butoxycarbonyl)amino)propyl 4-methylbenzenesulfonate, NaH, DMF, r.t., 16 h; (b) TFA, DCM, 16 h, 22% over two steps; (c) 3-ethylpent-1-yn-3-ol, DIPA, PdCl_2_(PPh_3_)_2_, CuI, 1-propanol, 85°C, 16 h, 30%; (d) NanoBRET590 SE, DIPEA, DMF, r.t., 30 min, 7*–*30%.

## Results and Discussion

### Benchmarking Target Engagement of Commercial TTBK Inhibitors

Based on the promising data generated for TTBK1-IN-1 coupled with its synthetic accessibility, we opted to use the 5-azaindole core and published structure–activity relationship (SAR) data to support our optimization campaign. Initially, we benchmarked the affinity of published TTBK inhibitors in our previously developed TTBK1 and TTBK2 NanoBRET (NB) cellular target engagement assays.^3^ These assays benefit from simultaneous quantification of on-target affinity and confirmation of cellular permeability because they are executed in intact cells. Thus, use of these assays as part of a chemical probe development campaign is proposed to expedite delivery of a cell-active compound that binds its intended protein target. When evaluated in these NanoBRET assays, TTBK1-IN-1 demonstrated an IC_50_ of 500 nM for TTBK1 and 420 nM for TTBK2 (Figure 1). TTBK1-IN-2, in contrast, exhibited IC_50_ values of greater than 10 µM for both isoforms (Figure 1). We previously reported that TTBK1/2-IN-3 also demonstrates weaker affinity for TTBK1 and TTBK2 when compared to TTBK1-IN-1 (Figure 1).^3^ For comparison, Figure 1 also includes published cell-free and functional cellular data. The NanoBRET assay data supported our approach centered around optimization of TTBK1-IN-1 for TTBK1/2 affinity and selectivity.

### Published Co-Crystal Structure Informs SAR Expansion

The co-crystal structure of TTBK1 bound to BGN18, a structurally similar precursor to TTBK1-IN-1, showed that it adopts a unique type 1.5 binding pose (Figure 2).^6^ A portion of the 2-aminopyrimdine ring of BGN18, highlighted in purple, makes critical hydrogen bonds to Glu110 and Glu108 within the hinge region. The BGN18 propargylic alcohol, colored red, extends past the gate keeper residue Met107, with the DFG motif flipped in, and hydrogen bonds to Phe177 in the back pocket. The aliphatic chains on the alkyne terminus make hydrophobic interactions with aliphatic residues in the back pocket (Leu81) that support its high affinity binding. Finally, the C5-nitrogen of the 5-azaindazole core hydrogen bonds with Lys63 to anchor the scaffold in the binding pocket. Due to the high sequence similarity of the kinase domains, it is proposed that similar interactions dictate binding of BGN18, TTBK1-IN-1, and related analogs to TTBK2.

**Figure 2.**
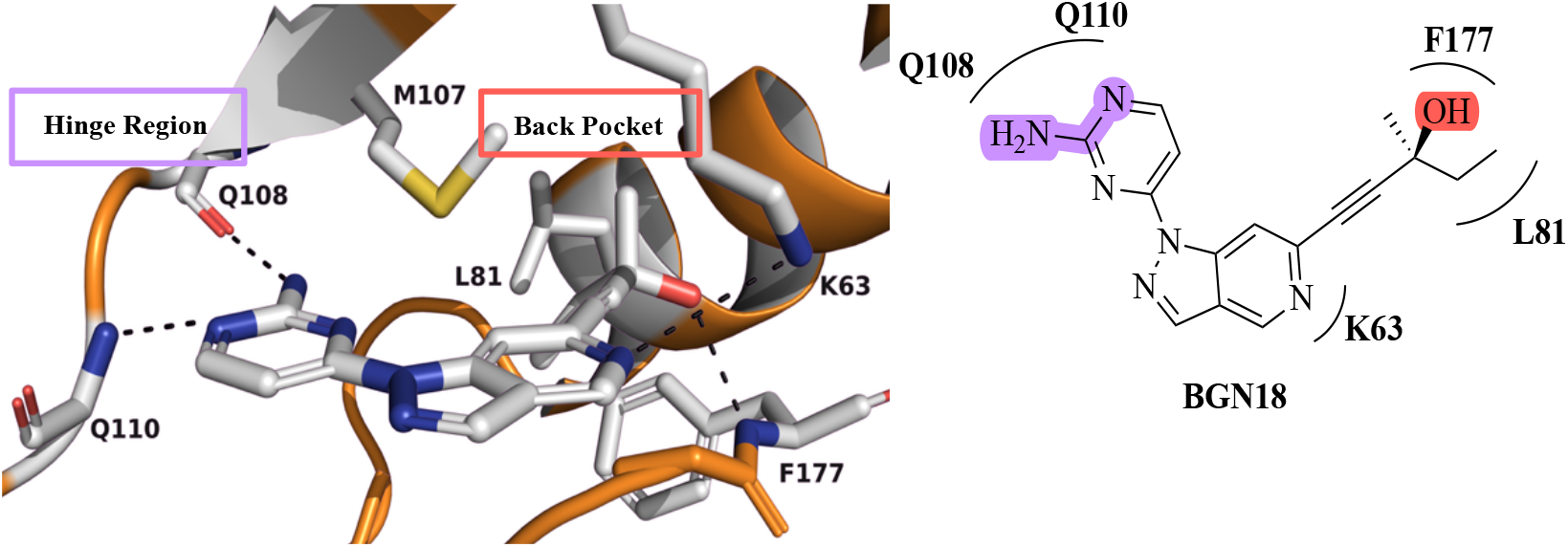
Co-crystal structure of BGN18 bound to TTBK1 (PDB code: 7JXY). Key residue interactions highlighted.

### C5-Aminopyrimidine Position Hinge-Binding Modifications are Tolerated

Our design strategy was guided by the published co-crystal structure (Figure 2) and aimed to generate compounds that maintained key hydrogen bonds with Glu110 and Glu108 in the hinge region and Lys53 as well as hydrophobic contacts in the back pocket. We sought to explore regions of the molecule where structural modifications had previously been exemplified without reducing TTBK1 affinity/activity and those where limited SAR had been probed. Our first set of analogs, compounds **1** and **2**, were designed to explore whether the hinge region of these kinases would bind compounds with a small aliphatic group at the C5 position of the 2-aminopyrimidine ring (methyl compared to methoxy in TTBK1-IN-1). Based on the SAR established by Biogen, TTBK1 tolerates small aliphatic groups in the hinge region, but incorporation of these groups did not impart significant increases in affinity based on cell-free activity data.^6^ As cell-free and cell-based data do not always align and limited examples of analogs employing this design strategy were exemplified by Biogen,^3, 6, 14, 15^ we expanded the scope through synthesis of two additional analogs. Compound **1**, bearing a methyl and ethyl group flanking the propargylic alcohol, exhibited IC_50_ values of 1200 nM for TTBK1 and 980 nM for TTBK2 (Table 1). Compound **2**, which was designed to replace the methyl group with an ethyl group on the alkyne terminus when compared to compound **1**, demonstrated a slight increase in affinity for TTBK1 (IC_50_ = 500 nM) but maintained a similar IC_50_ for TTBK2 (970 nM) versus compound **1**. That these analogs still bind with sub-micromolar affinity to TTBK1/2 in a range that is similar to TTBK1-IN-1 supports that modification of the C5 position of the 2-aminopyrimidine ring is tolerated and that it may be further tuned to result in analogs with enhanced affinity.

**Table 1.**
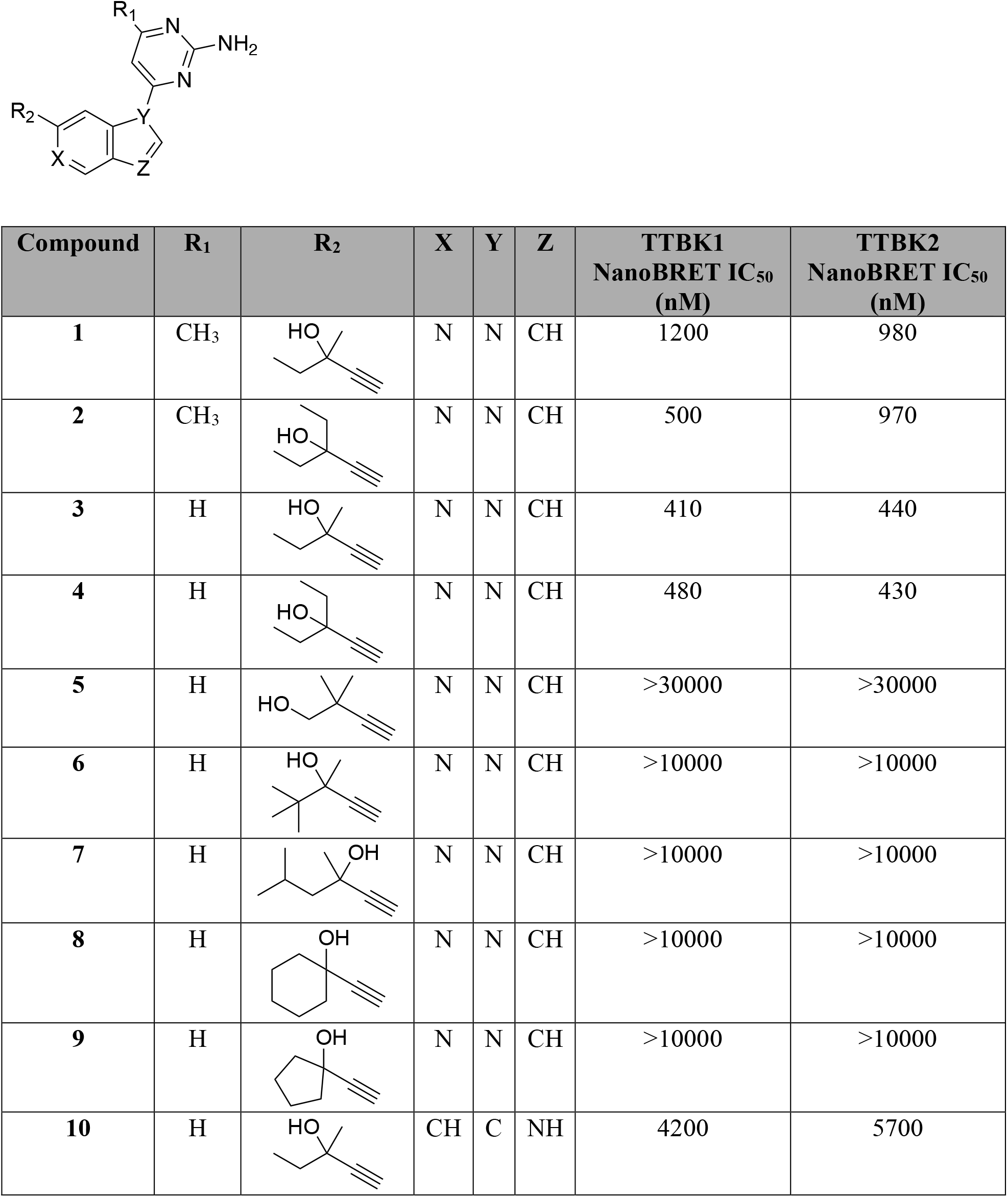
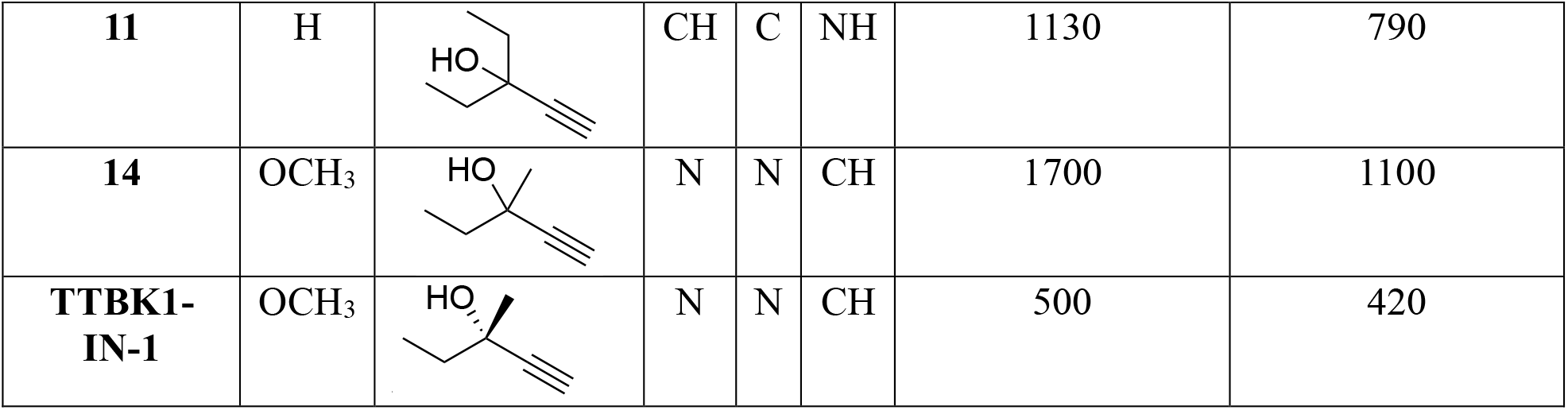
TTBK1 and TTBK2 NanoBRET affinity data for synthesized analogs and TTBK1-IN-1.

| Compound | R<sub>1</sub> | R<sub>2</sub> | X | Y | Z | TTBK1<br>NanoBRET IC<sub>50</sub><br>(nM) | TTBK2<br>NanoBRET IC<sub>50</sub><br>(nM) |
| --- | --- | --- | --- | --- | --- | --- | --- |
| 1 | CH<sub>3</sub> |  | N | N | CH | 1200 | 980 |
| 2 | CH<sub>3</sub> |  | N | N | CH | 500 | 970 |
| 3 | H |  | N | N | CH | 410 | 440 |
| 4 | H |  | N | N | CH | 480 | 430 |
| 5 | H |  | N | N | CH | >30000 | >30000 |
| 6 | H |  | N | N | CH | >10000 | >10000 |
| 7 | H |  | N | N | CH | >10000 | >10000 |
| 8 | H |  | N | N | CH | >10000 | >10000 |
| 9 | H |  | N | N | CH | >10000 | >10000 |
| 10 | H |  | CH | C | NH | 4200 | 5700 |
| **11** | H |  | CH | C | NH | 1130 | 790 |
| **14** | OCH<sub>3</sub> |  | N | N | CH | 1700 | 1100 |
| **TTBK1-IN-1** | OCH<sub>3</sub> |  | N | N | CH | 500 | 420 |

### Hinge Region Simplification Improves Affinity

Next, we explored analogs that removed the electron-donating group from the C5 position of the 2-aminopyrimdine ring (methoxy in TTBK1-IN-1), resulting in less bulky hinge-binding moieties in compounds **3**–**9**. For this series of analogs, the simplified hinge-binding 2-aminopyrimidine was paired with different alkynes that probed the space of the TTBK1/2 back pocket. Variable hydrophobic bulk was incorporated at the alkyne terminus, while maintaining the essential alcohol group. This alcohol was kept in the propargylic position in all analogs with the exception of compound **5**, which was homologated with a methylene. Analogs **3** and **4** demonstrated improved affinity for the TTBKs (Table 1). A direct comparison of analogs **1** and **3** supports that the simplified hinge-binding group was preferred by TTBK1/2. This trend is maintained when comparing analogs **2** and **4**. Incorporation of the methyl-ethyl propargylic alcohol found in TTBK1-IN-1 and TTBK1/2-IN-3 resulted in analogs with higher affinity than those with a diethyl propargylic alcohol.^3, 6^ These results support the previously reported conclusion that the TTBK1 back pocket is sensitive to hydrophobic bulk, as just a simple methyl to ethyl swap reduced binding affinity. The remainder of the analogs in this series (**5**–**9**) reinforce this finding, as all have additional hydrophobic bulk at the alkyne terminus and binding of these analogs to TTBK1 and TTBK2 has been abolished.

### 5-Azaindole Core Identified as Optimal

Compounds **10** and **11** were designed to evaluate whether an alternative core, an indole, could be used to generate analogs with similar TTBK1/2 affinity to those built upon a central 5-azaindole. These indole analogs were weaker binders to TTBK1 and TTBK2 (Table 1). When the same 2-aminopyrimidine hinge binder and alkyne are appended, the 5-azaindole analogs display better affinity than the indole. This is exemplified when considering the comparison of the matched pairs of compounds **10** and **3** as well as **11** and **4**. There is a more considerable drop in affinity when considering compounds **10** and **3** versus compounds **11** and **4**. The high affinity of compound **3** supports that it achieves optimal binding to TTBK1/2. Overall, the 5-azaindole core is proposed to make key interactions that are lost when moving to the indole core.

### Lead Compound Confirmed as TTBK Inhibitor

Our SAR campaign generated compound **3** as a lead to move forward for further characterization. Compound **3** demonstrated the highest in-cell affinity for TTBK1 and TTBK2 of all the published and novel analogs we evaluated in the TTBK1/2 NanoBRET assays. Next, to orthogonally assess whether binding to these kinases in cells correlated with inhibition of their enzymatic activity, compound **3** and TTBK1-IN-1 were profiled in TTBK1 and TTBK2 radiometric enzyme inhibition assays. TTBK1-IN-1 was found to potently inhibit TTBK1 and TTBK2 enzymatic activity with IC_50_ values of 84 nM and 30 nM for TTBK1 and TTBK2, respectively. Compound **3** was equipotent to TTBK1-IN-1 in these enzymatic assays, exhibiting IC_50_ values of 80 nM for TTBK1 and 28 nM for TTBK2. For both compounds, a slight bias in inhibition of TTBK2 over TTBK1 was noted. This trend was not echoed in the corresponding NanoBRET assay data.

### Compound 3 Demonstrates Excellent In-Cell Kinome-Wide Selectivity

Based on its high TTBK affinity, compound **3** was evaluated to ascertain its in-cell kinome-wide selectivity. Binding to a panel of 192 wild type human kinases (Figure 3A) was analyzed at a single concentration (1 µM) to generate percent occupancy values. TTBK1 and TTBK2 were not included in this panel.^16^ Only three of the 192 kinases assessed, MAPK6, MET, and CDKL5, demonstrated percent occupancy >30%, which is below the 50% threshold suggested by Promega to trigger follow-up assays in dose–response format (Figure 3 and Table S1). When these three follow-up NanoBRET assays were run in dose–response format, compound **3** did not bind (IC_50_ >10 µM) (Figure 3B). Based on the combined NanoBRET data for 194 kinases, compound **3** selectively engages TTBK1 and TTBK2 in cells and does not demonstrate appreciable binding to the 192 additional kinases profiled. To further assess the kinome-wide selectivity of **3**, we collaborated with Promega to have it evaluated in a panel of 300 wild type human kinase NanoBRET assays (K300) at 1 µM (Figure 4A and Table S2). TTBK1 and TTBK2 were also not included in this panel. To be rigorous, follow-up studies were executed for high affinity binding kinases from the K192 and K300 assays. Only four of the 300 kinases assayed, PIP4K2C, CDK11B, CDK11A, and CK1δ, exhibited a percent occupancy value >50% and thus binding was further evaluated in dose–response format (Figure 4B and 4D). Gratifyingly, the data from Figure 3 repeated. Based on the IC_50_ values in Figures 3B and 4D, we concluded that compound **3** demonstrates excellent in-cell, kinome-wide selectivity and represents a promising TTBK chemical tool for TTBK1 and TTBK2.

**Figure 3.**
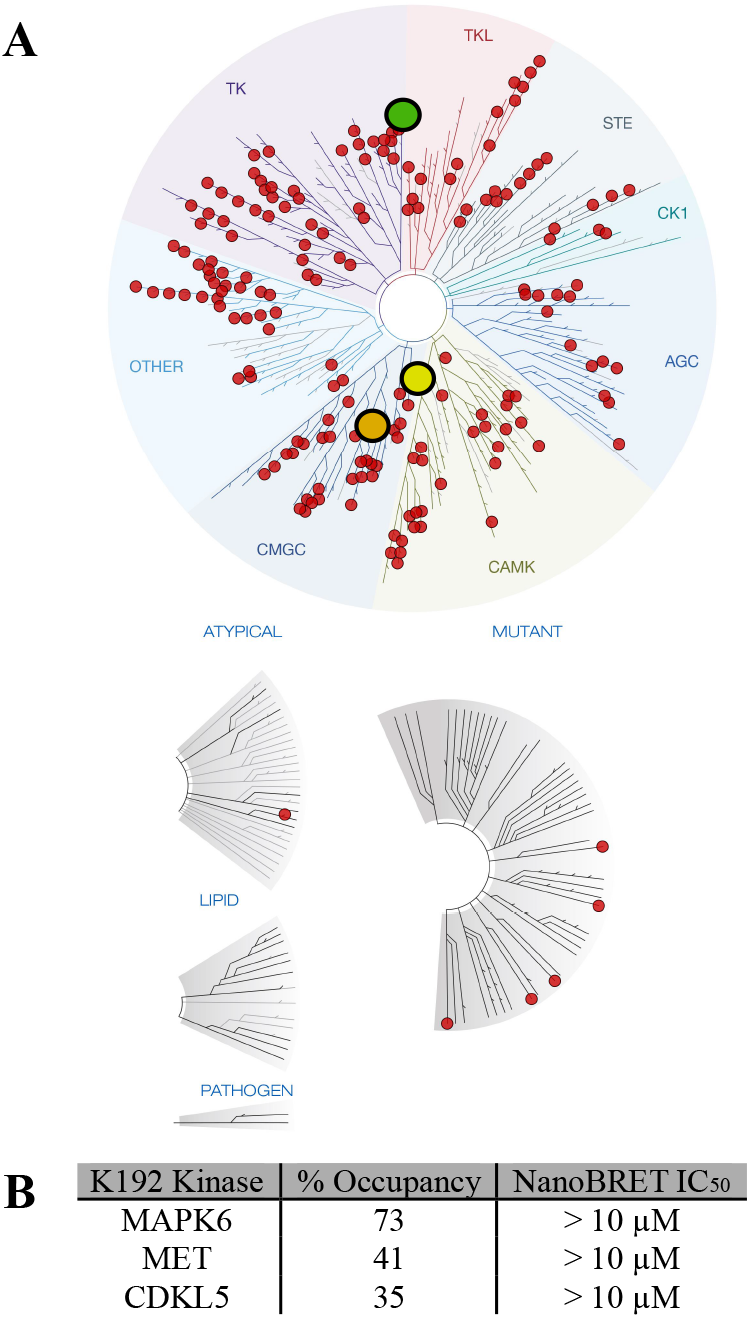
Compound **3** in-cell selectivity (K192) results when analyzed at 1 µM. (A) Dendrogram showing all kinases in K192 panel as red dots and high affinity binding kinases (above >30%) as colored dots: green dot is MET, yellow dot is CDKL5, and orange dot is MAPK6. (B) Summary of the follow-up, full dose–response NanoBRET IC_50_ values generated for the kinases that demonstrated >30% occupancy in the K192 panel.

**Figure 4.**
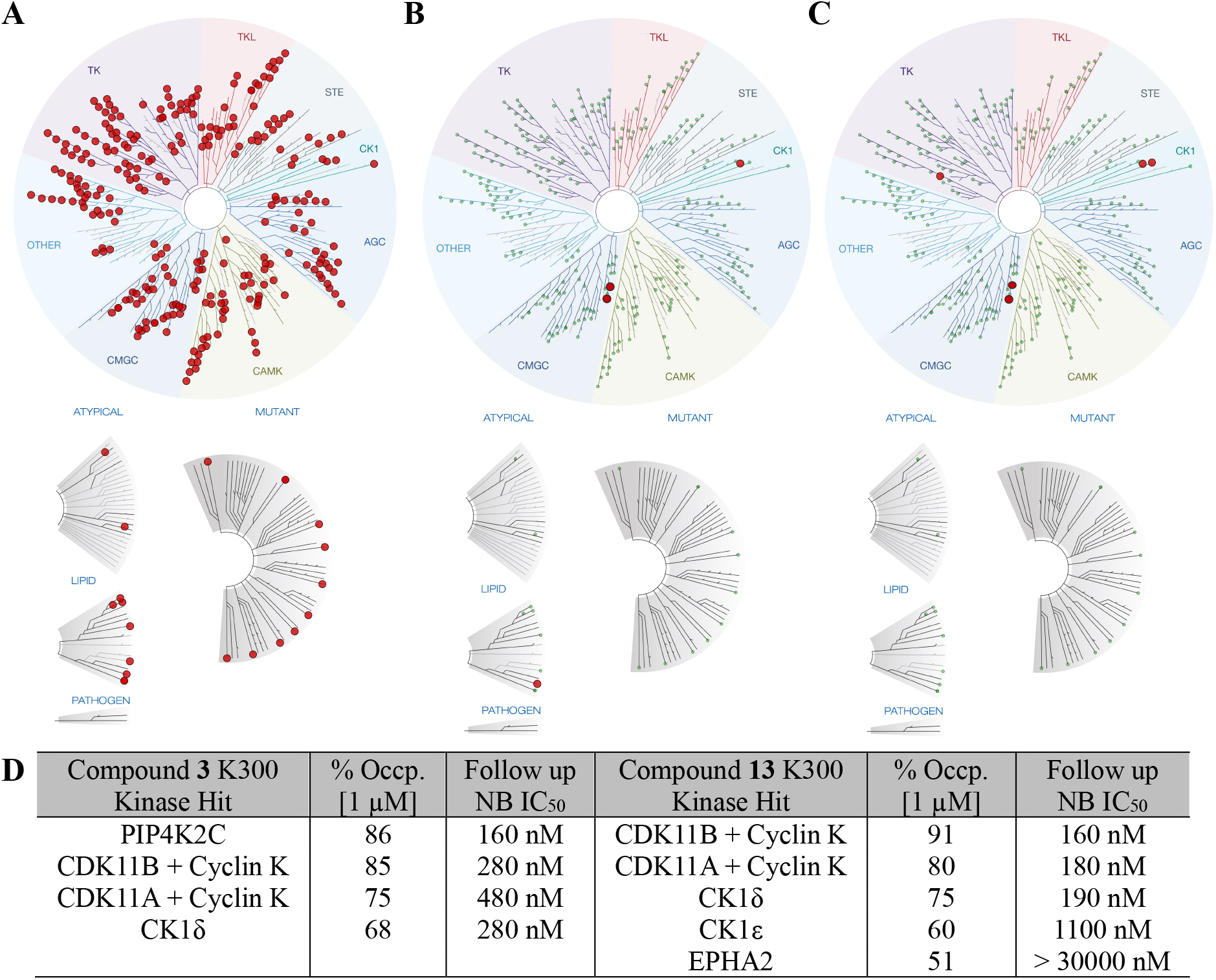
In-cell selectivity profiling in K300 panel. (A) Dendrogram that shows all kinases in K300 panel marked with red dots. (B) Dendrogram highlighting kinases in K300 that bind compound **3** with >50% occupancy (occp.) at 1 µM: PIP4K2C (86%), CDK11B (85%), CDK11A (75%), and CK1δ (68%). (C) Dendrogram depicting kinases in K300 that bind compound **13** with >50% occupancy at 1 µM: CDK11B (91%), CDK11A (80%), CK1δ (75%), CK1ε (60%), and EPHA2 (51%). Data in panels B and C summarize K300 panel profiling in duplicate. (D) Table capturing kinases that bind with >50% occupancy in the K300 when **3** and **13** were profiled at 1 µM and NanoBRET assay IC_50_ values generated via dose–response follow-up.

### Compound 3 is Highly Soluble, Permeable, and Non-Toxic

Based on its promising potency and selectivity data, we next ascertained the physicochemical and “drug-like” properties of **3**. We first quantified the aqueous kinetic solubility of compound **3**, which proved it to be highly soluble in aqueous buffer with an experimental solubility value of 141.8 µM (43.6 µg/mL). To next profile its permeability, compound **3** was sent for PAMPA analysis. This compound is quite permeable, as reflected by its permeability coefficient (P_e_) of 5.45 x 10^-6^ cm/s. This P_e_ value, which correlates with the human fraction absorbed >80%, puts this compound in the permeable category.^17^ Furthermore, compound **3** was evaluated for its ability to impact cell viability and found to be non-toxic at concentrations up to 10 µM for 8 hours (Figure S1). Collectively, our data support that compound **3** is potent in cell-free and cellular target engagement assays, selective across the kinome when profiled in cells, does not impact cell viability at an extended high dose, and demonstrates good physicochemical properties. These qualities solidify compound **3** as an excellent TTBK inhibitor, satisfying multiple of the established SGC chemical probe criteria.^11^

### (*S*)-Enantiomer Outperforms Racemic Mixture

TTBK1-IN-1 is a single (*S*)-enantiomer, defined by the propargylic alcohol geometry on the alkyne terminus. As we had synthesized several analogs with a chiral center (**1, 3, 5, 6, 7, 10**), we sought to understand the influence of chiral separation on TTBK binding affinity. To start, compound **14** was synthesized to assess the TTBK1/2 affinity of racemic TTBK1-IN-1. Comparison of the TTBK1/2 affinities of compound **14** and TTBK1-IN-1 reveals a >2.5-fold loss in affinity upon racemization of the alkyne terminus. This supports that the (*S*)-enantiomer preferentially engages the back pocket of TTBK1/2, resulting in improved cellular affinity of TTBK1-IN-1 versus **14**. It also suggests improved TTBK1/2 affinity may be achieved via chiral separation of our confirmed binders bearing a chiral center. We proposed that a more sensitive assay may be advantageous to distinguish the affinities of independent enantiomers and thus pursued synthesis and characterization of a novel TTBK1/2 NanoBRET tracer to realize this goal.

### Indole Core Affords Access to a Novel TTBK1/2 NanoBRET Tracer

Our first generation NanoBRET tracer enabled the TTBK1/2 target engagement assays used to generate our preliminary data (Table 1).^3^ The recommended working concentration of this tracer to afford a maximal assay window for both TTBK1 and TTBK2 is 2 µM.^16^ The weak in-cell TTBK1/2 affinity of the parent compound (12–16 µM for TTBK1 and TTBK2) is proposed to drive this high concentration requirement.^3^ Compound **11** differs from the parent of the current tracer only at the alkyne terminus. Extension of the methyl on the tracer parent to an ethyl group in compound **11** resulted in slightly improved IC_50_ values of 1130 nM for TTBK1 and 790 nM for TTBK2. Similar to our first-generation tracer, the core scaffold of **11** positions the indole-NH, which is a synthetically tractable site for synthesis, toward solvent. Due to its improved binding to TTBK1/2, compound **11** was chosen as the tracer precursor to deliver a NanoBRET tracer that can be used at a lower concentration, maintain or improve the window of the TTBK1/2 NanoBRET assays, and yield more robust and sensitive assays.

We synthesized the NanoBRET tracers via standard procedures,^3, 18^ and isolated two novel NanoBRET tracers from the same reaction mixture: tracer candidates **23** and **24** (Table 2). When evaluated for their ability to generate BRET when in proximity of NLuc-TTBK1 or NLuc-TTBK2, both tracer candidates produced a robust BRET response for TTBK1 and TTBK2 that could be competed away with introduction of an excess of compound **13** (Figure S2). An optimal signal was generated by tracer candidate **24**, allowing calculation of an EC_50_ of 660 nM for TTBK1 and 710 nM for TTBK2 (Figure S3). The same workflow was used to calculate micromolar EC_50_ values for tracer candidate **23** in both assays (Figure S3). Next, tracer titration experiments were performed in dose–response format with compound **13** at varied concentrations of tracer: 0.125–1 µM. The optimal concentration at which to use our new tracer (Figure S4) was identified as 1.0 µM for TTBK1 and 0.5 µM for TTBK2. Furthermore, we calculated apparent K_D_ values of compound **13** using the linear Cheng–Prusoff relationship: 69 nM and 9 nM for TTBK1 and TTBK2, respectively (Figure S5). Tracer **24** is functionally identical to our first-generation tracer and can be used at lower concentrations in TTBK NanoBRET assays.

**Table 2.**
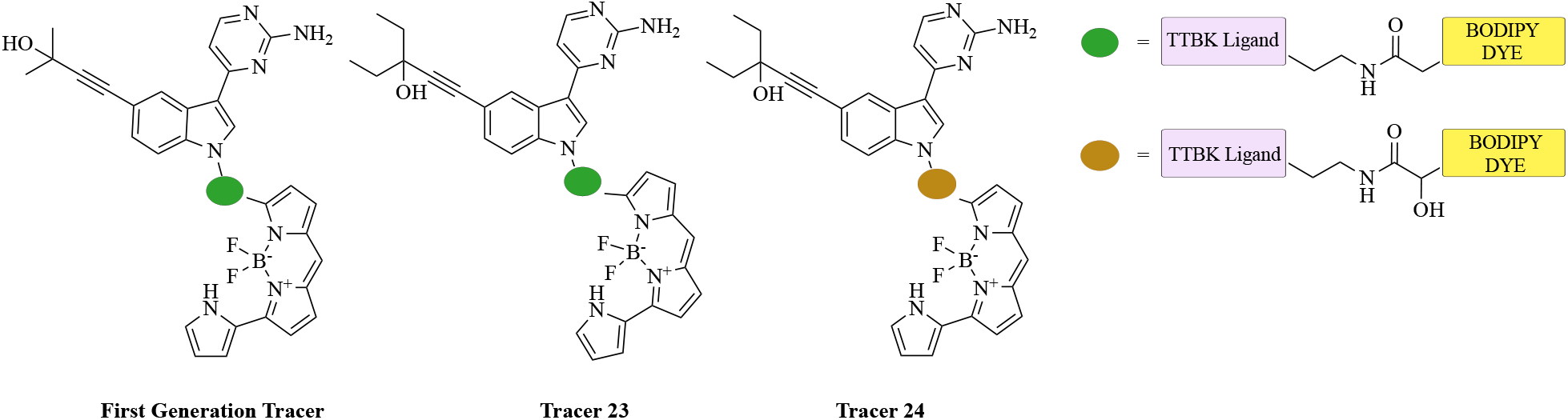
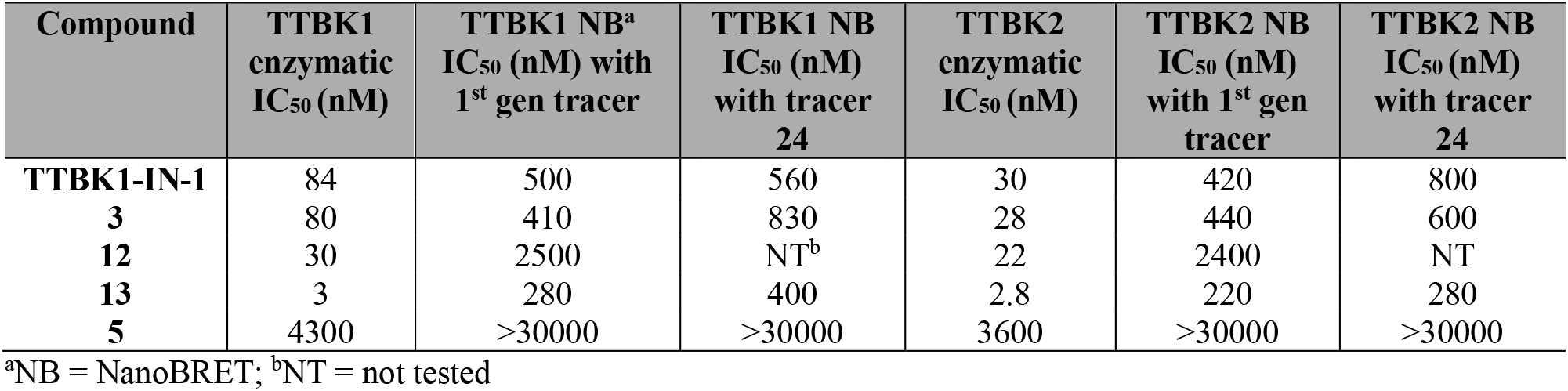
Enzymatic and NanoBRET data for commercial inhibitor TTBK1-IN-1, racemic compound **3**, enantiomers **12** and **13**, and negative control, **5.** NB = NanoBRET.

### (*S*)-Enantiomer of Compound 3 is a Potent, Non-Toxic, and Cell-Active TTBK1/2 Inhibitor

TTBK1-IN-1 is a single (*S*)-enantiomer of compound **14** that demonstrates improved TTBK1/2 NanoBRET affinity. As compound **3** bears a chiral center at its alkyne terminus, we sought to resolve the enantiomers of **3** and evaluate them in the TTBK1/2 NanoBRET assays. Following separation via chiral chromatography and determination of absolute configuration via vibrational circular dichroism (VCD), compound **12** was assigned as the (*R*)-enantiomer and **13** as the (*S*)-enantiomer (Figure S11). With these compounds in hand, **12** and **13** were first assessed for their inhibition of TTBK1 and TTBK2 enzymatic activity. Compound **12** exhibited an IC_50_ value of 30 nM in the TTBK1 enzymatic assay and an IC_50_ value of 22 nM in the TTBK2 enzymatic assay. Compound **13**, in comparison, was 10-fold more active than compound **3** and TTBK1-IN-1, with enzymatic IC_50_ values of 3 nM and 2.8 nM for TTBK1 and TTBK2, respectively (Table 2).

We used both tracer **24** and our first-generation tracer to evaluate **13** alongside a panel of our key compounds in the TTBK1 and TTBK2 NanoBRET assays. When comparing the data generated using our first-generation tracer to that with tracer **24**, we see conserved trends in analog affinity for TTBK1 and TTBK2. There was good agreement between the IC_50_ values observed for TTBK1-IN-1, compound **3**, and compound **13** in the NanoBRET assays (Table 2). Utilizing a lower concentration of tracer can reduce oversaturation of the binding site, providing a more sensitive assay via which we can gauge compound affinity. A 2-fold increase in affinity for both TTBK1 and TTBK2 were observed for the eutomer (**13**, (*S*)-enantiomer) compared to the racemate (**3**). When assayed in the TTBK1 and TTBK2 NanoBRET assays, **12** demonstrated micromolar IC_50_ values, reinforcing it as the distomer. Importantly, compound **13** maintains IC_50_ values of ≤400 nM in all TTBK NanoBRET assays, supporting its potent cellular target engagement of TTBK1 and TTBK2. Finally, compound **13** was found to have no influence on cell viability at concentrations up to 10 µM for 8 hours (Figure S1).

### Compound 13 Exhibits In-Cell Selectivity

Given its promising potency, compound **13** was also analyzed for its in-cell kinome-wide selectivity by Promega in the K300 panel (Figure 4C). Only five of the 300 kinases evaluated demonstrated >50% percent occupancy when compound **13** was profiled at 1 µM (Table S3). These kinases include CDK11B, CDK11A, CK1δ, CK1ε, and EPHA2 (Figure 4C). Unsurprisingly, three of these off-target kinases overlap with those identified for compound **3**. Full dose–response follow-up in corresponding NanoBRET assays illuminated that compound **13** exhibits excellent kinome-wide selectivity, with only four of the five occupied kinases binding with IC_50_ values within 30-fold of the TTBK1 and TTBK2 NanoBRET IC_50_ values (Figure 4D).

To fully appreciate the selectivity profile of TTBK1-IN-1 versus compound **13**, these compounds were screened against 24 kinases with available enzymatic assays. These 24 kinases were selected based on high affinity binding to TTBK1-IN-1 and/or **13** in the published cell-free selectivity profile of TTBK1-IN-1 and the K300 selectivity profile of compound **13** (Figure 4C).^6^ Full dose– response NanoBRET follow-up assays were run, where available and warranted based on enzymatic data (Table S4). Both TTBK1-IN-1 and **13** were found to inhibit many of the screened kinases with IC_50_ values within 30-fold of the TTBK1/2 enzymatic IC_50_ values (Table S4). TTBK1-IN-1 potently inhibited the enzymatic activity of 17 of the 24 kinases. Compound **13** demonstrated an improved cell-free selectivity profile, inhibiting the enzymatic activity of only 13 of the 24 assayed kinases. While **13** is a more potent inhibitor of CDK11B than TTBK1-IN-1, this represents the only increased potency interaction. Conversely, TTBK1-IN-1 is a more potent inhibitor of CK1α1, CK1γ1, JNK1, PRKD2, and TSSK3, and these interactions have been significantly decreased for **13**. NanoBRET assays were available for nine of 13 kinases for compound **13** and only eight of 17 for TTBK1-IN-1, limiting the in-cell selectivity profiling. For both compounds, however, the in-cell selectivity was greatly improved with only three off-target kinases identified for **13** and two kinases for TTBK1-IN-1 binding with an IC_50_ value <1 µM (Table S4).

### Molecular Docking Experiments Substantiate the Enhanced Affinity of Compound 13

We employed molecular docking to better understand the binding of eutomer **13** when compared to distomer **12**, and, additionally, to identify high affinity interactions made by **13**. Utilizing the co-crystal structure of TTBK1 bound to BGN18 (PDB code: 7JXY), we docked **12** and **13** into the TTBK1 active site and evaluated their binding.^6^ Like BGN18, compound **13** can place its propargylic hydroxyl group in a favorable position to act as a hydrogen bond donor to Glu77 and as an acceptor from a backbone nitrogen of Phe177 (Figure 5). The corresponding propargylic hydroxyl group on compound **12**, in contrast, can only act as a hydrogen bond donor to Glu77 but cannot form a second hydrogen bond due to its unfavorable conformation that positions the hydroxyl group far from Phe177. The increased number of back pocket hydrogen bonds made by compound **13** compared to compound **12** correlates with its enhanced enzymatic potency and in-cell affinity (Table 2).

**Figure 5.**
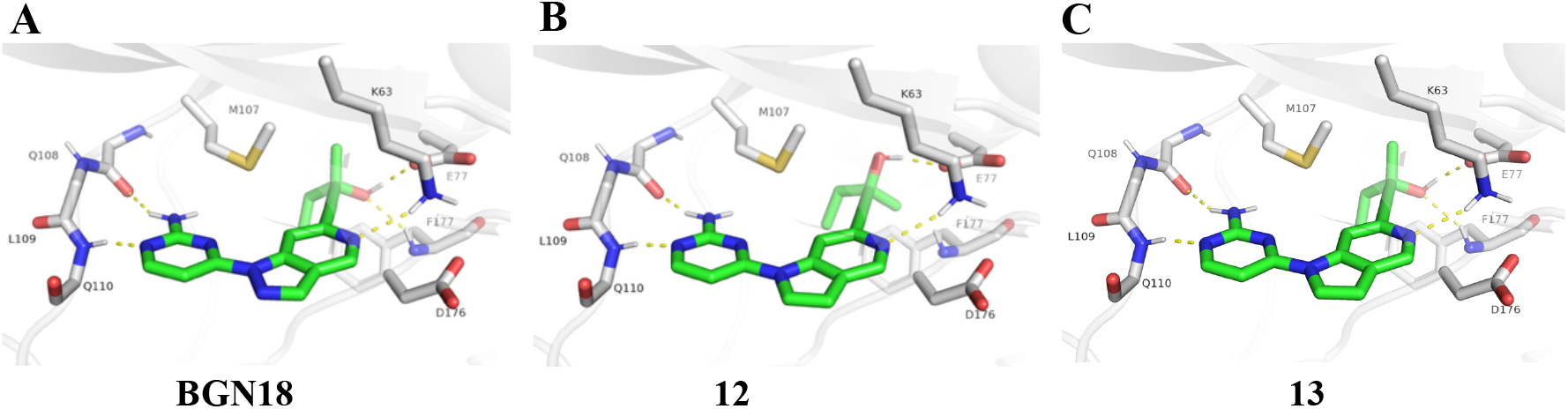
Molecular docking results support preferential binding of (*S*)-enantiomer. (A) Co-crystal structure of BGN18 bound to TTBK1 active site (PDB code: 7JXY). (B) Molecular docking pose of **12** bound to the TTBK1 active site. (C) Molecular docking pose of **13** bound to the TTBK1 active site. Hydrogen bond interactions shown as dotted lines.

### A Negative Control is Identified

A critical component of chemical probe development is the preparation and characterization of a structurally related negative control that lacks on-target affinity. In the co-crystal structure of BGN18 bound to TTBK1 (Figure 2), the propargylic alcohol hydrogen bonds with backbone residues (Glu 77 and Phe 177) in the back pocket. Compound **5** bears a secondary alcohol that extends deeper than the BGN18 propargylic alcohol and creates steric clash with residues that define the small back pocket. This structural modification is proposed to eliminate the hydrogen bonds with Phe 177 and Glu 77. Consistent with these structural hypotheses, compound **5** demonstrated no affinity for TTBK1 or TTBK2 (IC_50_ >30 µM) in the corresponding NanoBRET assays (Table 2). To further confirm that compound **5** lacks TTBK1/2 inhibitory activity, it was profiled in the TTBK1 and TTBK2 radiometric enzyme inhibition assays. Compound **5** was only weakly active in these assays, exhibiting a TTBK1 enzymatic IC_50_ of 4.3 µM and a TTBK2 enzymatic IC_50_ of 3.6 µM. A >1400-fold loss in TTBK1 enzymatic activity and a ∼1300-fold loss in TTBK2 enzymatic activity exist between compounds **13** and **5**. Its complete loss of cellular affinity confirms that compound **5** can serve as a negative control compound and be used in tandem with compound **13** when exploring the consequences of TTBK1/2 inhibition in cells. To confirm its suitability for cellular studies, compound **5** was evaluated for its impact on cell viability at concentrations up to 10 µM for an 8-hour treatment and found to be non-toxic (Figure S1).

### Compounds 3 and 13 Efficiently Inhibit Primary Cilia Formation

To further assess the cellular effects of the TTBK leads, we used hTERT-RPE-1 cells, a widely employed cellular model of ciliogenesis. TTBK2 undergoes a prominent autophosphorylation, which is reflected in its mobility shift when analyzed by Western blot (WB).^8, 19^ Indeed, 4 hour treatment with 10 µM of TTBK1-IN-1 (used as reference compound in these experiments), **3**, and **13**, led to a visible mobility shift of TTBK2 when compared to vehicle-(DMSO) or compound **5**-treated conditions (Figure 6A). These data demonstrate that both compounds **3** and **13** efficiently ablate TTBK2 autophosphorylation. Next, we examined the impact of these compounds on recruitment of IFT88 using immunofluorescence (IF) microscopy. Importantly, when dosed for 4 hours at 10 µM, both compound **3** and **13** caused a prominent reduction in IFT88 levels recruited to one of the two centrioles (Figure 6B-C). Notably, compound **5** showed no difference from the vehicle-treated condition, in agreement with our WB and NanoBRET assay data showing that it lacks binding affinity for and an ability to inhibit TTBK1/2. Finally, we examined these compounds for their cilia-ablating capabilities. To this end, we treated hTERT-RPE-1 cells for 24 hours with 1 μM of the lead compounds and analyzed cilia using IF microscopy. In vehicle- or compound **5**-treated conditions, approximately 70% of cells were ciliated (Figure 6D-E). Remarkably, compounds **3** and **13** caused a significant reduction in the percentage of ciliated cells. Furthermore, the effect of compounds **3** and **13** on cilia was more prominent than that of our reference compound, TTBK1-IN-1. In summary, these findings demonstrate that compounds **3** and **13** effectively inhibit TTBK2 activity and, in turn, cilia formation in a cellular model of ciliogenesis.

**Figure 6.**
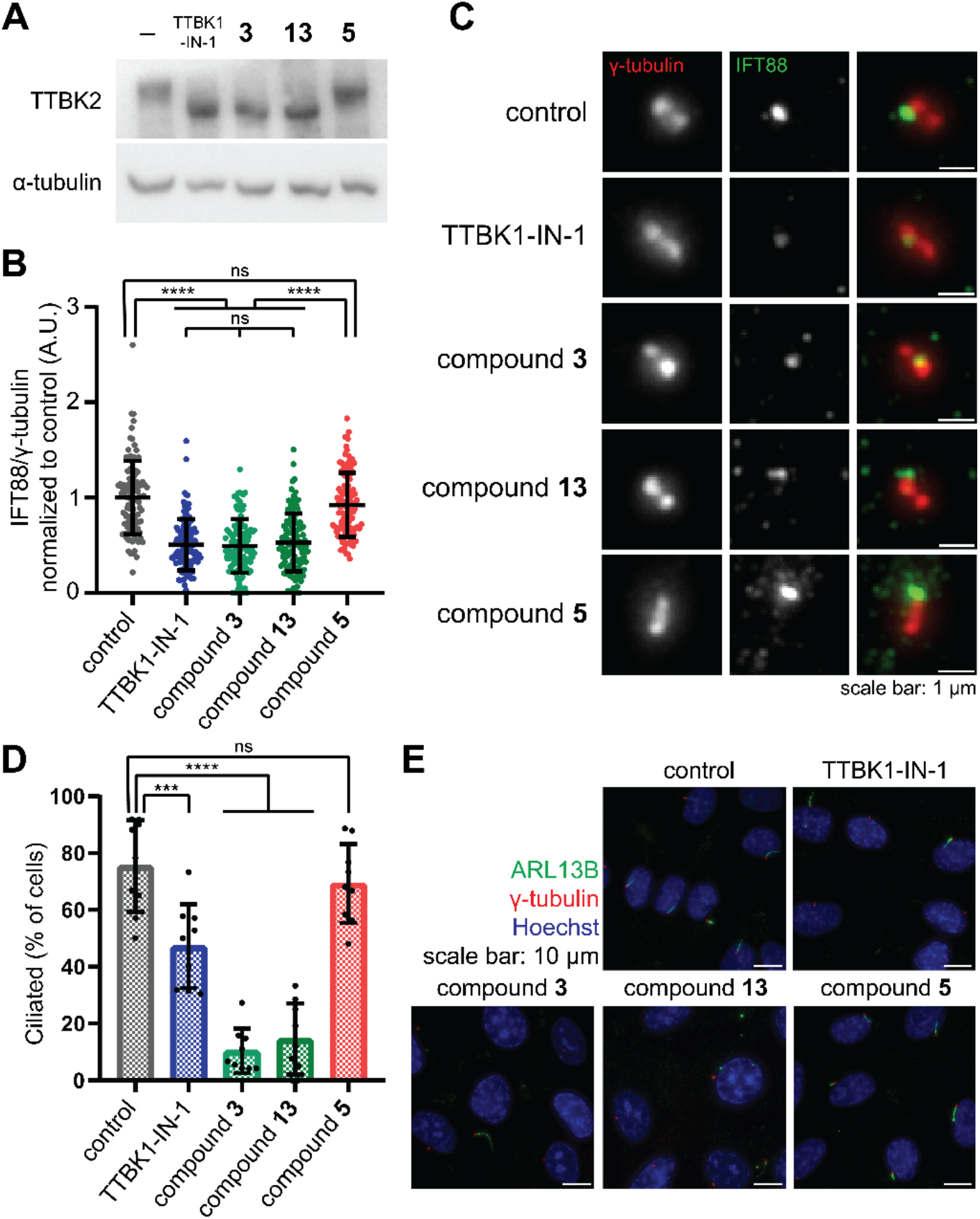
Compounds **3** and **13** successfully block TTBK2 cellular activity, leading to disruption of ciliogenesis. (A) 4-hour treatment of RPE1 cells with 10 µM TTBK1-IN-1, **3**, and **13**, but not **5**, results in ablation of autophosphorylation of TTBK2, reflected in mobility shift on WB. (B, C) IFT88 levels at the mother centriole, ±SD, individual data points shown. Treatment with 10 µM TTBK1-IN-1, **3**, and **13** for 4 hours significantly (p<0.0001) reduces levels of IFT88 residing at the mother centriole compared to both vehicle- and compound **5**-treated cells. γ-tubulin staining was used to detect centrioles (D, E). Percentage of ciliated cells, ±SD, each datapoint shows the average value per image frame. Treatment with 1 µM TTBK1-IN-1 (p=0.0005), **3** (p<0.0001) and **13** (p<0.0001) for 24 hours significantly reduced number of ciliated cells (cilia were detected by ARL13B staining), while compound **5** showed no significant change (p=0.8651).

### Conclusion

As part of our focused SAR campaign, we designed, synthesized, and evaluated 14 novel TTBK1/2 inhibitors. This effort identified compounds **3** and **13** as optimal TTBK inhibitors based on their potent in-cell affinity and enzymatic inhibition of both TTBK isoforms coupled with their selective in-cell kinome profiles and lack of toxicity at the timepoints measured. Additionally, compound **5** was designed as a negative control compound and was confirmed to abolish in-cell TTBK binding affinity and inhibitory activity without associated toxicity. Second generation TTBK1 and TTBK2 NanoBRET assays were developed herein and used to characterize the binding of our lead compounds. Finally, compounds **3** and **13** were found to outperform TTBK1-IN-1 in assays defining the role of TTBK2 in ciliogenesis. Compound **5** functioned as expected and did not differentiate from the vehicle, confirming its utility as a negative control in cell-based phenotypic studies. Our TTBK inhibitors and negative control compounds are available to the scientific community and represent powerful tools to interrogate TTBK1/2-mediated biology as it relates to ciliogenesis, neurodegenerative diseases, and beyond.

## Experimental Section

### Chemistry

#### General Information

All reagents and solvents were used as supplied from commercial suppliers and used without further purification. Temperatures are reported in degrees Celsius (°C), and reactions were run at room temperature (25°C) where no temperature is listed. Solvent was removed via a rotary evaporator under reduced pressure. Column chromatography was performed using preloaded silica gel cartridges or manually loaded cartridges packed with celite on a Biotage automated purification system. The following abbreviations are used in schemes and/or experimental procedures: equiv (equivalent(s)), h (hours), mmol (millimoles), mg (milligrams), min (minutes), µmol (micromoles), and r.t. (room temperature). ^1^H NMR and additional microanalytical data was collected for key intermediates and all final compounds to confirm their identity and assess purity. ^1^H and ^13^C NMR spectra were obtained in CD_3_OD or DMSO-*d*_6_ and recorded using Bruker instruments. The magnet strength of the spectrometer used is included in each line listing. Chemical shifts are reported in parts per million (ppm) and calibrated versus the shift of the deuterated solvent used. Coupling constants (*J* values) are reported in hertz (Hz) and spin multiplicities are listed as follows: singlet (s), doublet (d), doublet of doublets/triplets/quartets (dd/dt/dq), doublet of doublet of doublets (ddd), triplet (t), triplet of doublets/triplets (td/tt), quartet (q), quartet of doublets (qd), pentent (p), and multiplet (m). Preparative HPLC was performed using an Agilent 1260 Infinity II LC System equipped with a Phenomenex Gemini C18 (30°C, 5 μm particle size, 75 x 30 mm). LCMS analyses were executed using an Agilent 1290 Infinity II LC System equipped with an Agilent Infinity Lab PoroShell 120 EC-C18 column (30°C, 2.7 μm particle size, 2.1 × 50 mm), eluent 5−95% ACN in water with 0.2% formic acid (v/v), and flow rate of 1 mL/min. All final compounds were determined to be >95% pure by HPLC analysis.

#### General Procedure A

To a microwave vial, a mixture of heteroaryl halide (1.0 equiv), alkyne (3-5 equiv), K_2_CO_3_ (1.5 equiv), 1,3-bis(diphenylphosphino)propane (0.15 equiv), and Pd(OAc)_2_ (0.1 equiv) in DMF (0.2 M) was added and degassed using argon. The vial was heated in a microwave reactor to 120°C for 1 h. The reaction was cooled, diluted with EtOAc, filtered over celite and concentrated *in vacuo*. The residue was purified by flash chromatography (0–100% EtOAc in hexanes and 0–20% MeOH in EtOAc) and if needed, further purified via preparative HPLC (column: Phenomenex Gemini C18 75 x 30 mm x 5 µm; 30 mL/min; 10–100% ACN in H_2_O + 0.2% NH_4_OH).

#### General Procedure B

To a flask, a mixture of heteroaryl halide (1.0 equiv), alkyne (3.0–5.0 equiv), K_2_CO_3_ (1.5 equiv), 1,3-bis(diphenylphosphino)propane (0.15 equiv), and Pd(OAc)_2_ (0.1 equiv) in DMF (0.2 M) was added and charged with argon. The mixture was heated to 85°C for 16 h. The reaction was cooled, diluted with EtOAc, filtered over celite, and concentrated *in vacuo*. The residue was purified by flash chromatography (0–100% EtOAc in hexanes and 0–20% MeOH in EtOAc) and if needed, further purified by preparative-HPLC (column: Phenomenex Gemini C18 75 x 30 mm x 5 µm; 30 mL/min; 10–100% ACN in H_2_O + 0.2% NH_4_OH 10–100%).

#### General Procedure C

To a flask, a mixture of heteroaryl halide (1.0 equiv), alkyne (3.0–5.0 equiv), DIPA (2.0 equiv), CuI (0.15 equiv), and PdCl_2_(PPh_3_)_2_) (0.1 equiv) in DMF (0.2 M) was added and charged with argon. The mixture was heated to 85°C for 16 h. The reaction was cooled, diluted with EtOAc, filtered over celite, and concentrated *in vacuo*. The crude residue was purified by reverse phase flash chromatography (ACN in H_2_O plus 0.2% NH_4_OH 10–100%) and if needed, further purified by preparative-HPLC (column: Phenomenex Gemini C18 75 x 30 mm x 5 µm; 30 mL/min; 10– 100% ACN in H_2_O + 0.2% NH_4_OH).

#### 1-(1-(2-amino-6-methylpyrimidin-4-yl)-1*H*-pyrrolo[3,2-*c*]pyridin-6-yl)-3-methylpent-1-yn-3-ol (1)

The reaction was prepared according to General Procedure B starting from 4-(6-chloro-1*H*-pyrrolo[3,2-*c*]pyridin-1-yl)-6-methylpyrimidin-2-amine (**26**) (150 mg, 0.58 mmol, 1.0 equiv), Pd(OAc)_2_ (13.0 mg, 0.06 mmol, 0.1 equiv), 1,3-bis(diphenylphosphino)propane (35.7 mg, 0.09 mmol, 0.15 equiv), K_2_CO_3_ (120 mg, 0.87 mmol, 1.5 equiv), 3-methylpent-1-yn-3-ol (262 µL, 2.3 mmol, 4.0 equiv) and DMF (2.89 mL, 0.2 M) were combined under anhydrous conditions and heated to 80°C for 16 h. The reaction was cooled, diluted with EtOAc, filtered over celite, and concentrated. The residue was purified by flash chromatography (0–100% EtOAc in hexanes and 0–20% MeOH in EtOAc) followed by preparative-HPLC (10–100% ACN can in H_2_O (0.2% NH_4_OH)) to afford 1-(1-(2-amino-6-methylpyrimidin-4-yl)-1*H*-pyrrolo[3,2-*c*]pyridin-6-yl)-3-methylpent-1-yn-3-ol as a white solid (9.4 mg, 0.03 mmol, 5.1%). ^1^H NMR (500 MHz, CD_3_OD δ 8.77 (d, *J* = 1.0 Hz, 1H), 8.73 (d, *J* = 1.0 Hz, 1H), 8.05 (d, *J* = 3.7 Hz, 1H), 6.90 (dd, *J* = 3.6, 0.9 Hz, 1H), 6.84 (s, 1H), 2.40 (s, 3H), 1.89 – 1.76 (m, 2H), 1.58 (s, 3H), 1.14 (t, *J* = 7.5 Hz, 3H). ^13^C NMR (126 MHz, MeOD) δ 171.5, 164.7, 160.7, 144.2, 140.7, 136.7, 129.3, 128.2, 106.9, 99.0, 93.3, 83.9, 69.5, 37.6, 29.3, 24.0, 9.4. HPLC Purity 100%. MS (ESI): *m/z* calculated for C_18_H_19_N_5_O for [M+H]^+^: 322.2. Found 322.3.

#### 1-(1-(2-amino-6-methylpyrimidin-4-yl)-1*H*-pyrrolo[3,2-*c*]pyridin-6-yl)-3-ethylpent-1-yn-3-ol (2)

The reaction was prepared according to General Procedure B starting from 4-(6-chloro-1*H*-pyrrolo[3,2-*c*]pyridin-1-yl)-6-methylpyrimidin-2-amine (**26**) (150 mg, 0.58 mmol, 1.0 equiv), Pd(OAc)_2_ (13.0 mg, 0.06 mmol, 0.1 equiv), 1,3-bis(diphenylphosphino)propane (35.7 mg, 0.09 mmol, 0.15 equiv), K_2_CO_3_ (120 mg, 0.87 mmol, 1.5 equiv), 3-ethylpent-1-yn-3-ol (262 µL, 2.3 mmol, 4.0 equiv) and DMF (2.89 mL, 0.2 M) were combined under anhydrous conditions and heated to 80°C for 16 h. The reaction was cooled, diluted with EtOAc, filtered over celite, and concentrated. The residue was purified by flash chromatography (0–100% EtOAc in hexanes and 0–20% MeOH in EtOAc) followed by preparative-HPLC (10–100% ACN in H_2_O (0.2% NH_4_OH)) to afford 1-(1-(2-amino-6-methylpyrimidin-4-yl)-1*H*-pyrrolo[3,2-*c*]pyridin-6-yl)-3-ethylpent-1-yn-3-ol as a white solid (7.5 mg, 0.02 mmol, 4%). ^1^H NMR (500 MHz, MeOD) δ 8.77 (d, *J* = 1.0 Hz, 1H), 8.72 (d, *J* = 0.9 Hz, 1H), 8.04 (d, *J* = 3.7 Hz, 1H), 6.90 (dd, *J* = 3.7, 0.9 Hz, 1H), 6.83 (s, 1H), 2.40 (s, 3H), 1.88 – 1.74 (m, 4H), 1.13 (t, *J* = 7.4 Hz, 6H). ^13^C NMR (126 MHz, MeOD) δ 171.5, 164.7, 160.7, 144.2, 140.7, 136.7, 129.3, 128.1, 115.7, 106.9, 99.1, 92.5, 85.0, 73.1, 35.2, 24.0, 9.0. HPLC Purity 100%. MS (ESI): *m/z* calculated for C_19_H_21_N_5_O for [M+H]^+^: 336.2. Found 336.5.

#### 1-(1-(2-aminopyrimidin-4-yl)-1*H*-pyrrolo[3,2-*c*]pyridin-6-yl)-3-methylpent-1-yn-3-ol (3)

The reaction was prepared according to General Procedure B starting from 4-(6-chloro-1*H*-pyrrolo[3,2-*c*]pyridin-1-yl)pyrimidin-2-amine (**17**) (200 mg, 0.8 mmol, 1.0 equiv), Pd(OAc)_2_ (18.3 mg, 0.08 mmol, 0.1 equiv), 1,3-bis(diphenylphosphino)propane (50.4 mg, 0.12 mmol, 0.15 equiv), K_2_CO_3_ (169 mg, 1.22 mmol, 1.5 equiv), 3-methylpent-1-yn-3-ol (369 µL, 3.26 mmol, 4.0 equiv) and DMF (4.07 mL, 0.2 M) were combined under anhydrous conditions and heated to 85°C for 16 h. The reaction was cooled, diluted with EtOAc, filtered over celite, and concentrated. The residue was purified by flash chromatography (0–100% EtOAc in hexanes and 0–20% MeOH in EtOAc) to afford 1-(1-(2-aminopyrimidin-4-yl)-1*H*-pyrrolo[3,2-*c*]pyridin-6-yl)-3-methylpent-1-yn-3-ol as a white solid (150 mg, 0.49 mmol, 60%). ^1^H NMR (500 MHz, DMSO-*d*_6_) δ 8.85 (d, *J* = 1.1 Hz, 1H), 8.61 (t, *J* = 1.0 Hz, 1H), 8.34 (d, *J* = 5.5 Hz, 1H), 8.19 (d, *J* = 3.6 Hz, 1H), 7.06 (s, 2H), 6.99 (d, *J* = 5.6 Hz, 1H), 6.93 (dd, *J* = 3.6, 0.9 Hz, 1H), 5.40 (s, 1H), 1.76 – 1.64 (m, 2H), 1.47 (s, 3H), 1.04 (t, J = 7.4 Hz, 3H). ^13^C NMR (126 MHz, DMSO-*d*_6_) δ 163.4, 160.4, 158.3, 143.7, 138.2, 135.7, 128.1, 126.1, 113.8, 105.6, 98.3, 92.5, 83.0, 67.1, 36.3, 29.2, 9.1. HPLC Purity 100%. MS (ESI): *m/z* calculated for C_17_H_17_N_5_O for [M+H]^+^: 308.2. Found 308.4.

#### 1-(1-(2-aminopyrimidin-4-yl)-1*H*-pyrrolo[3,2-c]pyridin-6-yl)-3-ethylpent-1-yn-3-ol (4)

The reaction was prepared according to General Procedure A starting from 4-(6-chloro-1*H*-pyrrolo[3,2-*c*]pyridin-1-yl)pyrimidin-2-amine (**17**) (150 mg, 0.61 mmol, 1.0 equiv), Pd(AcO)_2_ (13.7 mg, 0.06 mmol, 0.1 equiv), 1,3-bis(diphenylphosphino)propane (37.8 mg, 0.09 mmol, 0.15 equiv), K_2_CO_3_ (127 mg, 0.92 mmol, 1.5 equiv), 3-Ethylpent-1-yn-3-ol (314 µL, 274 mg, 2.4 mmol, 4.0 equiv) in DMF (3.05 mL, 0.2 M) were combined under anhydrous conditions and heated to 120°C in a microwave reactor for 1 h. The reaction was cooled, diluted with EtOAc, filtered over celite, and concentrated. The residue was purified by flash chromatography (0–100% EtOAc in hexanes and 0–20% MeOH in EtOAc) followed by preparative-HPLC (10–100% ACN in H_2_O (0.2% NH_4_OH)) to afford 1-(1-(2-aminopyrimidin-4-yl)-1*H*-pyrrolo[3,2-*c*]pyridin-6-yl)-3-ethylpent-1-yn-3-ol as a white solid (20 mg, 0.06 mmol, 10%). ^1^H NMR (500 MHz, MeOD) δ 8.82 – 8.70 (m, 2H), 8.31 (d, *J* = 5.7 Hz, 1H), 8.06 (d, *J* = 3.7 Hz, 1H), 6.96 – 6.89 (m, 2H), 1.88 – 1.73 (m, 4H), 1.13 (t, *J* = 7.4 Hz, 6H). ^13^C NMR (126 MHz, MeOD) δ 164.9, 161.0, 160.5, 144.3, 140.7, 136.9, 129.2, 128.2, 115.8, 107.2, 99.8, 92.6, 85.0, 73.1, 35.2, 9.0. HPLC Purity 100%. MS (ESI): *m/z* calculated for C_18_H_19_N_5_O for [M+H]^+^: 322.2. Found 322.0.

#### 4-(1-(2-aminopyrimidin-4-yl)-1*H*-pyrrolo[3,2-*c*]pyridin-6-yl)-2,2-dimethylbut-3-yn-1-ol (5)

The reaction was prepared according to General Procedure A starting from 4-(6-chloro-1*H*-pyrrolo[3,2-*c*]pyridin-1-yl)pyrimidin-2-amine (**17**) (200 mg, 0.8 mmol, 1.0 equiv), Pd(OAc)_2_ (18.3 mg, 0.08 mmol, 0.1 equiv), 1,3-bis(diphenylphosphino)propane (50.4 mg, 0.12 mmol, 0.15 equiv), K_2_CO_3_ (169 mg, 1.22 mmol, 1.5 equiv), 2,2-dimethylbut-3-yn-1-ol (300 µL, 240 mg, 2.44 mmol, 3.0 equiv) and DMF (4.07 mL, 0.2 M) were combined under anhydrous conditions and heated to 120°C in a microwave reactor for 1 h. The reaction was cooled, diluted with EtOAc, filtered over celite, and concentrated. The residue was purified by flash chromatography (0–100% EtOAc in hexanes and 0–20% MeOH in EtOAc) followed by preparative-HPLC (10–100% ACN in H_2_O (0.2% NH_4_OH)) to afford 4-(1-(2-aminopyrimidin-4-yl)-1*H*-pyrrolo[3,2-*c*]pyridin-6-yl)-2,2-dimethylbut-3-yn-1-ol as white solid (12 mg, 0.04 mmol, 4.8%). ^1^H NMR (500 MHz, MeOD) δ 8.76 (m, 2H), 8.30 (d, *J* = 5.8 Hz, 1H), 8.05 (d, *J* = 3.7 Hz, 1H), 6.99 – 6.85 (m, 2H), 3.55 (s, 2H), 1.33 (s, 6H). ^13^C NMR (126 MHz, MeOD) δ 164.9, 160.9, 160.5, 144.1, 140.8, 137.5, 129.0, 128.0, 115.5, 107.2, 99.6, 95.8, 82.2, 71.9, 35.4, 25.7. HPLC Purity 100%. MS (ESI): *m/z* calculated for C_17_H_17_N_5_O for [M+H]^+^: 308.2. Found 308.4.

#### 1-(1-(2-aminopyrimidin-4-yl)-1*H*-pyrrolo[3,2-*c*]pyridin-6-yl)-3,4,4-trimethylpent-1-yn-3-ol (6)

The reaction was prepared according to General Procedure A starting from 4-(6-chloro-1*H*-pyrrolo[3,2-*c*]pyridin-1-yl)pyrimidin-2-amine (**17**) (200 mg, 0.81 mmol, 1.0 equiv), Pd(AcO)_2_(18.3 mg, 0.08 mmol, 0.1 equiv), 1,3-bis(diphenylphosphino)propane (50.4 mg, 0.12 mmol, 0.15 equiv), K_2_CO_3_ (169 mg, 1.2 mmol, 1.5 equiv), 3,4,4-trimethylpent-1-yn-3-ol (355 µL, 308 mg, 2.4 mmol, 3.0 equiv) in DMF (4.07 mL, 0.2 M) were combined under anhydrous conditions and heated to 120°C in a microwave reactor for 1 h. The reaction was cooled, diluted with EtOAc, filtered over celite, and concentrated. The residue was purified by flash chromatography (0–100% EtOAc in hexanes and 0–20% MeOH in EtOAc) followed by preparative-HPLC (10–100% ACN in H_2_O (0.2% NH_4_OH)) to afford 1-(1-(2-aminopyrimidin-4-yl)-1*H*-pyrrolo[3,2-*c*]pyridin-6-yl)-3,4,4-trimethylpent-1-yn-3-ol as a white solid (2.8 mg, 0.008 mmol, 1.0%). ^1^H NMR (500 MHz, MeOD) δ 8.79 (d, *J* = 1.0 Hz, 1H), 8.74 (t, *J* = 1.0 Hz, 1H), 8.31 (d, *J* = 5.7 Hz, 1H), 8.06 (d, *J* = 3.7 Hz, 1H), 6.96 – 6.91 (m, 2H), 1.57 (s, 3H), 1.15 (s, 9H). ^13^C NMR (214 MHz, MeOD) δ 161.0, 160.5, 144.3, 137.0, 129.2, 115.6, 107.2, 99.9, 57.5, 39.5, 25.9, 25.2, 17.4, 17.3, 17.2, 17.1. HPLC Purity 100%. MS (ESI): *m/z* calculated for C_19_H_21_N_5_O for [M+H]^+^: 336.2. Found 336.1.

#### 1-(1-(2-aminopyrimidin-4-yl)-1*H*-pyrrolo[3,2-*c*]pyridin-6-yl)-3,5-dimethylhex-1-yn-3-ol (7)

The reaction was prepared according to General Procedure A starting from 4-(6-chloro-1*H*-pyrrolo[3,2-*c*]pyridin-1-yl)pyrimidin-2-amine (**17**) (150 mg, 0.61 mmol, 1.0 equiv), Pd(AcO)_2_ (13.7 mg, 0.06 mmol, 0.1 equiv), 1,3-bis(diphenylphosphino)propane (37.8 mg, 0.09 mmol, 0.15 equiv), K_2_CO_3_ (127 mg, 0.92 mmol, 1.5 equiv), 3,5-dimethylhex-1-yn-3-ol (262 µL, 231 mg, 1.8 mmol, 3.0 equiv), in DMF (3.05 mL, 0.2 M) were combined under anhydrous conditions and heated to 120°C in a microwave reactor for 1 h. The reaction was cooled, diluted with EtOAc, filtered over celite, and concentrated. The residue was purified by flash chromatography (0–100% EtOAc in hexanes and 0–20% MeOH in EtOAc) followed by preparative-HPLC (10–100% ACN in H_2_O (0.2% NH_4_OH)) to afford 1-(1-(2-aminopyrimidin-4-yl)-1*H*-pyrrolo[3,2-*c*]pyridin-6-yl)-3,5-dimethylhex-1-yn-3-ol as a white solid (3.8 mg, 0.01 mmol, 1.9%). ^1^H NMR (500 MHz, MeOD) δ 8.80 – 8.72 (m, 2H), 8.31 (d, *J* = 5.7 Hz, 1H), 8.06 (d, *J* = 3.7 Hz, 1H), 6.95 – 6.90 (m, 2H), 2.08 (dp, *J* = 13.0, 6.5 Hz, 1H), 1.72 (dd, *J* = 6.0, 2.5 Hz, 2H), 1.60 (s, 3H), 1.08 (dd, *J* = 6.7, 1.5 Hz, 6H). ^13^C NMR (126 MHz, MeOD) δ 164.9, 161.0, 160.5, 144.3, 140.7, 136.9, 129.2, 128.2, 115.7, 107.2, 99.8, 94.1, 84.0, 68.9, 53.1, 30.9, 26.2, 24.8. HPLC Purity 100%. MS (ESI): *m/z* calculated for C_19_H_21_N_5_O for [M+H]^+^: 336.2. Found 336.5

#### 1-((1-(2-aminopyrimidin-4-yl)-1*H*-pyrrolo[3,2-*c*]pyridin-6-yl)ethynyl)cyclohexan-1-ol (8)

The reaction was prepared according to General Procedure A starting from 4-(6-chloro-1*H*-pyrrolo[3,2-*c*]pyridin-1-yl)pyrimidin-2-amine (**17**) (200 mg, 0.81 mmol, 1.0 equiv), Pd(AcO)_2_ (18.3 mg, 0.08 mmol, 0.1 equiv), 1,3-bis(diphenylphosphino)propane (50.4 mg, 0.12 mmol, 0.15 equiv), K_2_CO_3_ (169 mg, 1.2 mmol, 1.5 equiv), 1-ethynylcyclohexan-1-ol (416 µL, 404 mg, 3.3 mmol, 4.0 equiv) in DMF (4.07 mL, 0.2 M) were combined under anhydrous conditions and heated to 120°C in a microwave reactor for 1 h. The reaction was cooled, diluted with EtOAc, filtered over celite, and concentrated. The residue was purified by flash chromatography (0–100% EtOAc in hexanes and 0–20% MeOH in EtOAc) followed by preparative-HPLC (10–100% ACN in H_2_O (0.2% NH_4_OH)) to afford 1-((1-(2-aminopyrimidin-4-yl)-1*H*-pyrrolo[3,2-*c*]pyridin-6-yl)ethynyl)cyclohexan-1-ol as a white solid (5.7 mg, 0.02 mmol, 2.1%). ^1^H NMR (500 MHz, MeOD) δ 8.77 (dt, *J* = 15.1, 1.1 Hz, 2H), 8.30 (d, *J* = 5.7 Hz, 1H), 8.05 (dd, *J* = 3.7, 1.8 Hz, 1H), 6.92 (td, *J* = 5.0, 1.2 Hz, 2H), 2.10 – 2.00 (m, 2H), 1.80 – 1.66 (m, 6H), 1.60 (ddd, *J* = 9.8, 6.7, 3.2 Hz, 1H), 1.41 – 1.30 (m, 1H). ^13^C NMR (126 MHz, MeOD) δ 164.9, 160.9, 160.5, 144.3, 140.6, 136.9, 129.2, 128.2, 115.8, 107.2, 99.8, 93.6, 84.7, 69.4, 40.7, 26.4, 24.2. HPLC Purity 100%. MS (ESI): *m/z* calculated for C_19_H_19_N_5_O for [M+H]^+^: 334.2. Found 334.4.

#### 1-((1-(2-aminopyrimidin-4-yl)-1*H*-pyrrolo[3,2-*c*]pyridin-6-yl)ethynyl)cyclopentan-1-ol (9)

The reaction was prepared according to General Procedure A starting from 4-(6-chloro-1*H*-pyrrolo[3,2-*c*]pyridin-1-yl)pyrimidin-2-amine (**17**) (200 mg, 0.81 mmol, 1.0 equiv), Pd(AcO)_2_ (18.3 mg, 0.08 mmol, 0.1 equiv), 1,3-bis(diphenylphosphino)propane (50.4 mg, 0.12 mmol, 0.15 equiv), K_2_CO_3_ (169 mg, 1.2 mmol, 1.5 equiv), 1-ethynylcyclopentan-1-ol (280 µL, 269 mg, 2.4 mmol, 3.0 equiv) in DMF (4.07 mL, 0.2 M) were combined under anhydrous conditions and heated to 120°C in a microwave reactor for 1 h. The reaction was cooled, diluted with EtOAc, filtered over celite, and concentrated. The residue was purified by flash chromatography (0–100% EtOAc in hexanes and 0–20% MeOH in EtOAc) followed by preparative-HPLC (10–100% ACN in H_2_O (0.2% NH_4_OH)) to afford 1-((1-(2-aminopyrimidin-4-yl)-1*H*-pyrrolo[3,2-*c*]pyridin-6-yl)ethynyl)cyclopentan-1-ol as a white solid (3.8 mg, 0.012 mmol, 1.5%). ^1^H NMR (500 MHz, MeOD) δ 8.78 (dd, *J* = 2.1, 1.0 Hz, 2H), 8.31 (d, *J* = 5.7 Hz, 1H), 8.07 (d, *J* = 3.7 Hz, 1H), 6.97 – 6.88 (m, 2H), 2.19 – 2.00 (m, 4H), 1.97 – 1.77 (m, 4H). ^13^C NMR (126 MHz, MeOD) δ 164.9, 160.9, 160.5, 144.3, 140.7, 137.0, 129.2, 128.2, 115.8, 107.2, 99.7, 93.6, 83.5, 75.3, 43.1, 24.4. HPLC Purity 100%. MS (ESI): *m/z* calculated for C_18_H_17_N_5_O for [M+H]^+^: 320.2. Found 320.0.

#### 1-(3-(2-aminopyrimidin-4-yl)-1*H*-indol-5-yl)-3-methylpent-1-yn-3-ol (10)

The reaction was prepared according to General Procedure B starting from 4-(5-bromo-1*H*-indol-3-yl)pyrimidin-2-amine (**18**) (100 mg, 0.3 mmol, 1.0 equiv), PdCl_2_(PPh_3_)_2_ (24.3 mg, 0.03 mmol, 0.1 equiv), CuI (6.59 mg, 0.03 mmol, 0.1 equiv) 3-methylpent-1-yn-3-ol (118 µL, 102 mg, 1.0 mmol, 3.0 equiv), DIPA (98 µL, 70 mg, 0.7 mmol, 2.0 equiv), in propanol (1.73 mL, 0.2 M) was charged with argon and heated to 85°C for 16 h. The reaction was cooled, diluted with EtOAc, filtered over celite, and concentrated. The residue was purified by flash chromatography (0–100% EtOAc in hexanes and 0–20% MeOH in EtOAc) followed by preparative-HPLC (10–100% ACN in H_2_O (0.2% NH_4_OH)) to afford 1-(3-(2-aminopyrimidin-4-yl)-1*H*-indol-5-yl)-3-methylpent-1-yn-3-ol as a white solid (5.2 mg, 0.02 mmol, 4.8%). The analytical data for compound **10** matches that previously published.^31^ ^1^H NMR (500 MHz, MeOD) δ 8.54 (dd, *J* = 1.5, 0.7 Hz, 1H), 8.09 (d, *J* = 24.2 Hz, 1H), 8.06 (s, 1H), 7.39 (dd, *J* = 8.4, 0.7 Hz, 1H), 7.25 (dd, *J* = 8.4, 1.5 Hz, 1H), 7.04 (d, *J* = 5.4 Hz, 1H), 1.85 – 1.73 (m, 2H), 1.55 (s, 3H), 1.14 (t, *J* = 7.4 Hz, 3H). ^13^C NMR (126 MHz, MeOD) δ 165.3, 164.7, 157.5, 138.4, 130.1, 127.1, 126.6, 126.5, 116.7, 115.5, 112.9, 107.5, 91.6, 85.6, 69.7, 37.8, 29.7, 9.6. HPLC Purity 100%. MS (ESI): *m/z* calculated for C_18_H_18_N_4_O for [M+H]^+^: 307.1. Found 307.4

#### 1-(3-(2-aminopyrimidin-4-yl)-1*H*-indol-5-yl)-3-ethylpent-1-yn-3-ol (11)

The reaction was prepared according to General Procedure B starting from 4-(5-bromo-1*H*-indol-3-yl)pyrimidin-2-amine (**18**) (200 mg, 0.7 mmol, 1.0 equiv), PdCl_2_(PPh_3_)_2_ (48.6 mg, 0.07 mmol, 0.1 equiv), CuI (13.2 mg, 0.07 mmol, 0.1 equiv), 3-ethylpent-1-yn-3-ol (267 µL, 233 mg, 2.1 mmol, 3.0 equiv), DIPA (195 µL, 140 mg, 1.4 mmol, 2.0 equiv), in propanol (3.46 mL, 0.2 M) was charged with argon and heated to 85°C for 16 h. The reaction was cooled, diluted with EtOAc, filtered over celite, and concentrated. The residue was purified by flash chromatography (0–100% EtOAc in hexanes and 0–20% MeOH in EtOAc) followed by preparative-HPLC (10–100% ACN in H_2_O (0.2% NH_4_OH)) to afford 1-(3-(2-aminopyrimidin-4-yl)-1*H*-indol-5-yl)-3-ethylpent-1-yn-3-ol as a white solid (18.9 mg, 0.06 mmol, 8.5%). The analytical data for compound **11** matches that previously published.^31^ ^1^H NMR (500 MHz, MeOD) δ 8.52 (dd, *J* = 1.6, 0.7 Hz, 1H), 8.10 (d, *J* = 5.5 Hz, 1H), 8.05 (s, 1H), 7.39 (dd, *J* = 8.4, 0.7 Hz, 1H), 7.26 (dd, *J* = 8.4, 1.5 Hz, 1H), 7.03 (d, *J* = 5.5 Hz, 1H), 1.77 (m, 4H), 1.13 (t, *J* = 7.4 Hz, 6H). ^13^C NMR (214 MHz, MeOD) δ 165.2, 164.7, 157.6, 138.4, 130.1, 127.1, 126.6, 126.4, 116.8, 115.5, 112.9, 107.6, 90.6, 86.7, 73.3, 35.6, 9.1. HPLC Purity 100%. MS (ESI): *m/z* calculated for C_19_H_21_N_4_O for [M+H]^+^: 321.2. Found 321.1.

#### (*R*)-1-(1-(2-aminopyrimidin-4-yl)-1*H*-pyrrolo[3,2-*c*]pyridin-6-yl)-3-methylpent-1-yn-3-ol (12)

The reaction was prepared according General Procedure B starting from 4-(6-chloro-1*H*-pyrrolo[3,2-*c*]pyridin-1-yl)pyrimidin-2-amine (**17**) (250 mg, 1.02 mmol, 1.0 equiv), Pd(OAc)_2_ (22.8 mg, 0.102 mmol, 0.1 equiv), 1,3-bis(diphenylphosphino)propane (63.0 mg, 0.153 mmol, 0.15 equiv), K_2_CO_3_ (211 mg, 1.53 mmol, 1.5 equiv), 3-methylpent-1-yn-3-ol (461 µL, 4.07 mmol, 4.0 equiv) and DMF (5.09 mL, 0.2 M) were combined under anhydrous conditions and heated to 85°C for 16 h. The reaction was cooled, diluted with EtOAc, filtered over celite, and concentrated. The residue was purified by flash chromatography (0–100% EtOAc in hexanes and 0–20% MeOH in EtOAc) to afford racemic 1-(1-(2-aminopyrimidin-4-yl)-1*H*-pyrrolo[3,2-*c*]pyridin-6-yl)-3-methylpent-1-yn-3-ol which was chirally resolved via chiral chromatography to afford (*R*)-1-(1-(2-aminopyrimidin-4-yl)-1*H*-pyrrolo[3,2-*c*]pyridin-6-yl)-3-methylpent-1-yn-3-ol as a white solid in 99.9% ee (82 mg, 0.27 mmol, 26%).^1^H NMR (500 MHz, MeOD) δ 8.80 – 8.74 (m, 2H), 8.31 (d, *J* = 5.7 Hz, 1H), 8.06 (d, *J* = 3.7 Hz, 1H), 7.00 – 6.89 (m, 2H), 1.86 – 1.79 (m, 2H), 1.58 (s, 3H), 1.14 (t, *J* = 7.4 Hz, 3H). ^13^C NMR (126 MHz, MeOD) δ 164.9, 160.9, 160.5, 144.3, 140.7, 136.8, 129.2, 128.2, 115.8, 107.2, 99.8, 93.4, 83.9, 69.5, 37.5, 29.3, 9.4. HPLC Purity 100%. MS (ESI): *m/z* calculated for C_17_H_17_N_5_O for [M+H]^+^: 308.2. Found 308.0.

#### *(S*)-1-(1-(2-aminopyrimidin-4-yl)-1*H*-pyrrolo[3,2-*c*]pyridin-6-yl)-3-methylpent-1-yn-3-ol (13)

The reaction was prepared according General Procedure B starting from 4-(6-chloro-1*H*-pyrrolo[3,2-*c*]pyridin-1-yl)pyrimidin-2-amine (**17**) (250 mg, 1.02 mmol, 1.0 equiv), Pd(OAc)_2_ (22.8 mg, 0.102 mmol, 0.1 equiv), 1,3-bis(diphenylphosphino)propane (63.0 mg, 0.153 mmol, 0.15 equiv), K_2_CO_3_ (211 mg, 1.53 mmol, 1.5 equiv), 3-methylpent-1-yn-3-ol (461 µL, 4.07 mmol, 4.0 equiv) and DMF (5.09 mL, 0.2 M) were combined under anhydrous conditions and heated to 85°C for 16 h. The reaction was cooled, diluted with EtOAc, filtered over celite, and concentrated. The residue was purified by flash chromatography (0–100% EtOAc in hexanes and 0–20% MeOH in EtOAc) to afford racemic 1-(1-(2-aminopyrimidin-4-yl)-1*H*-pyrrolo[3,2-*c*]pyridin-6-yl)-3-methylpent-1-yn-3-ol which was chirally resolved via chiral chromatography to afford (*S*)-1-(1-(2-aminopyrimidin-4-yl)-1*H*-pyrrolo[3,2-*c*]pyridin-6-yl)-3-methylpent-1-yn-3-ol as a white solid in 99.5% ee (88 mg, 0.29 mmol, 28%).^1^H NMR (500 MHz, MeOD) δ 8.78 (d, *J* = 9.4 Hz, 2H), 8.31 (d, *J* = 5.7 Hz, 1H), 8.07 (d, *J* = 3.7 Hz, 1H), 6.98 – 6.88 (m, 2H), 1.87 – 1.78 (m, 2H), 1.58 (s, 3H), 1.14 (t, *J* = 7.5 Hz, 3H). ^13^C NMR (126 MHz, MeOD) δ 164.9, 160.9, 160.5, 144.3, 140.7, 136.8, 129.2, 128.2, 115.8, 107.2, 99.8, 93.4, 83.9, 69.5, 37.5, 29.3, 9.4. HPLC Purity 100%. MS (ESI): *m/z* calculated for C_17_H_17_N_5_O for [M+H]^+^: 308.2. Found 308.1.

#### 1-(1-(2-amino-6-methoxypyrimidin-4-yl)-1*H*-pyrrolo[3,2-*c*]pyridin-6-yl)-3-methylpent-1-yn-3-ol (14)

The reaction was prepared according to General Procedure B starting from 4-(6-chloro-1*H*-pyrrolo[3,2-*c*]pyridin-1-yl)pyrimidin-2-amine (**17**) (120 mg, 0.435 mmol, 1.0 equiv), Pd(OAc)_2_ (9.77 mg, 0.043 mmol, 0.1 equiv), 1,3-bis(diphenylphosphino)propane (26.9 mg, 0.065 mmol, 0.15 equiv), K_2_CO_3_ (90.2 mg, 0.653 mmol, 1.5 equiv), 3-methylpent-1-yn-3-ol (197 µL, 1.74 mmol, 4.0 equiv) and DMF (2.18 mL, 0.2 M) were combined under anhydrous conditions and heated to 85°C for 16 h. The reaction was cooled, diluted with EtOAc, filtered over celite, and concentrated. The residue was purified by flash chromatography (0–100% EtOAc in hexanes and 0–20% MeOH in EtOAc) followed by preparative-HPLC (10–100% ACN in H_2_O (0.2% NH_4_OH)) to afford 1-(1-(2-amino-6-methoxypyrimidin-4-yl)-1*H*-pyrrolo[3,2-*c*]pyridin-6-yl)-3-methylpent-1-yn-3-ol as a white solid (5.4 mg, 0.016 mmol, 3.7%). ^1^H NMR (500 MHz, MeOD) δ 8.75 (d, *J* = 1.1 Hz, 1H), 8.62 (t, *J* = 0.9 Hz, 1H), 7.97 (d, *J* = 3.6 Hz, 1H), 6.85 (dd, *J* = 3.5, 0.8 Hz, 1H), 6.25 (s, 1H), 3.94 (s, 3H), 1.82 (m, 2H), 1.57 (s, 3H), 1.14 (t, *J* = 7.4 Hz, 3H). ^13^C NMR (126 MHz, MeOD) δ 173.9, 165.0, 161.1, 144.2, 140.6, 136.3, 129.5, 128.0, 115.4, 106.4, 93.2, 85.1, 83.9, 69.5, 54.3, 37.5, 29.3, 9.4. HPLC Purity 98%. MS (ESI): *m/z* calculated for C_18_H_19_N_5_O_2_ for [M+H]^+^: 338.2. Found 338.4.

#### 4-(6-chloro-1*H*-pyrrolo[3,2-*c*]pyridin-1-yl)pyrimidin-2-amine (17)

The reaction was prepared starting from 6-chloro-1*H*-pyrrolo[3,2-*c*]pyridine (**16**) (1.50 g, 9.83 mmol, 1.0 equiv) in DMF (65.5 mL, 0.15 M) on ice was added NaH (590 mg, 14.7 mmol, 60% in mineral oil, 1.5 equiv) and allowed to stir for 30 min and warm to r.t. after which 4-chloropyrimidin-2-amine (**15**) (1.91 g, 14.7 mmol, 1.5 equiv) was added. The reaction was allowed to stir for 16 h and then quenched with water (25 mL). The precipitate was filtered and the filter cake washed with water (25 mL) and allowed to dry to afford 4-(6-chloro-1*H*-pyrrolo[3,2-*c*]pyridin-1-yl)pyrimidin-2-amine as an off-white solid (2.17 g, 8.83 mmol, 90%). ^1^H NMR (500 MHz, DMSO-*d*_6_) δ 8.76 (d, *J* = 0.9 Hz, 1H), 8.72 (d, *J* = 18.9 Hz, 1H), 8.33 (d, *J* = 5.6 Hz, 1H), 8.21 (d, *J* = 3.7 Hz, 1H), 7.11 (s, 2H), 7.00 (d, *J* = 5.6 Hz, 1H), 6.94 (dd, *J* = 3.7, 0.9 Hz, 1H). ^13^C NMR (126 MHz, DMSO*-d*_6_) δ 163.3, 160.4, 158.2, 144.1, 142.6, 140.3, 128.2, 126.6, 110.9, 105.4, 97.6. HPLC Purity 98.5%. MS (ESI): *m/z* calculated for C_11_H_8_ClN_5_ for [M+H]^+^: 246.1. Found 246.0.

#### 4-(6-chloro-1*H*-pyrrolo[3,2-*c*]pyridin-1-yl)-6-methoxypyrimidin-2-amine (20)

The reaction was prepared starting from 6-chloro-1*H*-pyrrolo[3,2-*c*]pyridine (**16**) (1.50 g, 9.83 mmol, 1.0 equiv) in DMF (65.5 mL, 0.15 M) on ice was added NaH (590 mg, 14.7 mmol, 60% in mineral oil, 1.5 equiv) and allowed to stir for 1 h and warm to r.t. after which 4-chloro-6-methoxypyrimidin-2-amine (**19**) (2.4 g, 15 mmol, 1.5 equiv) was added. The reaction was stirred at 90°C for 12 h, cooled to r.t. and quenched with water (20 mL) and extracted with EtOAc (15 mL x 3). The organic layers were combined and washed with brine (15 mL), dried over sodium sulfate and concentrated *in vacuo*. The crude residue was purified by flash chromatography (0-15% MeOH in DCM) to afford 4-(6-chloro-1*H*-pyrrolo[3,2-*c*]pyridin-1-yl)-6-methoxypyrimidin-2-amine as a white solid (1.85 g, 6.71 mmol, 68%). ^1^H NMR (500 MHz, DMSO*-d*_6_) δ 8.77 – 8.68 (m, 2H), 8.19 (d, *J* = 3.7 Hz, 1H), 6.91 (dd, *J* = 3.7, 0.8 Hz, 1H), 6.44 (s, 1H), 3.88 (s, 3H). Exchangeable protons not observed. ^13^C NMR (126 MHz, DMSO-*d*_6_) δ 171.8, 163.0, 159.4, 143.9, 142.5, 140.4, 128.4, 126.5, 110.8, 105.1, 82.6, 53.5. HPLC Purity 96.8%. MS (ESI): *m/z* calculated for C_12_H_10_ClN_5_O for [M+H]^+^: 276.1. Found 275.9.

#### 4-(1-(3-aminopropyl)-5-bromo-1*H*-indol-3-yl)pyrimidin-2-amine (21)

The reaction was prepared starting from 4-(5-bromo-1*H*-indol-3-yl)pyrimidin-2-amine (**18**) (600 mg, 2.08 mmol, 1.0 equiv) in DMF (20.8 mL, 0.1 M) on ice to which sodium hydride was added (166 mg, 4.15 mmol, 60% in mineral oil, 2.0 equiv) and stirred for 30 min and allowed to warm to r.t. after which 3-((*tert*-butoxycarbonyl)amino)propyl 4-methylbenzenesulfonate (1.37 g, 4.15 mmol, 2.0 equiv) was added and stirred at r.t. for 16 h. The reaction was diluted with water (100 mL) and extracted with EtOAc (3 x 30 mL) and concentrated *in vacuo*. The residue was taken up in DCM (10.4 mL, 0.2 M) and TFA (3.55 g, 2.38 mL, 31.1 mmol. 15 equiv) was added and allowed to stir for 12 h. The reaction was concentrated *in vacuo* and purified by reverse phase flash chromatography (10–100% ACN in H_2_O (0.2% NH_4_OH)) to afford 4-(1-(3-aminopropyl)-5-bromo-1*H*-indol-3-yl)pyrimidin-2-amine as a yellow solid (160 mg, 0.462 mmol, 22.3% over two steps). ^1^H NMR (500 MHz, MeOD) δ 8.72 (d, *J* = 1.9 Hz, 1H), 8.17 – 8.06 (m, 2H), 7.46 (d, *J* = 8.7 Hz, 1H), 7.38 (dd, *J* = 8.7, 2.0 Hz, 1H), 6.99 (d, *J* = 5.5 Hz, 1H), 4.36 (t, *J* = 7.0 Hz, 2H), 2.91 – 2.81 (m, 2H), 2.18 (dt, *J* = 16.3, 7.1 Hz, 2H). ^13^C NMR (126 MHz, MeOD) δ 164.7, 164.6, 157.6, 137.3, 133.1, 129.2, 126.7, 126.2, 115.8, 114.7, 112.7, 107.2, 44.9, 38.5, 30.1. HPLC Purity 100%. MS (ESI): *m/z* calculated for C_15_H_16_BrN_5_ for [M+H]^+^: 346.1. Found 348.0.

#### 1-(1-(3-aminopropyl)-3-(2-aminopyrimidin-4-yl)-1*H*-indol-5-yl)-3-ethylpent-1-yn-3-ol (22)

The reaction was prepared according to General Procedure C starting from 4-(1-(3-aminopropyl)-5-bromo-1*H*-indol-3-yl)pyrimidin-2-amine (**21**) (50 mg, 0.14 mmol, 1.0 equiv), 3-ethylpent-1-yn-3-ol (93 µL, 0.72 mmol, 5.0 equiv), DIPA (41 µL, 0.29 mmol, 2.0 equiv), PdCl_2_(PPh_3_)_2_) (10 mg, 0.014 mmol, 0.1 equiv) and CuI (4.1 mg, 0.022 mmol, 0.15 equiv) in DMF (0.72 mL, 0.2 M) was charged with argon. The mixture was heated to 85°C for 16 h. The reaction was cooled, diluted with EtOAc, filtered over celite and concentrated *in vacuo*. The crude residue was purified by reverse phase flash chromatography (10–100% ACN in H_2_O (0.2% NH_4_OH)) to afford 1-(1-(3-aminopropyl)-3-(2-aminopyrimidin-4-yl)-1*H*-indol-5-yl)-3-ethylpent-1-yn-3-ol as a white solid (18 mg, 0.048 mmol, 33%).^1^H NMR (500 MHz, MeOD) δ 8.56 (d, *J* = 1.5 Hz, 1H), 8.10 (d, *J* = 10.0 Hz, 2H), 7.48 (d, *J* = 8.5 Hz, 1H), 7.31 (dd, *J* = 8.5, 1.6 Hz, 1H), 7.01 (d, *J* = 5.5 Hz, 1H), 4.31 (q, *J* = 8.3 Hz, 2H), 2.70 (m, 2H), 2.05 (p, *J* = 7.2 Hz, 2H), 1.86 – 1.66 (m, 4H), 1.13 (t, *J* = 7.4 Hz, 6H). ^13^C NMR (126 MHz, MeOD) δ 164.8, 164.7, 157.6, 138.1, 133.1, 127.3, 126.8, 117.1, 115.0, 111.3, 107.5, 91.0, 86.4, 73.3, 35.5, 30.7, 22.6, 9.2. HPLC 98%. MS (ESI): *m/z* calculated for C_22_H_27_N_5_O for [M+H]^+^: 378.2. Found 378.5.

#### *N*-(3-(3-(2-aminopyrimidin-4-yl)-5-(3-ethyl-3-hydroxypent-1-yn-1-yl)-1*H*-indol-1-yl)propyl)-3-(5,5-difluoro-7-(1*H*-pyrrol-2-yl)-5*H*-5λ^4^,6λ^4^-dipyrrolo[1,2-*c*:2’,1’-*f*][1,3,2]diazaborinin-3-yl)propenamide (23)

The reaction was prepared starting from NanoBRET590 SE dye (10 mg, 0.023 mmol, 1.0 equiv) and 1-(1-(3-aminopropyl)-3-(2-aminopyrimidin-4-yl)-1*H*-indol-5-yl)-3-ethylpent-1-yn-3-ol (**22**) (9.7 mg, 0.026 mmol, 1.1 equiv) in DMF (0.12 mL, 0.2 M) was added DIPEA (0.02 mL, 0.12 mmol, 5.0 equiv) and allowed to stir at r.t. for 30 min. The reaction was dried *in vacuo* and the residue purified by reverse phase flash chromatography (10–100% ACN in H_2_O (0.2% NH_4_OH)) to afford *N*-(3-(3-(2-aminopyrimidin-4-yl)-5-(3-ethyl-3-hydroxypent-1-yn-1-yl)-1*H*-indol-1-yl)propyl)-3-(5,5-difluoro-7-(1*H*-pyrrol-2-yl)-5*H*-5λ^4^,6λ^4^-dipyrrolo[1,2-*c*:2’,1’-*f*][1,3,2]diazaborinin-3-yl)propenamide as a purple solid (1.2 mg, 1.7 µmol, 7.4%). ^1^H NMR (400 MHz, MeOD) δ 8.55 (dd, *J* = 1.6, 0.7 Hz, 1H), 8.06 – 7.97 (m, 2H), 7.42 – 7.33 (m, 1H), 7.26 (dd, *J* = 8.5, 1.6 Hz, 1H), 7.23 – 7.16 (m, 2H), 7.16 – 7.11 (m, 2H), 6.99 (d, *J* = 4.6 Hz, 1H), 6.96 (d, *J* = 5.5 Hz, 1H), 6.87 (d, *J* = 4.0 Hz, 1H), 6.40 – 6.30 (m, 2H), 4.15 (t, *J* = 7.0 Hz, 2H), 3.33 (s, 2H), 3.23 (t, *J* = 6.4 Hz, 2H), 2.66 (t, *J* = 7.5 Hz, 2H), 2.04 (p, *J* = 6.7 Hz, 2H), 1.90 – 1.63 (m, 4H), 1.12 (t, *J* = 7.4 Hz, 6H). HPLC Purity 100%. MS (ESI): *m/z* calculated for C_38_H_39_BF_2_N_8_O_2_ for [M+H]^+^: 689.3. Found 689.8.

#### *N*-(3-(3-(2-aminopyrimidin-4-yl)-5-(3-ethyl-3-hydroxypent-1-yn-1-yl)-1*H*-indol-1-yl)propyl)-3-(5,5-difluoro-7-(1*H*-pyrrol-2-yl)-5*H*-5λ^4^,6λ^4^-dipyrrolo[1,2-*c*:2’,1’-*f*][1,3,2]diazaborinin-3-yl)-2-hydroxypropanamide (24)

The reaction was prepared starting from NanoBRET590 SE dye (10 mg, 0.023 mmol, 1.0 equiv) and 1-(1-(3-aminopropyl)-3-(2-aminopyrimidin-4-yl)-1*H*-indol-5-yl)-3-ethylpent-1-yn-3-ol (**22**) (9.7 mg, 0.026 mmol, 1.1 equiv) in DMF (0.12 mL, 0.2 M) was added DIPEA (0.02 mL, 0.12 mmol, 5.0 equiv) and allowed to stir at r.t. for 30 min. The reaction was dried *in vacuo* and the residue purified by reverse phase flash chromatography (10–100% ACN in H_2_O (0.2% NH_4_OH)) to afford *N*-(3-(3-(2-aminopyrimidin-4-yl)-5-(3-ethyl-3-hydroxypent-1-yn-1-yl)-1*H*-indol-1-yl)propyl)-3-(5,5-difluoro-7-(1*H*-pyrrol-2-yl)-5*H*-5λ^4^,6λ^4^-dipyrrolo[1,2-*c*:2’,1’-*f*][1,3,2]diazaborinin-3-yl)-2-hydroxypropanamide as a purple solid (4.2 mg, 23 µmol, 25%). ^1^H NMR (400 MHz, MeOD) δ 8.55 (dd, *J* = 1.6, 0.7 Hz, 1H), 8.10 – 7.97 (m, 2H), 7.42 (dd, *J* = 8.6, 0.7 Hz, 1H), 7.27 (dd, *J* = 8.5, 1.6 Hz, 1H), 7.25 – 7.19 (m, 2H), 7.19 – 7.14 (m, 2H), 7.03 (d, *J* = 4.7 Hz, 1H), 6.98 (d, *J* = 5.5 Hz, 1H), 6.90 (d, *J* = 4.0 Hz, 1H), 6.60 (d, *J* = 4.0 Hz, 1H), 6.35 (dd, *J* = 3.9, 2.6 Hz, 1H), 5.72 (dd, *J* = 9.2, 4.1 Hz, 1H), 4.29 – 4.11 (m, 2H), 3.29 – 3.18 (m, 2H), 2.85 (dd, *J* = 14.1, 4.2 Hz, 1H), 2.65 (dd, *J* = 14.1, 9.1 Hz, 1H), 2.06 (p, *J* = 6.9 Hz, 2H), 1.86 – 1.66 (m, 4H), 1.12 (t, *J* = 7.4 Hz, 6H). HPLC Purity 100%. MS (ESI): *m/z* calculated for C_38_H_39_BF_2_N_8_O_3_ for [M+H]^+^: 705.3. Found 705.7.

#### 4-(6-chloro-1*H*-pyrrolo[3,2-*c*]pyridin-1-yl)-6-methylpyrimidin-2-amine (26)

The reaction was prepared starting from 6-chloro-1*H*-pyrrolo[3,2-*c*]pyridine (**16**) (2.0 g, 13 mmol, 1.0 equiv) in DMF (87 mL, 0.15 M) on ice was added NaH (790 mg, 20 mmol, 60% in mineral oil, 1.5 equiv) and allowed to stir for 30 min and warm to r.t. after which 4-chloro-6-methylpyrimidin-2-amine (**25**) (2.8 g, 20 mmol, 1.5 equiv) was added. The reaction was heated to 60°C and allowed to stir for 16 h and then quenched with water (50 mL) and extracted with EtOAC (3 x 30 mL). The combined organics were dried over sodium sulfate, filtered, and concentrated *in vacuo* to afford 4-(6-chloro-1*H*-pyrrolo[3,2-*c*]pyridin-1-yl)-6-methylpyrimidin-2-amine as a white solid (2.2 g, 8.5 mmol, 65%). ^1^H NMR (500 MHz, DMSO*-d*_6_) δ 8.76 – 8.68 (m, 1H), 6.99 (d, *J* = 8.1 Hz, 2H), 6.93 (d, *J* = 2.7 Hz, 1H), 5.83 (s, 2H), 5.77 (s, 1H), 2.06 (s, 3H). HPLC Purity 95%. MS (ESI): *m/z* calculated for C_12_H_10_ClN_5_ for [M+H]^+^: 260.1. Found 260.4.

### Biological Evaluation

#### NanoBRET Assays

Human embryonic kidney (HEK293) cells obtained from ATCC were cultured in Dulbecco’s Modified Eagle’s medium (DMEM, Gibco) supplemented with 10% (v/v) fetal bovine serum (FBS, Corning). These cells were incubated in 5% CO_2_ at 37°C and passaged every 72 h with trypsin (Gibco). Cells were not allowed to reach confluency.

Constructs for NanoBRET measurements of TTBK1 (NLuc-TTBK1), TTBK2 (NLuc-TTBK2), MAPK6 (NLuc-MAPK6), CDKL5 (NLuc-CDKL5), MET (MET-NLuc), PAK4 (PAK4-NLuc), PAK5 (PAK7-NLuc), PAK6 (PAK6-NLuc), PIP4K2C (PIP4K2C-NLuc), PIKFYVE (PIKFYVE-NLuc), CDK11B/Cyclin K (CDK11B-NLuc), CDK11A/Cyclin K (CDK11A-NLuc), CK1δ (NLuc-CSNK1D), CK1ε (NLuc-CSNK1E), MAP3K19 (MAP3K19-NLuc) MAP4K5 (MAP4K5-NLuc), and, EPHA2 (EPHA2-NLuc) included in Tables 1–2 and Figure 3-4 were kindly provided by Promega. The NLuc orientations (N- or C-terminally linked) used in the respective assays are indicated in parentheses after each construct. NanoBRET assays were run in dose–response format (11-pt curves) as described previously to generate IC_50_ values.^32^ mBRET units were calculated by dividing the acceptor emission (600 nm) values by the donor emission values (450 nm). Assays were carried out as described by the manufacturer using 0.5 µM tracer K10 for PAK4 and PAK6, 2.0 µM tracer K5 for PAK5, 0.063 µM tracer K8 for PIP4K2C, 0.13 µM tracer K8 for CK1δ and PIKFYVE, 0.5 µM tracer K8 for CK1ε, 0.13 µM tracer K12 for CDK11A/Cyclin K and CDK11B/Cyclin K, 0.031 µM tracer K4 for EPHA2, 0.0078 μM tracer K10 for MAPK6, 0.13 µM tracer K10 for MAP3K19, 1.0 µM tracer K10 for MAP4K5, and 0.25 μM tracer K10 for CDKL5 and MET. Representative curves generated for TTBK1 and TTBK2 are included in Figure S6. NanoBRET curves generated to further refine the in-cell selectivity of compound **3** versus MAPK6, CDKL5, MET, PIP4K2C, CDK11A/Cyclin K, CDK11B/Cyclin K, and CK1δ are found in Figure S7. Representative curves generated to refine the in-cell selectivity of compound **13** versus CDK11A/Cyclin K, CDK11B/Cyclin K, CK1δ, CK1ε, PIP4K2C, PIKFYVE, MAP4K5, PAK4, PAK5, PAK6, MAP3K19, and EPHA2 versus are included in Figure S8.

### In-cell Selectivity Profiling

#### K192 Selectivity Panel

The NanoBRET Target Engagement K192 Kinase Selectivity assay was performed as previously published with modifications detailed below.^33, 34, 16, 35^ Kinases were grouped according to their required K10 tracer concentration. A top control well (tracer plus DMSO) defined the maximum BRET/0% fractional occupancy, and a sample well (tracer plus compound) was prepared for each kinase. Bottom control wells (expressing NanoLuc control vector pNL1.1.CMV[Nluc/CMV]) defined zero BRET/100% fractional occupancy. Reagents were provided by Promega (Promega #NP4060).

#### K300 Selectivity Panel

The NanoBRET Target Engagement K300 Kinase Selectivity assay was performed at Promega.^36^

#### Enzyme Assays

Several Reaction Biology Corp. (RBC) radiometric HotSpot kinase assays were carried out at the K_m_ value for ATP in dose–response (10-pt curve), specifically for TTBK1 and TTBK2. The corresponding IC_50_ values are included in Table 2. Details related to the substrate used, protein constructs, controls, and assay protocol for these HotSpot kinase assays can be accessed via the RBC website: https://www.reactionbiology.com/list-kinase-targets-us-facility.

#### Kinetic Solubility

Analiza, Inc. analyzed the kinetic solubility of a 10 mM DMSO stock solution of compound **3** dissolved in a neutral phosphate buffered saline (PBS) solution (pH 7.4) as previously described.^17^ The reported solubility value has been corrected for background nitrogen present in the media as well as for the presence of DMSO.

#### PAMPA

Analiza, Inc. analyzed the permeability of a 10mM DMSO stock solution of compound **3** dissolved in neutral phosphate buffered saline (PBS) solution (pH 7.4) as previously described.^17^ CLND was used to analyze and quantify permeability.

#### Cell Titer-Glo Cell Viability Assay

Cell Titer-Glo viability assays were performed as previously described.^18^ Briefly, HEK293 cells cultured in DMEM (Gibco) with 10% FBS or SH-SY5Y cells cultured in DMEM/F12 (Gibco) with 10% FBS or at 37 °C in 5% CO_2_, were plated at 12,000 cells/well in a 96-well plate (Corning) and incubated overnight (37 °C, 5% CO_2_). The DMSO percentage was kept consistent across treatment conditions. Compound **3, 13, 5**, or TTBK1-IN-1 were added to wells at 0.5 µM, 1.0 µM, or 10 µM in quadruplicate and treated cells were incubated for 8 hours. Luminescence was read on a GloMax Discover luminometer (Promega). GraphPad Prism was used to analyze the dose– response data.

#### EC_50_ Determination

HEK293 cells were transfected with NL-TTBK1 or NL-TTBK2. **23** or **24** was tested in an 11 or 10-point dose-response format with a top concentration of 10 μM. N=3. Data was plotted using GraphPad Prism Software and fit using a Sigmoidal three-parameter dose response logistical curve to determine the EC_50_. Error bars indicate the standard deviation.

#### NanoBRET Tracer Titration

Tracer titration experiments were performed as described previously.^3, 52^ Briefly, HEK293 cells were transfected with NLuc-TTBK1 or NLuc-TTBK2. Compound **13** was tested in 11-point dose– response format with a top concentration of 30 µM. Tracer **24** stocks were prepared in tracer dilution buffer containing 20% DMSO, for final concentrations of 0.125, 0.25, 0.5, and 1.0 µM.

#### Cell Culture and Treatment

hTERT-RPE1 cell line was cultured in DMEM/F-12 media supplemented with 10% FBS, 2 mM L-Glutamine and 100 IU/mL Penicillin-Streptomycin. For IFT88 recruitment analysis and TTBK2 autophosphorylation assessment, cells were cultivated for 48 hours prior to treatment, then treated with 10 µM concentration of the tested compounds for 4 hours. TTBK1-IN-1 was obtained from MedChemExpress (HY-134968). For ciliogenesis assay, cells were cultivated without treatment for 48 hours, then serum starved simultaneously with the addition of tested compounds at a concentration of 1 µM for 24 hours.

#### Immunofluorescence Microscopy

Cells were fixed in MeOH for 10 min at -20°C, washed with PBS, and blocked in a blocking buffer (2% BSA in PBS). Primary and secondary antibodies (listed below) were applied in the blocking buffer for 1 hour at room temperature, with 10 min PBS washing steps in between. Nuclei were stained with Hoechst for 10 min at room temperature. Following a final washing step with dH_2_O, coverslips were mounted using ProLong™ Glass Antifade Mountant (Thermo Fisher Scientific). Imaging was performed using Zeiss AxioImager.Z2 equipped with a Hamamatsu ORCA Flash 4.0 camera. Z-stack images were acquired using 100x/1.4 Plan-Apochromat oil immersion objective, controlled by Zen Blue software.

Antibodies used for immunofluorescence microscopy: rabbit anti-IFT88 (13967-1-AP; Proteintech; diluted 1:500), mouse anti-γ-tubulin (T6557; Sigma; diluted 1:1000), rabbit anti-ARL13B (17711-1-AP; Proteintech; diluted 1:1000), donkey anti-mouse IgG Alexa 568-linked (A10037; Thermo Fisher Scientific; diluted 1:1000), donkey anti-rabbit IgG Alexa 488-linked (A21206; Thermo Fisher Scientific, diluted 1:1000).

#### Image Analysis

For analysis, maximum projections of Z-stack images created in ImageJ/Fiji software were used.^37^ IFT88 levels were measured in ImageJ/Fiji software from a region of 15x15 pixel size around the γ-tubulin marked centrosome chosen manually, with background subtracted from an area of the same size at a different location within the cell. IFT88 signal intensity was normalized to γ-tubulin signal intensity. Cilia were counted and measured manually in ImageJ/Fiji software using the polyline tool. Each presented quantification represents a summary of three independent experiments.

#### Western Blotting

Cells were lysed in SDS lysis buffer (50 mM Tris-HCl pH 6.8, 10% glycerol, 2% SDS, 0.01% bromphenol blue, 5% β-mercaptoethanol). Lysates were denatured for 10 min at 95°C, loaded onto 8% SDS-polyacrylamide gel, electrophoresed and blotted on polyvinylidene difluoride membrane (Merck Millipore). Membrane was blocked in 5% nonfat milk in PBS-Tween buffer (10 mM Tris-HCl pH 7.4, 100 mM NaCl, 0.05% Tween), incubated with primary antibodies diluted in the blocking buffer overnight at 4°C, rinsed in PBS-Tween buffer and incubated with secondary antibodies diluted in the blocking buffer for 2 hours at room temperature. Detection was performed using Clarity Max™ Western ECL Substrate and Chemidoc Touch Imaging System (both from Bio-Rad).

Antibodies used for WB: rabbit anti-TTBK2 (HPA018113; Sigma; diluted 1:500), rabbit anti-α-tubulin (11224-1-AP; Proteintech; diluted 1:2000), goat anti-rabbit IgG HRP-linked (7074S; Cell signaling, diluted 1:500-4000).

#### Chiral Separation of Compound 3

Chiral separation of **3** was performed at Chiral Technologies. The HPLC-grade solvents used were all purchased from Scientific Equipment Company (SECO) and were used as received from the manufacturer. Specifically, the hexane (hex) used contained 95% *n*-hexane. Screening and optimization of the HPLC mode was performed on an Agilent 1200 configured with a high-pressure mixing quaternary mobile phase delivery system, vacuum degasser, autosampler, temperature-controlled column compartment, and photodiode array UV detector. The instrument was controlled by an Agilent ChemStation Version RevB.04.03. The chiral columns used for screening included CHIRALPAK® IA, IB N-5, IC, ID, IE, IF, IG, IH, IJ, IK, IM, and IN, and were 4.6 mm inner diameter (i.d.) by 250 mm in length and had a 5 µm particle size.

200 mg of compound **3**, was sent to Chiral Technologies for resolution by chiral chromatography. A retention check was performed prior to the start of the screening to ensure reasonable elution was achieved, before committing to a full screen. A single injection was made on the IB N-5 column with both Hex-EtOAc-DEA and Hex-IPA-DEA = 70-30-0.1 (v/v/v). It was observed that this produced reasonable retention and was thus used for screening.

Several partial separations were observed with both EtOH and IPA mobile phases on all columns. IM produced the most optimal resolution of the two enantiomers using a Hex-IPA mobile phase and without a need for further optimization (Figure S9). To determine the productivity under preparative conditions, 1.51 milligrams of material was dissolved in the mobile phase and a series of increasing injection volumes made (Figure S9) until the leading edge and trailing edge of the two peaks corresponding with the enantiomer met. Upon establishing the sample loading (150 µl @ 1.51 mg/ml) and the cycle time (6.5 mins), the hourly productivity was calculated at 2.09 mg on a 4.6x250 mm analytical column. This translates to 84 mg/hour on an equivalent length 30 cm i.d. preparative column.

Using this method, we were able to separate 200 milligrams of racemate, yielding 82 and 88 milligrams of enantiomer peak 1 and 2, respectively, with corresponding chiral purities of 99.9% and 99.5% (Figure S10). Peak 1 was arbitrarily assigned as compound **12** and peak 2 was arbitrarily assigned to compound **13**.

#### Absolute Configuration Assignment

Both compounds **12** and **13** were sent to BioTools to elucidate the absolute configuration using vibrational circular dichroism (VCD).^20, 21, 22, 23, 24, 25^ Quantitative comparisons of the experimental and density functional theory IR (DFT IR) and VCD spectra resulted in high similarity (Sfg) and ESI (enantiomeric similarity index) values and a high confidence level assignment (CompareVOA, BioTools, Jupiter, FL).^26, 27^ Compound **12** was assigned as the (*R*) enantiomer and **13** as the (*S*) enantiomer (Figure S11).

#### VCD Measurements

The CDCl_3_ solvent (Cambridge Isotope Labs (CIL) Silver Foil) was run through a plug of activated basic alumina immediately prior to use. CD_3_CN (CIL) was used as is without further purification.

To a small vial containing 7 mg of an enantiomer (**12** or **13**) was added 485 µL of 8% CD_3_CN in CDCl_3_. A 125 µL aliquot was then transferred to a liquid IR cell (BaF_2_, 100 µm cell path) and placed in the measurement chamber. Experimental spectra were collected on a BioTools, Inc. (Jupiter, FL) Chiral*IR*-*2X*™ Dual *PEM* FT-VCD spectrometer set to 4 cm^-1^ resolution with PEM (both 1 and 2) maximum frequency set to 1400 cm^-1^. The sample was then measured for 18 hours in one-hour blocks. The IR data from the first block was solvent and water vapor subtracted, then offset to zero at 2000 cm^-1^. The VCD data blocks were averaged, enantiomer corrected ((E1 – E2) / 2) and offset to zero at 2000 cm^-1^. The VCD noise data was block averaged and used without further processing.

#### VCD Calculation

The (*R*) enantiomer (chosen arbitrarily) of **3** was constructed using ComputeVOA (BioTools, Jupiter, FL). A thorough conformational search was performed at the MM (molecular mechanics) level using the MMF94 force field in a 7 kcal/mol energy window. All conformers found were subjected to density functional theory (DFT) level optimization and frequency calculation with Gaussian ‘09 (Wallingford, CT) using the B3LYP / 6-31G(d) and B3PW91 / 6-31G(d) methods with implicit solvation (CPCM, Chloroform).^28^ The resulting lowest energy unique conformations were then reoptimized using the B3LYP/cc-pVTZ and B3PW91/cc-pVTZ methods (also with CPCM/chloroform), and the IR and VCD frequencies recalculated at this level. The resulting spectra from all methods were (individually) Boltzmann averaged (using both free energy and electronic energy), plotted at 5 cm^-1^ resolution, and then x-axis scaled (range of 0.968 to 0.987 - values obtained using CompareVOA (BioTools, Jupiter, FL) and varied with basis set and functional) for comparison to the experimental IR and VCD spectra. The two different weighing methods (free energy and electronic energy) gave consistent results for stereochemistry, with the electronic energy weighting performing slightly better in reproducing the experimental data in each case. Of the four different DFT levels screened, the combination of B3PW91/cc-pVTZ gave the highest similarity values and closest appearance to the experimental data, so this was used in all comparisons and plots.

#### Molecular Docking

Protein Preparation Workflow^29^ in Schrödinger Maestro was utilized to prepare the TTBK1 structure (PDB code: 7JXY, chain A). The docking grid file was generated subsequently via Receptor Grid Generation. 3D ligands were produced using the LigPrep tool, then core constrained docking was executed based on maximum common substructure in Glide Ligand Docking^30, 31^ with the standard precision mode. The native ligand in the PDB structure was selected as a template and, for rejected poses, docking was attempted once again using a 1 Å tolerance. The final docking conformations were visualized using PyMol.

## Associated Content

### Supporting Information

The Supporting Information is available free of charge at {filler}.

Molecular Formula Strings file (CSV), NanoBRET curves, Tracer Titration, EC_50_ determination, Cheng-Prusoff relationship, cellular toxicity, in-cell selectivity % occupancy data, VCD calculations, chiral separation experimental, purity chromatograms, and NMR spectra are included (PDF).

## Author Information

**Raymond Flax** - *Structural Genomics Consortium, UNC Eshelman School of Pharmacy, University of North Carolina at Chapel Hill, Chapel Hill, North Carolina 27599, United States*;

**Andrea Lacigová** - *Laboratory of Cilia and Centrosome Biology, Department of Histology and Embryology, Faculty of Medicine, Masaryk University, Kamenice 3, Brno, 62500, Czech Republic;*

**Stefanie Howell** - *Structural Genomics Consortium, UNC Eshelman School of Pharmacy, University of North Carolina at Chapel Hill, Chapel Hill, North Carolina 27599, United States*;

**Frances M. Bashore** - *Structural Genomics Consortium, UNC Eshelman School of Pharmacy, University of North Carolina at Chapel Hill, Chapel Hill, North Carolina 27599, United States*;

**Haoxi Li** - *Laboratory for Molecular Modeling UNC Eshelman School of Pharmacy, University of North Carolina at Chapel Hill, Chapel Hill, NC 27599, USA*;

**Lukáš Čajánek** - *Laboratory of Cilia and Centrosome Biology, Department of Histology and Embryology, Faculty of Medicine, Masaryk University, Kamenice 3, Brno, 62500, Czech Republic;*

## Author Contributions

All authors have given approval to the final version of the manuscript.

## Notes

The authors declare no competing financial interest.

## Acknowledgment

Constructs for NanoBRET measurements of MAPK6, CDKL5, MET, PIP4K2C, PIKFYVE, CDK11A/Cyclin K, CDK11B/Cyclin K, CK1δ, CK1ε, EPHA2, MAP4K5, PAK4, PAK5, PAK6, MAP3K19, TTBK1 and TTBK2, were kindly provided by Promega. To expand kinome-wide profiling, Promega executed the K300 NanoBRET assay panel on compounds **3** and **13**. The TREEspot kinase mapping software from DiscoverX was used to prepare the kinome trees in Figures 3 and 4 (http://treespot.discoverx.com). The table of contents graphic was created with BioRender.com. Research reported in this publication was supported by the Office Of The Director, NIH under award number S10OD032476.

The SGC is a registered charity (number 1097737) that receives funds from Bayer AG, Boehringer Ingelheim, the Canada Foundation for Innovation, Eshelman Institute for Innovation, Genentech, Genome Canada through Ontario Genomics Institute, EU/EFPIA/OICR/McGill/KTH/Diamond, Innovative Medicines Initiative 2 Joint Undertaking, Janssen, Merck KGaA (aka EMD in Canada and USA), Pfizer, the São Paulo Research Foundation-FAPESP, and Takeda. Research reported in this publication was supported by the Department of Defense grant AL220105 and NIH U24DK116204. LC acknowledges support from the Czech Science Foundation (grant 24-10345S). We acknowledge the core facility CELLIM supported by MEYS CR (LM2023050 Czech-BioImaging).

## Abbreviations

ACN: acetonitrile
AcOH: acetic acid
BSA: bovine serum albumin
CPMC: conductor-like polarizable continuum model
DCM: dichloromethane
DEA: diethylamine
DFT: density functional theory
DIPA, *N*: *N*-diisopropylamine
DMA, *N*: *N*-dimethylacetamide
DMAP: 4-dimethylaminopyridine
DMF, *N*: *N*-dimethylformamide
DMSO: dimethyl sulfoxide
DPPP: 1,3-Bis(diphenylphosphino)propane
EDC: 1-ethyl-3-(3-dimethylaminopropyl)carbodiimide
EtOAc: ethyl acetate
Hex: hexanes
HPLC: high-performance liquid chromatography
IC_50_: half maximal inhibitory concentration
IPA: isopropylalcohol
K_2_CO_3_: potassium carbonate
K_m_: Michaelis constant
LC–MS: liquid chromatography–mass spectrometry
MeOH: methanol
NanoBRET: bioluminescence resonance energy transfer using NanoLuciferase
NaOMe: sodium methoxide
NLuc: NanoLuciferase
NMR: nuclear magnetic resonance
Pd(OAc)_2_: Palladium (II) acetate
PEM: photoelastic modulator
TEA: triethylamine
TFA: trifluoroacetic acid
SAR: structure– activity relationships
v/v: volume for volume.

## For Table of Contents Only

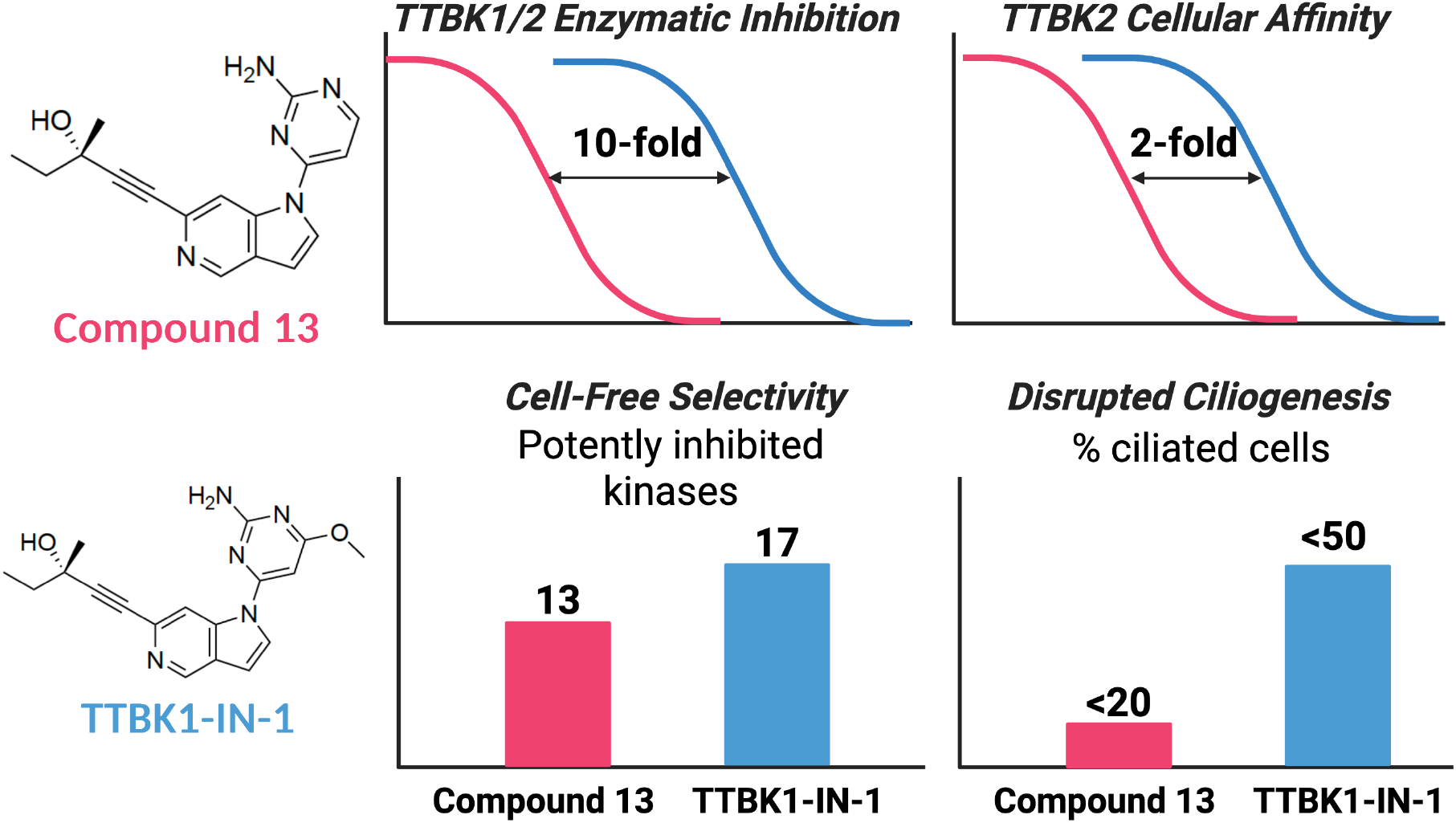

