## Supplemental Files for "Development of Potent and Cell Active 5-Azaindole-Based Tau Tubulin Kinase Inhibitors"

#### Table of Contents

| | |
|---|---|
| Figure S1 | S2 |
| Figure S2 | S3 |
| Figure S3 | S4 |
| Figure S4 | S5 |
| Figure S5 | S6 |
| Figure S6 | S7 |
| Figure S7 | S8 |
| Figure S8 | S9 |
| Figure S9 | S10 |
| Figure S10 | S11 |
| Figure S11 | S12–S39 |
| Table S1 | S40–S45 |
| Table S2 | S46–S53 |
| Table S3 | S54–S61 |
| Table S4 | S62–S63 |
| Purity traces and spectra for all compounds | S64–S123 |

### Figure S1.

A

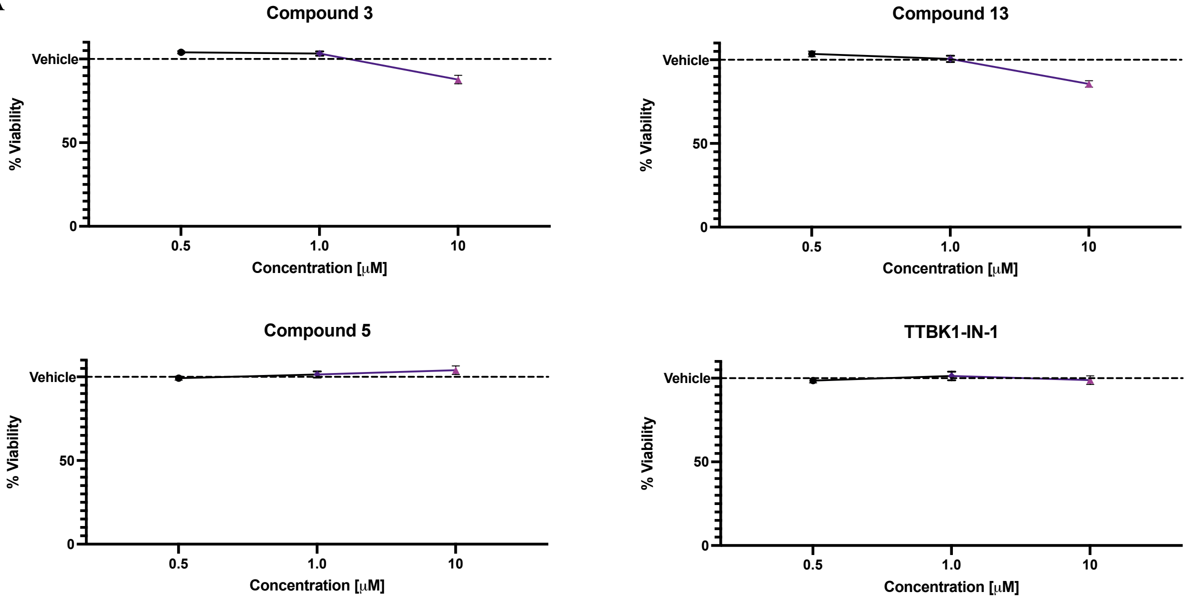


B

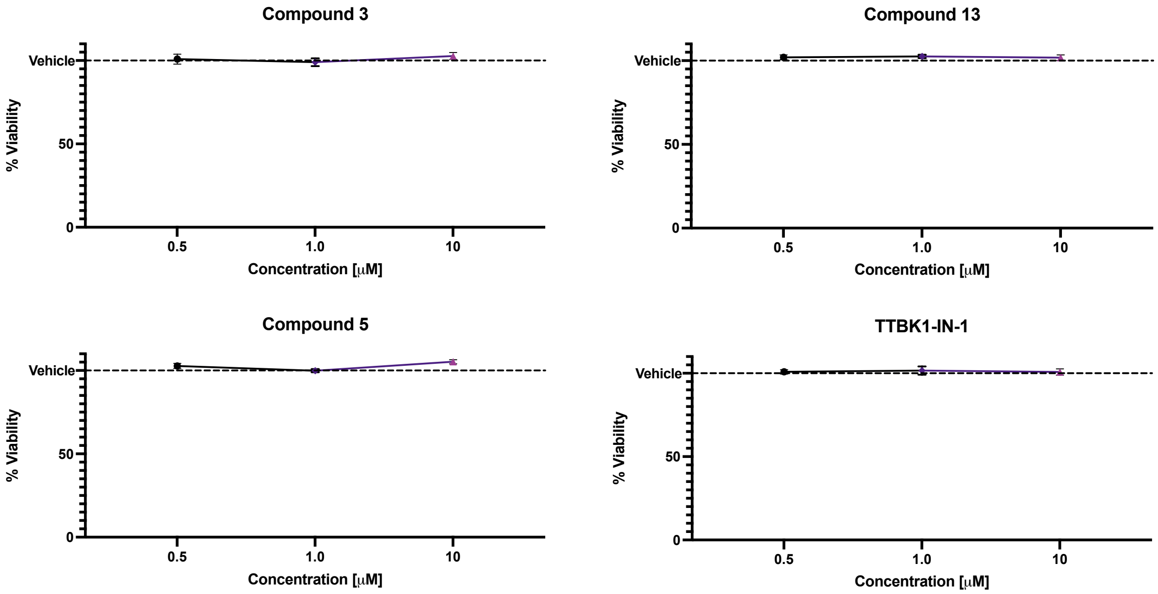


**Figure S1**. Cellular viability assay results for compounds **3**, **13**, **5**, and TTBK1-IN-1 in SH-SY5Y cells. a) Cell Titer Glo results from 8-hour treatment of SH-SY5Y cells at varying doses of compound **3**, **13**, **5**, and TTBK1-IN-1 (N=4). Error bars represent SD. b) Cell Titer Glo results from 8-hour treatment of HEK293 cells at varying doses of compound **3**, **13**, **5**, and TTBK1-IN-1 (N=4). Error bars represent SD.

### Figure S2.

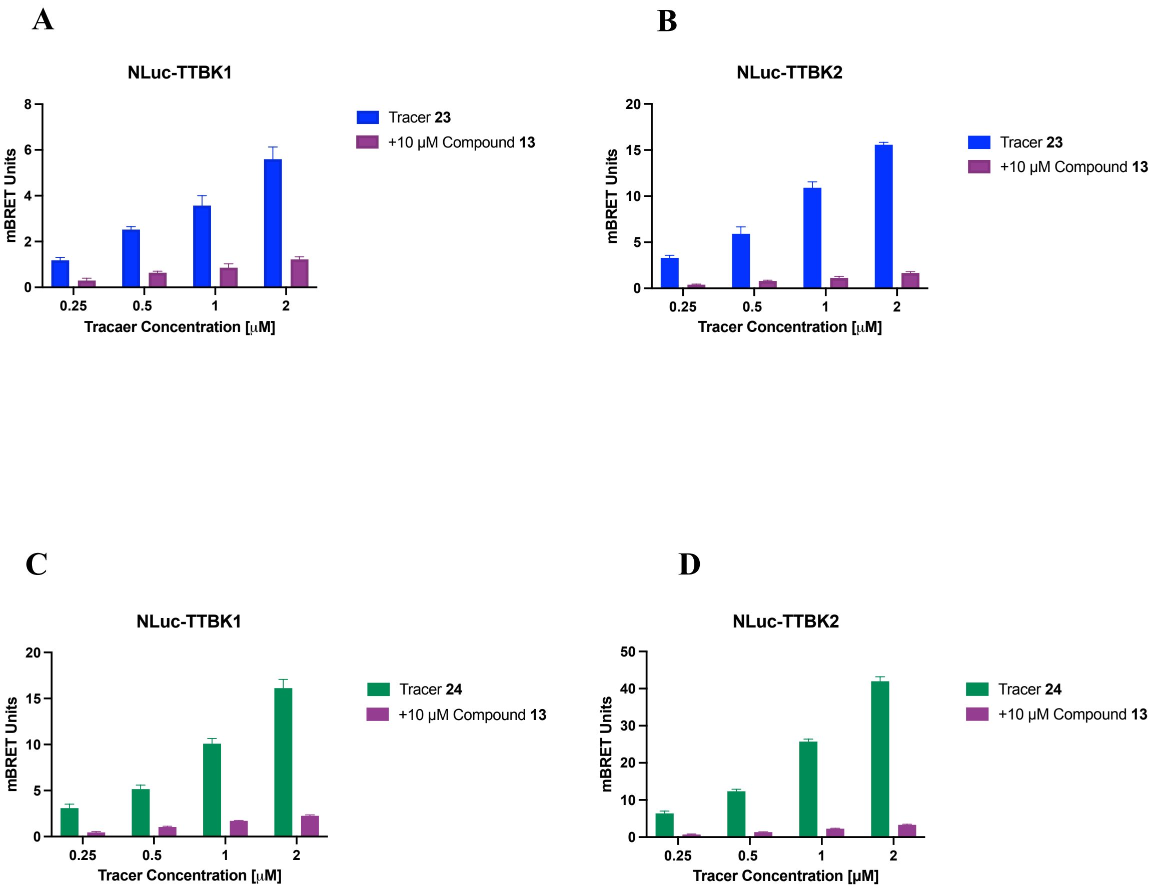


**Figure S2**. Cell-based NanoBRET tracer titration studies. a) NLuc-TTBK1 treated with 0.5–2.0 µM of tracer **23** with or without 10 µM of compound **13** (N=4). Error bars represent SD. b) NLuc-TTBK2 treated with 0.5–2.0 µM of tracer **23** with or without 10 µM of compound **13** (N=4). Error bars represent SD. c) NLuc-TTBK1 treated with 0.5–2.0 µM of tracer **24** with or without 10 µM of compound **13** (N=4). Error bars represent SD. d) NLuc-TTBK2 treated with 0.5–2.0 µM of tracer **24** with or without 10 µM of compound **13** (N=4). Error bars represent SD.

**Figure S3.**

A

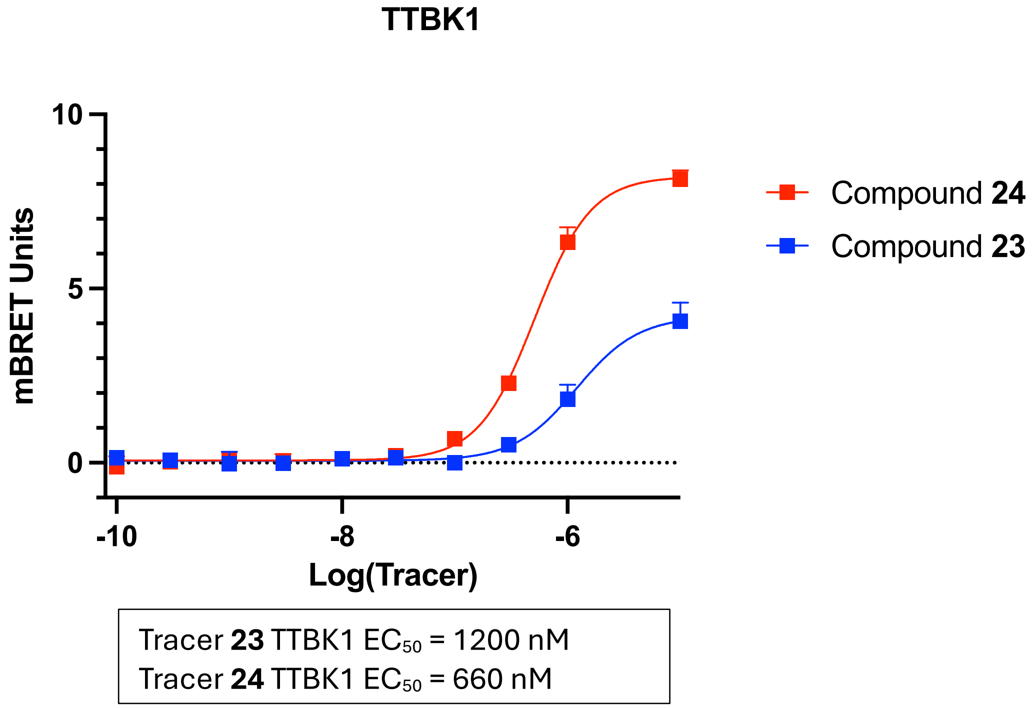


B

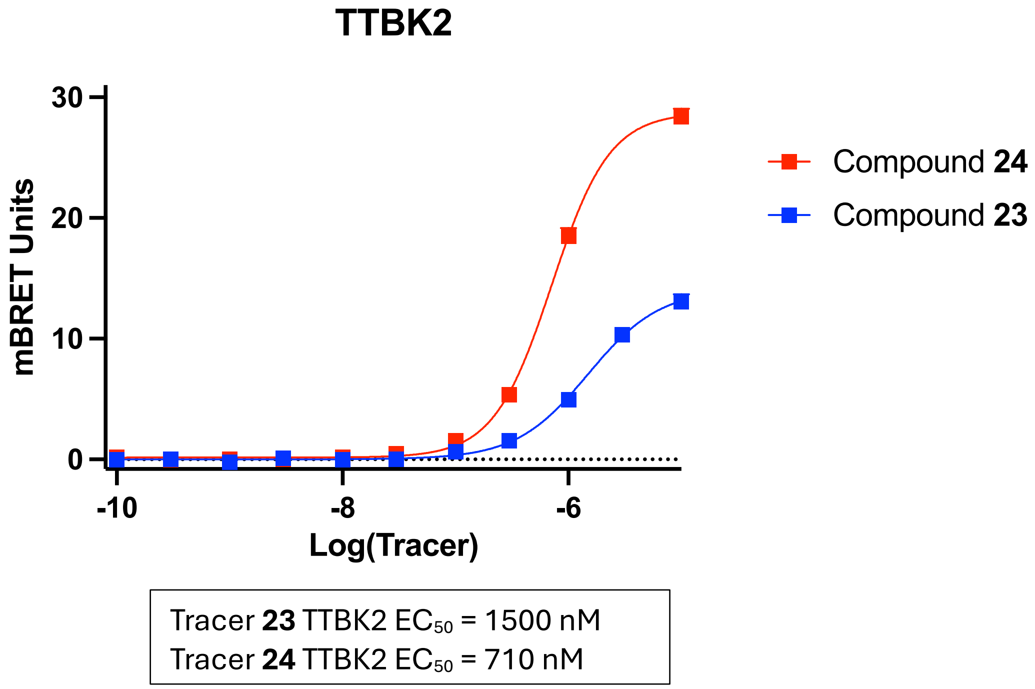


**Figure S3**. $EC_{50}$ calculations for compounds **23** and **24**. a) $EC_{50}$ curves for compound **23** (blue) and compound **24** (red) versus TTBK1 (N=3). b) $EC_{50}$ curves for compound **23** (blue) and compound **24** (red) versus TTBK2 (N=3). Dotted line shown represents the background. Error bars represent SD.

**Figure S4.**

**A**

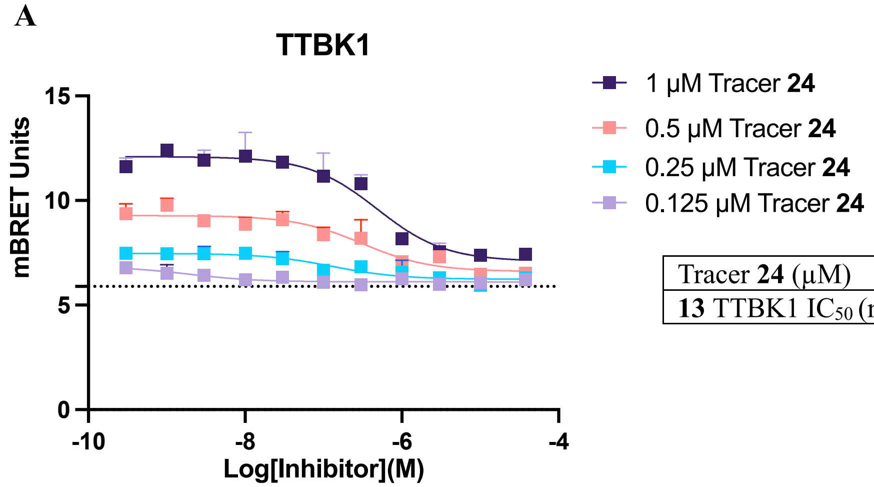


| Tracer **24** (μM) | 0.125 | 0.25 | 0.5 | 1.0 |
|---|---|---|---|---|
| **13** TTBK1 $IC_{50}$ (nM) | 2.0 | 130 | 310 | 400 |

**B**

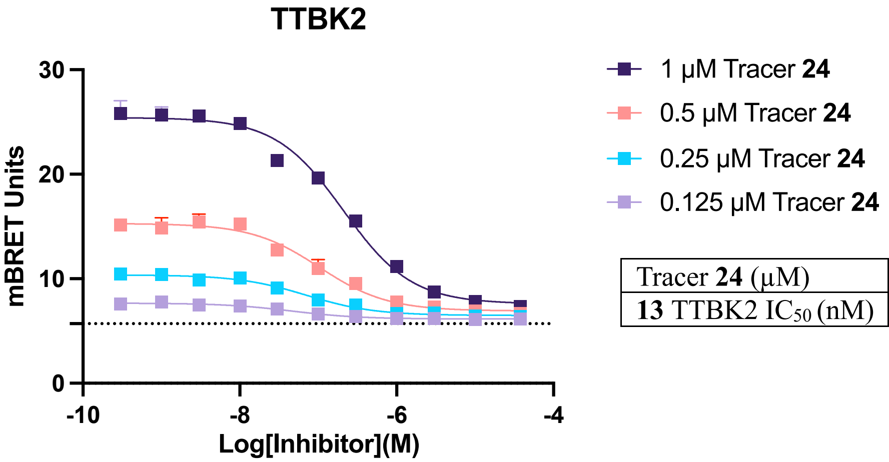


| Tracer **24** (μM) | 0.125 | 0.25 | 0.5 | 1.0 |
|---|---|---|---|---|
| **13** TTBK2 $IC_{50}$ (nM) | 46 | 70 | 100 | 210 |

**Figure S4**. Tracer titration studies using tracer **24** from 0.125–1.0 μM confirm a linear decline in $IC_{50}$ value of compound **13** with reduced tracer concentration. a) NLuc-TTBK1 11-point dose–response curve for compound **13** with varying concentrations of tracer **24** (N=3). Error bars represent SD. b) NLuc-TTBK2 11-point dose–response of compound **13** with varying concentrations of tracer **24** (N=3). Data was not subjected to background subtraction; dotted line indicates the background. Error bars represent SD.

**Figure S5.**

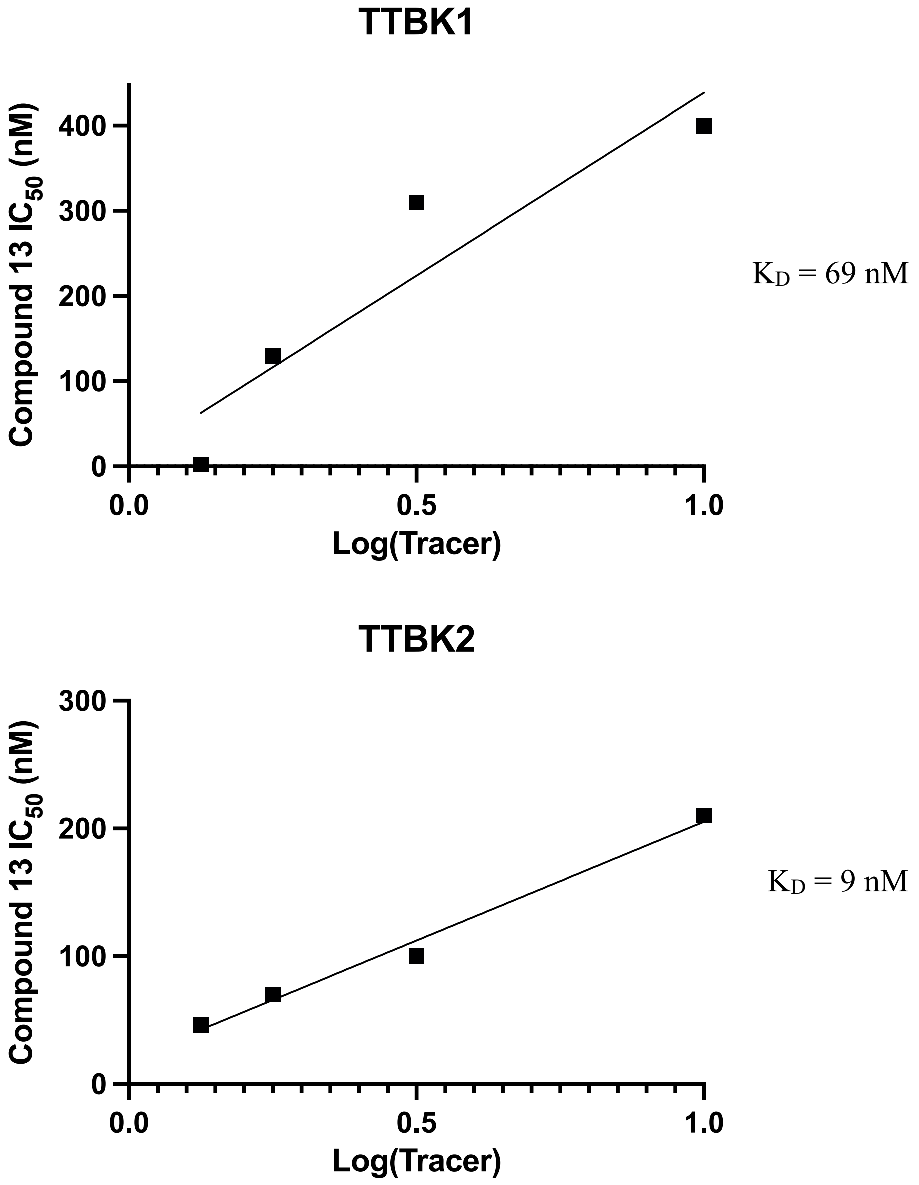


**Figure S5**. Tracer titration studies using tracer **24** from 0.125–1.0 μM confirm a linear decline in $IC_{50}$ value of compound **13** with reduced tracer concentration. Apparent $K_D$ calculated from the y-intercept using linear regression. a) Linear regression of tracer **24** and $IC_{50}$ of compound **13** in TTBK1 and TTBK2 NanoBRET assays (N=3). Error bars represent SD.

### Figure S6.

A

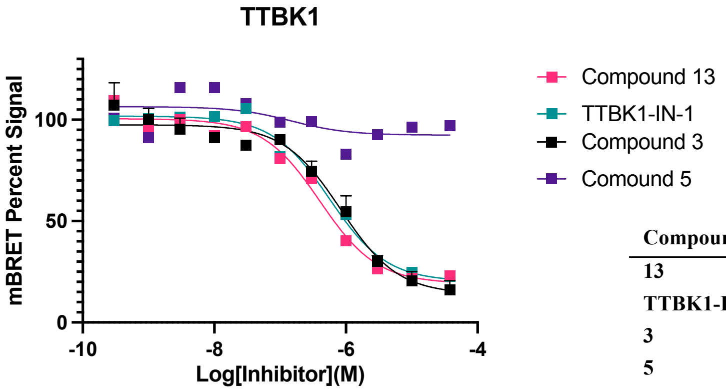


| Compound ID | TTBK1 NB $IC_{50}$ (nM) |
|---|---|
| **13** | 400 |
| **TTBK1-IN-1** | 560 |
| **3** | 830 |
| **5** | >30000 |

B

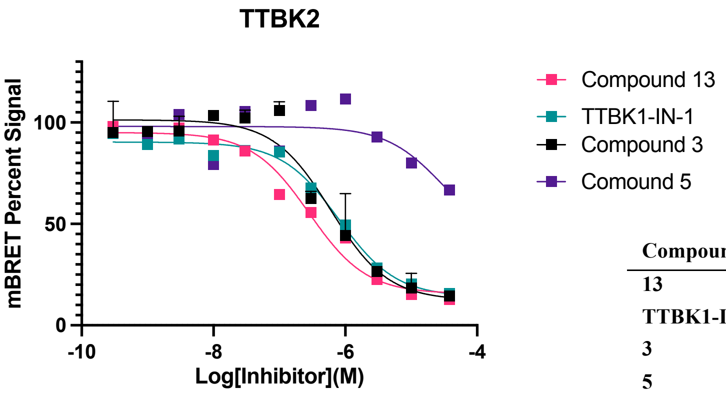


| Compound ID | TTBK2 NB $IC_{50}$ (nM) |
|---|---|
| **13** | 280 |
| **TTBK1-IN-1** | 800 |
| **3** | 600 |
| **5** | >30000 |

**Figure S6**. Normalized NanoBRET dose–response follow-up curves for TTBK1 and TTBK2. a) Representative TTBK1 NanoBRET full dose–response curves for compounds **13**, **3**, **5**, and TTBK1-IN-1 (N=2). b) Representative TTBK2 NanoBRET full dose–response curves for compounds **13**, **3**, **5**, and TTBK1-IN-1 (N=2).

#### Figure S7.

A

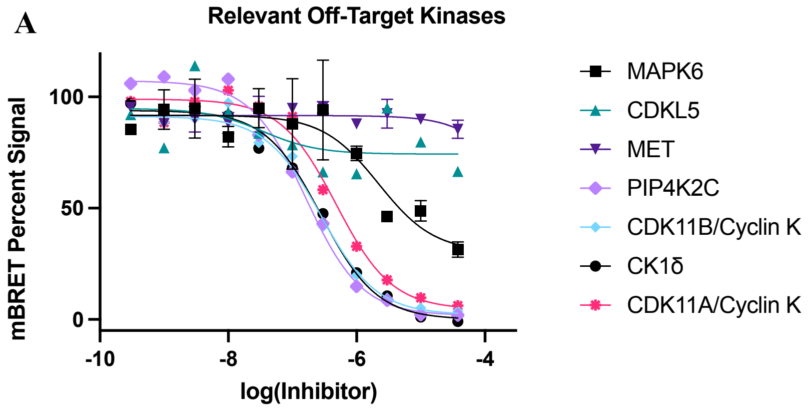


B

| Kinase | K300% Occupancy | NB $IC_{50}$ (nM) |
|---|---|---|
| MAPK6 | 5 | >10000 |
| CDKL5 | 1 | >10000 |
| MET | 2 | >30000 |
| PIP4K2C | 86 | 160 |
| CDK11A | 75 | 480 |
| CDK11B | 85 | 280 |
| CK1δ | 68 | 280 |

**Figure S7.** NanoBRET dose–response follow-up curves for off-target kinases based on kinome-wide selectivity screening in the K192 and K300 panels. a) NanoBRET full dose–response curves for compound **3** off-target kinases (N=1), MAPK6 and MET (N=2). Data have been normalized. b) NanoBRET $IC_{50}$ values for the off-target kinases of compound **3** and K300 % occupancy data, average of N=2.

#### Figure S8.

A

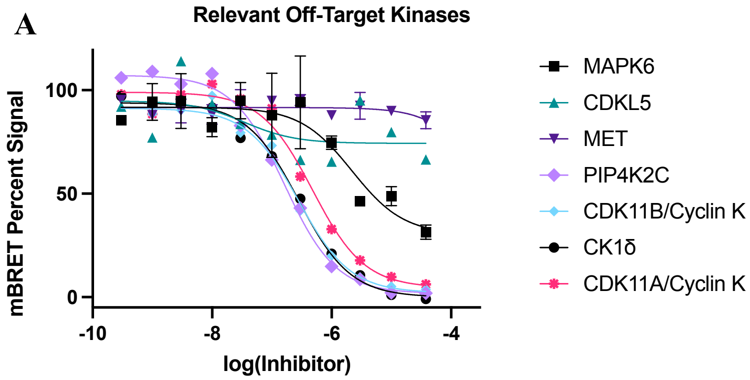


B

| Kinase | Compound **13** NB $IC_{50}$ (nM) |
|---|---|
| PIP4K2C | 1600 |
| PIKFYVE | 90 |
| CDK11B | 160 |
| CK1δ | 190 |
| CK1ε | 1100 |
| EPHA2 | >30000 |
| MAP4K5 | >10000 |
| PAK4 | >10000 |
| PAK5 | >10000 |
| PAK6 | >10000 |
| MAP3K19 | >30000 |

C

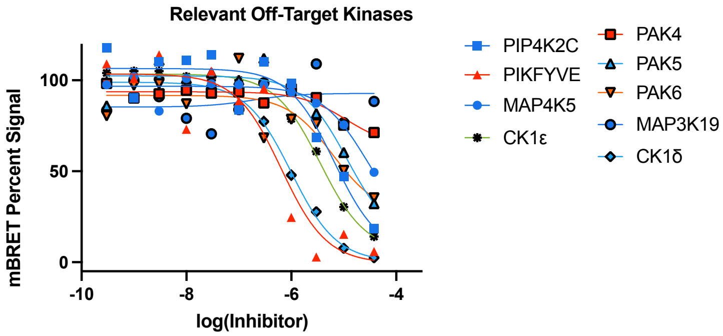


D

| Kinase | TTBK1-IN-1 NB $IC_{50}$ (nM) |
|---|---|
| PIP4K2C | 6600 |
| PIKFYVE | 640 |
| CK1δ | 990 |
| CK1ε | 3500 |
| MAP4K5 | >10000 |
| PAK4 | >10000 |
| PAK5 | >10000 |
| PAK6 | >10000 |
| MAP3K19 | >30000 |

**Figure S8.** NanoBRET dose–response follow-up curves for off-target kinases based on biochemical screening. a) NanoBRET full dose–response curves for compound **13** off-target kinases (N=1), PAK4, PAK5, and PAK6 (N=2). Data have been normalized. b) NanoBRET $IC_{50}$ values for the off-target kinases of compound **13**. c) NanoBRET full dose–response curves for TTBK1-IN-1 off-target kinases (N=1). Normalized mBRET signal. d) NanoBRET $IC_{50}$ values for the off-target kinases of TTBK1-IN-1.

### Figure S9.

**A**

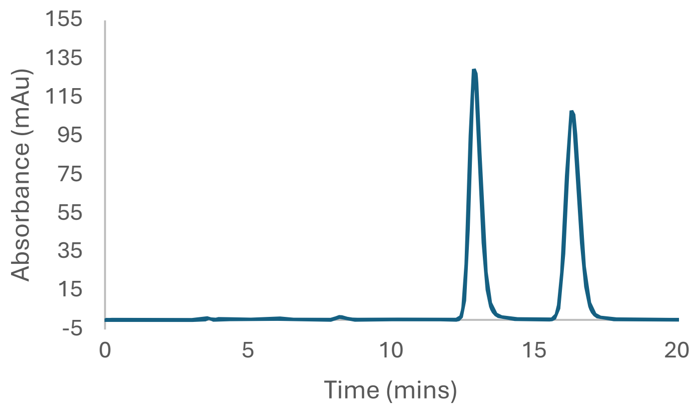


**B**

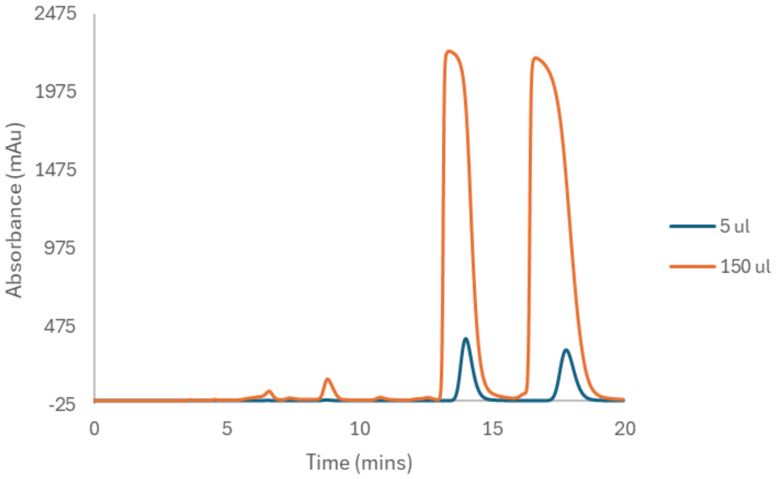


**Figure S9.** Chiral separation of compound **3**. a) CHIRALPAK IM normal phase separation of compound **3**. b) Overloading of **3** on CHIRALPAK IM.

#### Figure S10.

**A**

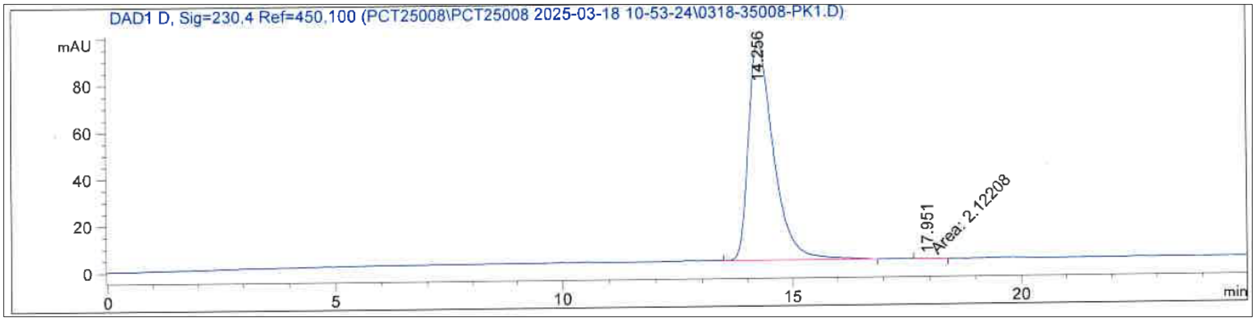


| Peak # | Retention Time (min) | Area (mAU*s) | Area % |
|---|---|---|---|
| 1 | 14.256 | 3572.75049 | 99.9406 |
| 2 | 17.951 | 2.12208 | 0.0594 |
| | Totals: | 3574.87257 | 100.0000 |

**B**

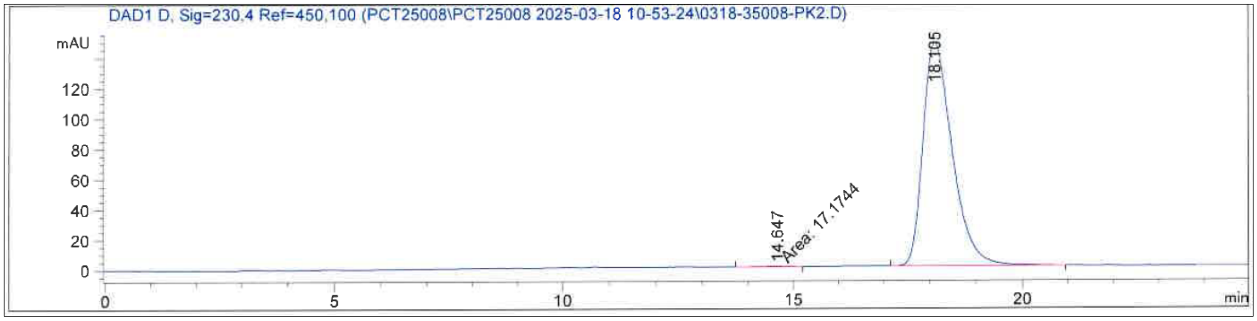


| Peak # | Retention Time (min) | Area (mAU*s) | Area % |
|---|---|---|---|
| 1 | 14.647 | 17.17438 | 0.2629 |
| 2 | 18.105 | 6516.59717 | 99.7371 |
| | Totals: | 6533.77154 | 100.0000 |

**Figure S10**. Analyses of isolated compounds after CHIRALPAK IM. a) Final analysis of isolated compound **3** enantiomer, compound **12** peak 1 CHIRALPAK IM. b) Final analysis of isolated compound **3** enantiomer, compound **13** peak 2 CHIRALPAK IM.

**Figure S11.**

**VCD Calculations Supporting Information:**

There were no imaginary frequencies in any of the conformers for any compounds. Weighted average spectra were produced using both free energy and electronic energy. While both gave the same answer for the stereochemistry, the results using electronic energy were a slightly better match to the experimental data, so this was used in all comparisons and plots.

Compound **3**

<u>(R) stereoisomer</u> (Compound**12**): DFT Level: B3PW91 / cc-pVTZ with CPCM (chloroform): 23 conformers used in Boltzmann average - weighted by electronic energy.

Conformer # 1

Electronic Energy = -1006.956658 Hartree

| Center Number | Atomic Number | Atomic Type | Coordinates (Angstroms) X | Y | Z |
|---|---|---|---|---|---|
| 1 | 6 | 0 | -0.189129 | 0.520211 | -0.013700 |
| 2 | 6 | 0 | 1.056093 | 1.147118 | -0.005962 |
| 3 | 7 | 0 | 1.232038 | 2.488679 | 0.041721 |
| 4 | 6 | 0 | 0.159681 | 3.259420 | 0.074528 |
| 5 | 6 | 0 | -1.138247 | 2.748395 | 0.066283 |
| 6 | 6 | 0 | -1.302311 | 1.347004 | 0.029025 |
| 7 | 6 | 0 | -2.440888 | 3.335834 | 0.079476 |
| 8 | 7 | 0 | -2.666530 | 1.090627 | 0.009731 |
| 9 | 6 | 0 | -3.330096 | 2.314847 | 0.040388 |
| 10 | 6 | 0 | 2.240274 | 0.348279 | -0.047529 |
| 11 | 6 | 0 | 3.235458 | -0.332851 | -0.085124 |
| 12 | 6 | 0 | -4.643828 | -0.259470 | -0.428663 |
| 13 | 6 | 0 | -3.305392 | -0.153712 | -0.046871 |
| 14 | 7 | 0 | -2.580063 | -1.208831 | 0.281486 |
| 15 | 6 | 0 | -3.189112 | -2.402858 | 0.234022 |
| 16 | 7 | 0 | -4.470676 | -2.623176 | -0.094781 |
| 17 | 6 | 0 | -5.164907 | -1.539811 | -0.422202 |
| 18 | 6 | 0 | 4.455587 | -1.152620 | -0.119381 |
| 19 | 7 | 0 | -2.433995 | -3.482903 | 0.528718 |
| 20 | 8 | 0 | 4.728628 | -1.673068 | 1.188045 |
| 21 | 6 | 0 | 4.240318 | -2.374651 | -1.007050 |

| | | | | | |
|---|---|---|---|---|---|
| 22 | 6 | 0 | 5.645235 | -0.321635 | -0.633777 |
| 23 | 6 | 0 | 5.997772 | 0.885882 | 0.220954 |
| 24 | 1 | 0 | -0.269569 | -0.553122 | -0.042066 |
| 25 | 1 | 0 | 0.329043 | 4.331623 | 0.106808 |
| 26 | 1 | 0 | -2.684490 | 4.384504 | 0.123987 |
| 27 | 1 | 0 | -4.404918 | 2.349079 | 0.063412 |
| 28 | 1 | 0 | -5.243575 | 0.581587 | -0.736068 |
| 29 | 1 | 0 | -6.201038 | -1.701471 | -0.703202 |
| 30 | 1 | 0 | -1.530853 | -3.344203 | 0.944025 |
| 31 | 1 | 0 | -2.895646 | -4.364443 | 0.660358 |
| 32 | 1 | 0 | 4.761270 | -0.933890 | 1.803135 |
| 33 | 1 | 0 | 4.030837 | -2.071627 | -2.032381 |
| 34 | 1 | 0 | 5.141168 | -2.990039 | -0.997202 |
| 35 | 1 | 0 | 3.404670 | -2.968065 | -0.637241 |
| 36 | 1 | 0 | 5.417545 | -0.001301 | -1.653633 |
| 37 | 1 | 0 | 6.499617 | -1.001118 | -0.693799 |
| 38 | 1 | 0 | 6.289272 | 0.593877 | 1.232623 |
| 39 | 1 | 0 | 5.160553 | 1.582364 | 0.293776 |
| 40 | 1 | 0 | 6.842634 | 1.423720 | -0.211521 |

---------------------------------------------------------------------

Conformer # 2

Electronic Energy = -1006.956444 Hartree

| Center Number | Atomic Number | Atomic Type | Coordinates (Angstroms) X | Y | Z |
|---|---|---|---|---|---|
| 1 | 6 | 0 | 0.128818 | 0.727141 | 0.023855 |
| 2 | 6 | 0 | -1.034894 | 1.494918 | 0.035097 |
| 3 | 7 | 0 | -1.052861 | 2.848567 | 0.050031 |
| 4 | 6 | 0 | 0.102214 | 3.489512 | 0.045540 |
| 5 | 6 | 0 | 1.331716 | 2.830743 | 0.030620 |
| 6 | 6 | 0 | 1.330903 | 1.419347 | 0.027312 |
| 7 | 6 | 0 | 2.694059 | 3.261993 | 0.007431 |
| 8 | 7 | 0 | 2.655331 | 1.004988 | -0.008130 |
| 9 | 6 | 0 | 3.457468 | 2.143387 | -0.019987 |
| 10 | 6 | 0 | -2.303725 | 0.837009 | 0.034174 |
| 11 | 6 | 0 | -3.366679 | 0.265923 | 0.031315 |
| 12 | 6 | 0 | 4.449423 | -0.576353 | -0.458819 |
| 13 | 6 | 0 | 3.143563 | -0.306461 | -0.046528 |
| 14 | 7 | 0 | 2.310251 | -1.261621 | 0.328553 |

| | | | | | |
|---|---|---|---|---|---|
| 15 | 6 | 0 | 2.775680 | -2.519225 | 0.298069 |
| 16 | 7 | 0 | 4.013268 | -2.895109 | -0.057182 |
| 17 | 6 | 0 | 4.818802 | -1.908186 | -0.431322 |
| 18 | 6 | 0 | -4.679116 | -0.397774 | 0.048626 |
| 19 | 7 | 0 | 1.909623 | -3.496730 | 0.641215 |
| 20 | 8 | 0 | -4.918796 | -0.971291 | 1.339991 |
| 21 | 6 | 0 | -5.771473 | 0.622307 | -0.279877 |
| 22 | 6 | 0 | -4.721678 | -1.564157 | -0.948401 |
| 23 | 6 | 0 | -3.677632 | -2.643928 | -0.714847 |
| 24 | 1 | 0 | 0.083970 | -0.348627 | 0.022645 |
| 25 | 1 | 0 | 0.059104 | 4.574671 | 0.051874 |
| 26 | 1 | 0 | 3.058989 | 4.275756 | 0.021217 |
| 27 | 1 | 0 | 4.529196 | 2.051623 | -0.014595 |
| 28 | 1 | 0 | 5.132812 | 0.182056 | -0.804430 |
| 29 | 1 | 0 | 5.820805 | -2.196050 | -0.734285 |
| 30 | 1 | 0 | 1.039669 | -3.244577 | 1.073556 |
| 31 | 1 | 0 | 2.269228 | -4.423094 | 0.782859 |
| 32 | 1 | 0 | -4.848286 | -0.268267 | 1.993385 |
| 33 | 1 | 0 | -5.617311 | 1.054071 | -1.269406 |
| 34 | 1 | 0 | -6.744067 | 0.128878 | -0.255199 |
| 35 | 1 | 0 | -5.769624 | 1.434030 | 0.449468 |
| 36 | 1 | 0 | -4.618166 | -1.148524 | -1.953165 |
| 37 | 1 | 0 | -5.725952 | -1.991718 | -0.881984 |
| 38 | 1 | 0 | -3.775223 | -3.076094 | 0.281178 |
| 39 | 1 | 0 | -2.666610 | -2.244751 | -0.813088 |
| 40 | 1 | 0 | -3.792164 | -3.446893 | -1.444923 |

---------------------------------------------------------------------

Conformer # 3
Electronic Energy = -1006.956113 Hartree

| Center Number | Atomic Number | Atomic Type | Coordinates (Angstroms) X | Y | Z |
|---|---|---|---|---|---|
| 1 | 6 | 0 | 0.185959 | 0.531036 | -0.084273 |
| 2 | 6 | 0 | -1.049999 | 1.171537 | -0.160896 |
| 3 | 7 | 0 | -1.207321 | 2.513779 | -0.242429 |
| 4 | 6 | 0 | -0.125219 | 3.271512 | -0.242741 |
| 5 | 6 | 0 | 1.164758 | 2.746203 | -0.167982 |
| 6 | 6 | 0 | 1.310007 | 1.344163 | -0.094613 |
| 7 | 6 | 0 | 2.473795 | 3.318632 | -0.137868 |

| | | | | | |
|---|---|---|---|---|---|
| 8 | 7 | 0 | 2.669164 | 1.072951 | -0.011520 |
| 9 | 6 | 0 | 3.348131 | 2.288917 | -0.039831 |
| 10 | 6 | 0 | -2.245113 | 0.387923 | -0.158811 |
| 11 | 6 | 0 | -3.250736 | -0.278862 | -0.153593 |
| 12 | 6 | 0 | 4.613220 | -0.287996 | 0.530579 |
| 13 | 6 | 0 | 3.290166 | -0.176621 | 0.099584 |
| 14 | 7 | 0 | 2.563815 | -1.231717 | -0.226808 |
| 15 | 6 | 0 | 3.156000 | -2.430952 | -0.127489 |
| 16 | 7 | 0 | 4.422876 | -2.657344 | 0.250538 |
| 17 | 6 | 0 | 5.118614 | -1.573845 | 0.574169 |
| 18 | 6 | 0 | -4.487505 | -1.074324 | -0.172642 |
| 19 | 7 | 0 | 2.397965 | -3.509730 | -0.419201 |
| 20 | 8 | 0 | -4.589509 | -1.774737 | -1.419731 |
| 21 | 6 | 0 | -4.436560 | -2.162913 | 0.894557 |
| 22 | 6 | 0 | -5.714131 | -0.157591 | -0.010095 |
| 23 | 6 | 0 | -5.765651 | 0.658210 | 1.272744 |
| 24 | 1 | 0 | 0.252112 | -0.542222 | -0.030301 |
| 25 | 1 | 0 | -0.279976 | 4.344734 | -0.303148 |
| 26 | 1 | 0 | 2.731685 | 4.363331 | -0.193883 |
| 27 | 1 | 0 | 4.423268 | 2.310972 | -0.018110 |
| 28 | 1 | 0 | 5.212610 | 0.553538 | 0.837188 |
| 29 | 1 | 0 | 6.142610 | -1.739756 | 0.894452 |
| 30 | 1 | 0 | 1.510639 | -3.372086 | -0.867599 |
| 31 | 1 | 0 | 2.852637 | -4.399866 | -0.511158 |
| 32 | 1 | 0 | -4.608253 | -1.119137 | -2.124657 |
| 33 | 1 | 0 | -4.311200 | -1.734550 | 1.887394 |
| 34 | 1 | 0 | -5.364053 | -2.736959 | 0.869267 |
| 35 | 1 | 0 | -3.601254 | -2.834405 | 0.698328 |
| 36 | 1 | 0 | -6.594466 | -0.800520 | -0.090646 |
| 37 | 1 | 0 | -5.734132 | 0.517772 | -0.871023 |
| 38 | 1 | 0 | -4.881027 | 1.288791 | 1.376699 |
| 39 | 1 | 0 | -5.834505 | 0.022302 | 2.156695 |
| 40 | 1 | 0 | -6.640545 | 1.310197 | 1.269486 |

---------------------------------------------------------------------

Conformer # 4
Electronic Energy = -1006.956073 Hartree

| Center Number | Atomic Number | Atomic Type | Coordinates (Angstroms) X | Y | Z |
|---|---|---|---|---|---|

---------------------------------------------------------------------

| | | | | | |
|---|---|---|---|---|---|
| 1 | 6 | 0 | 0.129751 | 0.728134 | -0.016273 |
| 2 | 6 | 0 | -1.033525 | 1.496404 | -0.032390 |
| 3 | 7 | 0 | -1.050528 | 2.849512 | -0.075001 |
| 4 | 6 | 0 | 0.104601 | 3.489695 | -0.094027 |
| 5 | 6 | 0 | 1.333832 | 2.830468 | -0.076072 |
| 6 | 6 | 0 | 1.332138 | 1.419499 | -0.044041 |
| 7 | 6 | 0 | 2.696487 | 3.261235 | -0.072527 |
| 8 | 7 | 0 | 2.656567 | 1.005153 | -0.010627 |
| 9 | 6 | 0 | 3.459510 | 2.142944 | -0.028028 |
| 10 | 6 | 0 | -2.304643 | 0.843106 | -0.006191 |
| 11 | 6 | 0 | -3.378240 | 0.292924 | 0.019126 |
| 12 | 6 | 0 | 4.451172 | -0.566827 | 0.466613 |
| 13 | 6 | 0 | 3.144986 | -0.305197 | 0.050185 |
| 14 | 7 | 0 | 2.312200 | -1.267341 | -0.308236 |
| 15 | 6 | 0 | 2.779226 | -2.523575 | -0.256774 |
| 16 | 7 | 0 | 4.016936 | -2.892244 | 0.104750 |
| 17 | 6 | 0 | 4.821869 | -1.898493 | 0.462027 |
| 18 | 6 | 0 | -4.692129 | -0.366506 | 0.058166 |
| 19 | 7 | 0 | 1.913752 | -3.508197 | -0.582944 |
| 20 | 8 | 0 | -5.188878 | -0.382803 | 1.402731 |
| 21 | 6 | 0 | -5.705109 | 0.448050 | -0.740804 |
| 22 | 6 | 0 | -4.596062 | -1.797157 | -0.502813 |
| 23 | 6 | 0 | -3.684418 | -2.730506 | 0.278398 |
| 24 | 1 | 0 | 0.084928 | -0.347458 | 0.006803 |
| 25 | 1 | 0 | 0.061823 | 4.574506 | -0.122364 |
| 26 | 1 | 0 | 3.061661 | 4.274301 | -0.110101 |
| 27 | 1 | 0 | 4.531097 | 2.049929 | -0.040119 |
| 28 | 1 | 0 | 5.133554 | 0.198472 | 0.798835 |
| 29 | 1 | 0 | 5.824091 | -2.180342 | 0.769803 |
| 30 | 1 | 0 | 1.048194 | -3.262882 | -1.027979 |
| 31 | 1 | 0 | 2.277577 | -4.434413 | -0.715283 |
| 32 | 1 | 0 | -4.512322 | -0.769211 | 1.967119 |
| 33 | 1 | 0 | -5.413234 | 0.501141 | -1.789123 |
| 34 | 1 | 0 | -6.684561 | -0.027162 | -0.669516 |
| 35 | 1 | 0 | -5.773647 | 1.459804 | -0.342665 |
| 36 | 1 | 0 | -4.260629 | -1.731153 | -1.540968 |
| 37 | 1 | 0 | -5.613476 | -2.197003 | -0.520103 |
| 38 | 1 | 0 | -4.036710 | -2.875544 | 1.302148 |
| 39 | 1 | 0 | -2.662271 | -2.349264 | 0.320067 |
| 40 | 1 | 0 | -3.654224 | -3.714681 | -0.191458 |

-----------------------------------------------------------------

Conformer # 5
Electronic Energy = -1006.955914 Hartree

| Center Number | Atomic Number | Atomic Type | Coordinates (Angstroms) X | Y | Z |
|---|---|---|---|---|---|
| 1 | 6 | 0 | -0.045748 | 0.397703 | -0.039266 |
| 2 | 6 | 0 | -1.293578 | 1.017612 | 0.013172 |
| 3 | 7 | 0 | -1.472359 | 2.358945 | 0.025389 |
| 4 | 6 | 0 | -0.402809 | 3.134640 | 0.007254 |
| 5 | 6 | 0 | 0.896980 | 2.630425 | -0.039722 |
| 6 | 6 | 0 | 1.064062 | 1.228459 | -0.087479 |
| 7 | 6 | 0 | 2.199443 | 3.218102 | -0.024353 |
| 8 | 7 | 0 | 2.424728 | 0.976736 | -0.107709 |
| 9 | 6 | 0 | 3.090886 | 2.196967 | -0.053006 |
| 10 | 6 | 0 | -2.473039 | 0.212979 | 0.061863 |
| 11 | 6 | 0 | -3.464767 | -0.472863 | 0.103853 |
| 12 | 6 | 0 | 2.526060 | -1.371425 | -0.786188 |
| 13 | 6 | 0 | 3.091628 | -0.255401 | -0.176699 |
| 14 | 7 | 0 | 4.305895 | -0.271341 | 0.350112 |
| 15 | 6 | 0 | 4.977088 | -1.429274 | 0.284746 |
| 16 | 7 | 0 | 4.521151 | -2.576854 | -0.243366 |
| 17 | 6 | 0 | 3.307648 | -2.514754 | -0.774083 |
| 18 | 6 | 0 | -4.681521 | -1.297581 | 0.143254 |
| 19 | 7 | 0 | 6.232383 | -1.437351 | 0.781369 |
| 20 | 8 | 0 | -4.958278 | -1.818073 | -1.163071 |
| 21 | 6 | 0 | -4.456988 | -2.519251 | 1.029090 |
| 22 | 6 | 0 | -5.871685 | -0.471194 | 0.663935 |
| 23 | 6 | 0 | -6.232310 | 0.736159 | -0.187631 |
| 24 | 1 | 0 | 0.015397 | -0.679142 | -0.020722 |
| 25 | 1 | 0 | -0.576708 | 4.206177 | 0.035969 |
| 26 | 1 | 0 | 2.440378 | 4.268036 | 0.001227 |
| 27 | 1 | 0 | 4.166549 | 2.203731 | -0.054974 |
| 28 | 1 | 0 | 1.569186 | -1.352244 | -1.280083 |
| 29 | 1 | 0 | 2.934180 | -3.428235 | -1.226472 |
| 30 | 1 | 0 | 6.541450 | -0.655745 | 1.329841 |
| 31 | 1 | 0 | 6.694814 | -2.321080 | 0.893983 |
| 32 | 1 | 0 | -5.002820 | -1.078216 | -1.776606 |
| 33 | 1 | 0 | -4.243538 | -2.216102 | 2.053550 |

| | | | | | |
|---|---|---|---|---|---|
| 34 | 1 | 0 | -5.355655 | -3.137835 | 1.023255 |
| 35 | 1 | 0 | -3.621135 | -3.109410 | 0.654572 |
| 36 | 1 | 0 | -5.640557 | -0.151294 | 1.683144 |
| 37 | 1 | 0 | -6.723336 | -1.153791 | 0.726960 |
| 38 | 1 | 0 | -6.527230 | 0.444798 | -1.198509 |
| 39 | 1 | 0 | -5.398134 | 1.436023 | -0.262878 |
| 40 | 1 | 0 | -7.077306 | 1.270155 | 0.249301 |

---------------------------------------------------------------------

Conformer # 6

Electronic Energy = -1006.955907 Hartree

| Center Number | Atomic Number | Atomic Type | Coordinates (Angstroms) X | Y | Z |
|---|---|---|---|---|---|
| 1 | 6 | 0 | -0.151844 | 0.673200 | -0.091256 |
| 2 | 6 | 0 | 1.025779 | 1.418857 | -0.123071 |
| 3 | 7 | 0 | 1.069991 | 2.771862 | -0.103970 |
| 4 | 6 | 0 | -0.071195 | 3.434285 | -0.042319 |
| 5 | 6 | 0 | -1.312234 | 2.798759 | 0.000044 |
| 6 | 6 | 0 | -1.339307 | 1.387996 | -0.033341 |
| 7 | 6 | 0 | -2.664006 | 3.254923 | 0.083837 |
| 8 | 7 | 0 | -2.669613 | 0.997942 | 0.039118 |
| 9 | 6 | 0 | -3.448006 | 2.150625 | 0.109691 |
| 10 | 6 | 0 | 2.280947 | 0.737780 | -0.180314 |
| 11 | 6 | 0 | 3.334104 | 0.150391 | -0.224141 |
| 12 | 6 | 0 | -4.477235 | -0.560697 | 0.514083 |
| 13 | 6 | 0 | -3.182100 | -0.304551 | 0.061263 |
| 14 | 7 | 0 | -2.382118 | -1.264999 | -0.368980 |
| 15 | 6 | 0 | -2.871032 | -2.513858 | -0.354393 |
| 16 | 7 | 0 | -4.101967 | -2.875727 | 0.036560 |
| 17 | 6 | 0 | -4.873628 | -1.884169 | 0.465717 |
| 18 | 6 | 0 | 4.624982 | -0.549983 | -0.297560 |
| 19 | 7 | 0 | -2.037365 | -3.498063 | -0.754592 |
| 20 | 8 | 0 | 4.863999 | -0.977241 | -1.645699 |
| 21 | 6 | 0 | 5.749911 | 0.381023 | 0.156778 |
| 22 | 6 | 0 | 4.584398 | -1.852173 | 0.515494 |
| 23 | 6 | 0 | 4.315124 | -1.687540 | 2.002310 |
| 24 | 1 | 0 | -0.128330 | -0.402898 | -0.119917 |
| 25 | 1 | 0 | -0.006733 | 4.518217 | -0.022650 |
| 26 | 1 | 0 | -3.008914 | 4.275434 | 0.109729 |

| | | | | | |
|---|---|---|---|---|---|
| 27 | 1 | 0 | -4.520862 | 2.079360 | 0.140620 |
| 28 | 1 | 0 | -5.132498 | 0.200922 | 0.904273 |
| 29 | 1 | 0 | -5.869440 | -2.161066 | 0.797881 |
| 30 | 1 | 0 | -1.180702 | -3.249829 | -1.214950 |
| 31 | 1 | 0 | -2.421419 | -4.412358 | -0.910032 |
| 32 | 1 | 0 | 4.866670 | -0.195124 | -2.206937 |
| 33 | 1 | 0 | 5.590889 | 0.731265 | 1.176393 |
| 34 | 1 | 0 | 6.701335 | -0.150592 | 0.106767 |
| 35 | 1 | 0 | 5.801178 | 1.256216 | -0.493164 |
| 36 | 1 | 0 | 5.545147 | -2.348338 | 0.354523 |
| 37 | 1 | 0 | 3.819635 | -2.487938 | 0.063884 |
| 38 | 1 | 0 | 3.370769 | -1.169894 | 2.180633 |
| 39 | 1 | 0 | 5.106538 | -1.126630 | 2.501834 |
| 40 | 1 | 0 | 4.254295 | -2.665136 | 2.482821 |

---------------------------------------------------------------------

Conformer # 7
Electronic Energy = -1006.955892 Hartree

| Center Number | Atomic Number | Atomic Type | Coordinates (Angstroms) X | Y | Z |
|---|---|---|---|---|---|
| 1 | 6 | 0 | 0.048241 | 0.397960 | -0.135751 |
| 2 | 6 | 0 | 1.294867 | 1.022573 | -0.140196 |
| 3 | 7 | 0 | 1.469240 | 2.364190 | -0.170098 |
| 4 | 6 | 0 | 0.396853 | 3.136153 | -0.176412 |
| 5 | 6 | 0 | -0.901889 | 2.627375 | -0.167121 |
| 6 | 6 | 0 | -1.065551 | 1.224166 | -0.169756 |
| 7 | 6 | 0 | -2.205129 | 3.212062 | -0.126036 |
| 8 | 7 | 0 | -2.425355 | 0.968662 | -0.138271 |
| 9 | 6 | 0 | -3.093517 | 2.188372 | -0.096373 |
| 10 | 6 | 0 | 2.478720 | 0.223631 | -0.105685 |
| 11 | 6 | 0 | 3.475398 | -0.455510 | -0.073496 |
| 12 | 6 | 0 | -2.541919 | -1.399472 | -0.740914 |
| 13 | 6 | 0 | -3.090303 | -0.266399 | -0.147492 |
| 14 | 7 | 0 | -4.286284 | -0.268548 | 0.419765 |
| 15 | 6 | 0 | -4.955573 | -1.429460 | 0.412858 |
| 16 | 7 | 0 | -4.513608 | -2.591947 | -0.094157 |
| 17 | 6 | 0 | -3.318726 | -2.543687 | -0.666972 |
| 18 | 6 | 0 | 4.697275 | -1.271808 | -0.019647 |
| 19 | 7 | 0 | -6.193817 | -1.424720 | 0.950625 |

| | | | | | |
|---|---|---|---|---|---|
| 20 | 8 | 0 | 4.738032 | -2.004566 | 1.211144 |
| 21 | 6 | 0 | 4.671137 | -2.327510 | -1.121063 |
| 22 | 6 | 0 | 5.943457 | -0.378993 | -0.165075 |
| 23 | 6 | 0 | 6.109211 | 0.669334 | 0.924033 |
| 24 | 1 | 0 | -0.008335 | -0.678190 | -0.086847 |
| 25 | 1 | 0 | 0.567727 | 4.208541 | -0.183588 |
| 26 | 1 | 0 | -2.448621 | 4.261713 | -0.122514 |
| 27 | 1 | 0 | -4.168670 | 2.192435 | -0.063178 |
| 28 | 1 | 0 | -1.602217 | -1.393462 | -1.267023 |
| 29 | 1 | 0 | -2.957504 | -3.470026 | -1.102848 |
| 30 | 1 | 0 | -6.487246 | -0.626957 | 1.484228 |
| 31 | 1 | 0 | -6.649380 | -2.305489 | 1.106232 |
| 32 | 1 | 0 | 4.645821 | -1.378290 | 1.935726 |
| 33 | 1 | 0 | 4.643820 | -1.856593 | -2.103130 |
| 34 | 1 | 0 | 5.568314 | -2.944105 | -1.049280 |
| 35 | 1 | 0 | 3.795212 | -2.966234 | -1.011714 |
| 36 | 1 | 0 | 5.898791 | 0.105765 | -1.143567 |
| 37 | 1 | 0 | 6.808266 | -1.047737 | -0.177233 |
| 38 | 1 | 0 | 6.219692 | 0.213916 | 1.910986 |
| 39 | 1 | 0 | 5.257363 | 1.351072 | 0.954800 |
| 40 | 1 | 0 | 7.006165 | 1.263997 | 0.745334 |

---------------------------------------------------------------------

Conformer # 8
Electronic Energy = -1006.955727 Hartree

| Center Number | Atomic Number | Atomic Type | Coordinates (Angstroms) X | Y | Z |
|---|---|---|---|---|---|
| 1 | 6 | 0 | -0.132655 | 0.629997 | 0.076809 |
| 2 | 6 | 0 | -1.321460 | 1.356601 | 0.021900 |
| 3 | 7 | 0 | -1.380562 | 2.707240 | -0.036657 |
| 4 | 6 | 0 | -0.246320 | 3.384463 | -0.065397 |
| 5 | 6 | 0 | 1.004436 | 2.768327 | -0.021573 |
| 6 | 6 | 0 | 1.047002 | 1.359713 | 0.076218 |
| 7 | 6 | 0 | 2.353694 | 3.235631 | -0.080406 |
| 8 | 7 | 0 | 2.380011 | 0.987717 | 0.083742 |
| 9 | 6 | 0 | 3.151103 | 2.140277 | -0.027989 |
| 10 | 6 | 0 | -2.567980 | 0.658174 | 0.022216 |
| 11 | 6 | 0 | -3.612811 | 0.054642 | 0.021767 |
| 12 | 6 | 0 | 2.281798 | -1.330337 | 0.856015 |

| | | | | | |
|---|---|---|---|---|---|
| 13 | 6 | 0 | 2.934173 | -0.296323 | 0.191277 |
| 14 | 7 | 0 | 4.130717 | -0.444631 | -0.355172 |
| 15 | 6 | 0 | 4.694191 | -1.656279 | -0.253491 |
| 16 | 7 | 0 | 4.146273 | -2.735008 | 0.329495 |
| 17 | 6 | 0 | 2.954756 | -2.540384 | 0.877978 |
| 18 | 6 | 0 | -4.904408 | -0.648760 | 0.041866 |
| 19 | 7 | 0 | 5.932875 | -1.799816 | -0.770924 |
| 20 | 8 | 0 | -5.110876 | -1.252451 | 1.324779 |
| 21 | 6 | 0 | -6.030477 | 0.343726 | -0.255795 |
| 22 | 6 | 0 | -4.923434 | -1.797583 | -0.976207 |
| 23 | 6 | 0 | -3.846461 | -2.850768 | -0.772573 |
| 24 | 1 | 0 | -0.166811 | -0.447957 | 0.096654 |
| 25 | 1 | 0 | -0.325301 | 4.465593 | -0.130760 |
| 26 | 1 | 0 | 2.686723 | 4.258090 | -0.149424 |
| 27 | 1 | 0 | 4.222976 | 2.050545 | -0.043732 |
| 28 | 1 | 0 | 1.341153 | -1.203190 | 1.364840 |
| 29 | 1 | 0 | 2.508892 | -3.396928 | 1.374069 |
| 30 | 1 | 0 | 6.299864 | -1.072675 | -1.357335 |
| 31 | 1 | 0 | 6.309089 | -2.726472 | -0.856372 |
| 32 | 1 | 0 | -5.061109 | -0.558730 | 1.989898 |
| 33 | 1 | 0 | -5.901289 | 0.796675 | -1.239367 |
| 34 | 1 | 0 | -6.987474 | -0.179124 | -0.228282 |
| 35 | 1 | 0 | -6.043930 | 1.142602 | 0.487467 |
| 36 | 1 | 0 | -4.842472 | -1.360960 | -1.974095 |
| 37 | 1 | 0 | -5.914212 | -2.255141 | -0.907566 |
| 38 | 1 | 0 | -3.919609 | -3.301177 | 0.217390 |
| 39 | 1 | 0 | -2.848217 | -2.421884 | -0.876482 |
| 40 | 1 | 0 | -3.947344 | -3.644877 | -1.514223 |

---------------------------------------------------------------------

Conformer # 9
Electronic Energy = -1006.955707 Hartree

| Center Number | Atomic Number | Atomic Type | Coordinates (Angstroms) X | Y | Z |
|---|---|---|---|---|---|
| 1 | 6 | 0 | -0.124572 | 0.611650 | -0.030229 |
| 2 | 6 | 0 | -1.312704 | 1.341306 | -0.013540 |
| 3 | 7 | 0 | -1.373077 | 2.690765 | -0.092752 |
| 4 | 6 | 0 | -0.240004 | 3.366036 | -0.171227 |
| 5 | 6 | 0 | 1.010062 | 2.747422 | -0.189326 |

| | | | | | |
|---|---|---|---|---|---|
| 6 | 6 | 0 | 1.053111 | 1.336273 | -0.140500 |
| 7 | 6 | 0 | 2.359270 | 3.217526 | -0.221259 |
| 8 | 7 | 0 | 2.386359 | 0.964612 | -0.150466 |
| 9 | 6 | 0 | 3.157478 | 2.122293 | -0.183656 |
| 10 | 6 | 0 | -2.557448 | 0.648611 | 0.098112 |
| 11 | 6 | 0 | -3.601855 | 0.051858 | 0.192846 |
| 12 | 6 | 0 | 2.270513 | -1.424001 | -0.665230 |
| 13 | 6 | 0 | 2.941623 | -0.323348 | -0.140637 |
| 14 | 7 | 0 | 4.158676 | -0.410964 | 0.372788 |
| 15 | 6 | 0 | 4.724638 | -1.625489 | 0.380437 |
| 16 | 7 | 0 | 4.161213 | -2.761544 | -0.061850 |
| 17 | 6 | 0 | 2.948862 | -2.628538 | -0.582204 |
| 18 | 6 | 0 | -4.889380 | -0.644253 | 0.338434 |
| 19 | 7 | 0 | 5.982834 | -1.710656 | 0.862507 |
| 20 | 8 | 0 | -4.877756 | -1.446280 | 1.525376 |
| 21 | 6 | 0 | -6.019134 | 0.385374 | 0.414054 |
| 22 | 6 | 0 | -5.129981 | -1.613352 | -0.827622 |
| 23 | 6 | 0 | -4.068565 | -2.688259 | -0.995832 |
| 24 | 1 | 0 | -0.157231 | -0.462592 | 0.061399 |
| 25 | 1 | 0 | -0.319274 | 4.448161 | -0.216686 |
| 26 | 1 | 0 | 2.691559 | 4.241414 | -0.269646 |
| 27 | 1 | 0 | 4.229508 | 2.034031 | -0.191433 |
| 28 | 1 | 0 | 1.310688 | -1.354204 | -1.148576 |
| 29 | 1 | 0 | 2.489085 | -3.534090 | -0.965817 |
| 30 | 1 | 0 | 6.367931 | -0.923931 | 1.352443 |
| 31 | 1 | 0 | 6.367400 | -2.622130 | 1.032310 |
| 32 | 1 | 0 | -4.680751 | -0.869562 | 2.270152 |
| 33 | 1 | 0 | -6.054278 | 0.992872 | -0.490930 |
| 34 | 1 | 0 | -6.971602 | -0.133337 | 0.531063 |
| 35 | 1 | 0 | -5.874080 | 1.051768 | 1.265887 |
| 36 | 1 | 0 | -5.218005 | -1.021349 | -1.741269 |
| 37 | 1 | 0 | -6.105994 | -2.073635 | -0.650635 |
| 38 | 1 | 0 | -3.974561 | -3.291483 | -0.092807 |
| 39 | 1 | 0 | -3.092654 | -2.251090 | -1.214491 |
| 40 | 1 | 0 | -4.329183 | -3.352727 | -1.821290 |

---------------------------------------------------------------------

Conformer # 10
Electronic Energy = -1006.955642 Hartree

| Center Number | Atomic Number | Atomic Type | Coordinates (Angstroms) X | Y | Z |
|---|---|---|---|---|---|
| 1 | 6 | 0 | -0.282629 | 0.602997 | 0.015286 |
| 2 | 6 | 0 | 0.917747 | 1.311622 | 0.042522 |
| 3 | 7 | 0 | 1.002695 | 2.661771 | 0.094256 |
| 4 | 6 | 0 | -0.119056 | 3.359158 | 0.111936 |
| 5 | 6 | 0 | -1.379707 | 2.762731 | 0.084736 |
| 6 | 6 | 0 | -1.448948 | 1.353622 | 0.043085 |
| 7 | 6 | 0 | -2.718940 | 3.261486 | 0.079904 |
| 8 | 7 | 0 | -2.792547 | 1.006441 | 0.003333 |
| 9 | 6 | 0 | -3.537064 | 2.183322 | 0.026399 |
| 10 | 6 | 0 | 2.153004 | 0.593034 | 0.018647 |
| 11 | 6 | 0 | 3.188407 | -0.026058 | -0.005282 |
| 12 | 6 | 0 | -4.668099 | -0.469425 | -0.471798 |
| 13 | 6 | 0 | -3.346037 | -0.277344 | -0.067216 |
| 14 | 7 | 0 | -2.558200 | -1.283693 | 0.270348 |
| 15 | 6 | 0 | -3.086735 | -2.514917 | 0.209392 |
| 16 | 7 | 0 | -4.345303 | -2.817808 | -0.141702 |
| 17 | 6 | 0 | -5.103879 | -1.781288 | -0.477990 |
| 18 | 6 | 0 | 4.459809 | -0.765275 | -0.013413 |
| 19 | 7 | 0 | -2.267382 | -3.544250 | 0.513605 |
| 20 | 8 | 0 | 4.627163 | -1.447835 | 1.235827 |
| 21 | 6 | 0 | 4.414406 | -1.860443 | -1.075218 |
| 22 | 6 | 0 | 5.609109 | 0.236798 | -0.245322 |
| 23 | 6 | 0 | 6.999412 | -0.381044 | -0.256849 |
| 24 | 1 | 0 | -0.291203 | -0.473221 | -0.015823 |
| 25 | 1 | 0 | -0.022043 | 4.440257 | 0.147252 |
| 26 | 1 | 0 | -3.032891 | 4.291374 | 0.122549 |
| 27 | 1 | 0 | -4.611958 | 2.144943 | 0.034306 |
| 28 | 1 | 0 | -5.316247 | 0.331915 | -0.786955 |
| 29 | 1 | 0 | -6.122280 | -2.009602 | -0.776959 |
| 30 | 1 | 0 | -1.381320 | -3.348758 | 0.942577 |
| 31 | 1 | 0 | -2.671679 | -4.455142 | 0.633286 |
| 32 | 1 | 0 | 4.596028 | -0.789916 | 1.938057 |
| 33 | 1 | 0 | 4.318206 | -1.425586 | -2.069692 |
| 34 | 1 | 0 | 5.326319 | -2.455408 | -1.034378 |
| 35 | 1 | 0 | 3.563883 | -2.517044 | -0.895155 |
| 36 | 1 | 0 | 5.546627 | 0.992809 | 0.543243 |
| 37 | 1 | 0 | 5.416865 | 0.761171 | -1.184837 |

| | | | | | |
|---|---|---|---|---|---|
| 38 | 1 | 0 | 7.148649 | -1.037976 | -1.114874 |
| 39 | 1 | 0 | 7.179418 | -0.961045 | 0.649029 |
| 40 | 1 | 0 | 7.756682 | 0.402822 | -0.310033 |

Conformer # 11
Electronic Energy = -1006.955390 Hartree

| Center Number | Atomic Number | Atomic Type | Coordinates (Angstroms) X | Y | Z |
|---|---|---|---|---|---|
| 1 | 6 | 0 | 0.044868 | 0.392450 | -0.014574 |
| 2 | 6 | 0 | 1.291722 | 1.013194 | 0.049749 |
| 3 | 7 | 0 | 1.469843 | 2.354662 | 0.058940 |
| 4 | 6 | 0 | 0.400327 | 3.129882 | 0.025091 |
| 5 | 6 | 0 | -0.898587 | 2.625029 | -0.032757 |
| 6 | 6 | 0 | -1.064740 | 1.222795 | -0.076441 |
| 7 | 6 | 0 | -2.201212 | 3.212610 | -0.033144 |
| 8 | 7 | 0 | -2.425210 | 0.970869 | -0.109900 |
| 9 | 6 | 0 | -3.092135 | 2.191338 | -0.066940 |
| 10 | 6 | 0 | 2.472674 | 0.211877 | 0.115047 |
| 11 | 6 | 0 | 3.469743 | -0.465402 | 0.168676 |
| 12 | 6 | 0 | -2.519864 | -1.379380 | -0.782369 |
| 13 | 6 | 0 | -3.091885 | -0.261029 | -0.183279 |
| 14 | 7 | 0 | -4.312730 | -0.274439 | 0.328348 |
| 15 | 6 | 0 | -4.984256 | -1.431797 | 0.257624 |
| 16 | 7 | 0 | -4.523054 | -2.581249 | -0.261775 |
| 17 | 6 | 0 | -3.302816 | -2.521806 | -0.777152 |
| 18 | 6 | 0 | 4.695755 | -1.272594 | 0.259254 |
| 19 | 7 | 0 | -6.245871 | -1.437070 | 0.738174 |
| 20 | 8 | 0 | 4.596474 | -2.176104 | 1.367449 |
| 21 | 6 | 0 | 4.839252 | -2.162623 | -0.971334 |
| 22 | 6 | 0 | 5.919500 | -0.359269 | 0.461291 |
| 23 | 6 | 0 | 6.169634 | 0.663607 | -0.636448 |
| 24 | 1 | 0 | -0.015980 | -0.684382 | 0.005870 |
| 25 | 1 | 0 | 0.573694 | 4.201607 | 0.050181 |
| 26 | 1 | 0 | -2.442392 | 4.262635 | -0.014246 |
| 27 | 1 | 0 | -4.167699 | 2.197806 | -0.080204 |
| 28 | 1 | 0 | -1.556807 | -1.362568 | -1.264136 |
| 29 | 1 | 0 | -2.924641 | -3.436893 | -1.222326 |
| 30 | 1 | 0 | -6.560467 | -0.653883 | 1.281244 |

| | | | | | |
|---|---|---|---|---|---|
| 31 | 1 | 0 | -6.710337 | -2.320095 | 0.847972 |
| 32 | 1 | 0 | 4.486477 | -1.651161 | 2.167062 |
| 33 | 1 | 0 | 4.877269 | -1.569546 | -1.883325 |
| 34 | 1 | 0 | 5.756831 | -2.747056 | -0.888303 |
| 35 | 1 | 0 | 3.991968 | -2.844606 | -1.035653 |
| 36 | 1 | 0 | 6.783449 | -1.019316 | 0.574225 |
| 37 | 1 | 0 | 5.788155 | 0.158146 | 1.416704 |
| 38 | 1 | 0 | 5.304943 | 1.314734 | -0.775062 |
| 39 | 1 | 0 | 6.392616 | 0.187775 | -1.592617 |
| 40 | 1 | 0 | 7.022415 | 1.292939 | -0.377238 |

---------------------------------------------------------------------

Conformer # 12
Electronic Energy = -1006.955387 Hartree

| Center Number | Atomic Number | Atomic Type | Coordinates (Angstroms) X | Y | Z |
|---|---|---|---|---|---|
| 1 | 6 | 0 | -0.049947 | 0.406440 | -0.147840 |
| 2 | 6 | 0 | -1.293285 | 1.037512 | -0.159187 |
| 3 | 7 | 0 | -1.460351 | 2.380265 | -0.181418 |
| 4 | 6 | 0 | -0.383977 | 3.146601 | -0.172418 |
| 5 | 6 | 0 | 0.911976 | 2.631007 | -0.157794 |
| 6 | 6 | 0 | 1.068370 | 1.227021 | -0.169213 |
| 7 | 6 | 0 | 2.217860 | 3.208691 | -0.104022 |
| 8 | 7 | 0 | 2.426674 | 0.964319 | -0.130258 |
| 9 | 6 | 0 | 3.100775 | 2.180288 | -0.075579 |
| 10 | 6 | 0 | -2.482069 | 0.245303 | -0.143125 |
| 11 | 6 | 0 | -3.484046 | -0.426702 | -0.127179 |
| 12 | 6 | 0 | 2.534984 | -1.401035 | -0.745689 |
| 13 | 6 | 0 | 3.085558 | -0.273893 | -0.143023 |
| 14 | 7 | 0 | 4.278522 | -0.285018 | 0.430537 |
| 15 | 6 | 0 | 4.942367 | -1.448977 | 0.420547 |
| 16 | 7 | 0 | 4.497549 | -2.606547 | -0.095118 |
| 17 | 6 | 0 | 3.305927 | -2.549360 | -0.673921 |
| 18 | 6 | 0 | -4.718461 | -1.226102 | -0.133527 |
| 19 | 7 | 0 | 6.177936 | -1.452970 | 0.964502 |
| 20 | 8 | 0 | -4.834246 | -1.921469 | -1.381925 |
| 21 | 6 | 0 | -4.650826 | -2.318645 | 0.928585 |
| 22 | 6 | 0 | -5.945095 | -0.312823 | 0.048535 |
| 23 | 6 | 0 | -5.982195 | 0.497563 | 1.335325 |

| | | | | | |
|---|---|---|---|---|---|
| 24 | 1 | 0 | 0.000708 | -0.670229 | -0.104292 |
| 25 | 1 | 0 | -0.549223 | 4.219909 | -0.172163 |
| 26 | 1 | 0 | 2.466637 | 4.257038 | -0.091665 |
| 27 | 1 | 0 | 4.175683 | 2.178521 | -0.035231 |
| 28 | 1 | 0 | 1.598147 | -1.387725 | -1.276746 |
| 29 | 1 | 0 | 2.942556 | -3.471535 | -1.116778 |
| 30 | 1 | 0 | 6.472416 | -0.659487 | 1.503876 |
| 31 | 1 | 0 | 6.628686 | -2.336665 | 1.117472 |
| 32 | 1 | 0 | -4.866368 | -1.263135 | -2.083820 |
| 33 | 1 | 0 | -4.514007 | -1.893975 | 1.921475 |
| 34 | 1 | 0 | -5.577113 | -2.894932 | 0.912677 |
| 35 | 1 | 0 | -3.816494 | -2.987242 | 0.718860 |
| 36 | 1 | 0 | -6.824739 | -0.957680 | -0.023491 |
| 37 | 1 | 0 | -5.977814 | 0.366016 | -0.809243 |
| 38 | 1 | 0 | -5.097927 | 1.129991 | 1.430640 |
| 39 | 1 | 0 | -6.038313 | -0.142089 | 2.217445 |
| 40 | 1 | 0 | -6.858678 | 1.147300 | 1.345827 |

---------------------------------------------------------------------

Conformer # 13
Electronic Energy = -1006.955344 Hartree

| Center Number | Atomic Number | Atomic Type | Coordinates (Angstroms) X | Y | Z |
|---|---|---|---|---|---|
| 1 | 6 | 0 | 0.269221 | 0.708170 | 0.032627 |
| 2 | 6 | 0 | -0.870220 | 1.511356 | 0.056741 |
| 3 | 7 | 0 | -0.846565 | 2.864978 | 0.062414 |
| 4 | 6 | 0 | 0.327340 | 3.470129 | 0.034898 |
| 5 | 6 | 0 | 1.535719 | 2.773941 | 0.005358 |
| 6 | 6 | 0 | 1.491832 | 1.363232 | 0.012335 |
| 7 | 6 | 0 | 2.909951 | 3.163342 | -0.042651 |
| 8 | 7 | 0 | 2.802373 | 0.908524 | -0.040757 |
| 9 | 6 | 0 | 3.638489 | 2.021858 | -0.073606 |
| 10 | 6 | 0 | -2.158527 | 0.892974 | 0.080690 |
| 11 | 6 | 0 | -3.240505 | 0.359156 | 0.098776 |
| 12 | 6 | 0 | 4.540796 | -0.729334 | -0.507007 |
| 13 | 6 | 0 | 3.250376 | -0.417239 | -0.076030 |
| 14 | 7 | 0 | 2.395461 | -1.344032 | 0.320780 |
| 15 | 6 | 0 | 2.823087 | -2.615115 | 0.294277 |

| | | | | | |
|---|---|---|---|---|---|
| 16 | 7 | 0 | 4.042975 | -3.030582 | -0.077610 |
| 17 | 6 | 0 | 4.871034 | -2.071166 | -0.473606 |
| 18 | 6 | 0 | -4.565135 | -0.280006 | 0.135804 |
| 19 | 7 | 0 | 1.934007 | -3.563534 | 0.659877 |
| 20 | 8 | 0 | -4.780073 | -0.866744 | 1.425503 |
| 21 | 6 | 0 | -5.642064 | 0.767556 | -0.154108 |
| 22 | 6 | 0 | -4.584695 | -1.443612 | -0.869811 |
| 23 | 6 | 0 | -5.891271 | -2.222488 | -0.922838 |
| 24 | 1 | 0 | 0.191101 | -0.365739 | 0.039515 |
| 25 | 1 | 0 | 0.317614 | 4.556133 | 0.033920 |
| 26 | 1 | 0 | 3.305667 | 4.165581 | -0.041905 |
| 27 | 1 | 0 | 4.706892 | 1.897488 | -0.084193 |
| 28 | 1 | 0 | 5.240649 | 0.005467 | -0.870271 |
| 29 | 1 | 0 | 5.858971 | -2.391201 | -0.790220 |
| 30 | 1 | 0 | 1.080282 | -3.281345 | 1.105862 |
| 31 | 1 | 0 | 2.269013 | -4.498473 | 0.805632 |
| 32 | 1 | 0 | -4.755436 | -0.160795 | 2.079162 |
| 33 | 1 | 0 | -5.521283 | 1.183029 | -1.155340 |
| 34 | 1 | 0 | -6.630966 | 0.317799 | -0.072540 |
| 35 | 1 | 0 | -5.573074 | 1.587268 | 0.562943 |
| 36 | 1 | 0 | -3.766385 | -2.114300 | -0.598035 |
| 37 | 1 | 0 | -4.344536 | -1.037257 | -1.854757 |
| 38 | 1 | 0 | -6.716457 | -1.617343 | -1.300973 |
| 39 | 1 | 0 | -6.167515 | -2.596732 | 0.063011 |
| 40 | 1 | 0 | -5.784278 | -3.079634 | -1.589856 |

---------------------------------------------------------------------

Conformer # 14
Electronic Energy = -1006.955200 Hartree

| Center Number | Atomic Number | Atomic Type | Coordinates (Angstroms) X | Y | Z |
|---|---|---|---|---|---|
| 1 | 6 | 0 | 0.103290 | 0.580034 | -0.142726 |
| 2 | 6 | 0 | 1.302652 | 1.289830 | -0.098462 |
| 3 | 7 | 0 | 1.381178 | 2.636757 | 0.005512 |
| 4 | 6 | 0 | 0.257242 | 3.326172 | 0.092131 |
| 5 | 6 | 0 | -1.002299 | 2.727115 | 0.064875 |
| 6 | 6 | 0 | -1.066127 | 1.323409 | -0.080622 |
| 7 | 6 | 0 | -2.342998 | 3.208112 | 0.180532 |
| 8 | 7 | 0 | -2.403452 | 0.967861 | -0.061056 |

| | | | | | |
|---|---|---|---|---|---|
| 9 | 6 | 0 | -3.155790 | 2.124879 | 0.113911 |
| 10 | 6 | 0 | 2.538941 | 0.576156 | -0.159544 |
| 11 | 6 | 0 | 3.575496 | -0.039489 | -0.209746 |
| 12 | 6 | 0 | -2.358343 | -1.322117 | -0.917431 |
| 13 | 6 | 0 | -2.977013 | -0.304968 | -0.196774 |
| 14 | 7 | 0 | -4.158278 | -0.458702 | 0.380582 |
| 15 | 6 | 0 | -4.740027 | -1.659257 | 0.253444 |
| 16 | 7 | 0 | -4.223790 | -2.722853 | -0.383741 |
| 17 | 6 | 0 | -3.047096 | -2.522650 | -0.961414 |
| 18 | 6 | 0 | 4.846299 | -0.775067 | -0.290467 |
| 19 | 7 | 0 | -5.964342 | -1.806605 | 0.802942 |
| 20 | 8 | 0 | 5.074354 | -1.192837 | -1.643189 |
| 21 | 6 | 0 | 5.996444 | 0.118125 | 0.176173 |
| 22 | 6 | 0 | 4.766971 | -2.085256 | 0.507073 |
| 23 | 6 | 0 | 4.497934 | -1.930582 | 1.995006 |
| 24 | 1 | 0 | 0.122744 | -0.496891 | -0.201646 |
| 25 | 1 | 0 | 0.352338 | 4.403304 | 0.192620 |
| 26 | 1 | 0 | -2.660567 | 4.231387 | 0.295110 |
| 27 | 1 | 0 | -4.227835 | 2.047538 | 0.158211 |
| 28 | 1 | 0 | -1.432061 | -1.188039 | -1.450307 |
| 29 | 1 | 0 | -2.627452 | -3.366041 | -1.501003 |
| 30 | 1 | 0 | -6.304410 | -1.096601 | 1.425456 |
| 31 | 1 | 0 | -6.350116 | -2.731098 | 0.866274 |
| 32 | 1 | 0 | 5.108681 | -0.404176 | -2.194125 |
| 33 | 1 | 0 | 5.846280 | 0.460480 | 1.199776 |
| 34 | 1 | 0 | 6.932323 | -0.439797 | 0.120462 |
| 35 | 1 | 0 | 6.073053 | 0.999294 | -0.463145 |
| 36 | 1 | 0 | 5.714021 | -2.605906 | 0.342563 |
| 37 | 1 | 0 | 3.986235 | -2.694124 | 0.045876 |
| 38 | 1 | 0 | 3.568551 | -1.387671 | 2.176855 |
| 39 | 1 | 0 | 5.303766 | -1.399170 | 2.503615 |
| 40 | 1 | 0 | 4.406944 | -2.911653 | 2.463487 |

---------------------------------------------------------------------

Conformer # 15
Electronic Energy = -1006.955168 Hartree

| Center Number | Atomic Number | Atomic Type | Coordinates (Angstroms) X | Y | Z |
|---|---|---|---|---|---|
| 1 | 6 | 0 | 0.081048 | 0.525153 | -0.039687 |

| | | | | | |
|---|---|---|---|---|---|
| 2 | 6 | 0 | 1.286133 | 1.224324 | -0.096623 |
| 3 | 7 | 0 | 1.382887 | 2.572258 | -0.028699 |
| 4 | 6 | 0 | 0.269994 | 3.276578 | 0.078554 |
| 5 | 6 | 0 | -0.994193 | 2.690050 | 0.137732 |
| 6 | 6 | 0 | -1.074195 | 1.280162 | 0.100089 |
| 7 | 6 | 0 | -2.329590 | 3.194435 | 0.205558 |
| 8 | 7 | 0 | -2.415448 | 0.942537 | 0.151506 |
| 9 | 6 | 0 | -3.155877 | 2.119577 | 0.198851 |
| 10 | 6 | 0 | 2.508986 | 0.499235 | -0.239393 |
| 11 | 6 | 0 | 3.535305 | -0.123723 | -0.359847 |
| 12 | 6 | 0 | -2.345070 | -1.445003 | 0.679002 |
| 13 | 6 | 0 | -3.002804 | -0.331061 | 0.165702 |
| 14 | 7 | 0 | -4.235251 | -0.391300 | -0.313853 |
| 15 | 6 | 0 | -4.830967 | -1.591420 | -0.298499 |
| 16 | 7 | 0 | -4.283847 | -2.738628 | 0.135413 |
| 17 | 6 | 0 | -3.054920 | -2.632785 | 0.621947 |
| 18 | 6 | 0 | 4.787531 | -0.874353 | -0.536026 |
| 19 | 7 | 0 | -6.103563 | -1.648086 | -0.745740 |
| 20 | 8 | 0 | 4.682177 | -1.716881 | -1.691233 |
| 21 | 6 | 0 | 5.956994 | 0.096232 | -0.707039 |
| 22 | 6 | 0 | 5.007716 | -1.848602 | 0.630891 |
| 23 | 6 | 0 | 5.132912 | -1.207724 | 2.003491 |
| 24 | 1 | 0 | 0.084113 | -0.550181 | -0.124517 |
| 25 | 1 | 0 | 0.377839 | 4.356596 | 0.114108 |
| 26 | 1 | 0 | -2.634455 | 4.226697 | 0.256343 |
| 27 | 1 | 0 | -4.229119 | 2.058531 | 0.238051 |
| 28 | 1 | 0 | -1.370939 | -1.396285 | 1.135623 |
| 29 | 1 | 0 | -2.607277 | -3.547296 | 0.998614 |
| 30 | 1 | 0 | -6.482037 | -0.854983 | -1.230544 |
| 31 | 1 | 0 | -6.514697 | -2.550752 | -0.899652 |
| 32 | 1 | 0 | 4.513059 | -1.153324 | -2.453135 |
| 33 | 1 | 0 | 6.881482 | -0.470206 | -0.828178 |
| 34 | 1 | 0 | 6.054800 | 0.758009 | 0.153121 |
| 35 | 1 | 0 | 5.806049 | 0.717234 | -1.591573 |
| 36 | 1 | 0 | 5.909231 | -2.419140 | 0.392446 |
| 37 | 1 | 0 | 4.174126 | -2.554214 | 0.620754 |
| 38 | 1 | 0 | 5.225807 | -1.978426 | 2.770075 |
| 39 | 1 | 0 | 6.012378 | -0.566186 | 2.077031 |
| 40 | 1 | 0 | 4.254756 | -0.604949 | 2.242695 |

---------------------------------------------------------------------

Conformer # 16

Electronic Energy = -1006.954905 Hartree

| Center Number | Atomic Number | Atomic Type | Coordinates (Angstroms) X | Y | Z |
|---|---|---|---|---|---|
| 1 | 6 | 0 | 0.035753 | 0.499970 | -0.068440 |
| 2 | 6 | 0 | -1.176026 | 1.189049 | -0.035942 |
| 3 | 7 | 0 | -1.279132 | 2.538240 | -0.027878 |
| 4 | 6 | 0 | -0.167535 | 3.252662 | -0.029200 |
| 5 | 6 | 0 | 1.102504 | 2.676359 | -0.055837 |
| 6 | 6 | 0 | 1.190952 | 1.267206 | -0.100842 |
| 7 | 6 | 0 | 2.435798 | 3.189522 | -0.020984 |
| 8 | 7 | 0 | 2.535452 | 0.939020 | -0.100748 |
| 9 | 6 | 0 | 3.268572 | 2.119698 | -0.036058 |
| 10 | 6 | 0 | -2.398970 | 0.450801 | -0.004540 |
| 11 | 6 | 0 | -3.423639 | -0.185574 | 0.025460 |
| 12 | 6 | 0 | 2.511420 | -1.409473 | -0.783130 |
| 13 | 6 | 0 | 3.131720 | -0.329014 | -0.162848 |
| 14 | 7 | 0 | 4.335507 | -0.415990 | 0.381024 |
| 15 | 6 | 0 | 4.939223 | -1.610838 | 0.322388 |
| 16 | 7 | 0 | 4.425138 | -2.729323 | -0.214634 |
| 17 | 6 | 0 | 3.224925 | -2.596237 | -0.762403 |
| 18 | 6 | 0 | -4.681745 | -0.947159 | 0.043756 |
| 19 | 7 | 0 | 6.184994 | -1.692222 | 0.836308 |
| 20 | 8 | 0 | -4.832736 | -1.656555 | -1.192206 |
| 21 | 6 | 0 | -4.619146 | -2.020487 | 1.126906 |
| 22 | 6 | 0 | -5.849449 | 0.038339 | 0.253384 |
| 23 | 6 | 0 | -7.228067 | -0.604838 | 0.274786 |
| 24 | 1 | 0 | 0.036514 | -0.578554 | -0.047348 |
| 25 | 1 | 0 | -0.281563 | 4.332330 | -0.003614 |
| 26 | 1 | 0 | 2.735260 | 4.224201 | 0.008074 |
| 27 | 1 | 0 | 4.342833 | 2.065682 | -0.022097 |
| 28 | 1 | 0 | 1.564412 | -1.334019 | -1.290433 |
| 29 | 1 | 0 | 2.805598 | -3.485951 | -1.221918 |
| 30 | 1 | 0 | 6.530539 | -0.930751 | 1.391316 |
| 31 | 1 | 0 | 6.593462 | -2.601489 | 0.954185 |
| 32 | 1 | 0 | -4.816532 | -1.011640 | -1.906886 |
| 33 | 1 | 0 | -4.533095 | -1.564694 | 2.112893 |
| 34 | 1 | 0 | -5.520197 | -2.632348 | 1.095782 |

| | | | | | |
|---|---|---|---|---|---|
| 35 | 1 | 0 | -3.756663 | -2.665217 | 0.961384 |
| 36 | 1 | 0 | -5.799170 | 0.779303 | -0.550220 |
| 37 | 1 | 0 | -5.668884 | 0.584966 | 1.182444 |
| 38 | 1 | 0 | -7.366936 | -1.246977 | 1.145621 |
| 39 | 1 | 0 | -7.395373 | -1.206172 | -0.619547 |
| 40 | 1 | 0 | -7.999780 | 0.165791 | 0.310776 |

---------------------------------------------------------------------

Conformer # 17
Electronic Energy = -1006.954870 Hartree

| Center Number | Atomic Number | Atomic Type | Coordinates (Angstroms) X | Y | Z |
|---|---|---|---|---|---|
| 1 | 6 | 0 | -0.036302 | 0.494478 | -0.089918 |
| 2 | 6 | 0 | 1.176480 | 1.182129 | -0.066367 |
| 3 | 7 | 0 | 1.281605 | 2.531140 | -0.067392 |
| 4 | 6 | 0 | 0.171038 | 3.247194 | -0.070912 |
| 5 | 6 | 0 | -1.099847 | 2.672584 | -0.086475 |
| 6 | 6 | 0 | -1.190618 | 1.263232 | -0.120040 |
| 7 | 6 | 0 | -2.432130 | 3.188176 | -0.048256 |
| 8 | 7 | 0 | -2.535661 | 0.937261 | -0.109911 |
| 9 | 6 | 0 | -3.266604 | 2.119661 | -0.050358 |
| 10 | 6 | 0 | 2.399199 | 0.443534 | -0.032741 |
| 11 | 6 | 0 | 3.425519 | -0.190178 | -0.003694 |
| 12 | 6 | 0 | -2.520445 | -1.417255 | -0.772074 |
| 13 | 6 | 0 | -3.134694 | -0.330055 | -0.157583 |
| 14 | 7 | 0 | -4.335419 | -0.409745 | 0.394178 |
| 15 | 6 | 0 | -4.942013 | -1.603708 | 0.349622 |
| 16 | 7 | 0 | -4.433494 | -2.727992 | -0.180552 |
| 17 | 6 | 0 | -3.236315 | -2.602276 | -0.736609 |
| 18 | 6 | 0 | 4.684096 | -0.948788 | 0.059568 |
| 19 | 7 | 0 | -6.184963 | -1.677843 | 0.871463 |
| 20 | 8 | 0 | 4.629535 | -1.879552 | 1.147523 |
| 21 | 6 | 0 | 4.841786 | -1.798337 | -1.198372 |
| 22 | 6 | 0 | 5.846422 | 0.047412 | 0.249912 |
| 23 | 6 | 0 | 7.222374 | -0.592667 | 0.358755 |
| 24 | 1 | 0 | -0.038339 | -0.583926 | -0.063631 |
| 25 | 1 | 0 | 0.286802 | 4.326862 | -0.055038 |
| 26 | 1 | 0 | -2.729748 | 4.223549 | -0.025658 |
| 27 | 1 | 0 | -4.340840 | 2.067419 | -0.030022 |

| | | | | | |
|---|---|---|---|---|---|
| 28 | 1 | 0 | -1.576388 | -1.348287 | -1.285740 |
| 29 | 1 | 0 | -2.821592 | -3.496902 | -1.190732 |
| 30 | 1 | 0 | -6.525494 | -0.910829 | 1.421926 |
| 31 | 1 | 0 | -6.594505 | -2.585187 | 0.999968 |
| 32 | 1 | 0 | 4.473725 | -1.380523 | 1.956035 |
| 33 | 1 | 0 | 4.920581 | -1.164607 | -2.081248 |
| 34 | 1 | 0 | 5.738743 | -2.411988 | -1.120324 |
| 35 | 1 | 0 | 3.980707 | -2.455802 | -1.313447 |
| 36 | 1 | 0 | 5.635279 | 0.626188 | 1.154223 |
| 37 | 1 | 0 | 5.820930 | 0.759201 | -0.579054 |
| 38 | 1 | 0 | 7.530520 | -1.063401 | -0.575799 |
| 39 | 1 | 0 | 7.242655 | -1.350623 | 1.142669 |
| 40 | 1 | 0 | 7.968532 | 0.164696 | 0.604429 |

---------------------------------------------------------------------

Conformer # 18
Electronic Energy = -1006.954635 Hartree

| Center Number | Atomic Number | Atomic Type | Coordinates (Angstroms) X | Y | Z |
|---|---|---|---|---|---|
| 1 | 6 | 0 | 0.003938 | 0.622458 | 0.083588 |
| 2 | 6 | 0 | -1.164442 | 1.382146 | 0.037309 |
| 3 | 7 | 0 | -1.186180 | 2.733414 | -0.032056 |
| 4 | 6 | 0 | -0.033590 | 3.377752 | -0.080313 |
| 5 | 6 | 0 | 1.199568 | 2.726698 | -0.045954 |
| 6 | 6 | 0 | 1.203755 | 1.318239 | 0.062913 |
| 7 | 6 | 0 | 2.560604 | 3.155550 | -0.123850 |
| 8 | 7 | 0 | 2.525982 | 0.909107 | 0.058293 |
| 9 | 6 | 0 | 3.327650 | 2.038849 | -0.071422 |
| 10 | 6 | 0 | -2.429969 | 0.719319 | 0.060713 |
| 11 | 6 | 0 | -3.493042 | 0.148949 | 0.081669 |
| 12 | 6 | 0 | 2.372357 | -1.401266 | 0.845417 |
| 13 | 6 | 0 | 3.046188 | -0.388710 | 0.169134 |
| 14 | 7 | 0 | 4.233912 | -0.571866 | -0.385989 |
| 15 | 6 | 0 | 4.766486 | -1.797091 | -0.280874 |
| 16 | 7 | 0 | 4.195376 | -2.857454 | 0.313436 |
| 17 | 6 | 0 | 3.013711 | -2.628275 | 0.869900 |
| 18 | 6 | 0 | -4.795883 | -0.533231 | 0.125609 |
| 19 | 7 | 0 | 5.996959 | -1.976025 | -0.806872 |
| 20 | 8 | 0 | -4.973871 | -1.145714 | 1.408498 |

| | | | | | |
|---|---|---|---|---|---|
| 21 | 6 | 0 | -5.909025 | 0.483976 | -0.134238 |
| 22 | 6 | 0 | -4.792730 | -1.681799 | -0.897591 |
| 23 | 6 | 0 | -6.076373 | -2.498115 | -0.946365 |
| 24 | 1 | 0 | -0.061135 | -0.453867 | 0.112974 |
| 25 | 1 | 0 | -0.082588 | 4.460142 | -0.153962 |
| 26 | 1 | 0 | 2.921208 | 4.167734 | -0.204825 |
| 27 | 1 | 0 | 4.396328 | 1.919077 | -0.098316 |
| 28 | 1 | 0 | 1.439405 | -1.246649 | 1.360717 |
| 29 | 1 | 0 | 2.549630 | -3.469725 | 1.375039 |
| 30 | 1 | 0 | 6.377987 | -1.262375 | -1.400853 |
| 31 | 1 | 0 | 6.348187 | -2.912690 | -0.889436 |
| 32 | 1 | 0 | -4.968822 | -0.448993 | 2.072372 |
| 33 | 1 | 0 | -5.815569 | 0.916513 | -1.131103 |
| 34 | 1 | 0 | -6.882001 | 0.002092 | -0.045226 |
| 35 | 1 | 0 | -5.855469 | 1.295544 | 0.593278 |
| 36 | 1 | 0 | -3.951602 | -2.331638 | -0.645612 |
| 37 | 1 | 0 | -4.577092 | -1.253866 | -1.879042 |
| 38 | 1 | 0 | -6.922915 | -1.912844 | -1.308139 |
| 39 | 1 | 0 | -6.330517 | -2.892112 | 0.037683 |
| 40 | 1 | 0 | -5.951632 | -3.343560 | -1.625081 |

---------------------------------------------------------------------

Conformer # 19
Electronic Energy = -1006.954614 Hartree

| Center Number | Atomic Number | Atomic Type | Coordinates (Angstroms) X | Y | Z |
|---|---|---|---|---|---|
| 1 | 6 | 0 | -0.164271 | 0.628925 | -0.102035 |
| 2 | 6 | 0 | 1.027065 | 1.351421 | -0.151086 |
| 3 | 7 | 0 | 1.098775 | 2.702698 | -0.152508 |
| 4 | 6 | 0 | -0.029273 | 3.388600 | -0.097037 |
| 5 | 6 | 0 | -1.282016 | 2.778651 | -0.042925 |
| 6 | 6 | 0 | -1.337095 | 1.368172 | -0.053012 |
| 7 | 6 | 0 | -2.624643 | 3.263099 | 0.033781 |
| 8 | 7 | 0 | -2.675235 | 1.006024 | 0.026612 |
| 9 | 6 | 0 | -3.430487 | 2.175676 | 0.077858 |
| 10 | 6 | 0 | 2.269127 | 0.646627 | -0.209582 |
| 11 | 6 | 0 | 3.309365 | 0.039625 | -0.258702 |
| 12 | 6 | 0 | -4.521304 | -0.504882 | 0.509043 |
| 13 | 6 | 0 | -3.214161 | -0.284807 | 0.070763 |

| | | | | | |
|---|---|---|---|---|---|
| 14 | 7 | 0 | -2.427636 | -1.271085 | -0.324132 |
| 15 | 6 | 0 | -2.941510 | -2.509463 | -0.287724 |
| 16 | 7 | 0 | -4.185490 | -2.837721 | 0.091568 |
| 17 | 6 | 0 | -4.943385 | -1.820922 | 0.484999 |
| 18 | 6 | 0 | 4.600528 | -0.651617 | -0.326297 |
| 19 | 7 | 0 | -2.121715 | -3.518710 | -0.651109 |
| 20 | 8 | 0 | 4.304266 | -2.028210 | -0.597018 |
| 21 | 6 | 0 | 5.433338 | -0.064633 | -1.470168 |
| 22 | 6 | 0 | 5.357699 | -0.513098 | 1.009210 |
| 23 | 6 | 0 | 4.634590 | -1.093314 | 2.213208 |
| 24 | 1 | 0 | -0.161681 | -0.447691 | -0.112547 |
| 25 | 1 | 0 | 0.056968 | 4.471175 | -0.093524 |
| 26 | 1 | 0 | -2.948960 | 4.290640 | 0.043049 |
| 27 | 1 | 0 | -4.504435 | 2.126997 | 0.110607 |
| 28 | 1 | 0 | -5.167879 | 0.278052 | 0.870246 |
| 29 | 1 | 0 | -5.949795 | -2.069917 | 0.807243 |
| 30 | 1 | 0 | -1.248490 | -3.300094 | -1.094893 |
| 31 | 1 | 0 | -2.518356 | -4.430896 | -0.784963 |
| 32 | 1 | 0 | 5.146238 | -2.490087 | -0.668519 |
| 33 | 1 | 0 | 6.393882 | -0.582365 | -1.527977 |
| 34 | 1 | 0 | 5.624619 | 0.996682 | -1.309858 |
| 35 | 1 | 0 | 4.909257 | -0.188080 | -2.417353 |
| 36 | 1 | 0 | 5.572427 | 0.546373 | 1.169558 |
| 37 | 1 | 0 | 6.325868 | -1.008646 | 0.877782 |
| 38 | 1 | 0 | 5.251631 | -1.003870 | 3.108720 |
| 39 | 1 | 0 | 3.695192 | -0.570184 | 2.400653 |
| 40 | 1 | 0 | 4.407093 | -2.149215 | 2.063150 |

---------------------------------------------------------------------

Conformer # 20
Electronic Energy = -1006.954577 Hartree

| Center Number | Atomic Number | Atomic Type | Coordinates (Angstroms) X | Y | Z |
|---|---|---|---|---|---|
| 1 | 6 | 0 | 0.006568 | 0.609589 | -0.032469 |
| 2 | 6 | 0 | -1.161326 | 1.370869 | -0.002625 |
| 3 | 7 | 0 | -1.185550 | 2.722265 | -0.066079 |
| 4 | 6 | 0 | -0.034754 | 3.367348 | -0.141277 |
| 5 | 6 | 0 | 1.197970 | 2.715371 | -0.171346 |

| | | | | | |
|---|---|---|---|---|---|
| 6 | 6 | 0 | 1.203089 | 1.303124 | -0.138902 |
| 7 | 6 | 0 | 2.559315 | 3.149278 | -0.203341 |
| 8 | 7 | 0 | 2.525853 | 0.895787 | -0.158408 |
| 9 | 6 | 0 | 3.327833 | 2.032649 | -0.181519 |
| 10 | 6 | 0 | -2.423022 | 0.709323 | 0.106835 |
| 11 | 6 | 0 | -3.481515 | 0.137945 | 0.201471 |
| 12 | 6 | 0 | 2.342734 | -1.482576 | -0.699737 |
| 13 | 6 | 0 | 3.046194 | -0.406627 | -0.166310 |
| 14 | 7 | 0 | 4.263149 | -0.533054 | 0.339299 |
| 15 | 6 | 0 | 4.796183 | -1.762400 | 0.329318 |
| 16 | 7 | 0 | 4.200069 | -2.877464 | -0.123439 |
| 17 | 6 | 0 | 2.988810 | -2.705715 | -0.635061 |
| 18 | 6 | 0 | -4.774966 | -0.549747 | 0.338934 |
| 19 | 7 | 0 | 6.054236 | -1.887152 | 0.803227 |
| 20 | 8 | 0 | -4.763629 | -1.360197 | 1.520001 |
| 21 | 6 | 0 | -5.895090 | 0.489531 | 0.421146 |
| 22 | 6 | 0 | -4.953681 | -1.519633 | -0.841517 |
| 23 | 6 | 0 | -6.242426 | -2.329195 | -0.814635 |
| 24 | 1 | 0 | -0.055479 | -0.464323 | 0.046965 |
| 25 | 1 | 0 | -0.084730 | 4.451695 | -0.174042 |
| 26 | 1 | 0 | 2.918945 | 4.164338 | -0.241311 |
| 27 | 1 | 0 | 4.397031 | 1.915719 | -0.194922 |
| 28 | 1 | 0 | 1.382437 | -1.381406 | -1.176548 |
| 29 | 1 | 0 | 2.502789 | -3.593949 | -1.026708 |
| 30 | 1 | 0 | 6.462694 | -1.117253 | 1.300925 |
| 31 | 1 | 0 | 6.414686 | -2.810662 | 0.960442 |
| 32 | 1 | 0 | -4.636097 | -0.779018 | 2.276381 |
| 33 | 1 | 0 | -6.851774 | -0.006846 | 0.579757 |
| 34 | 1 | 0 | -5.947011 | 1.078951 | -0.495083 |
| 35 | 1 | 0 | -5.716016 | 1.172251 | 1.253356 |
| 36 | 1 | 0 | -4.094631 | -2.194409 | -0.833908 |
| 37 | 1 | 0 | -4.889153 | -0.938237 | -1.763834 |
| 38 | 1 | 0 | -6.243277 | -3.050621 | -1.633487 |
| 39 | 1 | 0 | -6.341235 | -2.881656 | 0.119838 |
| 40 | 1 | 0 | -7.125931 | -1.700479 | -0.933833 |

---------------------------------------------------------------------

Conformer # 21
Electronic Energy = -1006.954046 Hartree
Center Atomic Atomic Coordinates (Angstroms)

| Number | Number | Type | X | Y | Z |
|---|---|---|---|---|---|
| 1 | 6 | 0 | 0.194502 | 0.512569 | 0.030773 |
| 2 | 6 | 0 | -1.050475 | 1.139868 | 0.034523 |
| 3 | 7 | 0 | -1.227844 | 2.480867 | -0.005494 |
| 4 | 6 | 0 | -0.155571 | 3.252312 | -0.040871 |
| 5 | 6 | 0 | 1.142059 | 2.741857 | -0.044874 |
| 6 | 6 | 0 | 1.306808 | 1.340219 | -0.016557 |
| 7 | 6 | 0 | 2.444685 | 3.329925 | -0.064394 |
| 8 | 7 | 0 | 2.671475 | 1.084342 | -0.009282 |
| 9 | 6 | 0 | 3.334482 | 2.309452 | -0.038226 |
| 10 | 6 | 0 | -2.234907 | 0.341464 | 0.080676 |
| 11 | 6 | 0 | -3.228213 | -0.339967 | 0.124053 |
| 12 | 6 | 0 | 4.656573 | -0.266567 | 0.390609 |
| 13 | 6 | 0 | 3.311317 | -0.159702 | 0.032838 |
| 14 | 7 | 0 | 2.580739 | -1.214287 | -0.285179 |
| 15 | 6 | 0 | 3.190456 | -2.408469 | -0.251417 |
| 16 | 7 | 0 | 4.477561 | -2.629701 | 0.054413 |
| 17 | 6 | 0 | 5.177254 | -1.546800 | 0.372047 |
| 18 | 6 | 0 | -4.458659 | -1.133863 | 0.189587 |
| 19 | 7 | 0 | 2.430006 | -3.487357 | -0.535454 |
| 20 | 8 | 0 | -4.088182 | -2.467957 | -0.187734 |
| 21 | 6 | 0 | -4.999742 | -1.136738 | 1.622144 |
| 22 | 6 | 0 | -5.493302 | -0.600209 | -0.820695 |
| 23 | 6 | 0 | -5.945535 | 0.837139 | -0.611887 |
| 24 | 1 | 0 | 0.275225 | -0.560804 | 0.054381 |
| 25 | 1 | 0 | -0.325757 | 4.324576 | -0.065744 |
| 26 | 1 | 0 | 2.687485 | 4.378979 | -0.104286 |
| 27 | 1 | 0 | 4.409020 | 2.344607 | -0.068496 |
| 28 | 1 | 0 | 5.262309 | 0.573532 | 0.688568 |
| 29 | 1 | 0 | 6.218227 | -1.708953 | 0.634296 |
| 30 | 1 | 0 | 1.517440 | -3.348110 | -0.929356 |
| 31 | 1 | 0 | 2.887594 | -4.369780 | -0.674481 |
| 32 | 1 | 0 | -4.885111 | -3.006237 | -0.137578 |
| 33 | 1 | 0 | -5.926772 | -1.714021 | 1.663677 |
| 34 | 1 | 0 | -5.207378 | -0.126718 | 1.973892 |
| 35 | 1 | 0 | -4.270316 | -1.593743 | 2.290027 |
| 36 | 1 | 0 | -6.357110 | -1.272164 | -0.773447 |
| 37 | 1 | 0 | -5.056555 | -0.715696 | -1.814996 |
| 38 | 1 | 0 | -6.626403 | 1.137014 | -1.409999 |

| | | | | | |
|---|---|---|---|---|---|
| 39 | 1 | 0 | -6.473357 | 0.965741 | 0.334279 |
| 40 | 1 | 0 | -5.097025 | 1.523444 | -0.621862 |

---------------------------------------------------------------------

Conformer # 22
Electronic Energy = -1006.953887 Hartree

| Center Number | Atomic Number | Atomic Type | Coordinates (Angstroms) X | Y | Z |
|---|---|---|---|---|---|
| 1 | 6 | 0 | 0.084171 | 0.523593 | -0.157431 |
| 2 | 6 | 0 | 1.294675 | 1.215243 | -0.138406 |
| 3 | 7 | 0 | 1.396639 | 2.562233 | -0.070147 |
| 4 | 6 | 0 | 0.284227 | 3.272405 | 0.002802 |
| 5 | 6 | 0 | -0.984652 | 2.694073 | -0.004837 |
| 6 | 6 | 0 | -1.072365 | 1.288067 | -0.111572 |
| 7 | 6 | 0 | -2.317035 | 3.200785 | 0.101172 |
| 8 | 7 | 0 | -2.415613 | 0.955959 | -0.078480 |
| 9 | 6 | 0 | -3.148072 | 2.130418 | 0.066449 |
| 10 | 6 | 0 | 2.519751 | 0.481770 | -0.192079 |
| 11 | 6 | 0 | 3.544759 | -0.150617 | -0.239867 |
| 12 | 6 | 0 | -2.410833 | -1.359396 | -0.866367 |
| 13 | 6 | 0 | -3.011738 | -0.309414 | -0.178127 |
| 14 | 7 | 0 | -4.197121 | -0.424400 | 0.400211 |
| 15 | 6 | 0 | -4.800680 | -1.617060 | 0.307707 |
| 16 | 7 | 0 | -4.303236 | -2.708806 | -0.295659 |
| 17 | 6 | 0 | -3.121738 | -2.547705 | -0.875898 |
| 18 | 6 | 0 | 4.817932 | -0.874330 | -0.309686 |
| 19 | 7 | 0 | -6.029244 | -1.724811 | 0.857820 |
| 20 | 8 | 0 | 4.483751 | -2.251052 | -0.529126 |
| 21 | 6 | 0 | 5.640245 | -0.344502 | -1.488720 |
| 22 | 6 | 0 | 5.605395 | -0.710491 | 1.005146 |
| 23 | 6 | 0 | 4.893542 | -1.231949 | 2.242285 |
| 24 | 1 | 0 | 0.086275 | -0.554728 | -0.186642 |
| 25 | 1 | 0 | 0.397791 | 4.350019 | 0.074829 |
| 26 | 1 | 0 | -2.616846 | 4.232114 | 0.188250 |
| 27 | 1 | 0 | -4.221072 | 2.072353 | 0.115548 |
| 28 | 1 | 0 | -1.480957 | -1.259196 | -1.400338 |
| 29 | 1 | 0 | -2.716450 | -3.414578 | -1.388577 |
| 30 | 1 | 0 | -6.354810 | -0.991355 | 1.460662 |
| 31 | 1 | 0 | -6.429857 | -2.640446 | 0.951163 |

| | | | | | |
|---|---|---|---|---|---|
| 32 | 1 | 0 | 5.312830 | -2.736013 | -0.598830 |
| 33 | 1 | 0 | 6.586165 | -0.887998 | -1.549616 |
| 34 | 1 | 0 | 5.861044 | 0.716101 | -1.366154 |
| 35 | 1 | 0 | 5.093282 | -0.484419 | -2.420546 |
| 36 | 1 | 0 | 5.849232 | 0.347990 | 1.125222 |
| 37 | 1 | 0 | 6.558081 | -1.234380 | 0.870941 |
| 38 | 1 | 0 | 5.530640 | -1.126691 | 3.121877 |
| 39 | 1 | 0 | 4.638514 | -2.286662 | 2.133641 |
| 40 | 1 | 0 | 3.970643 | -0.680271 | 2.429938 |

Conformer # 23
Electronic Energy = -1006.953196 Hartree

| Center Number | Atomic Number | Atomic Type | Coordinates (Angstroms) X | Y | Z |
|---|---|---|---|---|---|
| 1 | 6 | 0 | -0.281939 | 0.618991 | -0.075233 |
| 2 | 6 | 0 | 0.902678 | 1.351802 | -0.131487 |
| 3 | 7 | 0 | 0.961524 | 2.703436 | -0.158927 |
| 4 | 6 | 0 | -0.173279 | 3.379376 | -0.122061 |
| 5 | 6 | 0 | -1.420302 | 2.758620 | -0.059989 |
| 6 | 6 | 0 | -1.461948 | 1.347706 | -0.043729 |
| 7 | 6 | 0 | -2.767701 | 3.231542 | 0.004391 |
| 8 | 7 | 0 | -2.796683 | 0.974360 | 0.039241 |
| 9 | 6 | 0 | -3.563214 | 2.137424 | 0.067100 |
| 10 | 6 | 0 | 2.152318 | 0.658763 | -0.163943 |
| 11 | 6 | 0 | 3.200476 | 0.063909 | -0.187842 |
| 12 | 6 | 0 | -4.629000 | -0.544497 | 0.547328 |
| 13 | 6 | 0 | -3.324075 | -0.320486 | 0.104581 |
| 14 | 7 | 0 | -2.528094 | -1.306225 | -0.272084 |
| 15 | 6 | 0 | -3.031687 | -2.548383 | -0.216523 |
| 16 | 7 | 0 | -4.269172 | -2.882268 | 0.179997 |
| 17 | 6 | 0 | -5.036173 | -1.865537 | 0.554881 |
| 18 | 6 | 0 | 4.499964 | -0.615504 | -0.198351 |
| 19 | 7 | 0 | -2.224676 | -3.549939 | -0.622193 |
| 20 | 8 | 0 | 4.214579 | -2.020718 | -0.209246 |
| 21 | 6 | 0 | 5.267002 | -0.227076 | -1.465725 |
| 22 | 6 | 0 | 5.259182 | -0.227253 | 1.089737 |
| 23 | 6 | 0 | 6.607930 | -0.909453 | 1.269272 |
| 24 | 1 | 0 | -0.269085 | -0.457539 | -0.066386 |

| | | | | | |
|---|---|---|---|---|---|
| 25 | 1 | 0 | -0.097386 | 4.462594 | -0.139857 |
| 26 | 1 | 0 | -3.101828 | 4.255924 | -0.006393 |
| 27 | 1 | 0 | -4.636735 | 2.078917 | 0.098126 |
| 28 | 1 | 0 | -5.282336 | 0.238958 | 0.894974 |
| 29 | 1 | 0 | -6.037018 | -2.118735 | 0.890948 |
| 30 | 1 | 0 | -1.257224 | -3.359466 | -0.807055 |
| 31 | 1 | 0 | -2.520567 | -4.497708 | -0.478274 |
| 32 | 1 | 0 | 5.040306 | -2.480922 | -0.388372 |
| 33 | 1 | 0 | 6.221177 | -0.755615 | -1.508134 |
| 34 | 1 | 0 | 5.467544 | 0.844389 | -1.486197 |
| 35 | 1 | 0 | 4.684170 | -0.491912 | -2.347275 |
| 36 | 1 | 0 | 4.608916 | -0.474232 | 1.931702 |
| 37 | 1 | 0 | 5.383926 | 0.858365 | 1.094954 |
| 38 | 1 | 0 | 7.044873 | -0.620312 | 2.226261 |
| 39 | 1 | 0 | 6.512209 | -1.997491 | 1.277940 |
| 40 | 1 | 0 | 7.320725 | -0.634676 | 0.490343 |

---------------------------------------------------------------------

CompareVOA Results:

Compound **12** (R) configuration

Comparison to (R) calculated: 1900 – 950 $cm^{-1}$

DFT Level: B3PW91 / cc-pVTZ w/ CPCM (chloroform)

Uniform Scaling Factor: 0.982

TNS ($S_{fg}$) IR = 95.9

TNS ($S_{fg}$) VCD = 67.3

SNS (R) config VCD = 73.5

SNS (S) config VCD = 8.0

ESI (Enantiomeric Similarity Index) = SNS (R) – SNS (S) = 65.5

Confidence Level = 99 (Assigned Configuration)

**Table S1. K192 % occupancy data for compound 3 (N=1).**

| **Kinase** | Compound **3** % occupancy |
|---|---|
| LRRK2 | -4 |
| MAPK6 | 73 |
| IRAK3 | 1 |
| TEK | 6 |
| TNK1 | -3 |
| GAK | 2 |
| MAPK4 | -2 |
| AAK1 | 7 |
| AURKA | -8 |
| AURKC | -9 |
| AURKB | 2 |
| NUAK1 | 10 |
| LATS2 | 3 |
| RPS6KA3 | 24 |
| SNF1LK2 | 0 |
| MYLK2 | 3 |
| AXL | 6 |
| FGFR3 | -2 |
| FLT3 | 6 |
| IGF1R | -5 |
| INSR | 0 |
| LIMK2 | -3 |
| TEC | 7 |
| TIE1 | 4 |
| CLK1 | -13 |
| SBK3 | -2 |
| NEK9 | -5 |
| NEK3 | -3 |
| NIM1K | -7 |
| STK36 | -20 |
| ULK2 | -3 |
| ULK3v1 | -1 |
| BRSK1 | 7 |
| MAP3K10 | 0 |
| MAP3K9 | -2 |

| | |
|---|---|
| MYLK3 | 10 |
| PHKG1 | 4 |
| STK33 | -2 |
| STK4 | -35 |
| TLK1 | 1 |
| FGFR1 | 9 |
| FGFR2 | -1 |
| MUSK | 1 |
| NTRK1v1 | 1 |
| RET | 0 |
| NTRK2 | -1 |
| TNK2 | 17 |
| LTK | -8 |
| BRAF(V600E) | -11 |
| IRAK4 | -6 |
| ITK | -4 |
| JAK2 (V617F) | 6 |
| MAP3K11 | 2 |
| PTK2 | -6 |
| PTK6 | 2 |
| PTK2B | -10 |
| BMP2K | -7 |
| NEK5 | -9 |
| STK16 | 2 |
| TBK1 | -1 |
| ULK1 | 5 |
| MAP4K2 | -6 |
| WEE1 | 1 |
| MYLK4 | 8 |
| MARK4 | 2 |
| PRKAA1 | -1 |
| PRKAA2 | -2 |
| RPS6KA1 | 1 |
| RPS6KA2 | 3 |
| RPS6KA4 | 1 |
| RPS6KA6 | -2 |
| SIK1 | -7 |
| CLK4 | -13 |
| MAPK8 | 5 |

| | |
|---|---|
| MAPK9 | -8 |
| IKBKE | -12 |
| LATS1 | 7 |
| PRKX | 5 |
| CSNK2A2 | 12 |
| HIPK4 | 10 |
| STK10 | 3 |
| FGFR4 | 6 |
| MAP4K1 | 1 |
| MERTK | 0 |
| MET | 41 |
| RON | 0 |
| TYRO3 | -10 |
| LCK | -18 |
| LIMK1 | 0 |
| EPHA1 | 6 |
| EPHA4 | -2 |
| EPHA6 | -4 |
| EPHA7 | -6 |
| EPHB1 | 7 |
| EPHB4 | 3 |
| FYN | 14 |
| ABL2 | 1 |
| BMX | -15 |
| BTK | -4 |
| FER | -3 |
| FES | -1 |
| JAK3 | -3 |
| SRMS | 13 |
| TXK | -20 |
| CLK2 | 6 |
| DYRK1A | 1 |
| DYRK1B | 3 |
| ERN1 | 4 |
| ERN2 | 5 |
| HIPK2 | 1 |
| HIPK3 | 20 |
| ICK | 4 |
| CDK1 + B1 | -28 |

| | |
|---|---|
| CDK2 + E1 | 7 |
| CDK3 + E1 | 0 |
| CDK4 + D3 | -5 |
| CDK5 + CDK5R1 | 8 |
| CDK6 + D1 | -5 |
| CDK7 + pMB | -1 |
| CDK9 + K | 5 |
| CDK10 + L2 | 2 |
| CDK14 + Y | -3 |
| CDK15 + Y | 7 |
| CDK16 + Y | 0 |
| CDK17 + Y | -5 |
| CDK18 + Y | -2 |
| CDKL2 | -21 |
| CDK20 + H | -10 |
| CDKL1 | 1 |
| CSNK2A1 | -11 |
| CDKL3 | -3 |
| CDKL5 | 35 |
| JNK3 | 4 |
| MAPK11 | -5 |
| MAPK14 | 7 |
| NLK | -1 |
| NEK11 | -4 |
| NEK1 | -9 |
| NEK2 | -7 |
| NEK4 | -17 |
| PAK4 | 12 |
| MAP4K3 | 0 |
| STK11 | -2 |
| SLK | 2 |
| DAPK2 | 4 |
| MAP3K2 | -5 |
| PLK2 | -2 |
| PLK3 | 3 |
| PLK4 | 4 |
| STK35 | 11 |
| STK17B | -10 |
| TLK2 | 6 |

| | |
|---|---|
| BRSK2 | -9 |
| MARK2 | 13 |
| MELK | -2 |
| CSNK1A1L | 4 |
| CSNK1D | 27 |
| CSNK1G2 | 10 |
| SIK3 | 7 |
| SNRK | -5 |
| CAMK1 | 1 |
| CAMK2A | -3 |
| CAMK2D | -77 |
| CHEK2 | 17 |
| DCLK3 | -6 |
| MKNK2 | 1 |
| PHKG2 | -15 |
| MAP3K3 | 2 |
| RIOK2 | -1 |
| MAP4K5 | 6 |
| MAST3 | -2 |
| MAST4 | 1 |
| STK32B | 5 |
| STK3 | 2 |
| STK38 | 4 |
| STK38L | 8 |
| PAK6 | 15 |
| AKT2 | 23 |
| PKMYT1 | -13 |
| PRKACA | -8 |
| PRKACB | 15 |
| PRKCE | -1 |
| SGK1 | 4 |
| WEE2 | 2 |
| RIPK1 | -3 |
| RIPK2 | -9 |
| TNNI3K | -7 |
| MLTK | 10 |
| MAP3K12 | -12 |
| MAP3K19 | -5 |
| MAP3K21 | 6 |

| | |
|---|---|
| MAP3K4 | -10 |

**Table S2. K300 % occupancy data for compound 3 (N=2).**

| Kinase | %Occ 1 | %Occ 2 | %Occ Avg |
|---|---|---|---|
| PIP4K2C | 86 | 87 | 86 |
| CDK11B+Cyclin K | 86 | 85 | 85 |
| CDK11A+Cyclin K | 76 | 75 | 75 |
| CSNK1D | 69 | 67 | 68 |
| CSNK1E | 51 | 48 | 49 |
| NEK3 | 37 | 37 | 37 |
| EGFR | 50 | 19 | 35 |
| MYLK4 | 30 | 36 | 33 |
| CDK7 | 24 | 28 | 26 |
| MAP2K6 | 30 | 16 | 23 |
| PAK6 | 23 | 23 | 23 |
| HIPK3 | 22 | 20 | 21 |
| COQ8B | 24 | 17 | 20 |
| EPHA2 | 40 | 0 | 20 |
| PIP5K1B | 17 | 19 | 18 |
| DYRK2 | 28 | 7 | 17 |
| CDK20+Cyclin H | 18 | 15 | 16 |
| TEC | 2 | 29 | 16 |
| CAMK2B | 4 | 27 | 15 |
| CDK12 | 14 | 14 | 14 |
| PKN1 | 9 | 18 | 13 |
| IRAK1 | 13 | 12 | 12 |
| TLK2 | 9 | 13 | 11 |
| EPHA10 | 27 | -5 | 11 |
| STK36 | 11 | 11 | 11 |
| DCLK1 | 13 | 8 | 10 |
| MAP4K5 | 10 | 10 | 10 |
| EPHA8 | 6 | 14 | 10 |
| PIKFYVE | 3 | 17 | 10 |
| STK4 | 17 | 3 | 10 |
| CDK13 | 7 | 9 | 8 |
| CDK10+Cyclin L2 | 10 | 6 | 8 |
| SBK3 | 7 | 7 | 7 |
| MOK | 4 | 11 | 7 |
| TSSK1B | 9 | 6 | 7 |
| PIM2 | 11 | 3 | 7 |
| ICK | 5 | 9 | 7 |
| LTK | 5 | 8 | 7 |
| NUAK2 | 12 | 2 | 7 |
| ULK3 | 8 | 5 | 7 |

| | | | |
|---|---|---|---|
| CDK19+Cyclin C | 6 | 7 | 7 |
| PRKCH | 8 | 5 | 6 |
| PKMYT1 | 10 | 2 | 6 |
| CAMK1G | 6 | 6 | 6 |
| CIT | 6 | 6 | 6 |
| PAK4 | 6 | 5 | 6 |
| JAK2 | 8 | 3 | 6 |
| CAMK2G | 6 | 5 | 6 |
| TXK | 7 | 5 | 6 |
| ULK1 | 5 | 6 | 5 |
| MAPK4 | 6 | 5 | 5 |
| CSNK1G2 | 5 | 5 | 5 |
| RPS6KA6 | 5 | 5 | 5 |
| PIK3CD+PIK3R1 | -1 | 11 | 5 |
| MAPK6 | 3 | 7 | 5 |
| FER | 5 | 4 | 5 |
| PDPK1 | 5 | 4 | 5 |
| CDK17+Cyclin Y | 6 | 3 | 4 |
| ROCK1 | 5 | 4 | 4 |
| PIM1 | 1 | 8 | 4 |
| TTK | 1 | 7 | 4 |
| FGR | -5 | 13 | 4 |
| STK32A | -1 | 9 | 4 |
| STK17A | -16 | 23 | 4 |
| GAK | 3 | 4 | 4 |
| NIM1K | 2 | 6 | 4 |
| PKN2 | 3 | 4 | 4 |
| MYLK3 | 3 | 4 | 3 |
| PRKACB | 3 | 4 | 3 |
| CDKL2 | 3 | 3 | 3 |
| DDR2 | 4 | 2 | 3 |
| CSNK1A1L | 4 | 2 | 3 |
| SRC | 2 | 4 | 3 |
| CDK18+Cyclin Y | 3 | 3 | 3 |
| SGK2 | -3 | 9 | 3 |
| TIE1 | 4 | 2 | 3 |
| MAST4 | 5 | 1 | 3 |
| WEE2 | -2 | 7 | 3 |
| PRKCD | 3 | 3 | 3 |
| CDK9+Cyclin K | 4 | 2 | 3 |
| CDK15+Cyclin Y | 2 | 4 | 3 |
| TEK | 1 | 4 | 3 |
| BRSK1 | 2 | 4 | 3 |

| | | | |
|---|---|---|---|
| FLT1 | 8 | -3 | 2 |
| MET | 7 | -2 | 2 |
| CDK5+CDK5R1 | 1 | 4 | 2 |
| SIK1 | 3 | 1 | 2 |
| CRAF | 8 | -3 | 2 |
| PIK3C3 | 0 | 4 | 2 |
| STK24 | 4 | 0 | 2 |
| TLK1 | 4 | 1 | 2 |
| MLTK | 3 | 1 | 2 |
| PIM3 | 15 | -11 | 2 |
| EPHA5 | -7 | 11 | 2 |
| TBK1 | 3 | 1 | 2 |
| HCK | 2 | 2 | 2 |
| MAP3K21 | 5 | -2 | 2 |
| RIPK3 | 3 | 1 | 2 |
| GSK3A | 1 | 2 | 2 |
| MAP4K3 | 1 | 2 | 2 |
| ADK | -1 | 4 | 1 |
| KIT | -3 | 6 | 1 |
| PRKAA1 | 2 | 1 | 1 |
| CDK3+Cyclin E1 | 0 | 2 | 1 |
| NLK | 2 | -1 | 1 |
| DDR1 | -5 | 7 | 1 |
| GSK3B | -1 | 3 | 1 |
| CLK2 | 0 | 1 | 1 |
| CDKL5 | 1 | 1 | 1 |
| STK33 | -5 | 7 | 1 |
| SGK1 | 2 | -1 | 1 |
| RIOK2 | 2 | -1 | 1 |
| CDK2+Cyclin E1 | 2 | -1 | 0 |
| CDK16+Cyclin Y | -1 | 2 | 0 |
| LYN | -6 | 7 | 0 |
| STK32B | 1 | 0 | 0 |
| NEK2 | 3 | -2 | 0 |
| MKNK2 | 1 | -1 | 0 |
| MAP4K2 | 1 | -1 | 0 |
| CLK4 | 5 | -5 | 0 |
| FES | -3 | 3 | 0 |
| VRK2 | -2 | 2 | 0 |
| PLK2 | -3 | 2 | 0 |
| MAPK14 | -3 | 3 | 0 |
| HUNK | 6 | -6 | 0 |
| ERN2 | -1 | 0 | 0 |

| | | | |
|---|---|---|---|
| STK35 | 0 | -1 | 0 |
| INSR | 1 | -1 | 0 |
| RPS6KA3 | 0 | -1 | 0 |
| RPS6KA1 | -1 | 0 | 0 |
| CDK6+Cyclin D1 | -1 | 0 | 0 |
| CDK1+Cyclin B1 | -1 | 0 | 0 |
| MAP4K1 | 1 | -2 | -1 |
| CAMK2D | -2 | 1 | -1 |
| BMX | -2 | 0 | -1 |
| DCLK3 | -1 | 0 | -1 |
| MAP3K3 | -1 | 0 | -1 |
| DAPK2 | -3 | 1 | -1 |
| RIPK2 | -1 | -1 | -1 |
| TGFBR1 | -1 | -1 | -1 |
| CDK14+Cyclin Y | 2 | -4 | -1 |
| MAPK11 | -3 | 1 | -1 |
| MARK4 | -3 | 1 | -1 |
| STK11 | -3 | 0 | -1 |
| EPHA6 | 3 | -6 | -1 |
| MAP3K2 | -2 | -1 | -1 |
| MAP3K13 | -7 | 4 | -1 |
| LIMK2 | -2 | -1 | -1 |
| STK26 | -2 | -1 | -2 |
| NTRK2 | -3 | -1 | -2 |
| MELK | -2 | -1 | -2 |
| SRMS | 0 | -4 | -2 |
| ROCK2 | -7 | 4 | -2 |
| STK16 | 0 | -3 | -2 |
| STK38 | -2 | -1 | -2 |
| LATS2 | -2 | -1 | -2 |
| CDK2+Cyclin A1 | -2 | -2 | -2 |
| CAMK1 | -2 | -1 | -2 |
| PRKACA | -3 | -1 | -2 |
| PIK3CA+PIK3R1 | -2 | -2 | -2 |
| CDKL1 | 1 | -5 | -2 |
| NEK11 | 0 | -4 | -2 |
| MARK1 | -8 | 4 | -2 |
| BRSK2 | -3 | -1 | -2 |
| HIPK4 | -2 | -2 | -2 |
| RON | -7 | 2 | -2 |
| CDK8+Cyclin C | -4 | -1 | -2 |
| JNK3 | -3 | -3 | -3 |
| AURKB | 1 | -6 | -3 |

| | | | |
|---|---|---|---|
| MAP3K11 | -2 | -4 | -3 |
| EPHA3 | -9 | 3 | -3 |
| MARK3 | -3 | -3 | -3 |
| MYLK2 | -3 | -3 | -3 |
| ERN1 | -1 | -5 | -3 |
| GRK7 | 3 | -9 | -3 |
| TYK2 | -1 | -5 | -3 |
| CASK | -7 | 1 | -3 |
| PTK2B | -1 | -5 | -3 |
| NEK9 | -4 | -2 | -3 |
| MAP3K4 | -5 | -1 | -3 |
| PRKX | -6 | 0 | -3 |
| PRKAA2 | -1 | -6 | -3 |
| IGF1R | -4 | -3 | -3 |
| AXL | -3 | -4 | -3 |
| BMP2K | 0 | -7 | -4 |
| AKT2 | -1 | -6 | -4 |
| JAK3 | -6 | -1 | -4 |
| EPHB3 | -5 | -2 | -4 |
| PLK3 | -6 | -2 | -4 |
| BTK | -2 | -5 | -4 |
| EPHB4 | -6 | -1 | -4 |
| MAST3 | -1 | -8 | -4 |
| AURKC | -3 | -6 | -4 |
| MAPK9 | 0 | -9 | -4 |
| RIPK1 | -6 | -3 | -5 |
| STK17B | -11 | 2 | -5 |
| ARAF | 5 | -15 | -5 |
| BMPR1A | 12 | -21 | -5 |
| TNNI3K | -4 | -6 | -5 |
| CHEK1 | -7 | -3 | -5 |
| ABL1 | -15 | 5 | -5 |
| CSNK2A1 | -9 | -1 | -5 |
| MAP3K10 | -2 | -8 | -5 |
| ALK | -4 | -6 | -5 |
| STK3 | -8 | -2 | -5 |
| CHEK2 | -7 | -3 | -5 |
| PTK2 | -10 | -1 | -5 |
| EIF2AK4 K2 Domain | -2 | -8 | -5 |
| IRAK4 | -2 | -9 | -5 |
| NEK4 | -6 | -5 | -6 |
| PLK4 | -6 | -5 | -6 |

| | | | |
|---|---|---|---|
| RET | -3 | -8 | -6 |
| BRAF (V600E) | -6 | -6 | -6 |
| PRKCE | -8 | -4 | -6 |
| MAP2K5 | 0 | -11 | -6 |
| CLK1 | -4 | -7 | -6 |
| LIMK1 | -6 | -5 | -6 |
| PIK3CB+PIK3R1 | -9 | -3 | -6 |
| PTK6 | -7 | -5 | -6 |
| LATS1 | -5 | -7 | -6 |
| DYRK1A | -6 | -6 | -6 |
| MAPK1 | -4 | -9 | -6 |
| EPHA7 | -6 | -7 | -6 |
| FGFR2 | -8 | -6 | -7 |
| CAMK1D | -5 | -8 | -7 |
| EPHA4 | -5 | -8 | -7 |
| SIK2 | -7 | -6 | -7 |
| EPHB2 | -3 | -10 | -7 |
| ABL2 | -6 | -8 | -7 |
| CSF1R | -5 | -9 | -7 |
| CSK | -20 | 6 | -7 |
| ULK2 | -6 | -8 | -7 |
| MAP3K19 | -7 | -7 | -7 |
| NEK6 | -3 | -11 | -7 |
| BLK | -11 | -4 | -7 |
| FLT3 | -3 | -12 | -7 |
| SLK | -8 | -6 | -7 |
| CAMK2A | -8 | -7 | -7 |
| ITK | -9 | -7 | -8 |
| CDK4+Cyclin D3 | -9 | -7 | -8 |
| PRKCG | -15 | -2 | -8 |
| FGFR4 | -6 | -11 | -8 |
| SIK3 | -13 | -4 | -8 |
| TYRO3 | -9 | -7 | -8 |
| LRRK2 | -9 | -8 | -8 |
| CDKL3 | -3 | -13 | -8 |
| EPHA1 | -7 | -9 | -8 |
| MARK2 | -8 | -10 | -9 |
| EPHB1 | -8 | -9 | -9 |
| MERTK | -9 | -9 | -9 |
| NEK1 | -8 | -10 | -9 |
| STK38L | -11 | -7 | -9 |
| TESK1 | -9 | -9 | -9 |
| PLK1 | -7 | -11 | -9 |

| | | | |
|---|---|---|---|
| MAPK15 | -9 | -9 | -9 |
| FGFR1 | -11 | -8 | -9 |
| AURKA | -13 | -5 | -9 |
| MAP3K12 | -12 | -7 | -9 |
| HIPK2 | -11 | -9 | -10 |
| IKBKE | -9 | -11 | -10 |
| STK10 | -12 | -8 | -10 |
| YES1 | -13 | -7 | -10 |
| TNK2 | -6 | -15 | -11 |
| DYRK1B | -11 | -10 | -11 |
| PHKG1 | -12 | -11 | -11 |
| FRK | -13 | -10 | -12 |
| ACVR1B | -13 | -11 | -12 |
| TNK1 | -13 | -11 | -12 |
| RPS6KA4 | -16 | -8 | -12 |
| TGFBR2 | -15 | -9 | -12 |
| BRAF | -7 | -17 | -12 |
| MAPK3 | -5 | -20 | -13 |
| MAP3K9 | -24 | -1 | -13 |
| PRKCB | -7 | -19 | -13 |
| PHKG2 | -12 | -16 | -14 |
| RPS6KA2 | -14 | -14 | -14 |
| NTRK3 | -12 | -16 | -14 |
| CSNK2A2 | -1 | -27 | -14 |
| SGK3 | -12 | -17 | -14 |
| JAK2 (V617F) | -5 | -24 | -15 |
| MUSK | -9 | -20 | -15 |
| WEE1 | -18 | -12 | -15 |
| PRKG2 | -19 | -11 | -15 |
| LCK | -14 | -16 | -15 |
| FGFR3 | -16 | -15 | -15 |
| SNRK | -15 | -19 | -17 |
| PRKCA | -18 | -16 | -17 |
| PRKCQ | -13 | -21 | -17 |
| IRAK3 | -12 | -23 | -17 |
| AKT1 | -31 | -4 | -17 |
| PKN3 | -27 | -10 | -18 |
| MAPK8 | -17 | -21 | -19 |
| ACVR1 | -21 | -19 | -20 |
| ACVRL1 | -17 | -27 | -22 |
| NUAK1 | -37 | -11 | -24 |
| MAP3K7 | -34 | -19 | -26 |
| FYN | -31 | -35 | -33 |

| | | | |
|---|---|---|---|
| NTRK1 | -43 | -27 | -35 |
| AAK1 | -56 | -58 | -57 |
| NEK5 | -69 | -55 | -62 |

**Table S3. K300 % occupancy data for compound 13 (N=2).**

| Kinase | %Occ 1 | %Occ 2 | %Occ Avg |
|---|---|---|---|
| CDK11B+Cyclin K | 90 | 92 | 91 |
| CDK11A+Cyclin K | 82 | 79 | 80 |
| CSNK1D | 76 | 73 | 75 |
| CSNK1E | 60 | 59 | 60 |
| EPHA2 | 52 | 50 | 51 |
| PIP4K2C | 42 | 41 | 41 |
| NEK3 | 35 | 37 | 36 |
| STK4 | 43 | 19 | 31 |
| PAK6 | 28 | 25 | 26 |
| PRKCD | 36 | 16 | 26 |
| CDK7 | 24 | 25 | 24 |
| MYLK4 | 16 | 23 | 19 |
| MAPK6 | 14 | 21 | 18 |
| MAP2K6 | 35 | 0 | 17 |
| PRKCH | 27 | 2 | 15 |
| MOK | 14 | 14 | 14 |
| HIPK3 | 9 | 16 | 12 |
| PKN2 | 7 | 18 | 12 |
| IRAK1 | 10 | 14 | 12 |
| DCLK1 | 14 | 9 | 11 |
| MAP4K5 | 13 | 9 | 11 |
| CDK20+Cyclin H | 11 | 11 | 11 |
| PAK4 | 9 | 13 | 11 |
| FLT1 | 16 | 6 | 11 |
| PIP5K1B | 8 | 13 | 11 |
| CSNK1G2 | 9 | 11 | 10 |
| TSSK1B | 7 | 12 | 10 |
| STK36 | 8 | 10 | 9 |
| PKN1 | 7 | 10 | 9 |
| COQ8B | 7 | 10 | 9 |
| CLK4 | 11 | 5 | 8 |
| CDK13 | 13 | 2 | 8 |
| CSNK1A1L | 5 | 10 | 7 |
| ROCK2 | 0 | 14 | 7 |
| EPHA10 | 10 | 4 | 7 |
| CAMK2G | 8 | 6 | 7 |
| SGK3 | 12 | 1 | 7 |
| TLK2 | 11 | 2 | 6 |
| RPS6KA6 | 6 | 7 | 6 |
| STK24 | 6 | 5 | 6 |

| | | | |
|---|---|---|---|
| CDK10+Cyclin L2 | 6 | 4 | 5 |
| PHKG1 | 22 | -12 | 5 |
| SRC | 6 | 4 | 5 |
| TXK | 4 | 6 | 5 |
| ADK | 2 | 7 | 5 |
| GSK3A | 7 | 2 | 5 |
| ALK | 3 | 6 | 5 |
| MKNK2 | 6 | 3 | 4 |
| DDR2 | -2 | 11 | 4 |
| MARK4 | 5 | 3 | 4 |
| EGFR | 2 | 5 | 4 |
| TBK1 | 5 | 3 | 4 |
| MAP3K13 | 3 | 4 | 3 |
| GAK | 3 | 3 | 3 |
| ULK3 | 1 | 5 | 3 |
| CIT | 3 | 3 | 3 |
| MAP3K2 | 5 | 2 | 3 |
| GRK7 | 6 | 0 | 3 |
| MAPK4 | 2 | 4 | 3 |
| TTK | 3 | 3 | 3 |
| TIE1 | 3 | 3 | 3 |
| CDK9+Cyclin K | 2 | 4 | 3 |
| RIPK3 | -6 | 12 | 3 |
| RIPK2 | 2 | 4 | 3 |
| MAST4 | 1 | 4 | 3 |
| LTK | 9 | -4 | 3 |
| FES | 1 | 4 | 2 |
| DCLK3 | 7 | -2 | 2 |
| CDK15+Cyclin Y | 3 | 2 | 2 |
| MAPK14 | 3 | 1 | 2 |
| EPHA5 | 21 | -17 | 2 |
| SGK2 | -7 | 11 | 2 |
| STK26 | 0 | 5 | 2 |
| SBK3 | 3 | 1 | 2 |
| CASK | 4 | 1 | 2 |
| NIM1K | 3 | 1 | 2 |
| CDK16+Cyclin Y | 1 | 3 | 2 |
| PRKACB | 1 | 3 | 2 |
| CDK12 | 5 | -1 | 2 |
| STK33 | 4 | 0 | 2 |
| TLK1 | 1 | 3 | 2 |
| MYLK3 | 0 | 3 | 2 |
| RON | 0 | 4 | 2 |

| | | | |
|---|---|---|---|
| AURKB | 3 | 0 | 2 |
| BRAF (V600E) | -5 | 8 | 2 |
| CDK3+Cyclin E1 | 0 | 3 | 2 |
| BRSK1 | 1 | 2 | 2 |
| SGK1 | -3 | 5 | 1 |
| CAMK1G | 4 | -2 | 1 |
| CDK17+Cyclin Y | 1 | 1 | 1 |
| HUNK | -1 | 3 | 1 |
| NTRK2 | 4 | -2 | 1 |
| MAP3K21 | -1 | 3 | 1 |
| PRKCE | -3 | 5 | 1 |
| NLK | 1 | 1 | 1 |
| ICK | 7 | -5 | 1 |
| EPHB3 | 0 | 2 | 1 |
| CDK6+Cyclin D1 | -1 | 2 | 1 |
| FER | 5 | -3 | 1 |
| MET | 2 | 0 | 1 |
| PKMYT1 | 1 | 1 | 1 |
| NUAK2 | -12 | 13 | 1 |
| TYK2 | -2 | 3 | 1 |
| CLK2 | -1 | 1 | 0 |
| BRAF | 11 | -11 | 0 |
| STK11 | -1 | 1 | 0 |
| CDKL2 | -1 | 1 | 0 |
| CHEK1 | 7 | -7 | 0 |
| CAMK1 | 1 | -1 | 0 |
| TEK | 1 | -2 | 0 |
| AKT1 | -7 | 7 | 0 |
| CDK18+Cyclin Y | 0 | 0 | 0 |
| CDKL5 | 3 | -4 | 0 |
| RIOK2 | 0 | -1 | 0 |
| CDK14+Cyclin Y | -2 | 1 | 0 |
| MLTK | -1 | 0 | -1 |
| PIK3CD+PIK3R1 | -2 | 1 | -1 |
| STK3 | 4 | -6 | -1 |
| CDK5+CDK5R1 | -2 | 0 | -1 |
| STK32A | -4 | 3 | -1 |
| SIK1 | -1 | 0 | -1 |
| PDPK1 | 10 | -12 | -1 |
| ROCK1 | 6 | -8 | -1 |
| CDK8+Cyclin C | -2 | 0 | -1 |
| CDK1+Cyclin B1 | -1 | -1 | -1 |
| INSR | -1 | -2 | -1 |

| | | | |
|---|---|---|---|
| STK17B | -4 | 1 | -1 |
| CDK19+Cyclin C | 0 | -2 | -1 |
| JAK2 | -8 | 5 | -1 |
| ABL1 | -9 | 6 | -1 |
| PRKCG | -7 | 4 | -1 |
| EIF2AK4 K2 Domain | -1 | -2 | -1 |
| EPHB2 | -17 | 14 | -2 |
| PIK3C3 | 0 | -3 | -2 |
| MAP4K3 | -1 | -2 | -2 |
| MAP3K3 | -1 | -2 | -2 |
| CSF1R | -9 | 6 | -2 |
| CDK2+Cyclin E1 | -1 | -3 | -2 |
| CAMK2D | -3 | -1 | -2 |
| BTK | -4 | 0 | -2 |
| GSK3B | -4 | 0 | -2 |
| ERN2 | -3 | -1 | -2 |
| PTK2B | 0 | -4 | -2 |
| WEE2 | -2 | -3 | -2 |
| PIK3CA+PIK3R1 | -1 | -3 | -2 |
| MAP3K19 | -6 | 2 | -2 |
| STK38 | -2 | -2 | -2 |
| CDK2+Cyclin A1 | -2 | -2 | -2 |
| MAST3 | 1 | -5 | -2 |
| LIMK2 | -2 | -3 | -2 |
| PRKACA | -2 | -3 | -2 |
| BRSK2 | -2 | -3 | -3 |
| STK16 | -4 | -2 | -3 |
| IGF1R | -4 | -1 | -3 |
| STK32B | -3 | -2 | -3 |
| PIM3 | -8 | 2 | -3 |
| DAPK2 | -1 | -4 | -3 |
| CAMK1D | 1 | -7 | -3 |
| MAP3K9 | -5 | -1 | -3 |
| MAP4K2 | 0 | -6 | -3 |
| PIM1 | 3 | -8 | -3 |
| JNK3 | -2 | -4 | -3 |
| RPS6KA1 | -2 | -3 | -3 |
| PLK2 | -1 | -5 | -3 |
| JAK3 | -3 | -3 | -3 |
| MAPK11 | -2 | -4 | -3 |
| MAP4K1 | -3 | -3 | -3 |
| AXL | -5 | -2 | -3 |

| | | | |
|---|---|---|---|
| CHEK2 | -4 | -3 | -3 |
| PIM2 | 1 | -8 | -3 |
| KIT | -12 | 5 | -4 |
| TNNI3K | 0 | -7 | -4 |
| EPHB4 | -2 | -5 | -4 |
| TEC | -2 | -6 | -4 |
| MELK | -1 | -6 | -4 |
| STK35 | -6 | -2 | -4 |
| MYLK2 | -2 | -6 | -4 |
| IKBKE | -1 | -7 | -4 |
| BMX | -4 | -4 | -4 |
| PRKAA2 | -2 | -6 | -4 |
| CDK4+Cyclin D3 | -5 | -4 | -4 |
| NEK11 | -5 | -4 | -4 |
| MAP3K4 | -3 | -6 | -4 |
| LATS1 | -7 | -2 | -4 |
| RIPK1 | -4 | -5 | -4 |
| ULK2 | -4 | -5 | -4 |
| NEK4 | -5 | -4 | -5 |
| PLK4 | -3 | -6 | -5 |
| LATS2 | -4 | -6 | -5 |
| NTRK3 | -12 | 2 | -5 |
| AKT2 | -7 | -3 | -5 |
| NEK9 | -4 | -6 | -5 |
| MAP2K5 | -5 | -5 | -5 |
| BLK | -8 | -2 | -5 |
| DDR1 | -1 | -9 | -5 |
| PIKFYVE | -13 | 3 | -5 |
| IRAK4 | -1 | -10 | -5 |
| RPS6KA3 | -6 | -5 | -5 |
| CAMK2A | -5 | -6 | -5 |
| CLK1 | -7 | -3 | -5 |
| MAP3K11 | -5 | -7 | -6 |
| LIMK1 | -5 | -6 | -6 |
| PRKCQ | -8 | -3 | -6 |
| EPHA8 | 5 | -17 | -6 |
| FLT3 | 0 | -12 | -6 |
| MAP3K12 | -6 | -5 | -6 |
| MAP3K10 | -5 | -7 | -6 |
| SRMS | -6 | -6 | -6 |
| EPHA4 | -7 | -5 | -6 |
| EPHA3 | 0 | -13 | -6 |
| HIPK4 | -7 | -5 | -6 |

| | | | |
|---|---|---|---|
| CSNK2A2 | -7 | -5 | -6 |
| NUAK1 | -7 | -5 | -6 |
| PRKX | -4 | -9 | -6 |
| ABL2 | -9 | -4 | -7 |
| SLK | -9 | -4 | -7 |
| PLK3 | -6 | -7 | -7 |
| EPHB1 | -7 | -6 | -7 |
| PTK6 | -5 | -9 | -7 |
| PHKG2 | -8 | -5 | -7 |
| HCK | -9 | -4 | -7 |
| MARK2 | -8 | -6 | -7 |
| ITK | -9 | -5 | -7 |
| EPHA7 | -9 | -5 | -7 |
| CDKL1 | -5 | -9 | -7 |
| EPHA6 | -7 | -8 | -7 |
| DYRK1A | -8 | -7 | -7 |
| DYRK2 | -12 | -2 | -7 |
| NEK1 | -5 | -9 | -7 |
| MAPK9 | -7 | -7 | -7 |
| CDKL3 | -3 | -12 | -7 |
| MAPK1 | -10 | -5 | -8 |
| PRKAA1 | -7 | -8 | -8 |
| TYRO3 | -9 | -7 | -8 |
| RET | -9 | -7 | -8 |
| SIK2 | -7 | -9 | -8 |
| RPS6KA2 | -10 | -6 | -8 |
| MARK1 | -8 | -8 | -8 |
| AURKA | -6 | -10 | -8 |
| SIK3 | -12 | -5 | -8 |
| EPHA1 | -9 | -8 | -8 |
| LYN | -17 | 0 | -9 |
| STK38L | -6 | -11 | -9 |
| BMPR1A | -5 | -12 | -9 |
| CSNK2A1 | -11 | -7 | -9 |
| FGFR2 | -9 | -9 | -9 |
| BMP2K | -11 | -7 | -9 |
| ACVR1B | -13 | -5 | -9 |
| NEK2 | -9 | -9 | -9 |
| TESK1 | -10 | -8 | -9 |
| MERTK | -7 | -11 | -9 |
| ERN1 | -12 | -6 | -9 |
| YES1 | -12 | -7 | -9 |
| MAPK15 | -1 | -18 | -10 |

| | | | |
|---|---|---|---|
| NEK6 | -11 | -8 | -10 |
| FGFR4 | -7 | -12 | -10 |
| PIK3CB+PIK3R1 | -4 | -15 | -10 |
| MARK3 | -9 | -11 | -10 |
| RPS6KA4 | -12 | -9 | -10 |
| AURKC | -12 | -9 | -11 |
| DYRK1B | -10 | -11 | -11 |
| CRAF | -18 | -5 | -11 |
| STK10 | -22 | -1 | -11 |
| ACVR1 | -6 | -17 | -12 |
| FRK | -25 | 1 | -12 |
| LCK | -11 | -13 | -12 |
| TGFBR1 | -14 | -10 | -12 |
| HIPK2 | -13 | -11 | -12 |
| PLK1 | -6 | -19 | -12 |
| FGFR1 | -13 | -12 | -13 |
| JAK2 (V617F) | -20 | -6 | -13 |
| PRKG2 | -18 | -7 | -13 |
| SNRK | -12 | -15 | -13 |
| CAMK2B | -6 | -20 | -13 |
| ARAF | -8 | -20 | -14 |
| WEE1 | -13 | -15 | -14 |
| IRAK3 | -17 | -12 | -14 |
| TNK2 | -13 | -17 | -15 |
| TNK1 | -15 | -15 | -15 |
| ULK1 | -14 | -16 | -15 |
| MAPK3 | -15 | -15 | -15 |
| PTK2 | -14 | -17 | -15 |
| TGFBR2 | -24 | -9 | -17 |
| LRRK2 | -17 | -18 | -17 |
| FGR | 13 | -49 | -18 |
| FGFR3 | -23 | -16 | -20 |
| MUSK | -22 | -20 | -21 |
| PRKCB | -16 | -26 | -21 |
| MAP3K7 | -23 | -21 | -22 |
| CSK | -51 | 8 | -22 |
| AAK1 | -53 | 8 | -23 |
| ACVRL1 | -25 | -22 | -24 |
| MAPK8 | -23 | -24 | -24 |
| PRKCA | -29 | -24 | -27 |
| NTRK1 | -41 | -16 | -28 |
| VRK2 | -23 | -36 | -30 |
| FYN | -32 | -36 | -34 |

| | | | |
|---|---|---|---|
| PKN3 | -32 | -42 | -37 |
| NEK5 | -69 | -6 | -37 |
| STK17A | -100 | -6 | -53 |

**Table S4**. Enzymatic $IC_{50}$ and NanoBRET data for select off-targets of TTBK1-IN-1 and compound **13**. Green indicates outside 30-fold selectivity window and orange indicates within 30-fold selectivity window.

| Kinase | Compound 13 NB $IC_{50}$ | Compound 13 enzymatic $IC_{50}$ | Fold over TTBK1 enzymatic $IC_{50}$ | TTBK probe K300 % occup. | TTBK1-IN-1 NB | TTBK1-IN-1 RBC $IC_{50}$ | Fold over TTBK1 enzymatic $IC_{50}$ |
|---|---|---|---|---|---|---|---|
| PIP4K2C | 1600 nM | N/A | N/A | 41 | 6600 nM | N/A | N/A |
| PIKFYVE | 110 nM | 3.45 nM | 1 | -5 | 640 nM | 51.3 nM | 0.6 |
| MAP3K14 | N/A | 6.42 nM | 2 | N/A | N/A | 139 nM | 2 |
| CDK11B | 160 nM | 85.7 nM | 29 | 91 | N/A | >10,000 nM | > 30 |
| CK1a1 | N/A | 143 nM | 48 | N/A | N/A | 1890 nM | 22 |
| CK1δ | 190 nM | 7.68 nM | 3 | 75 | 990 nM | 50.0 nM | 0.6 |
| CK1ε | 1100 | 50.4 nM | 17 | 60 | 3500 nM | 164 nM | 2 |
| CK1g1 | N/A | 510 nM | 170 | N/A | N/A | 857 nM | 10 |
| EPHA2 | >30,000 nM | > 10,000 nM | > 30 | 51 | N/A | >10,000 nM | > 30 |
| JNK1 | N/A | 443 nM | 148 | N/A | N/A | 315 nM | 4 |
| MAP4K5 | >10,000 nM | 4.48 nM | 2 | 11 | >10,000 nM | 28.8 nM | 0.3 |
| MKK4 | N/A | 2500 nM | 833 | N/A | N/A | 3080 nM | 36 |
| NEK3 | N/A | > 10,000 nM | 3333 | 36 | N/A | 6750 nM | 80 |

| | | | | | | | |
|---|---|---|---|---|---|---|---|
| PAK4 | > 10,000 nM | 22.6 nM | 8 | 11 | >10,000 nM | 421 nM | 5 |
| PAK5 | > 10,000 nM | 83.7 nM | 28 | N/A | >10,000 nM | 1840 nM | 22 |
| PAK6 | > 10,000 nM | 37.1 nM | 12 | 26 | >10,000 nM | 315 nM | 3.8 |
| PRKD1 | N/A | 40.4 nM | 14 | N/A | N/A | 20.4 nM | 0.2 |
| PRKD3 | N/A | 28.3 nM | 9 | N/A | N/A | 16.3 nM | 0.2 |
| PRKD2 | N/A | 341 nM | 114 | N/A | N/A | 129 nM | 1.5 |
| RIPK5 | N/A | >10,000 nM | 3333 | N/A | N/A | 5650 nM | 67 |
| STK21 | N/A | 155 nM | 52 | 3 | N/A | 6460 nM | 77 |
| TAOK1 | N/A | 28.3 nM | 9 | N/A | N/A | 392 nM | 5 |
| TSSK3 | N/A | 139 nM | 46 | N/A | N/A | 435 nM | 5 |
| MAP3K19 | > 30,000 nM | 6.47 nM | 2 | -2 | >30,000 nM | 84.5 nM | 1 |

**Purity traces and spectra for all compounds.**

$^{1}H$ NMR (500 MHz, $CD_3OD$) for Compound **1**.

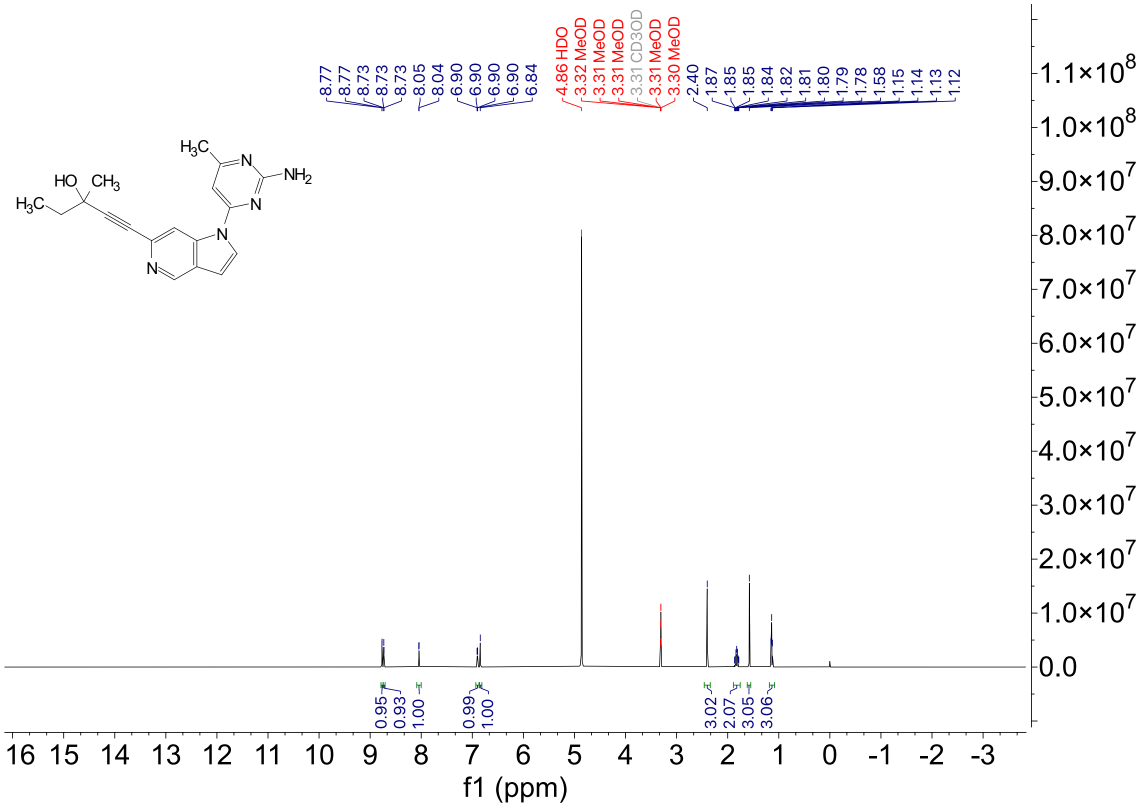

$^{13}C$ NMR (126 MHz, $CD_3OD$) for Compound **1.**

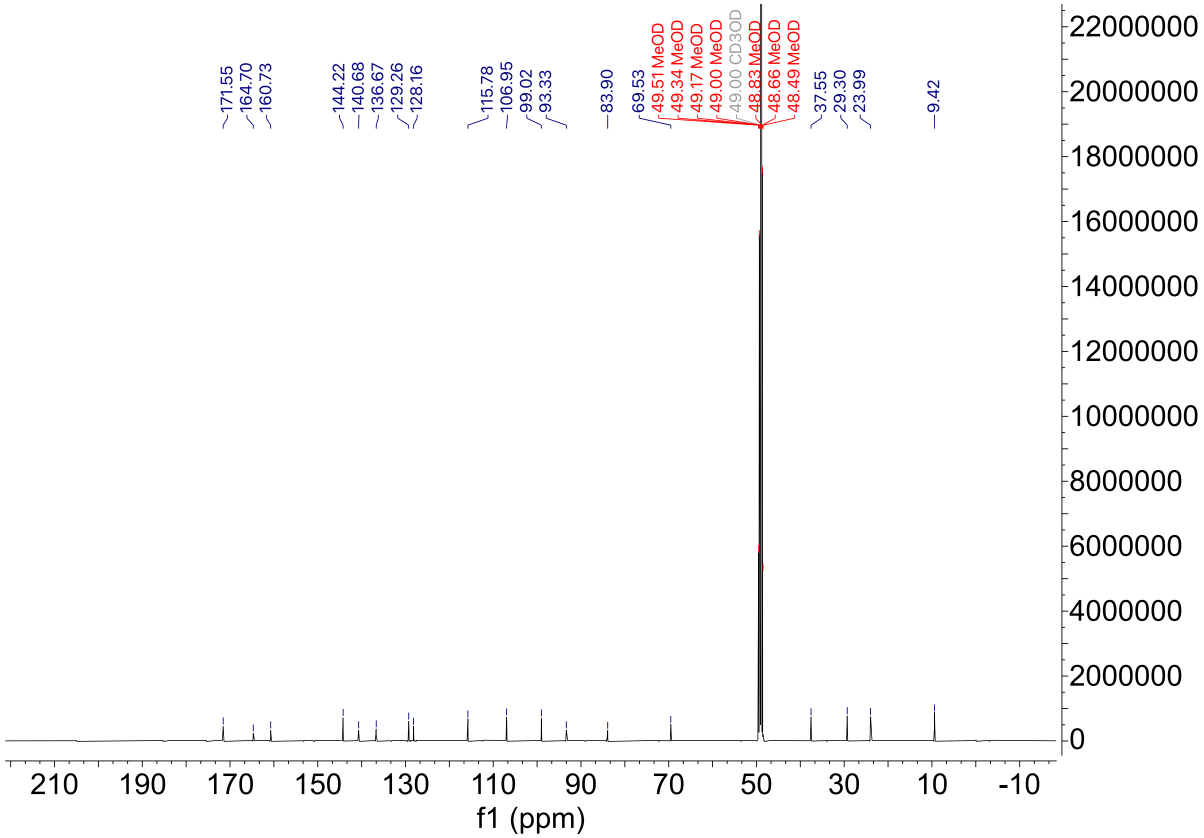

HPLC Tracer for compound **1**

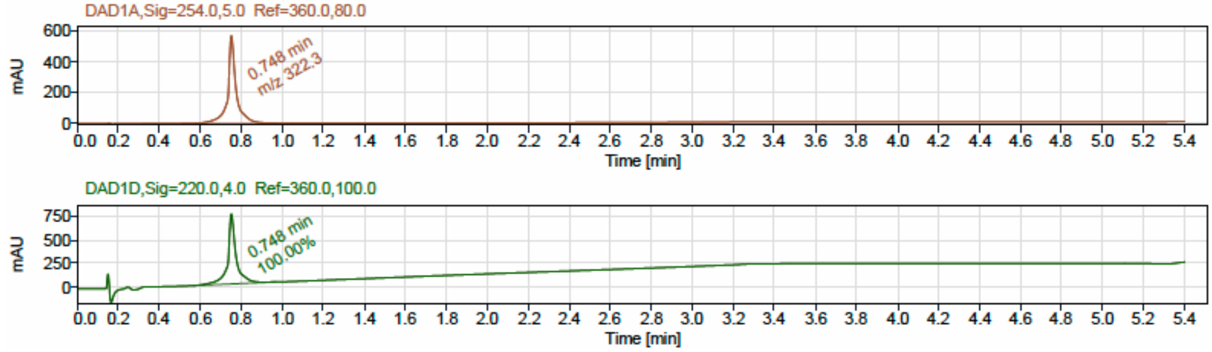


LCMS

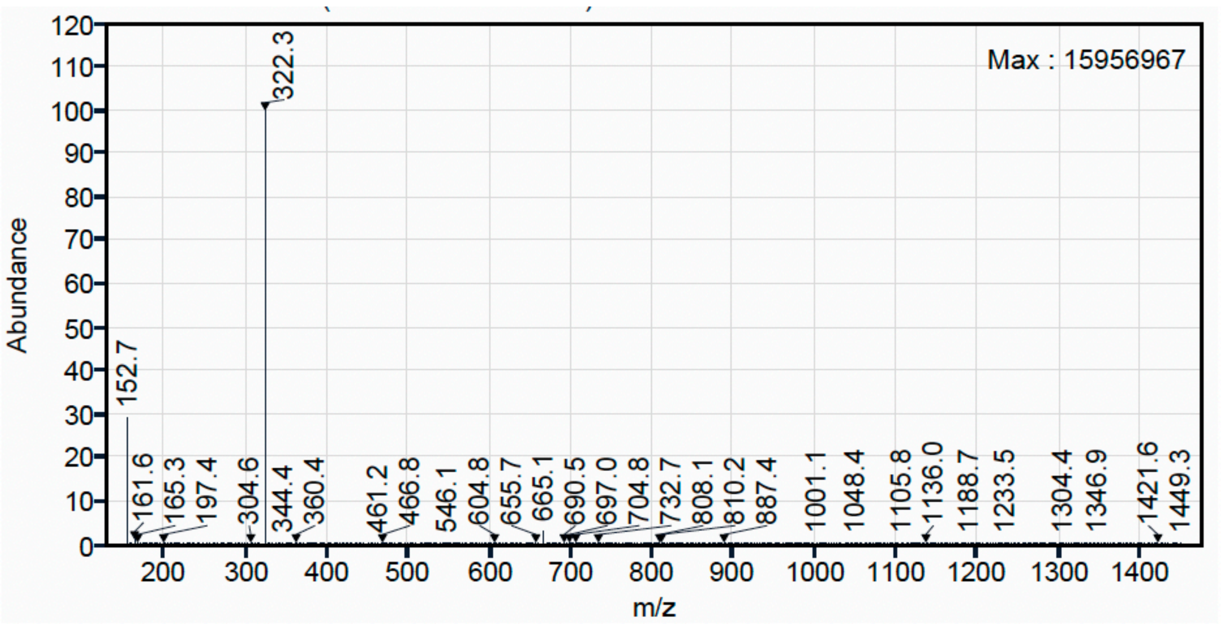

[1]H NMR (500 MHz, $CD_3OD$) for Compound **2**.

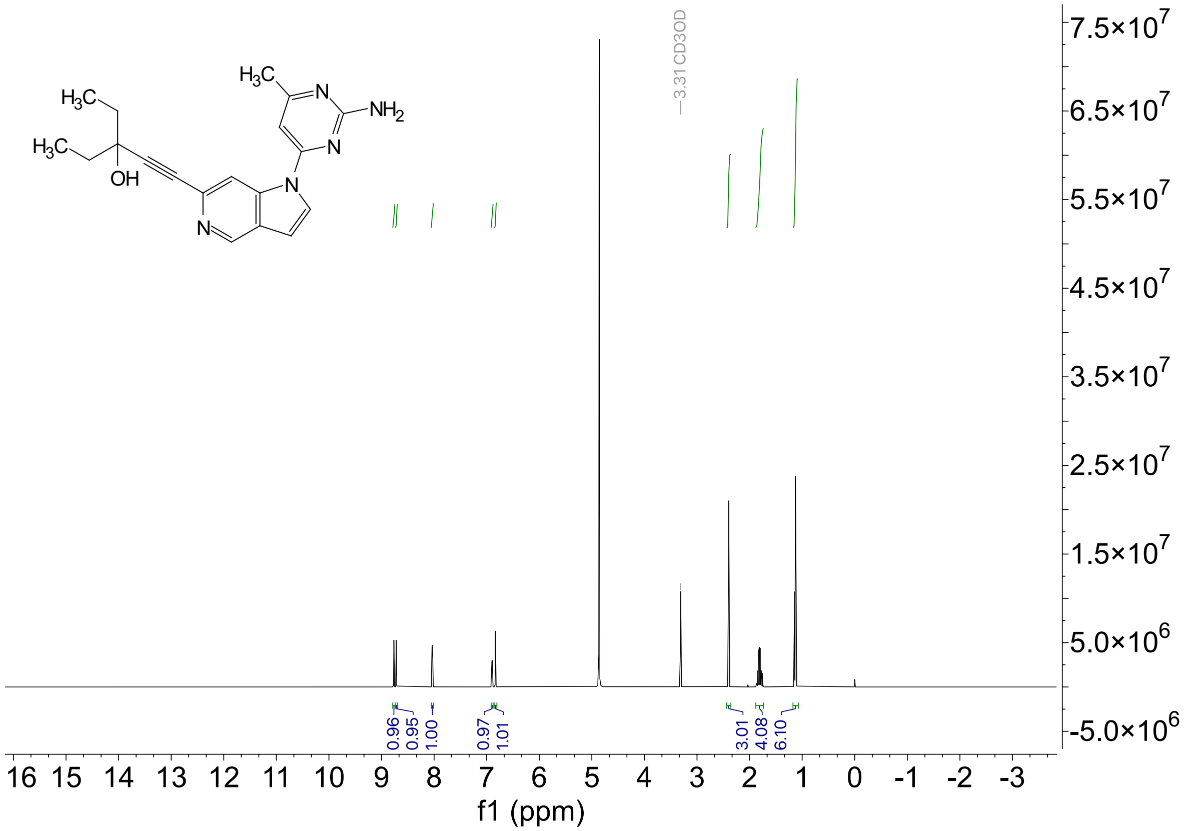

$^{13}$C NMR (126 MHz, $CD_3OD$) for Compound **2.**

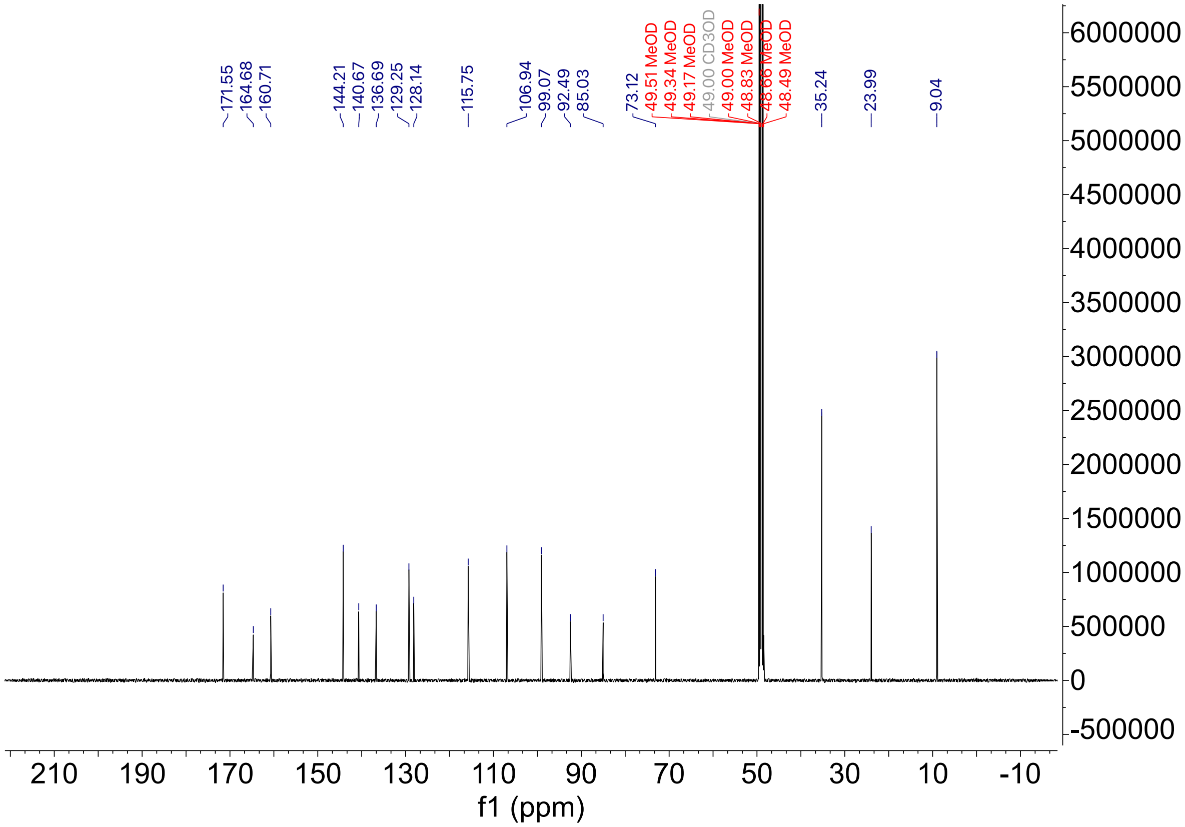

HPLC trace for compound **2**

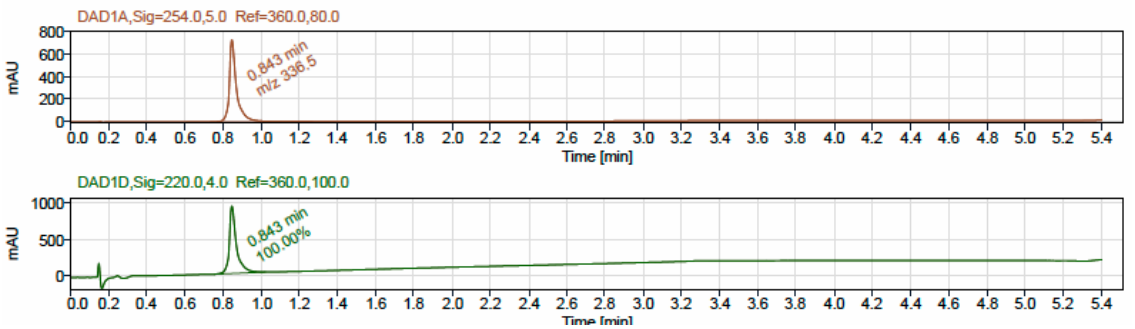


LCMS

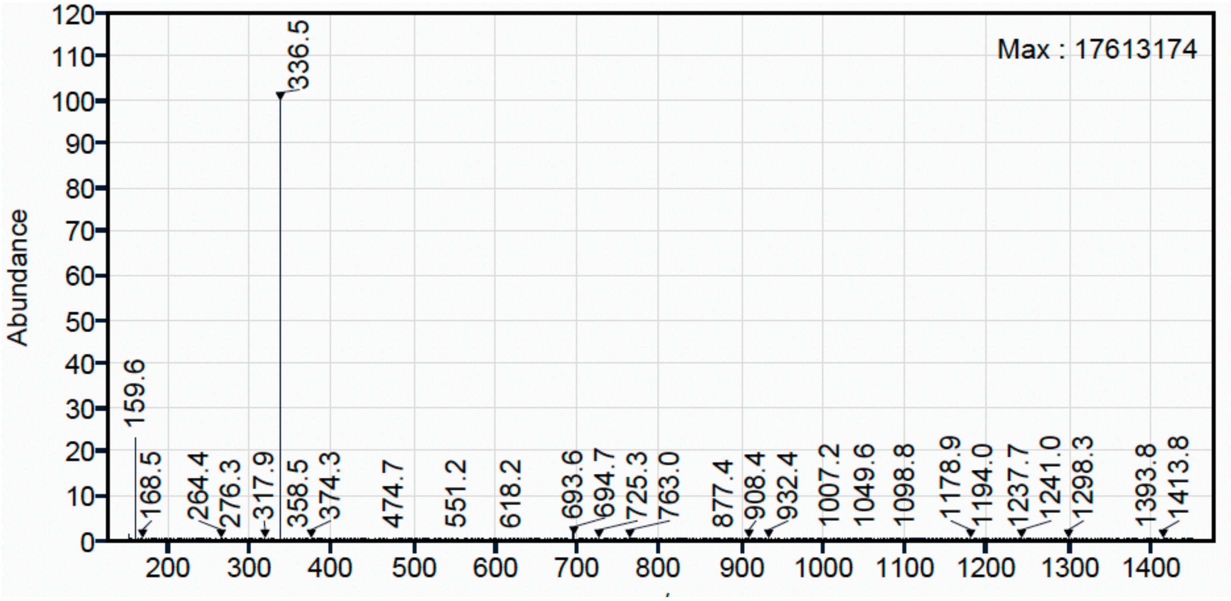

$^{1}H$ NMR (500 MHz, DMSO-$d_6$) for Compound **3**.

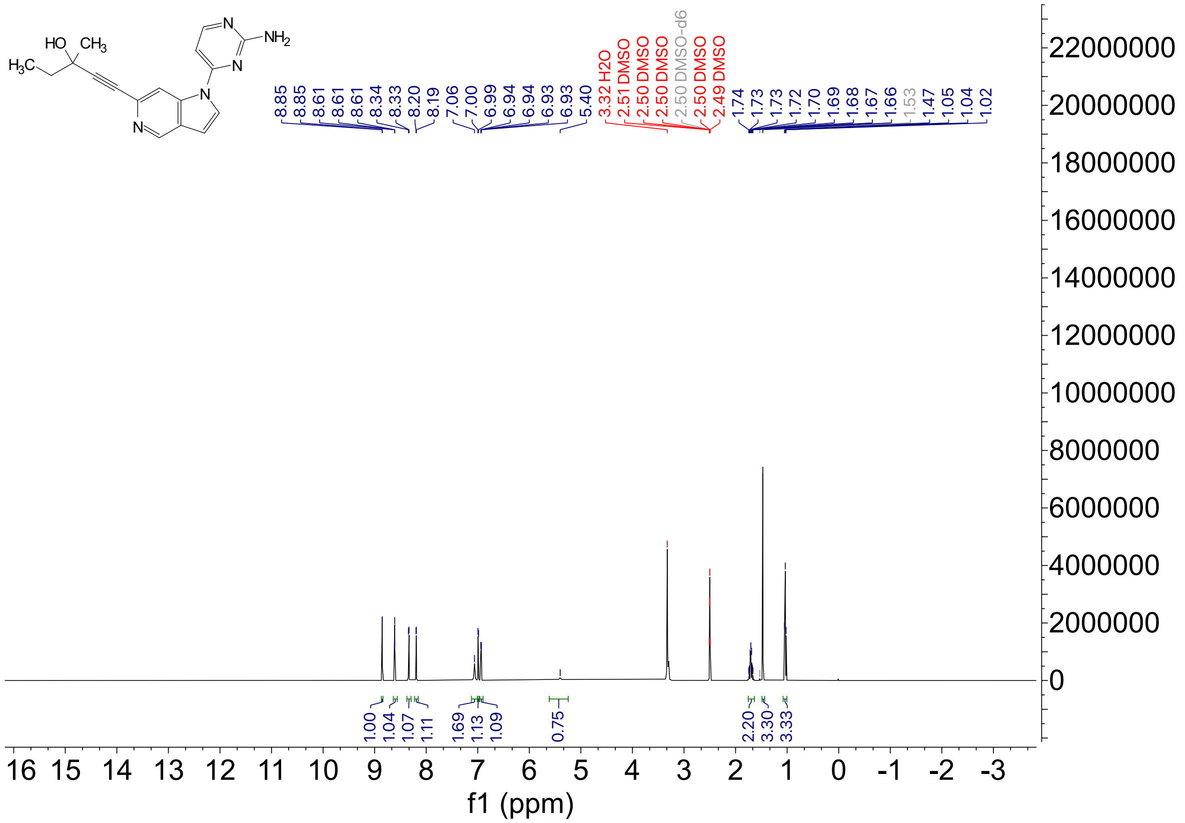

$^{13}C$ NMR (126 MHz, DMSO-$d_6$) for Compound **3.**

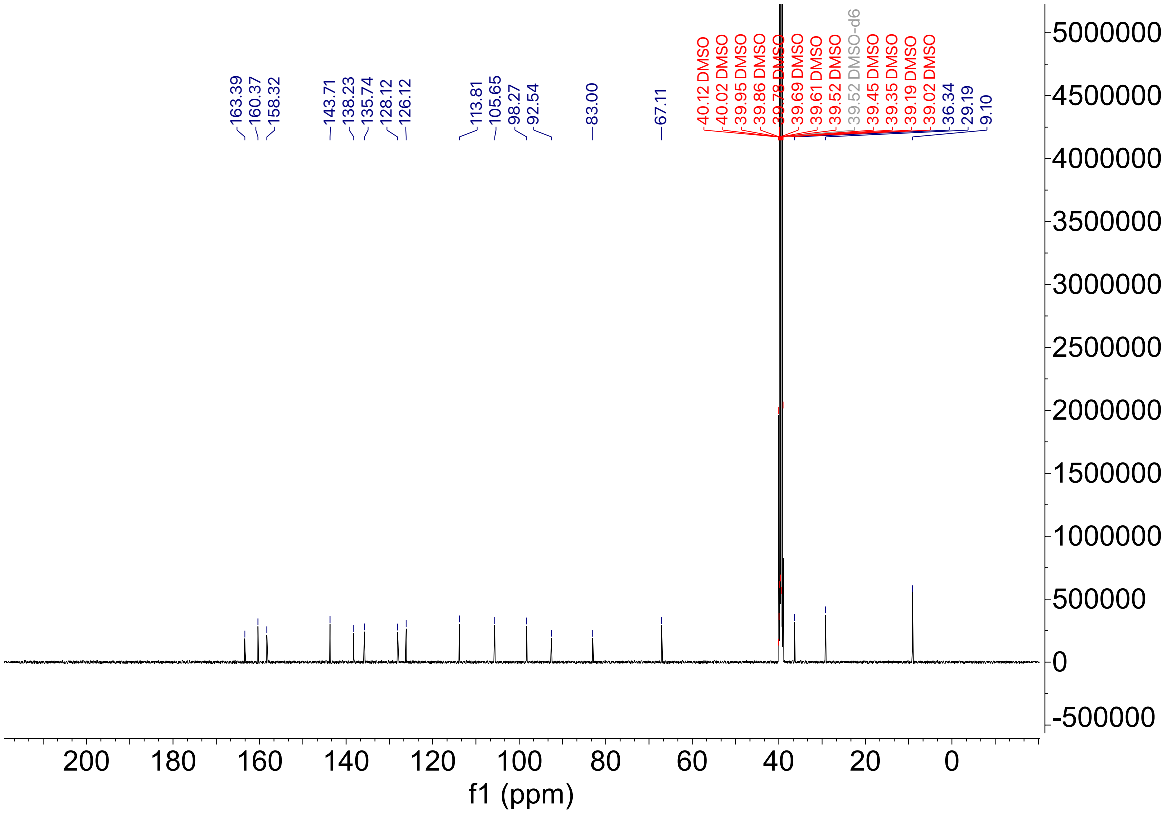

HPLC trace for compound **3**

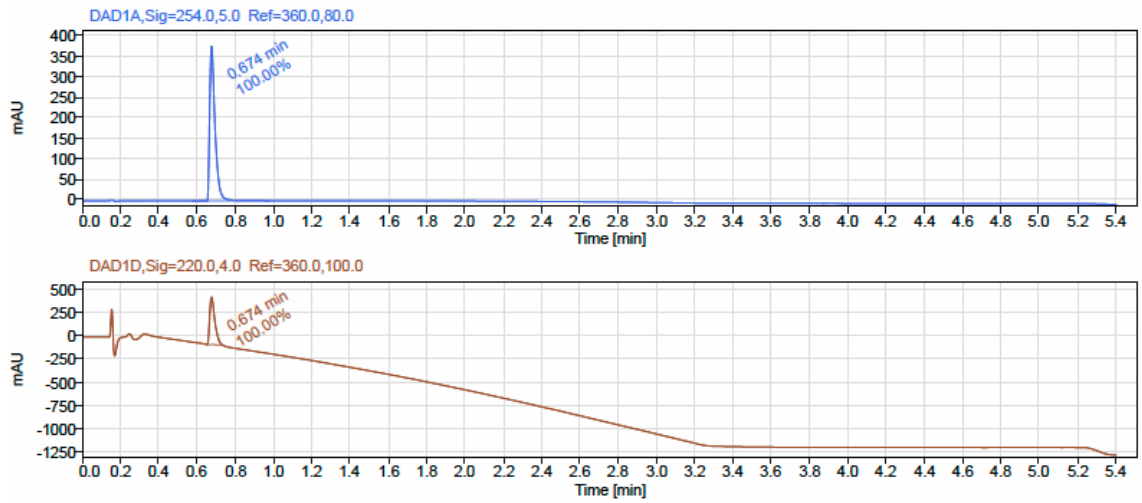


LCMS

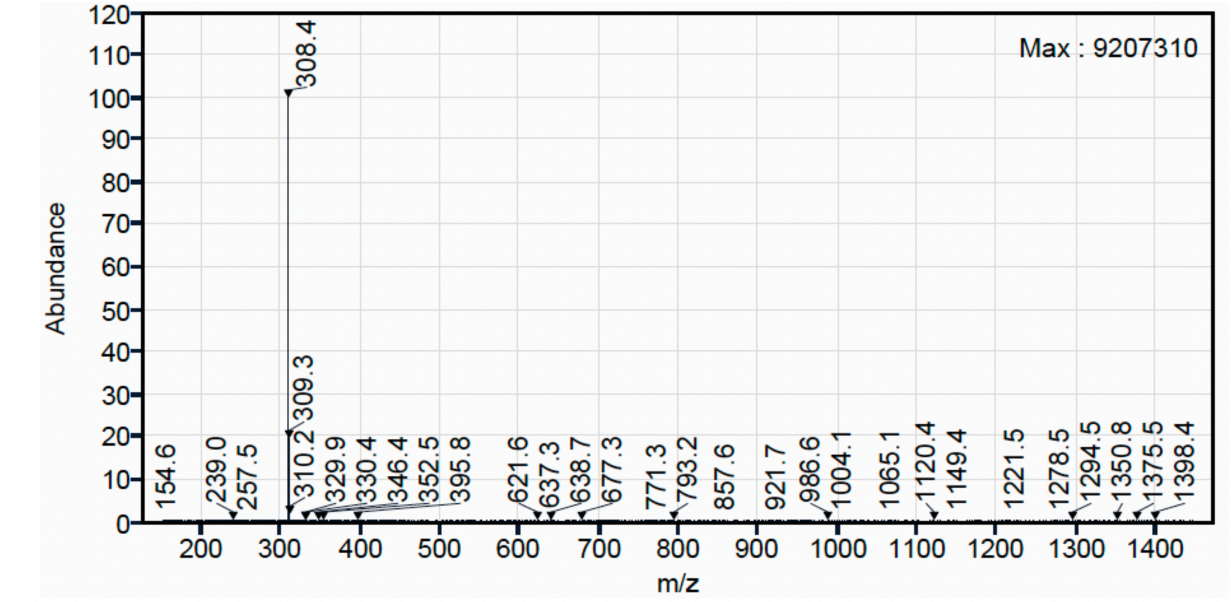

$^1H$ NMR (500 MHz, $CD_3OD$) for Compound **4**.

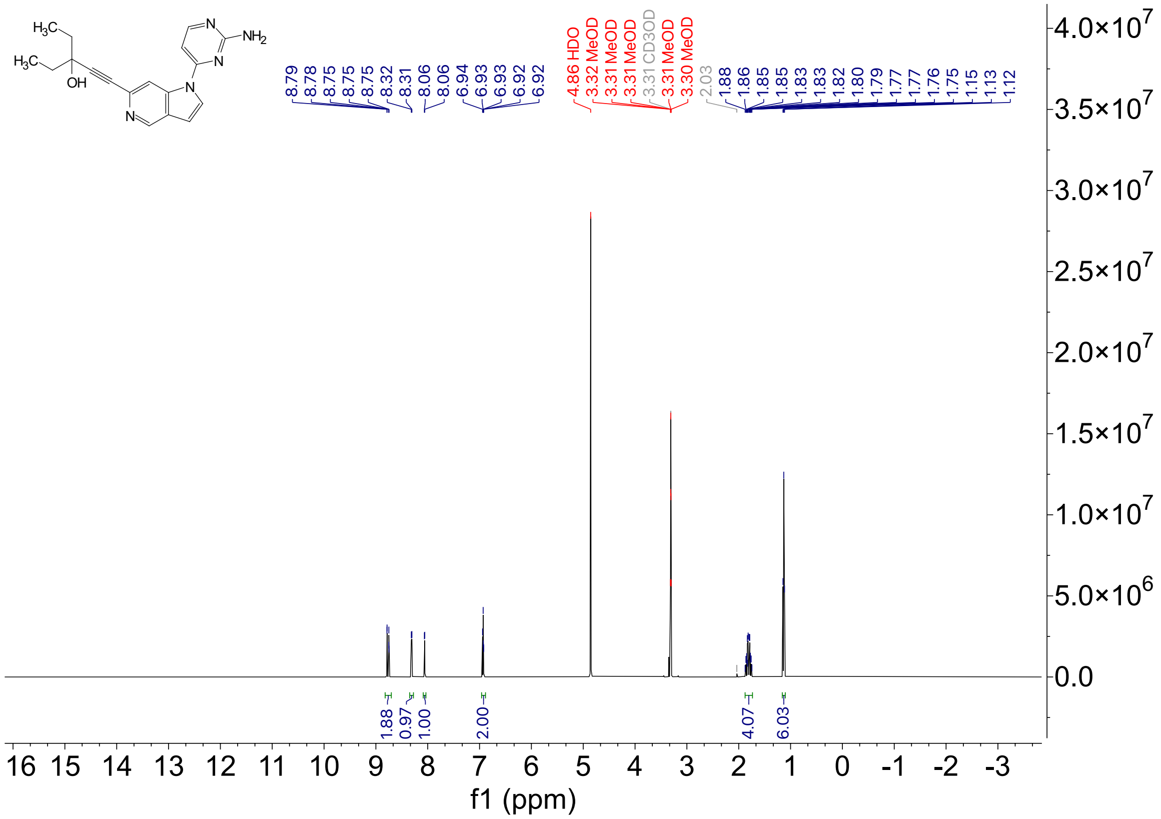

$^{13}C$ NMR (126 MHz, $CD_3OD$) for Compound **4.**

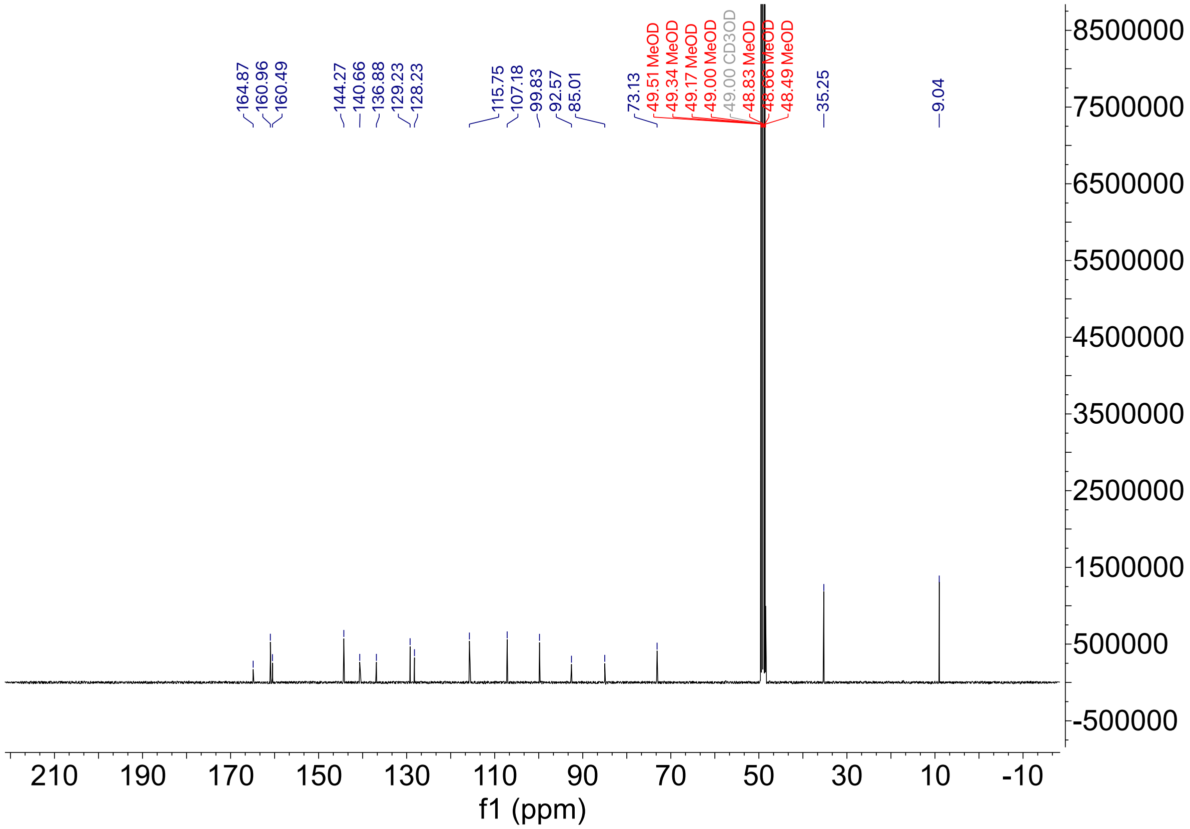

HPLC trace for compound **4**

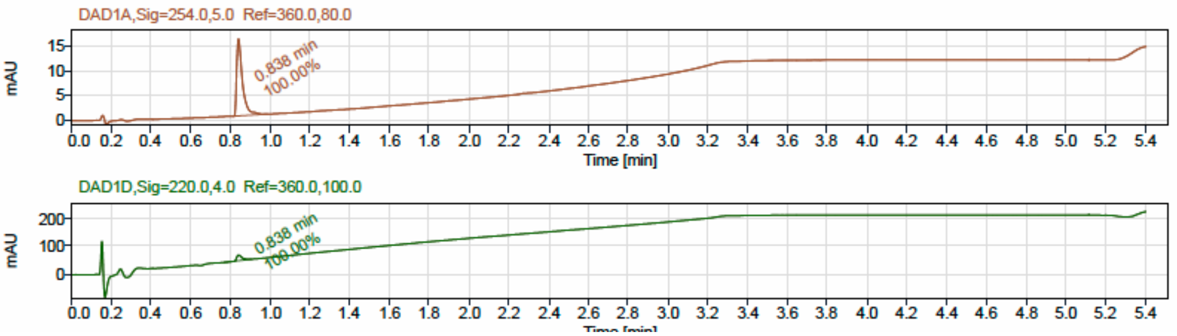


LCMS

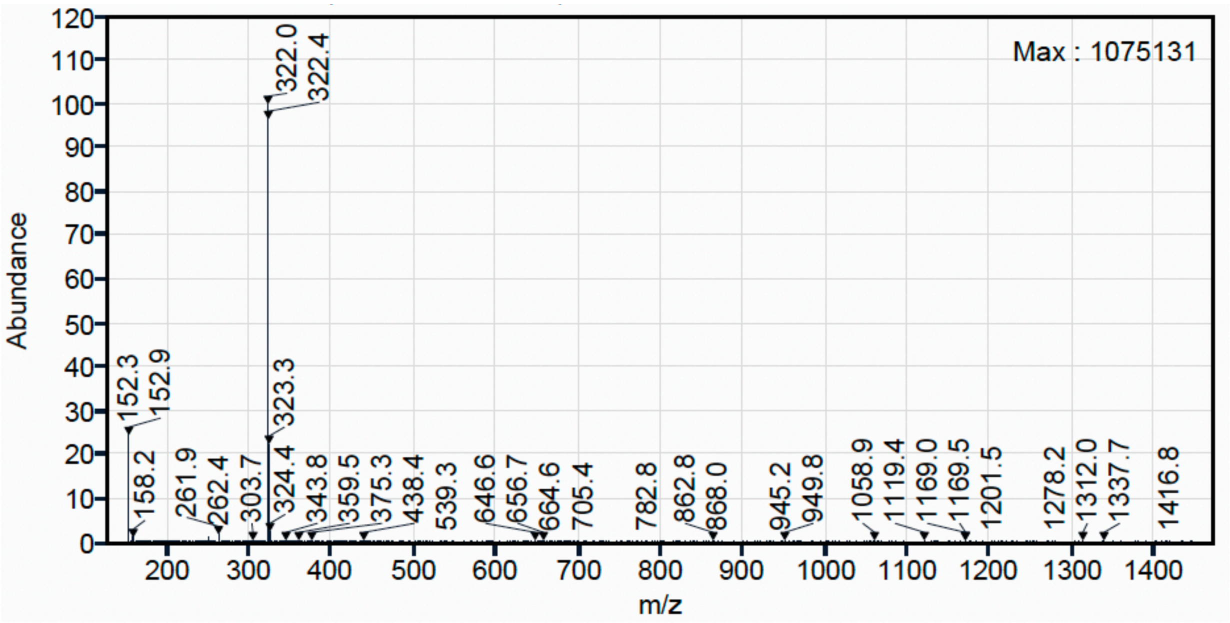

$^1$H NMR (500 MHz, $CD_3OD$) for Compound **5**.

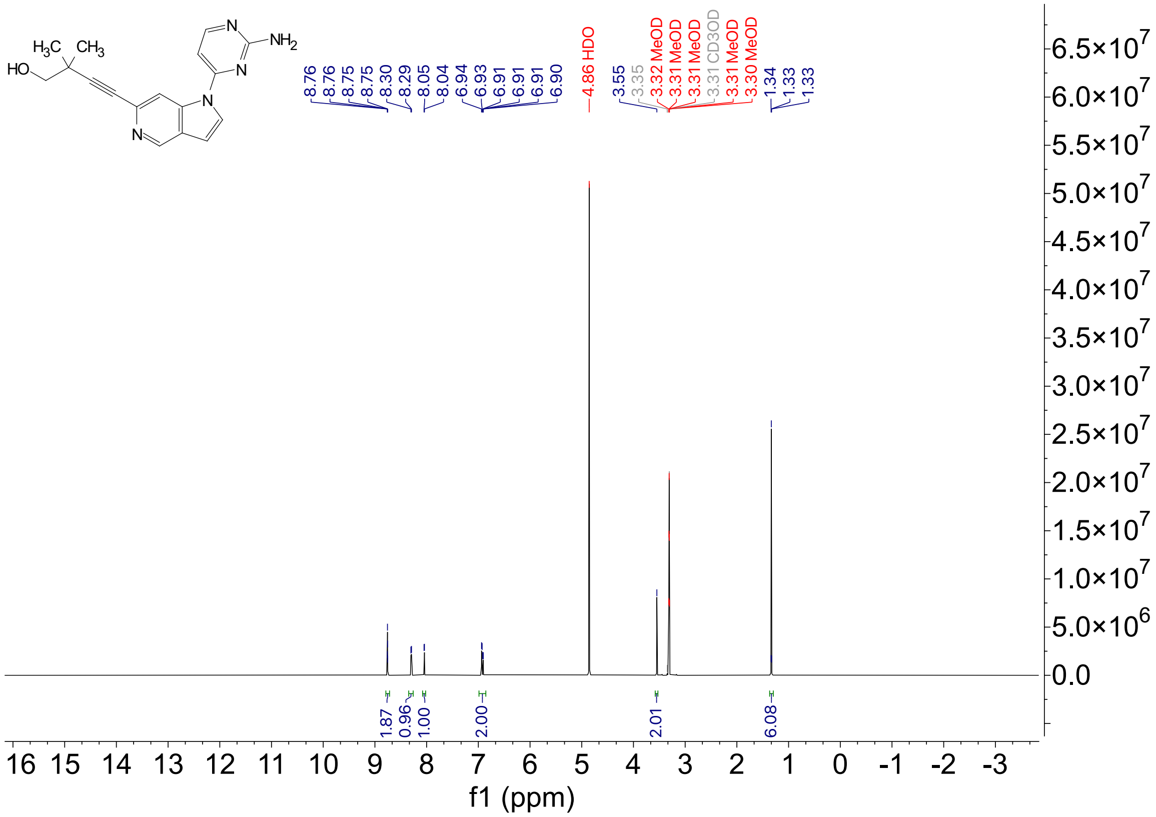

$^{13}C$ NMR (126 MHz, $CD_3OD$) for Compound **5.**

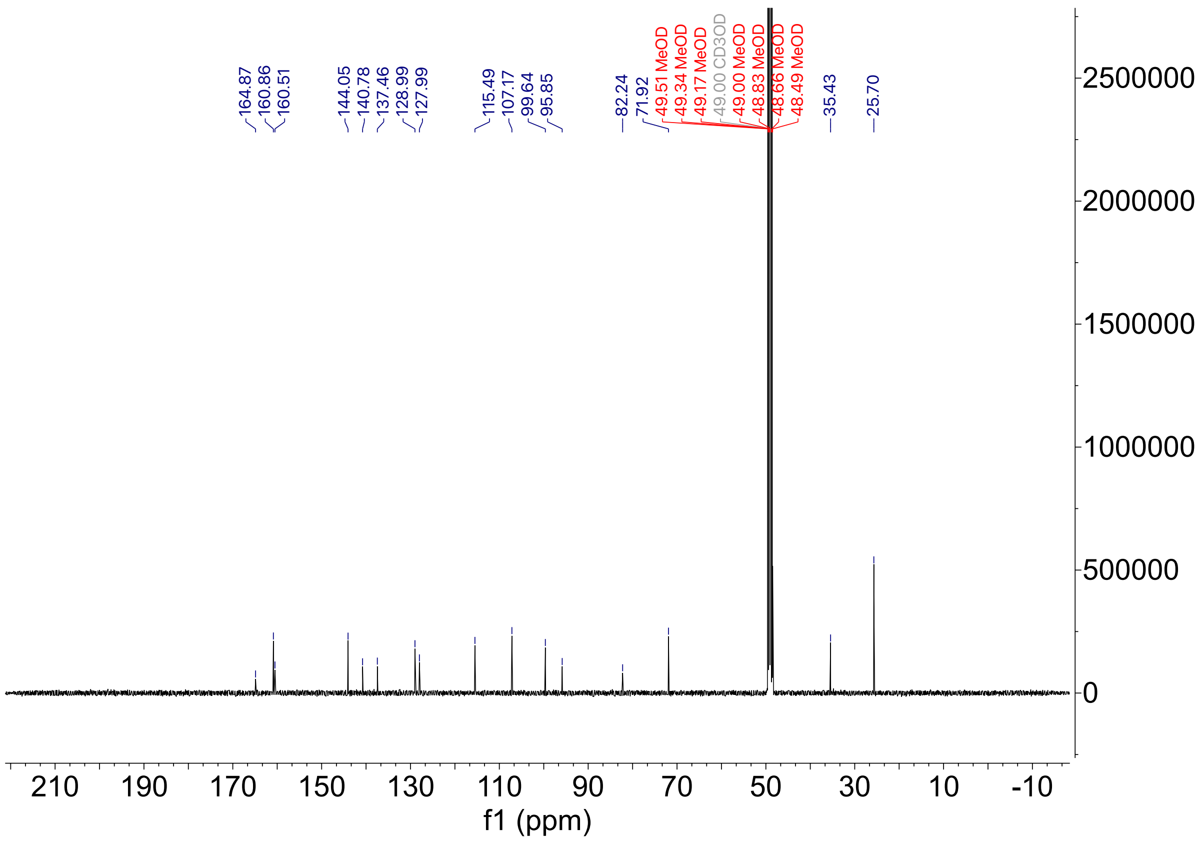

HPLC trace for compound **5**

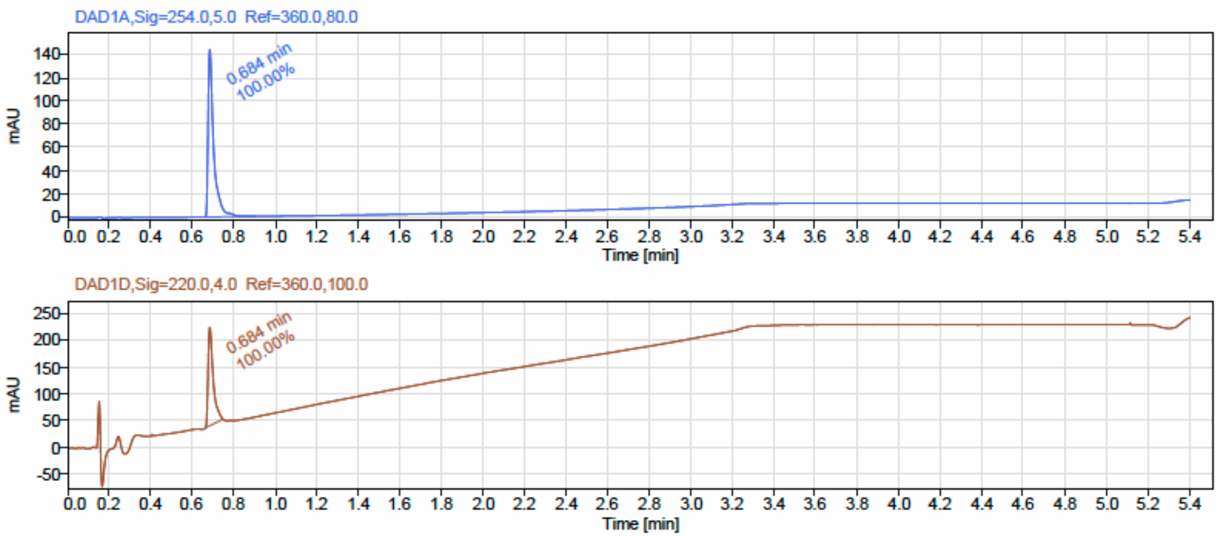


LCMS

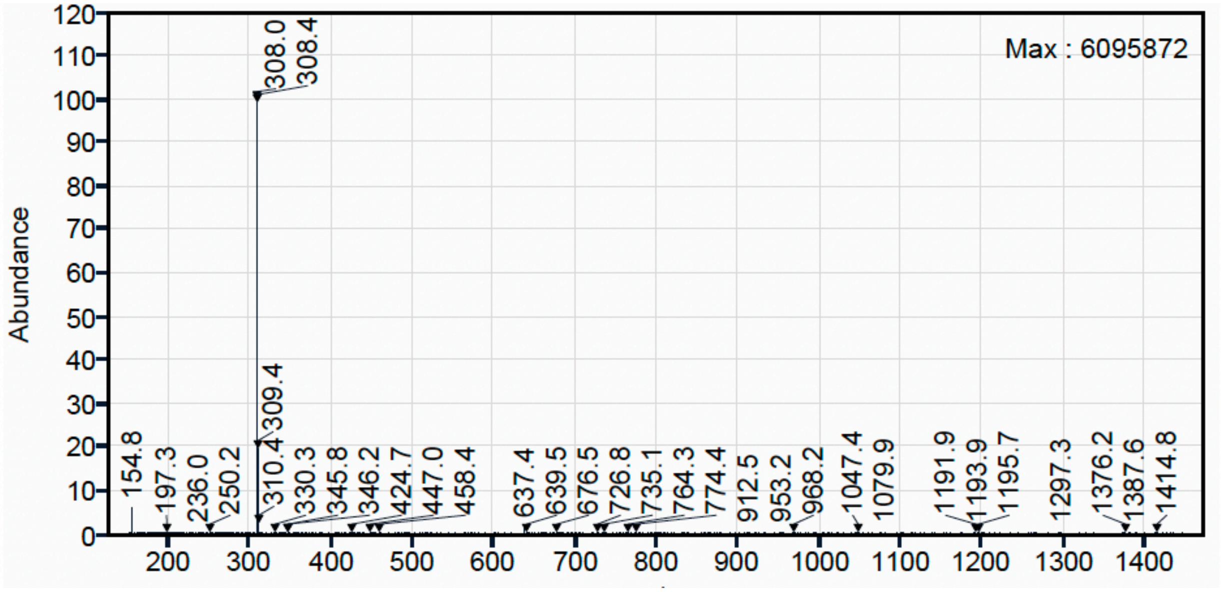

[1]H NMR (500 MHz, $CD_3OD$) for Compound **6**.

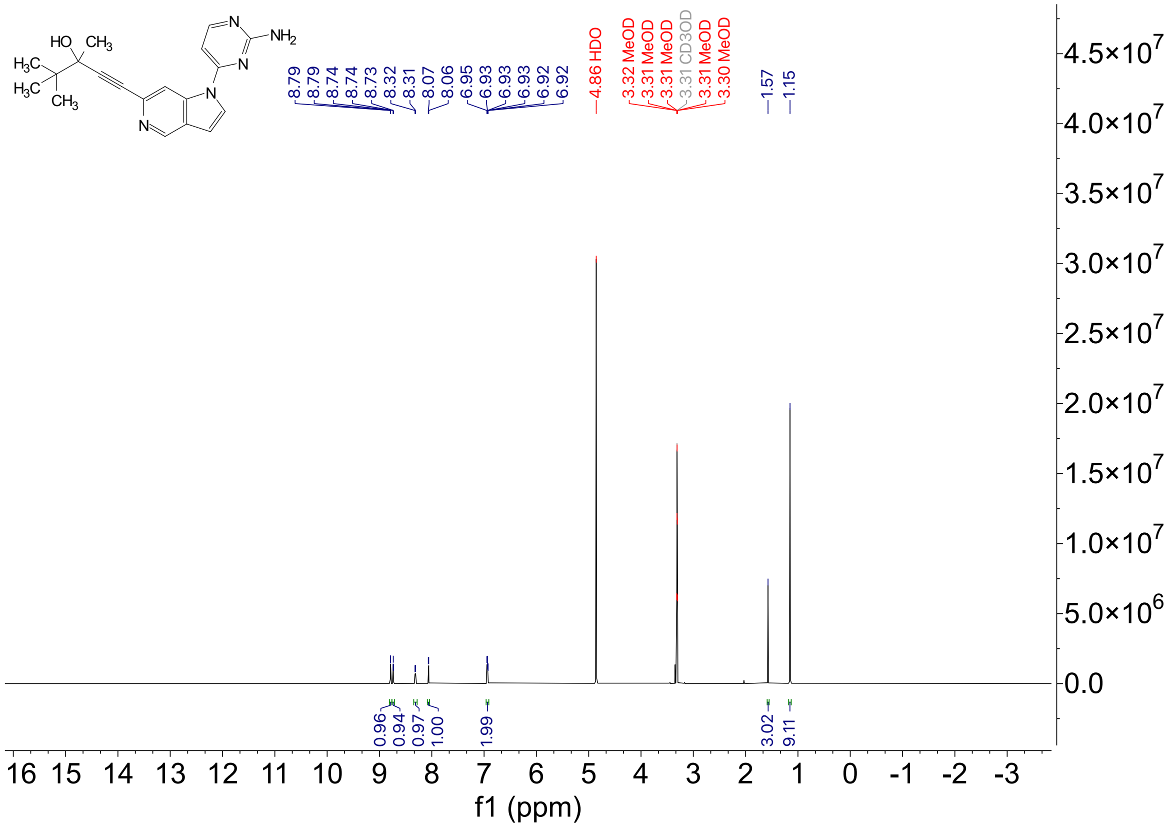

$^{13}C$ NMR (126 MHz, $CD_3OD$) for Compound **6.**

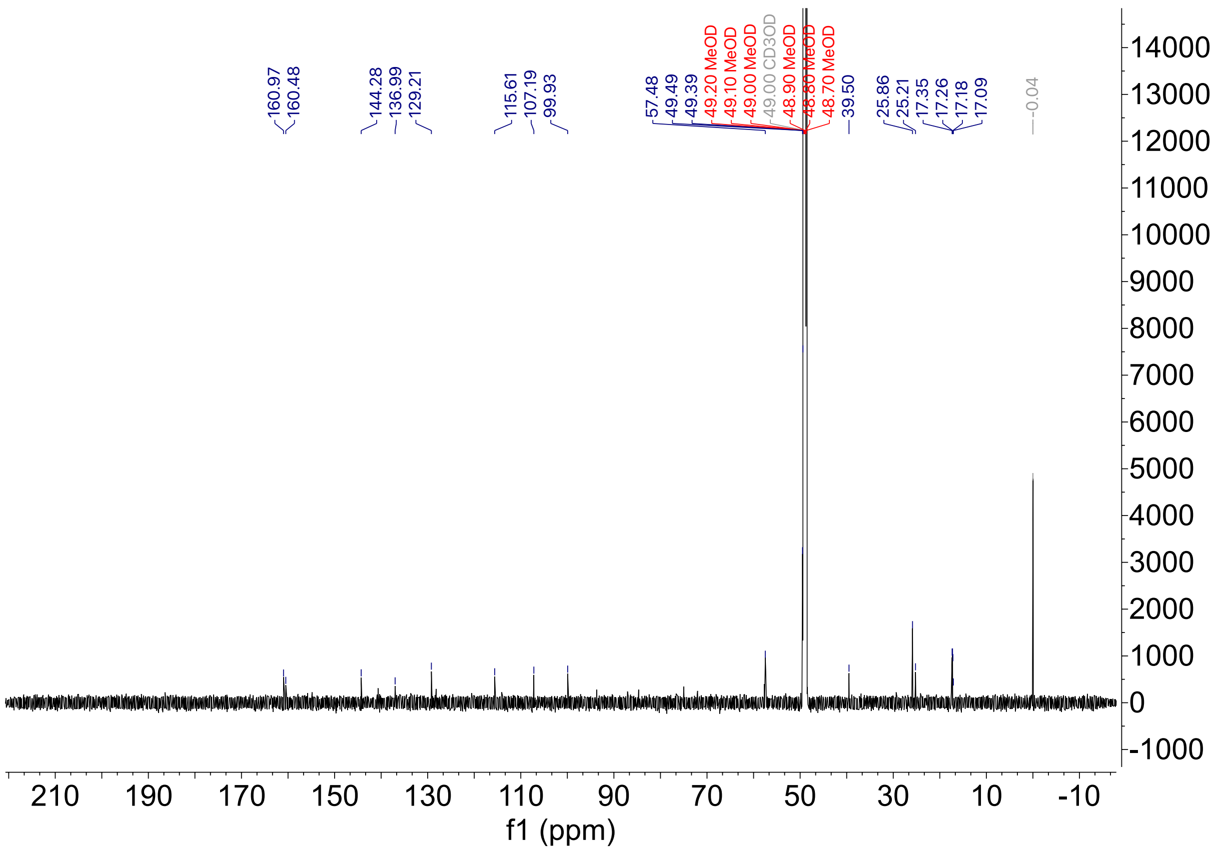

HPLC trace for compound **6**

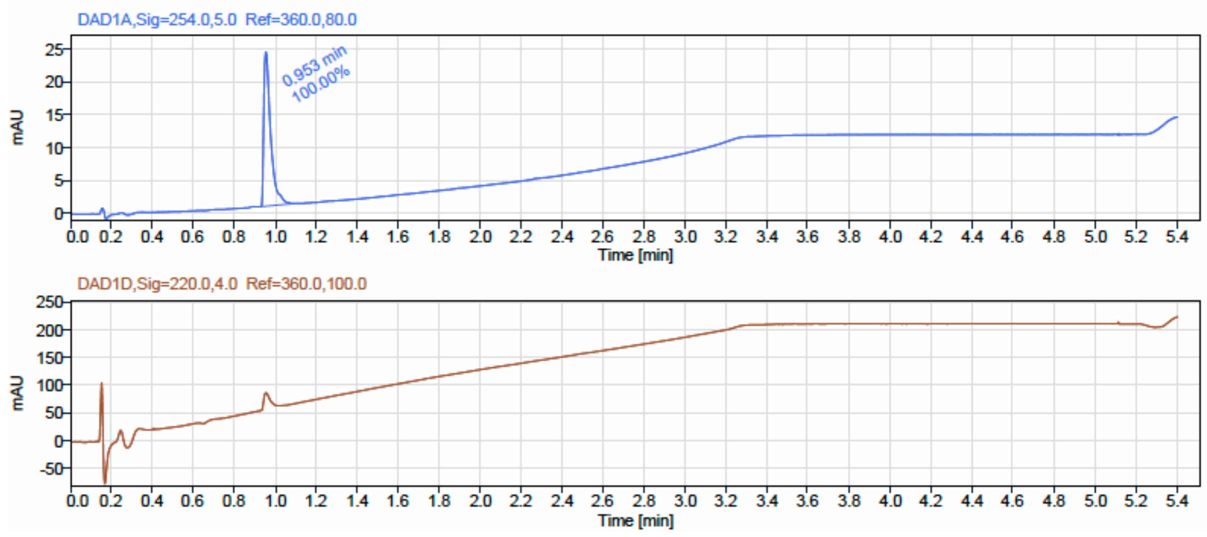


LCMS

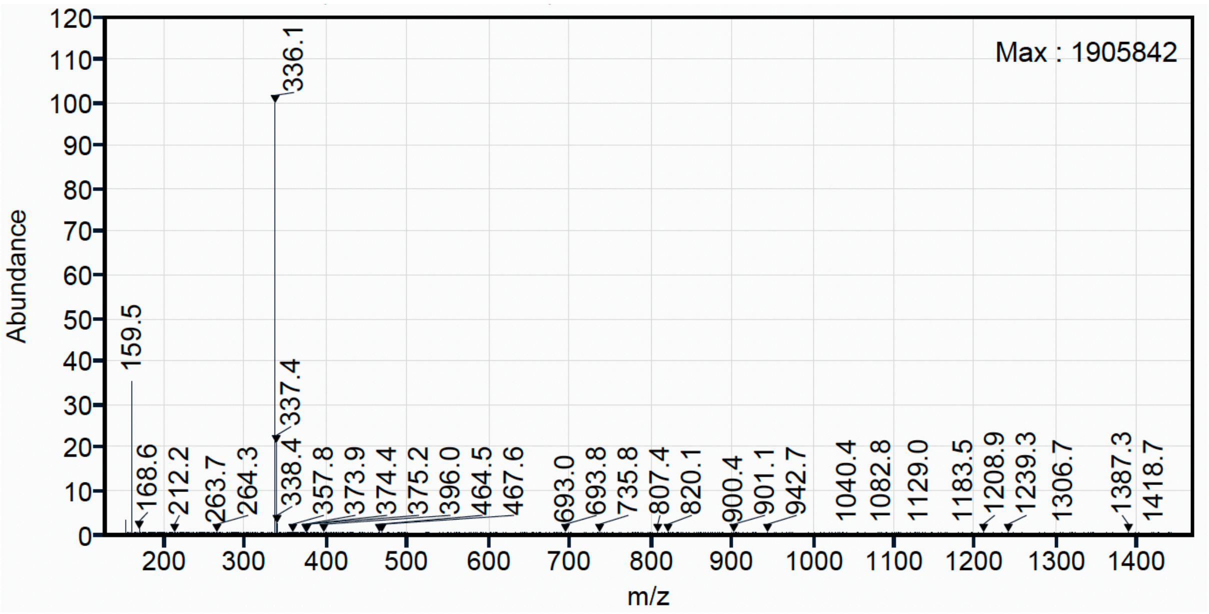

$^{1}H$ NMR (500 MHz, $CD_3OD$) for Compound **7**.

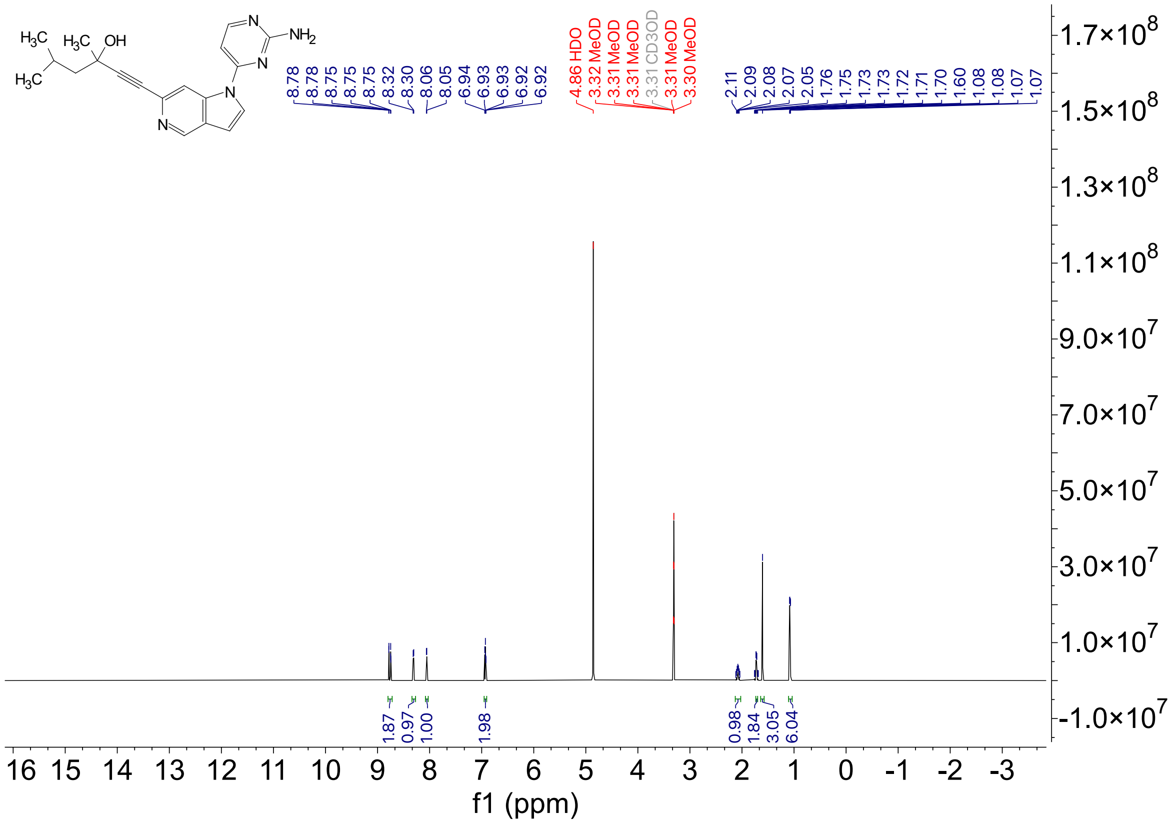

$^{13}C$ NMR (126 MHz, $CD_3OD$) for Compound **7.**

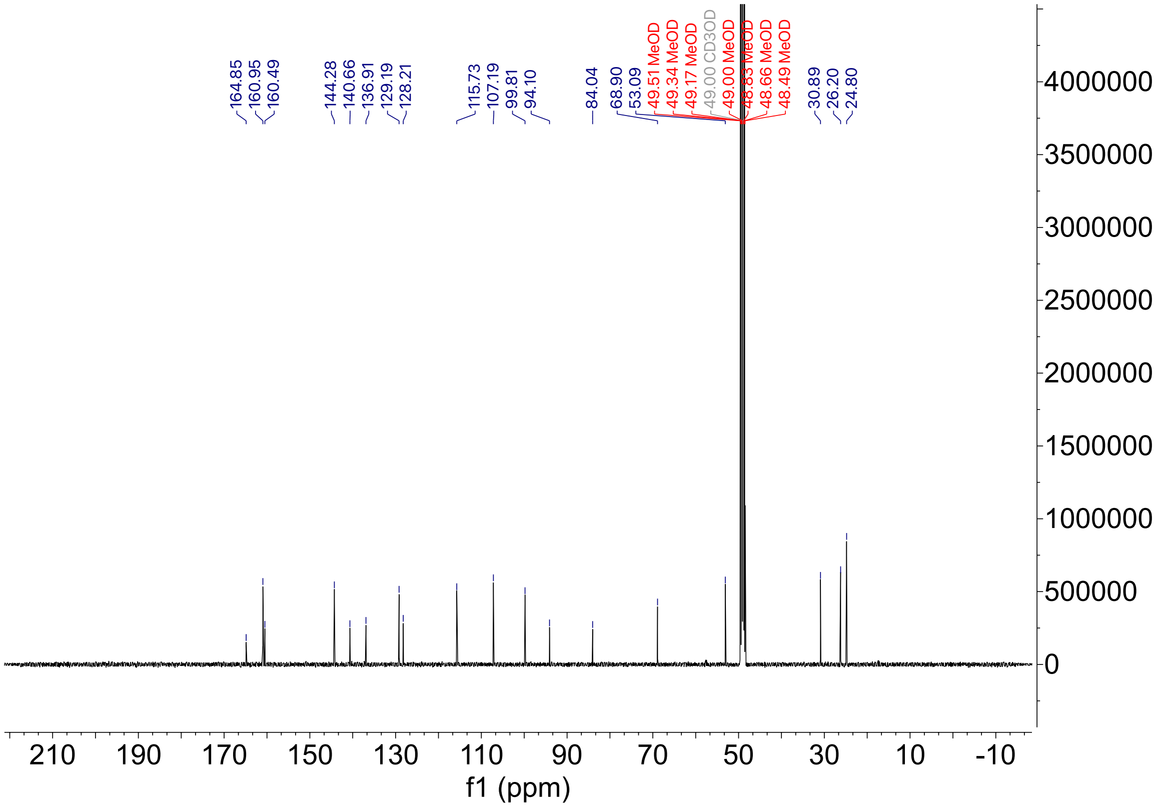

HPLC trace for compound **7**

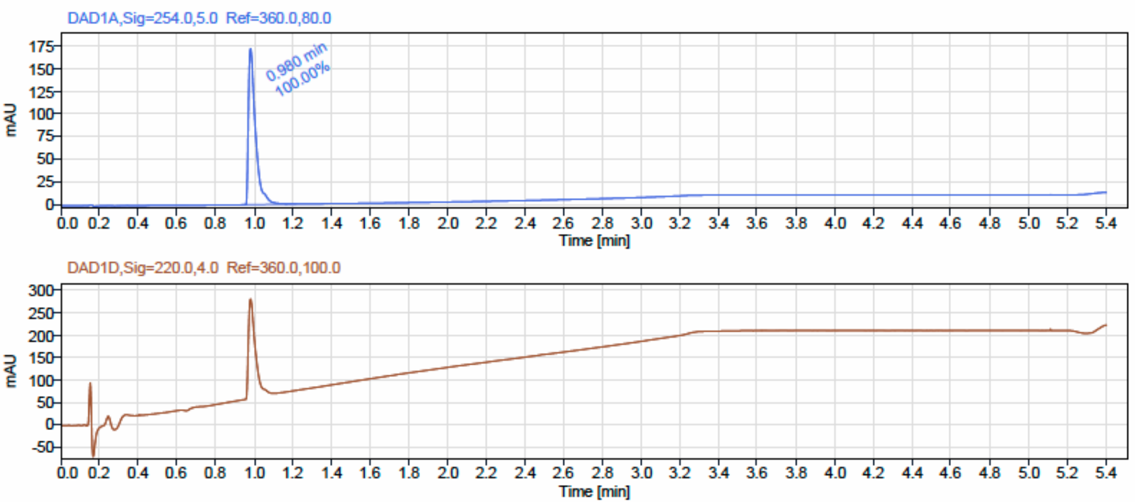


LCMS

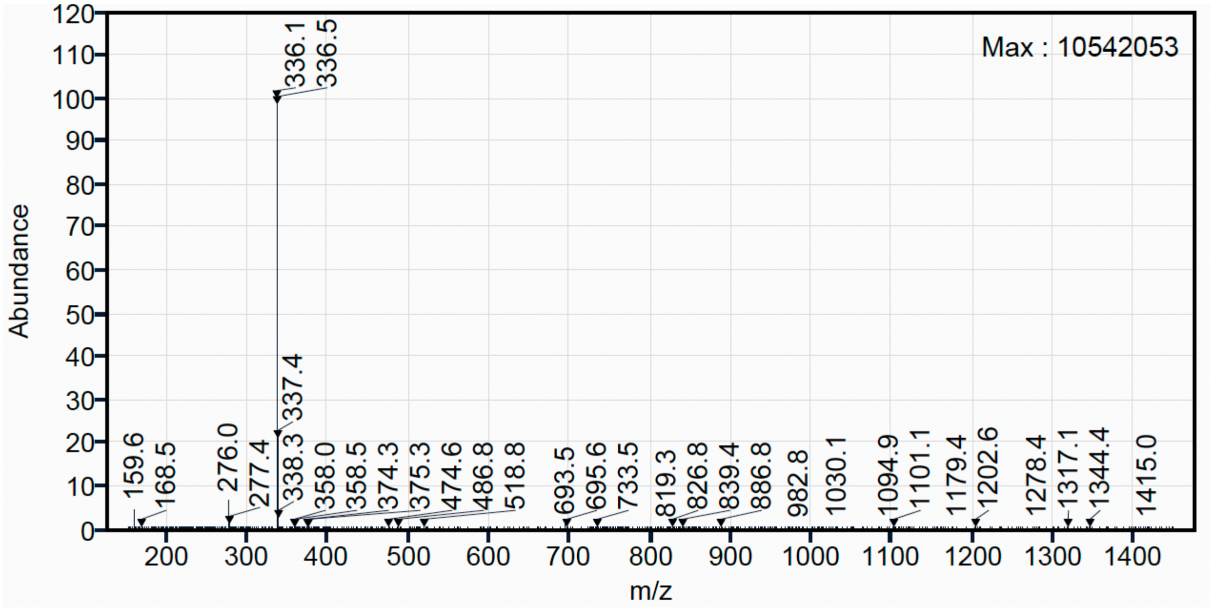

$^1H$ NMR (500 MHz, $CD_3OD$) for Compound **8**.

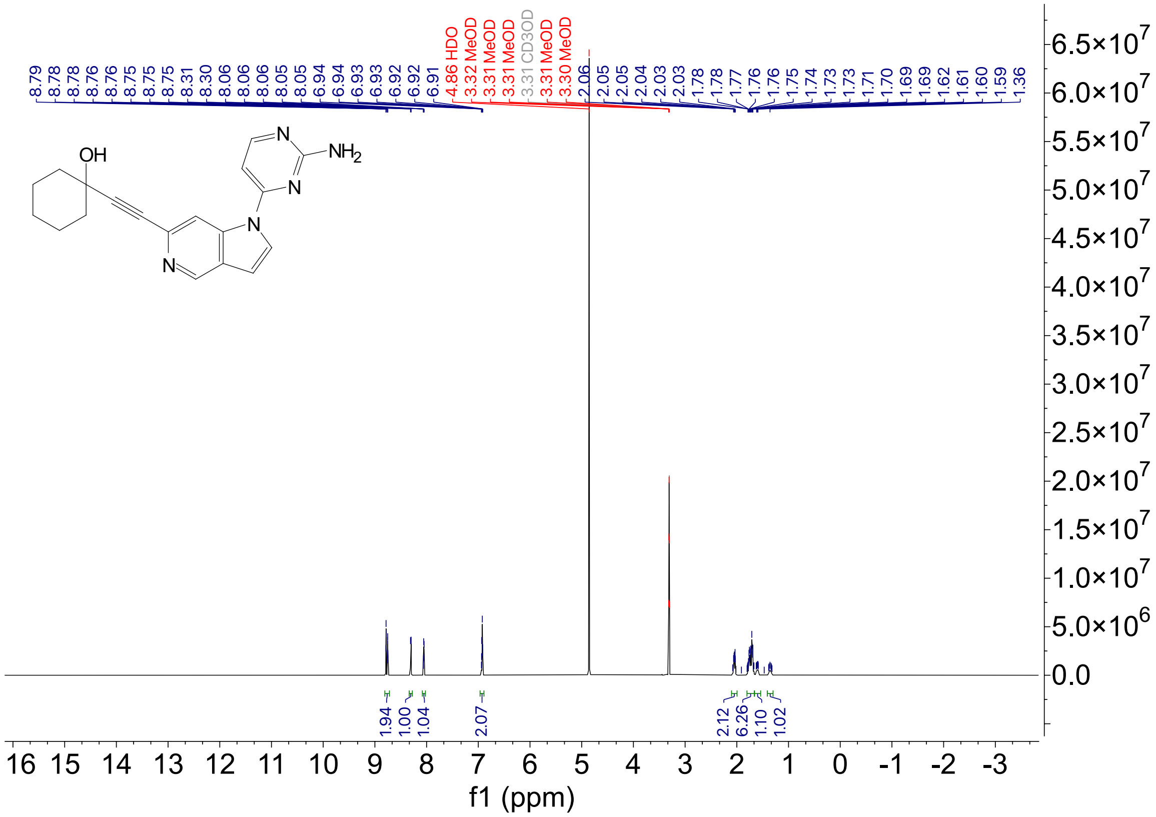

$^{13}C$ NMR (126 MHz, $CD_3OD$) for Compound **8.**

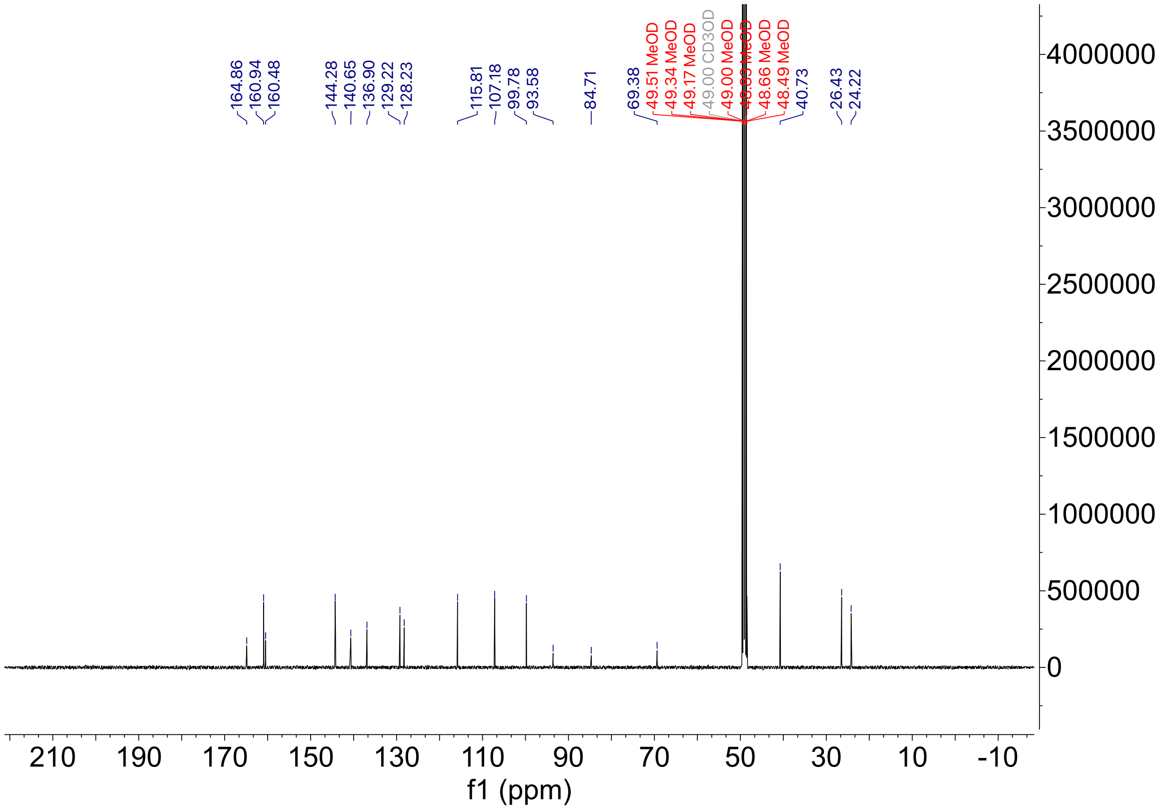

HPLC trace for compound **8**

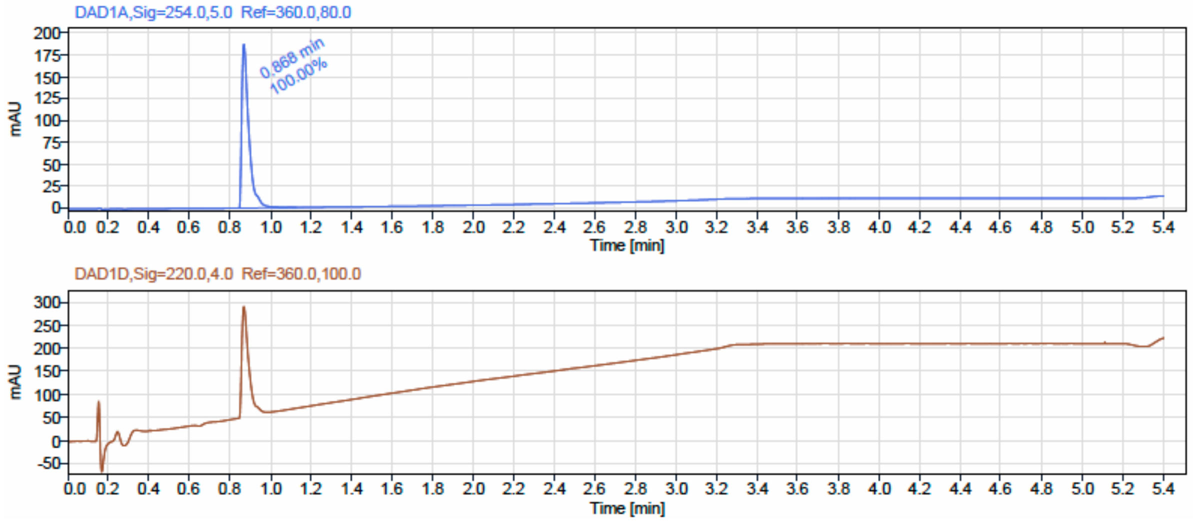


LCMS

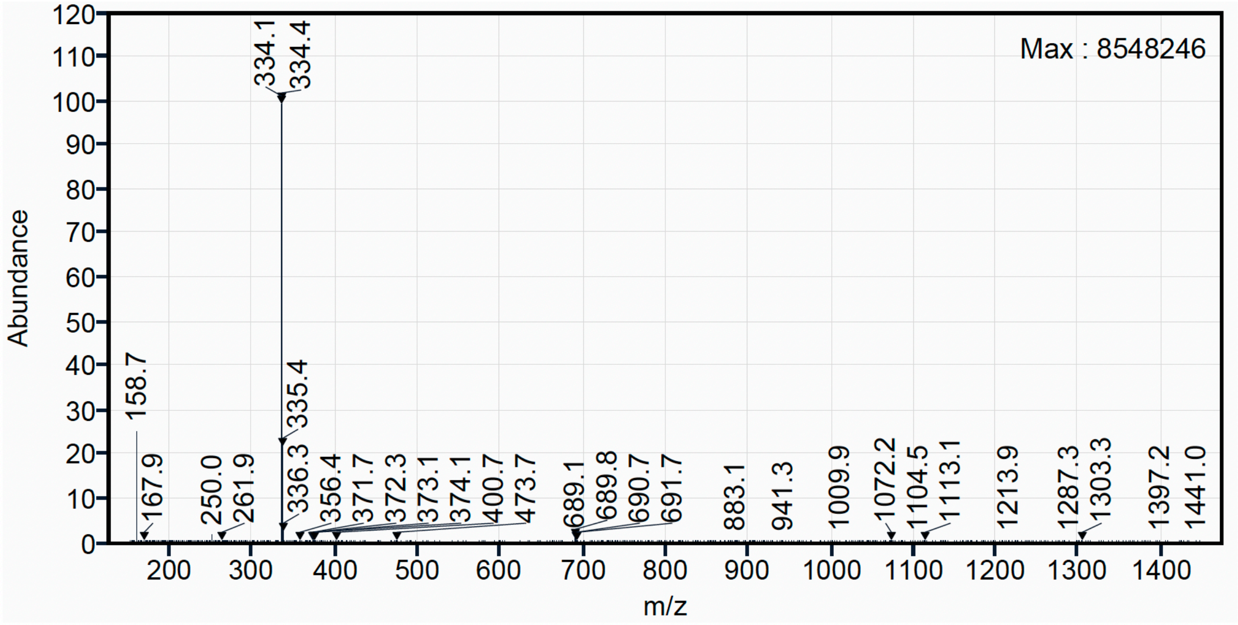

$^{1}H$ NMR (500 MHz, $CD_3OD$) for Compound **9.**

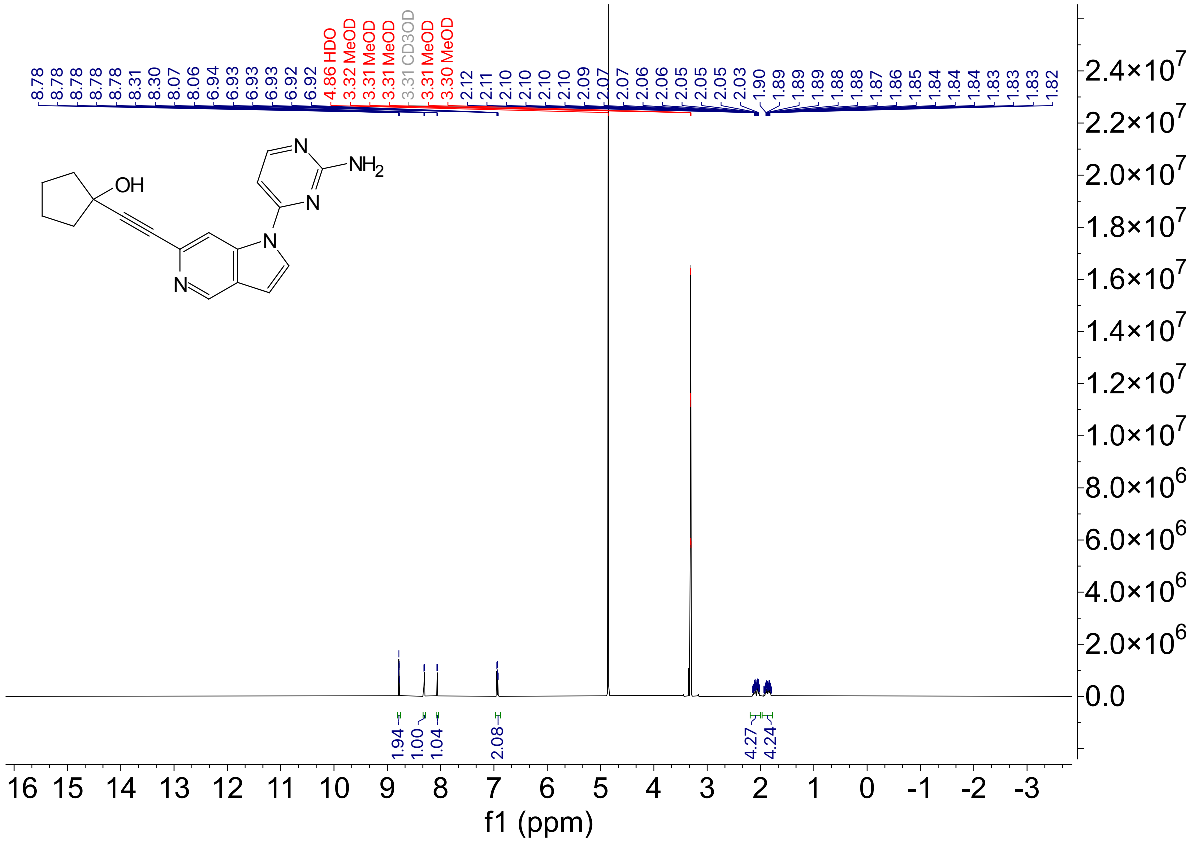

$^{13}C$ NMR (126 MHz, $CD_3OD$) for Compound **9.**

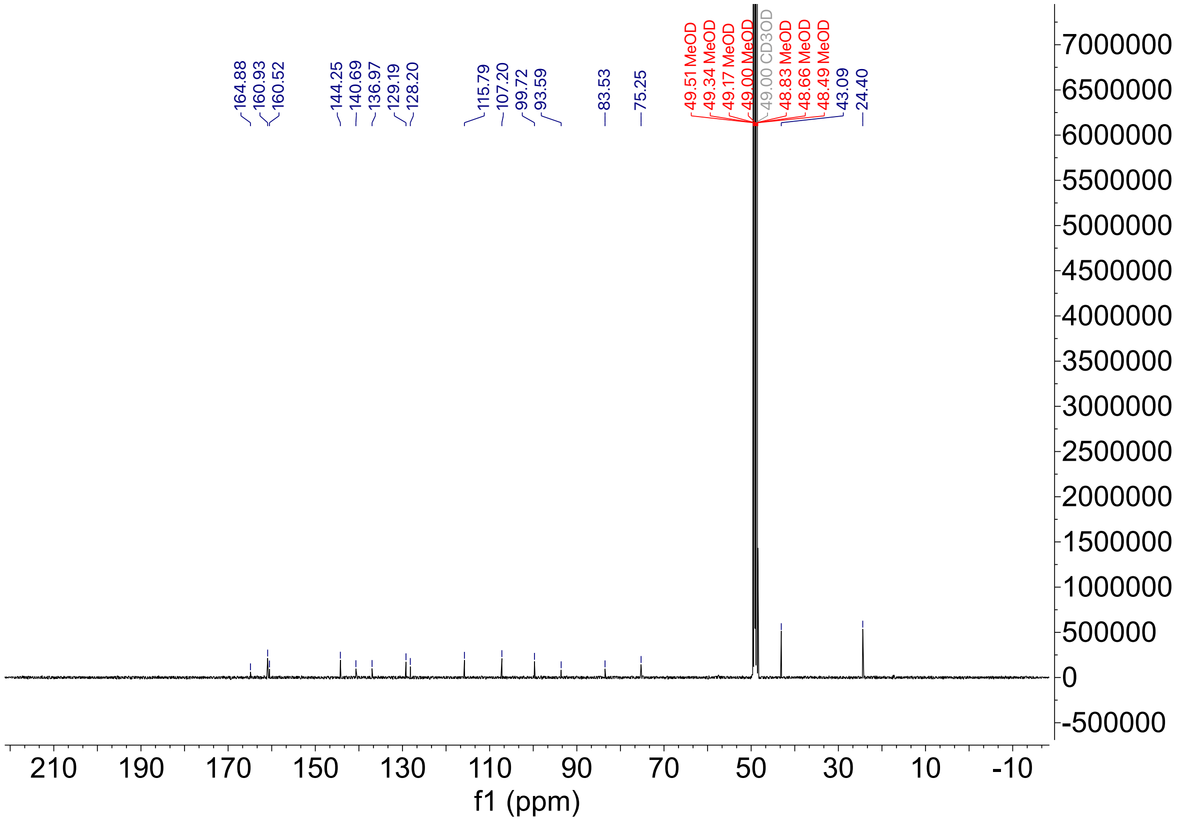

HPLC trace for compound **9**

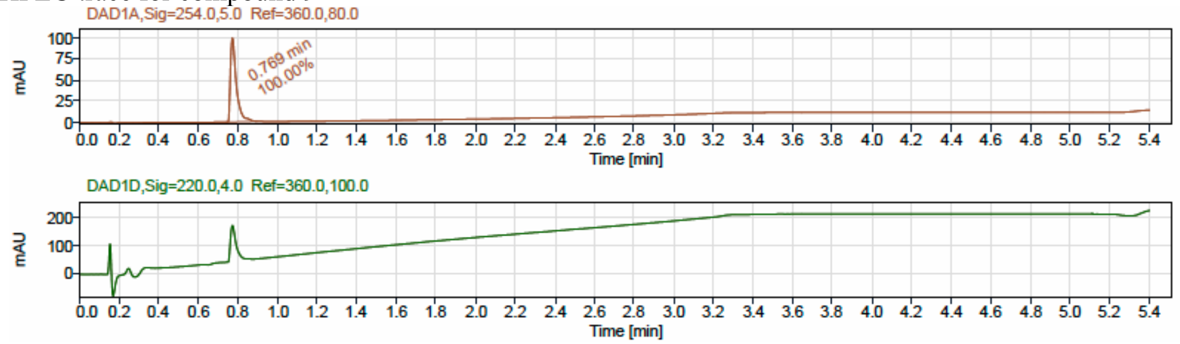


LCMS

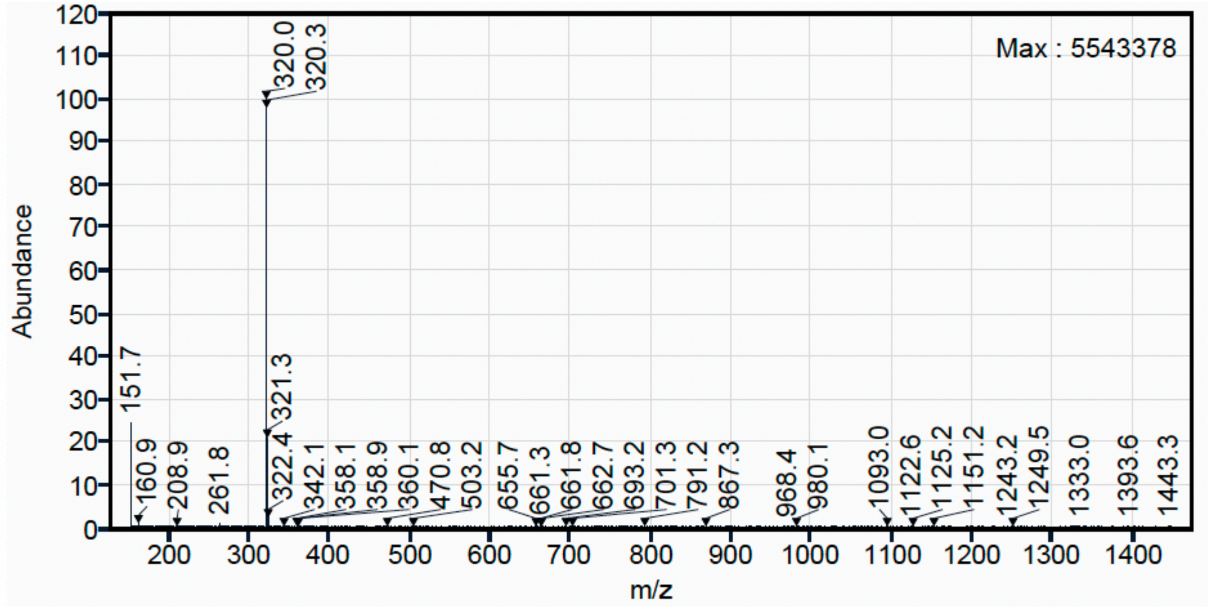

[1]H NMR (500 MHz, $CD_3OD$) for Compound **10.**

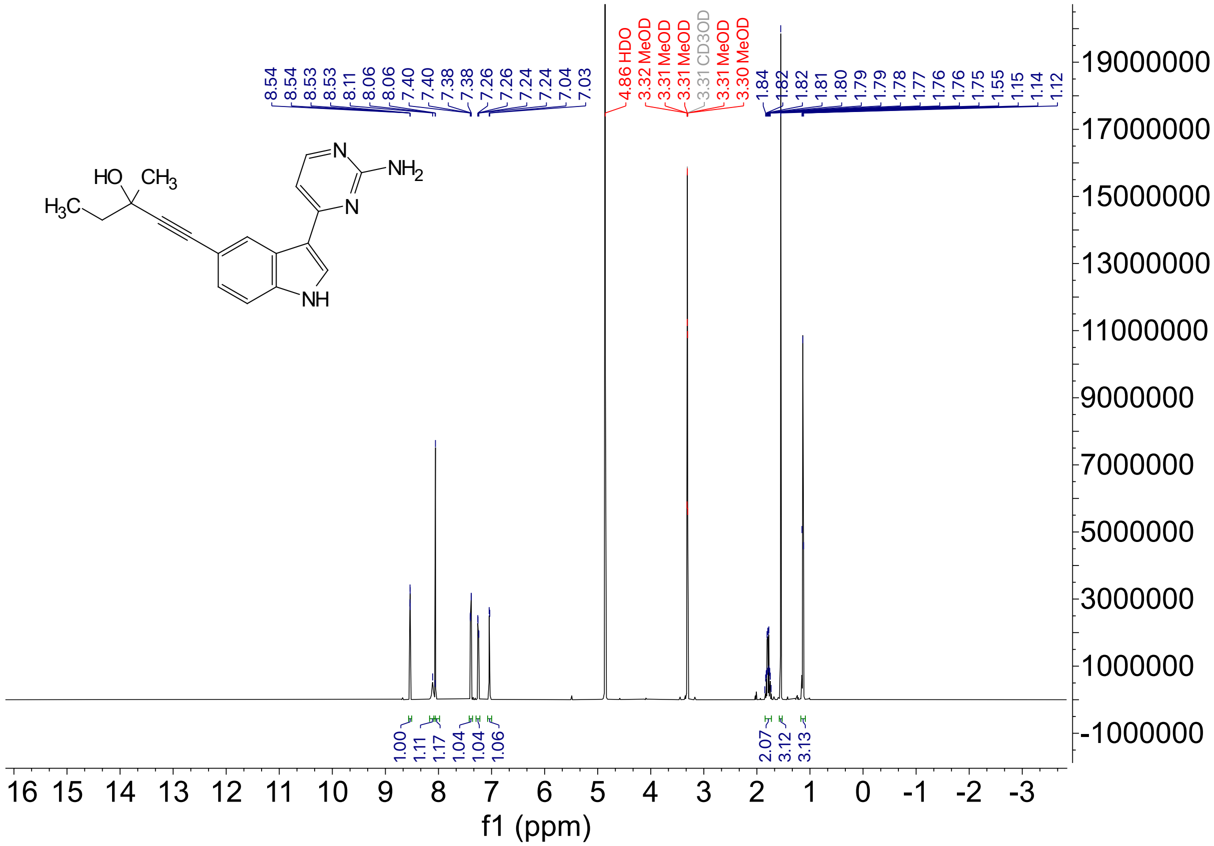

$^{13}C$ NMR (126 MHz, $CD_3OD$) for Compound **10**.

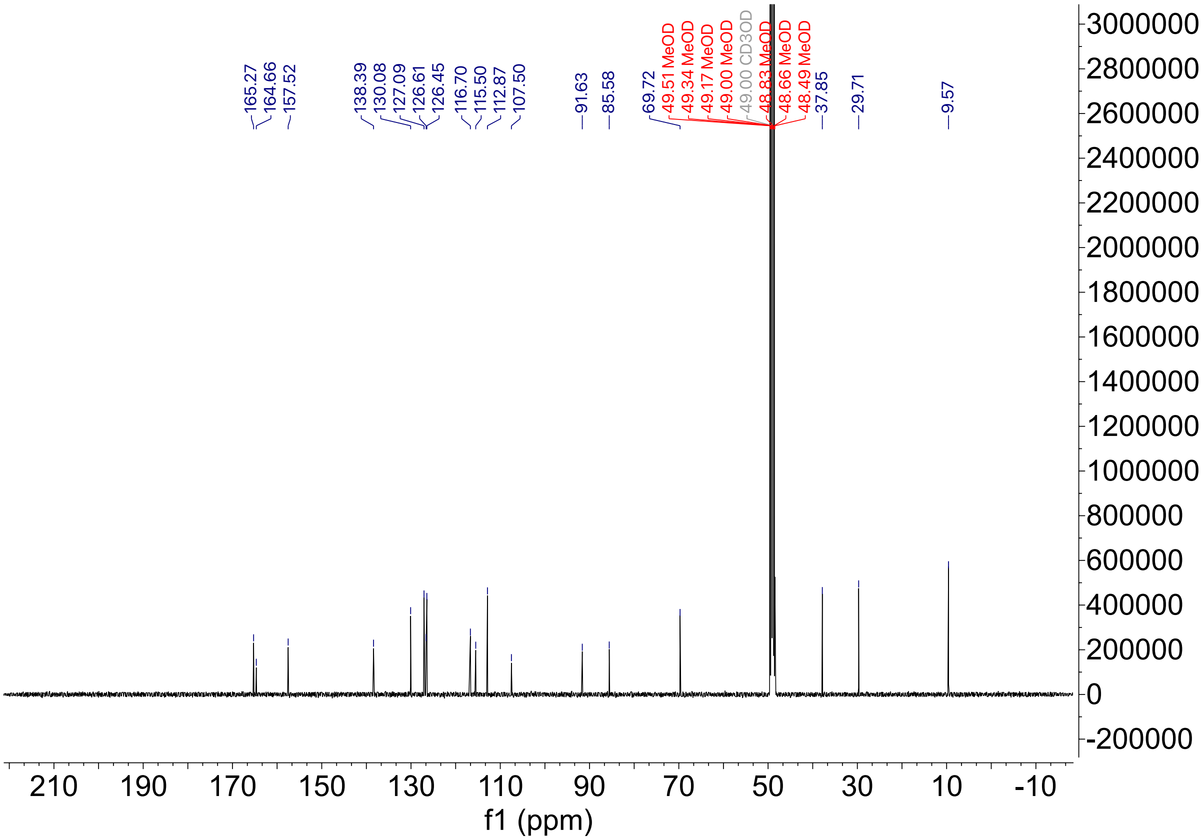

HPLC trace for compound **10**

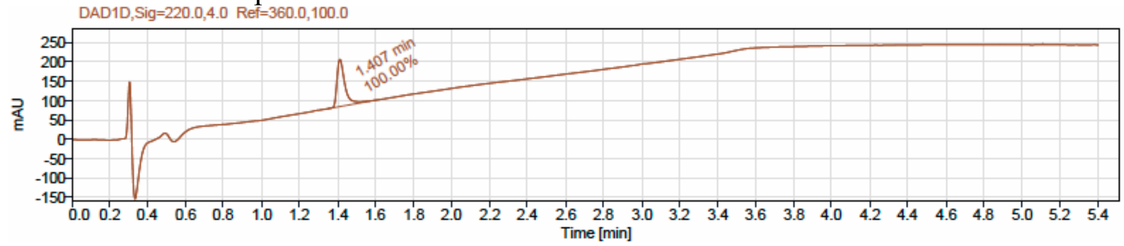


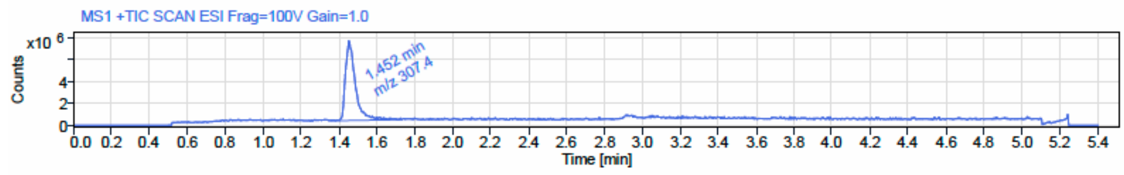


LCMS

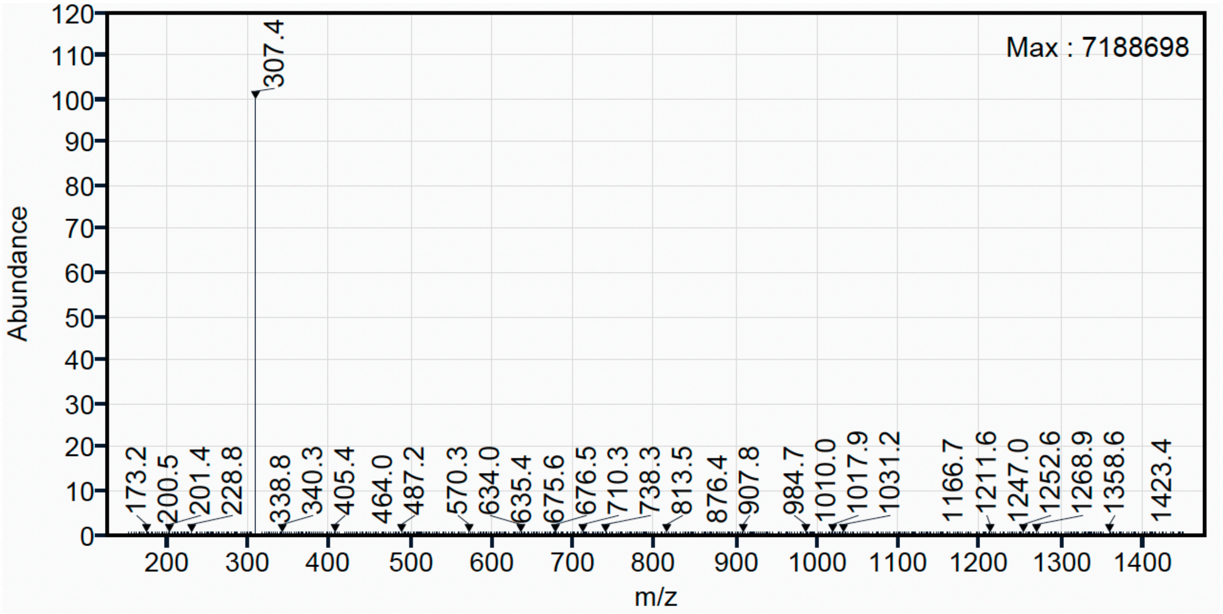

[1]H NMR (500 MHz, $CD_3OD$) for Compound **11.**

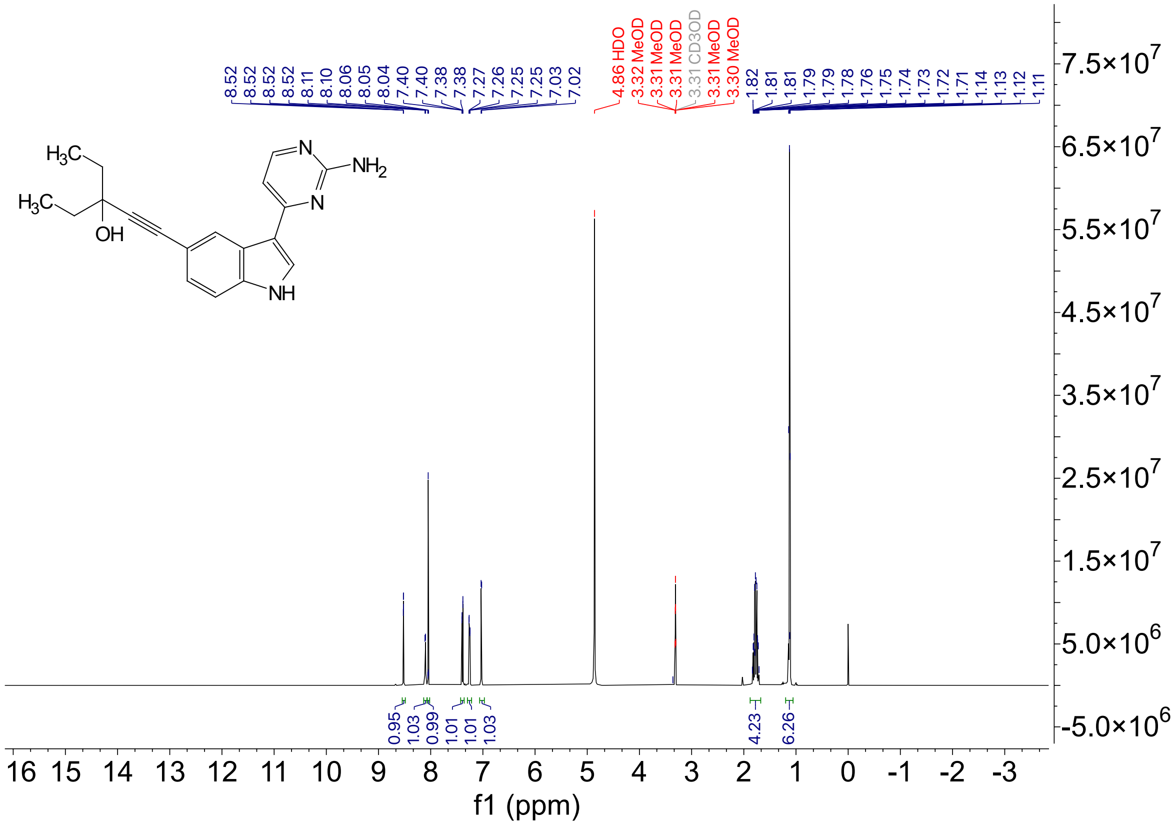

$^{13}C$ NMR (126 MHz, $CD_3OD$) for Compound **11**.

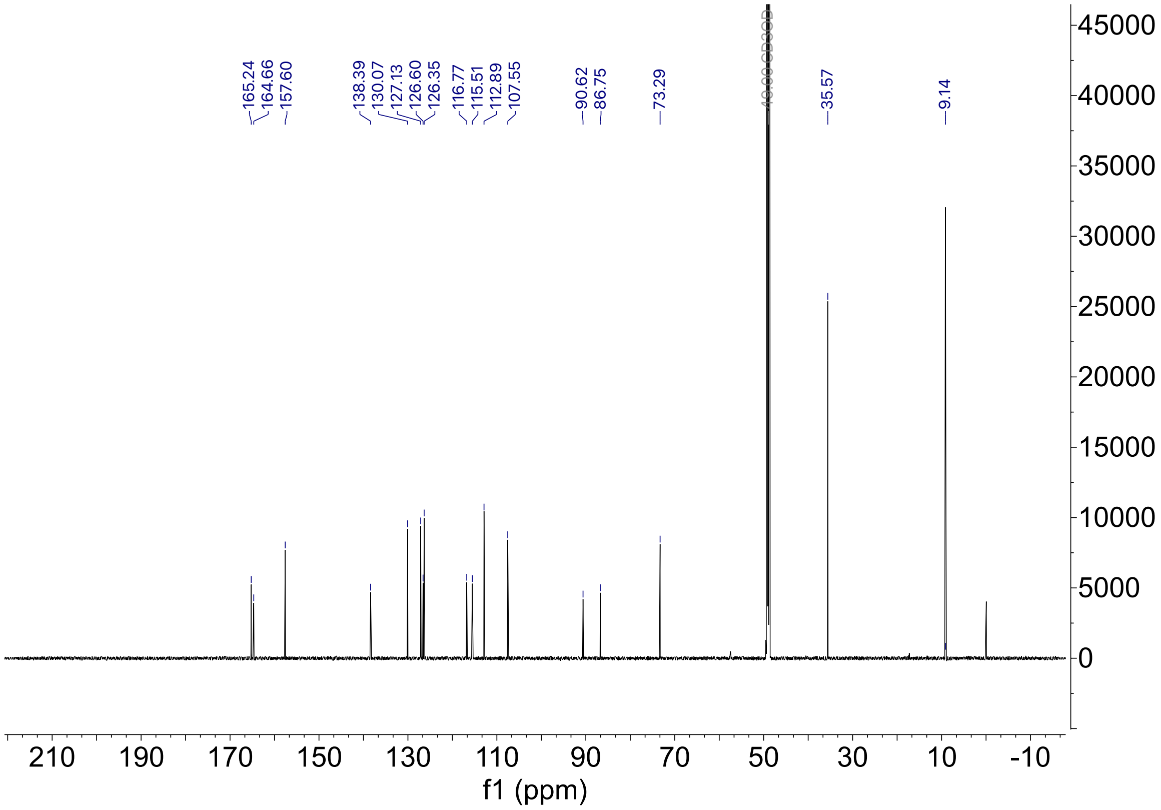

HPLC trace for compound **11**

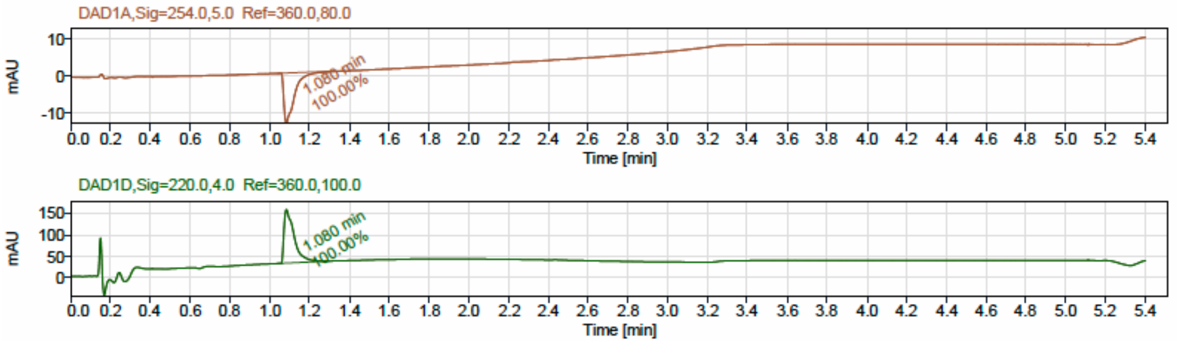


LCMS

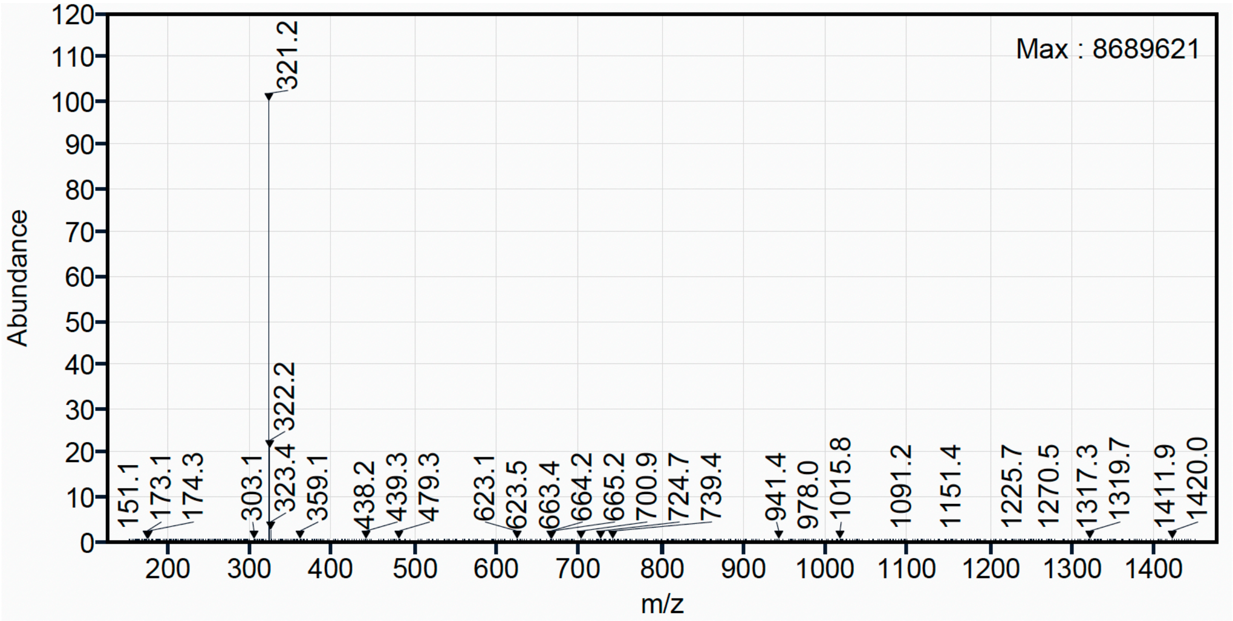

$^{1}H$ NMR (500 MHz, $CD_3OD$) for Compound **12**.

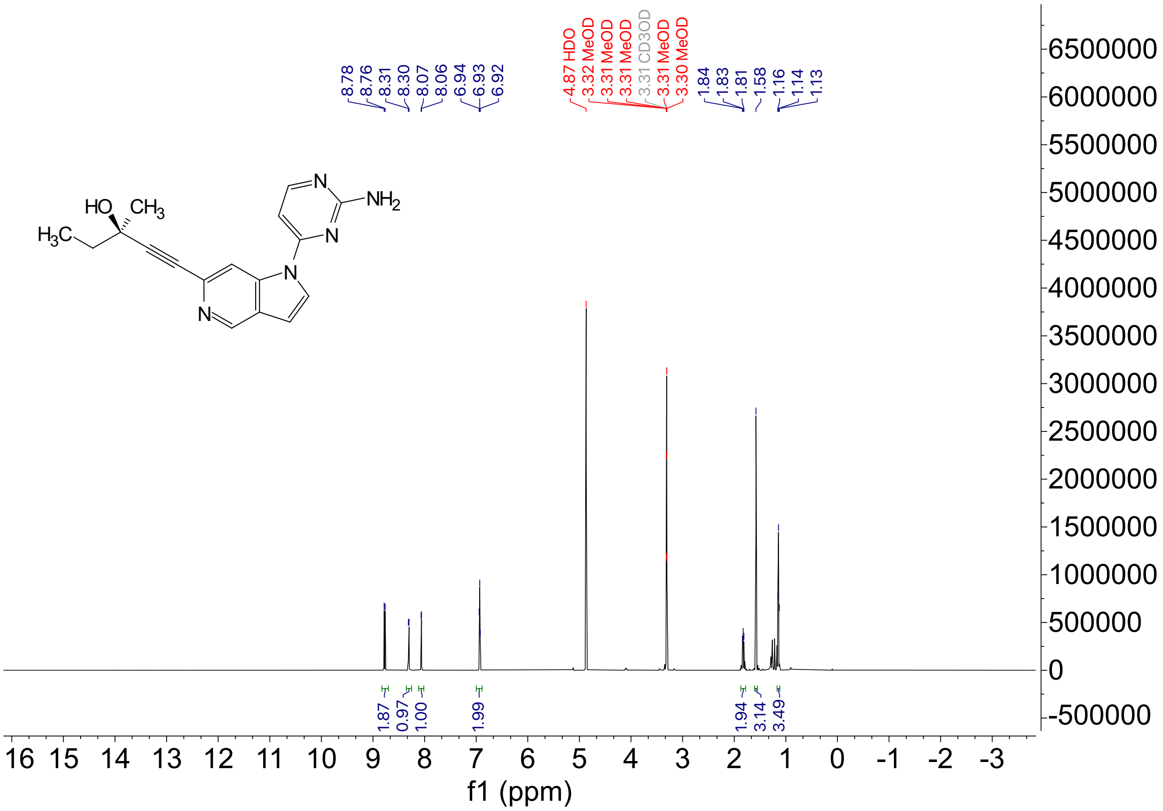

$^{13}C$ NMR (126 MHz, $CD_3OD$) for Compound **12.**

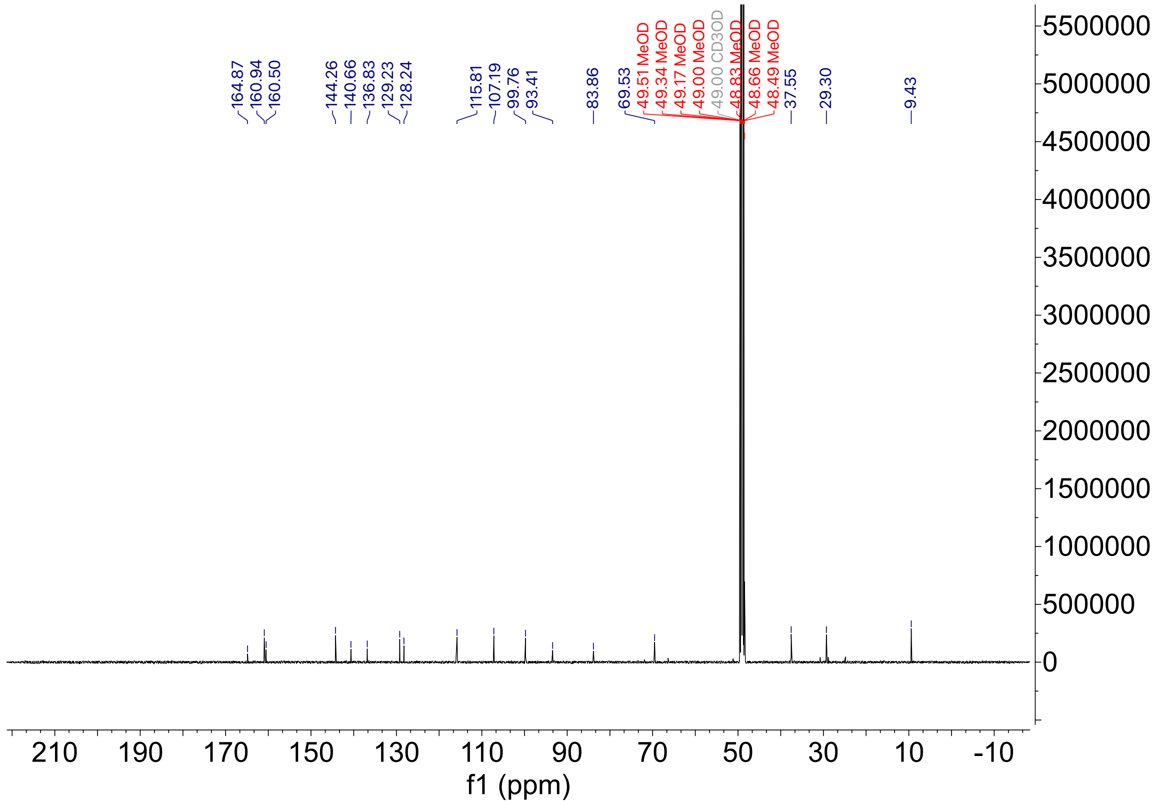

HPLC trace for compound **12**

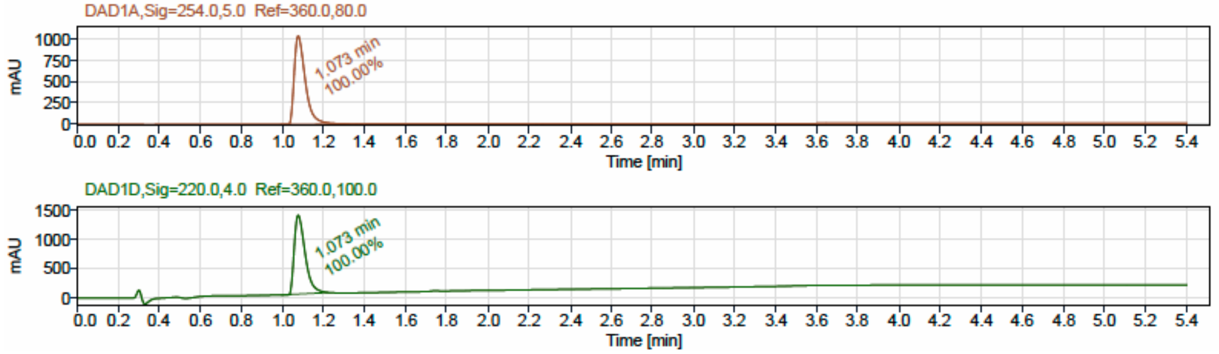


LCMS

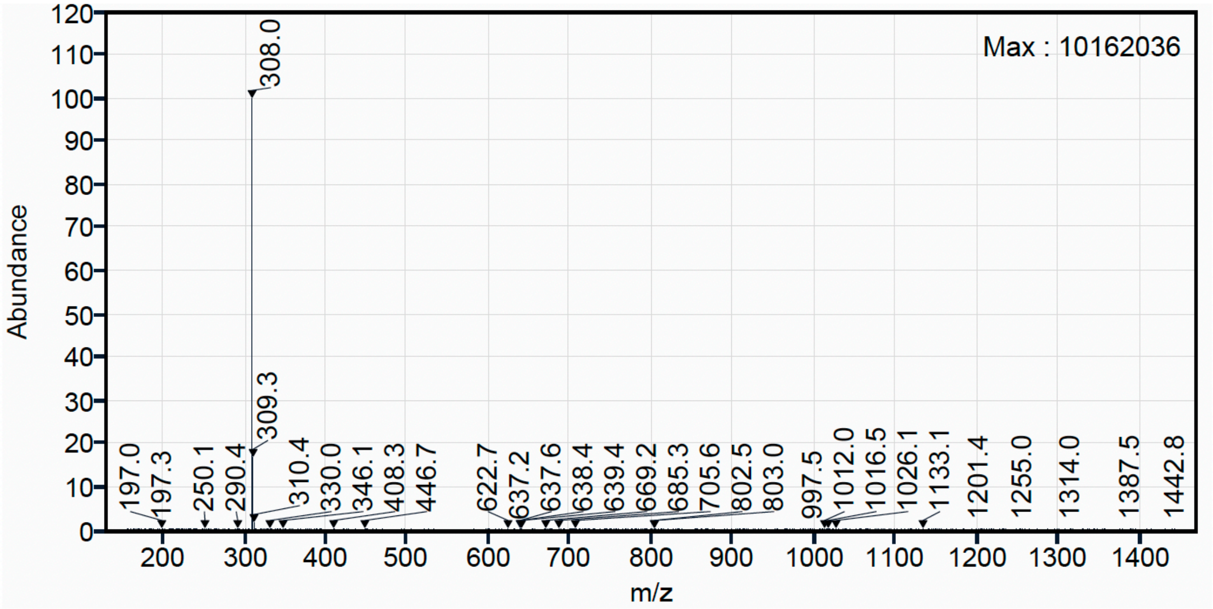

[1]H NMR (500 MHz, $CD_3OD$) for Compound **13**.

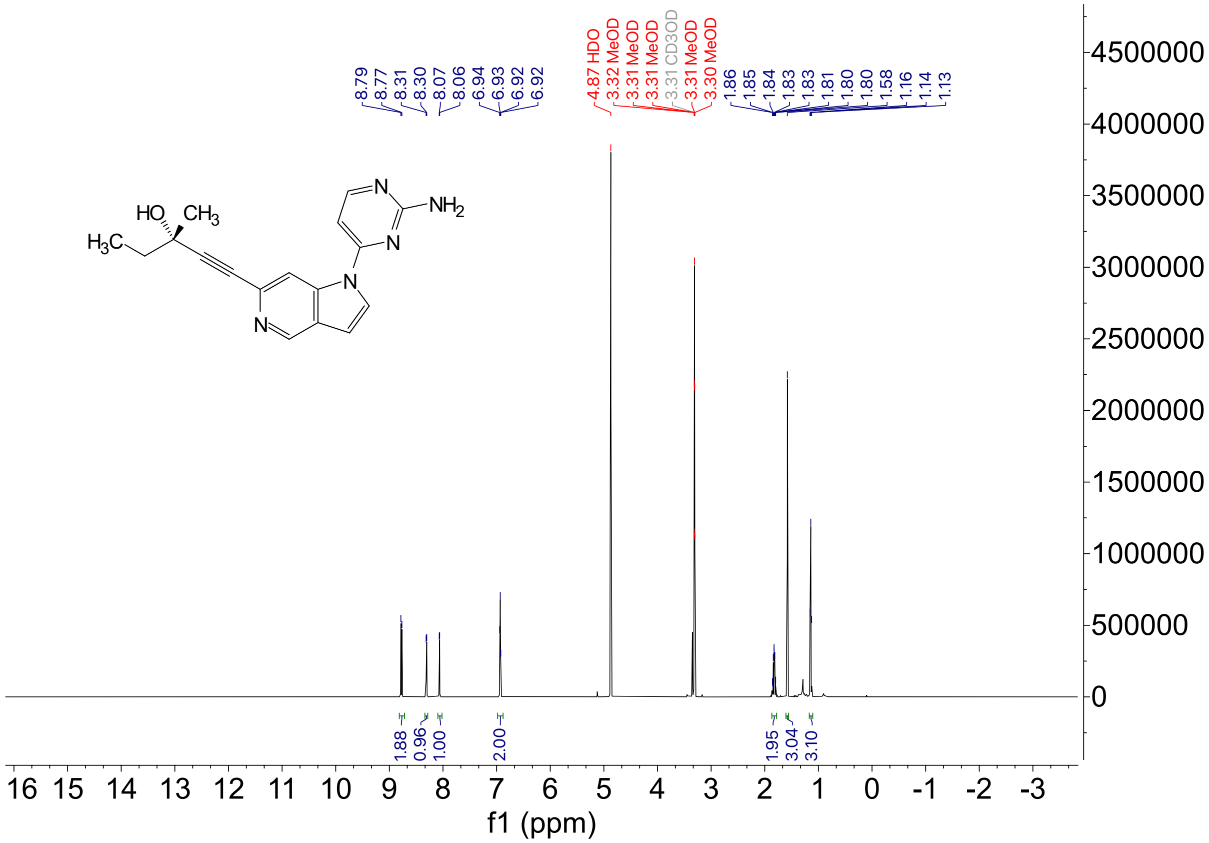

$^{13}C$ NMR (126 MHz, $CD_3OD$) for Compound **13**.

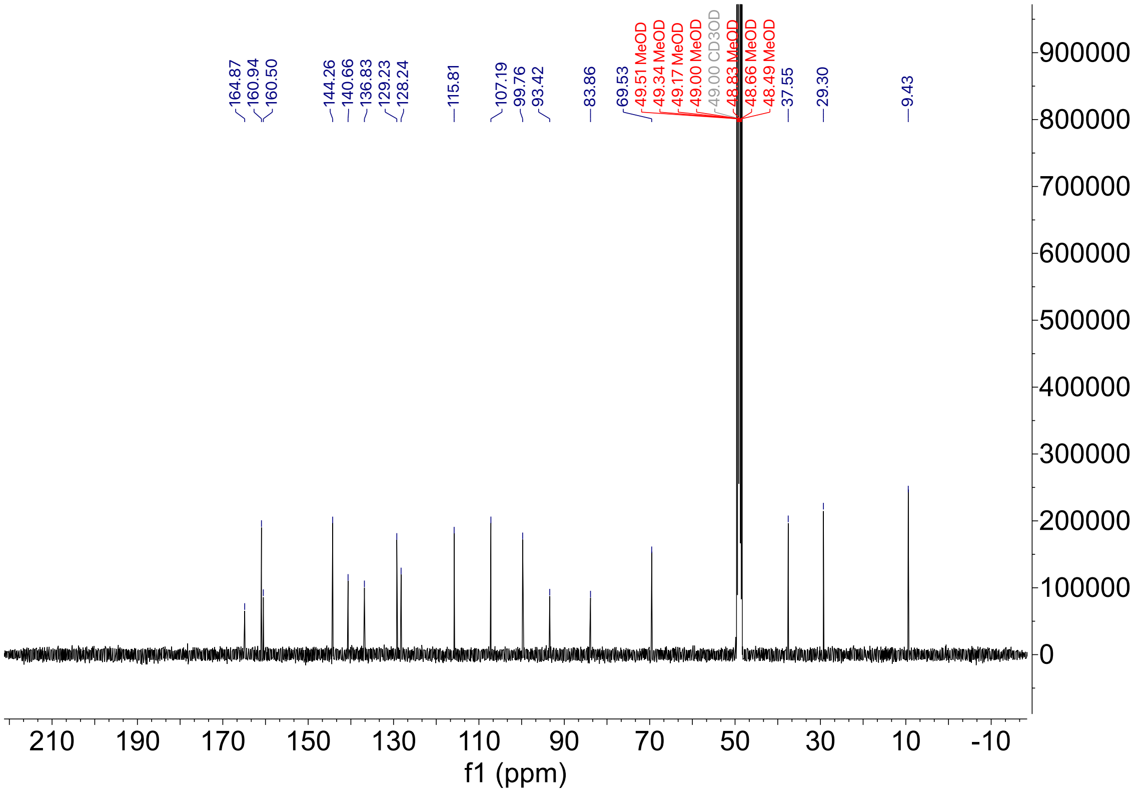

HPLC trace for compound **13**

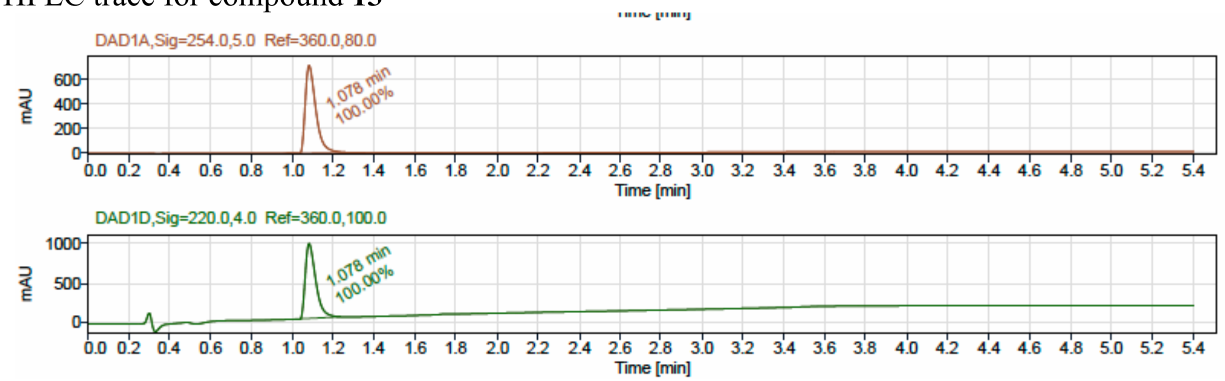


LCMS

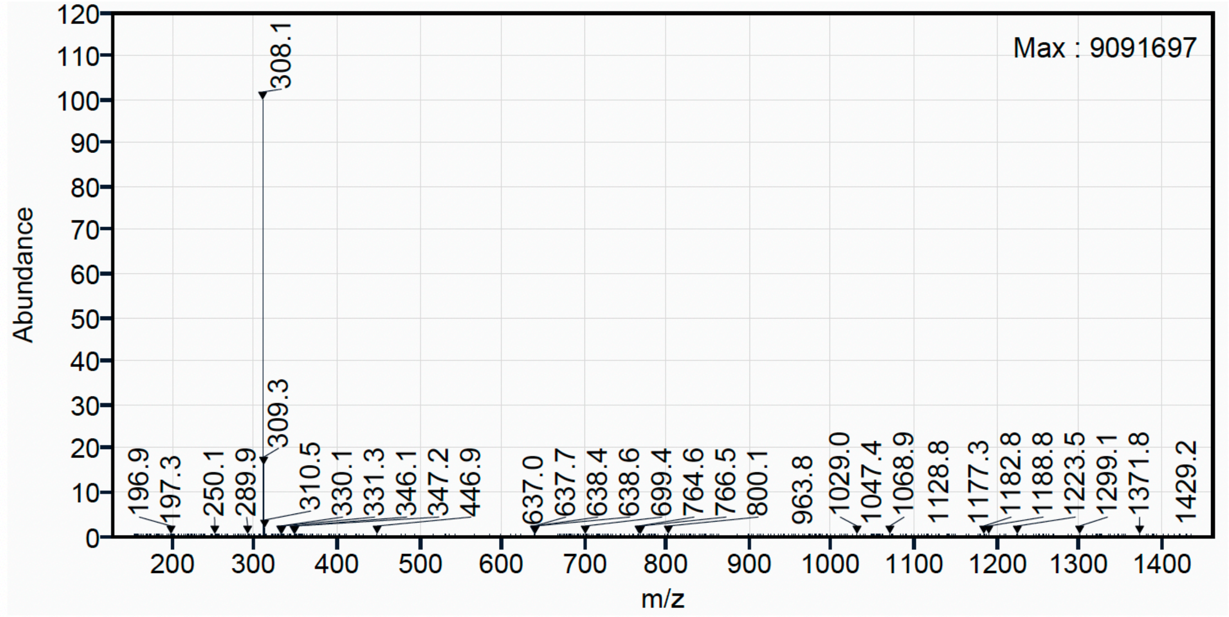

$^{1}H$ NMR (500 MHz, $CD_3OD$) for Compound **14**.

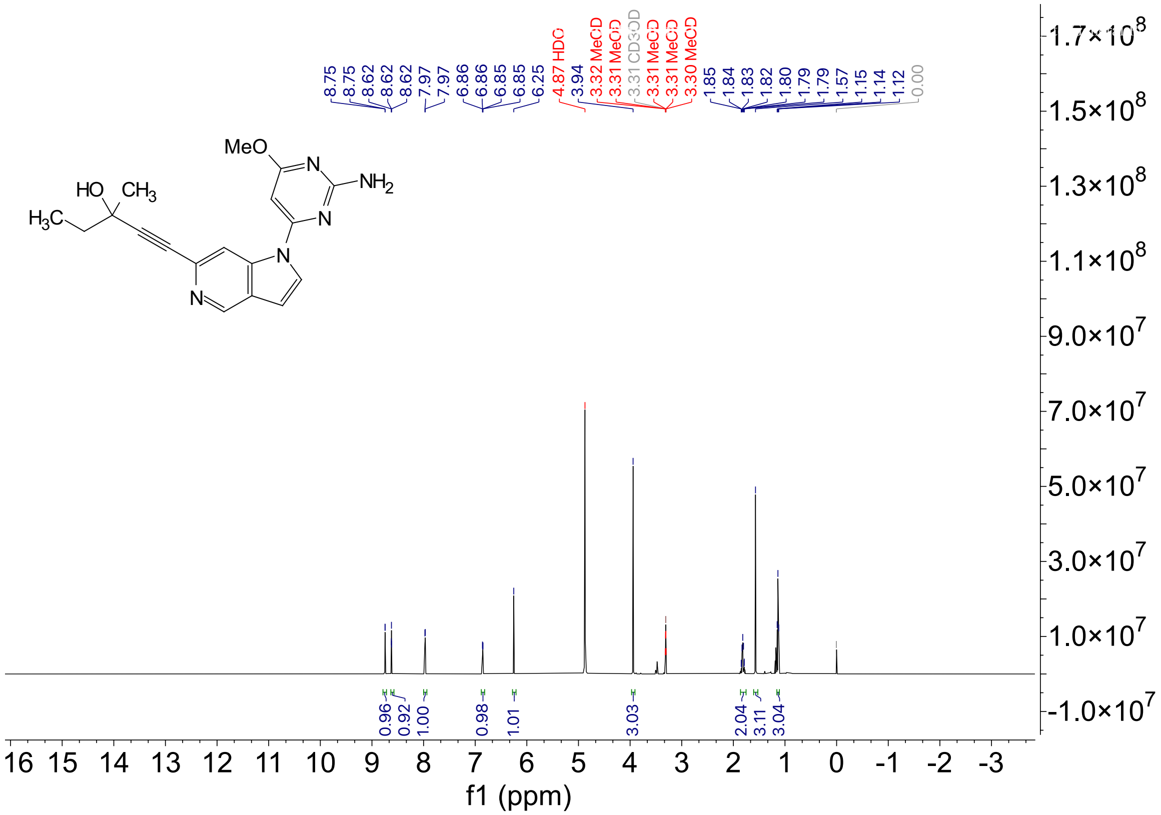

[13]C NMR (126 MHz, $CD_3OD$) for Compound **14**.

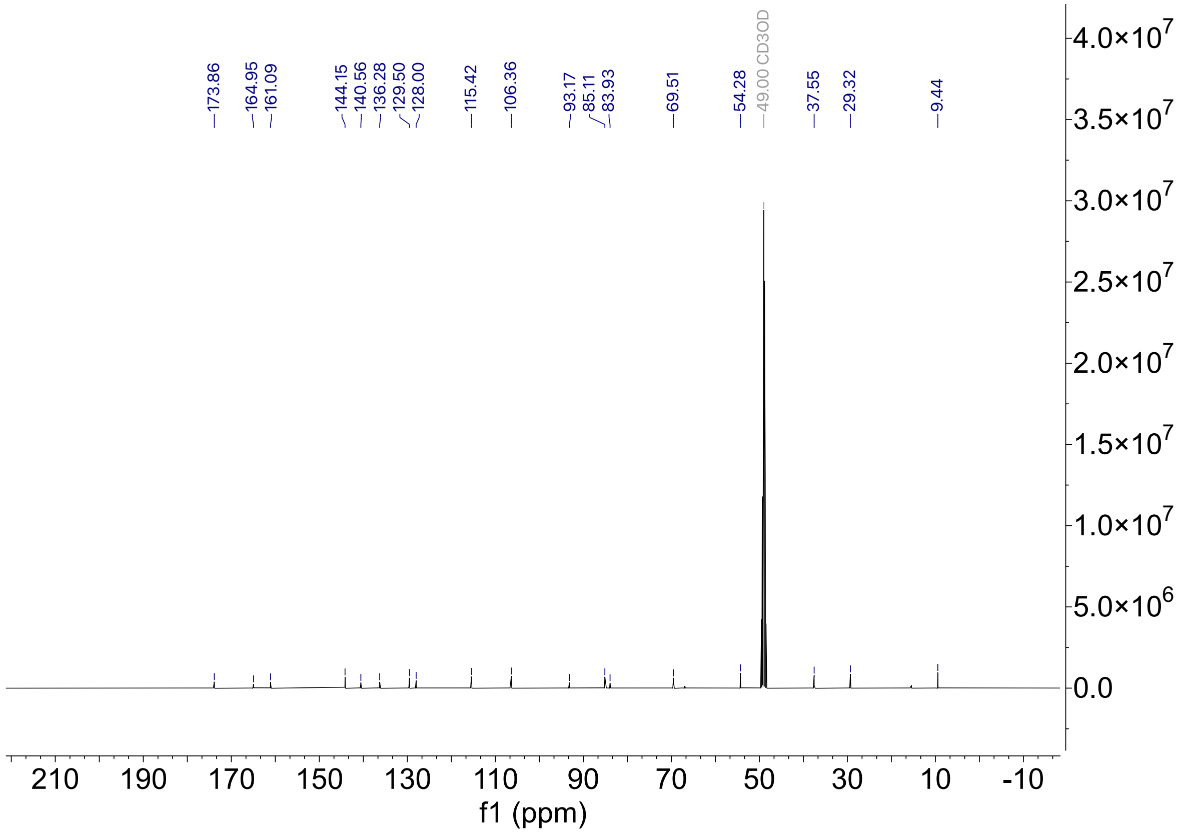

HPLC trace for compound **14**

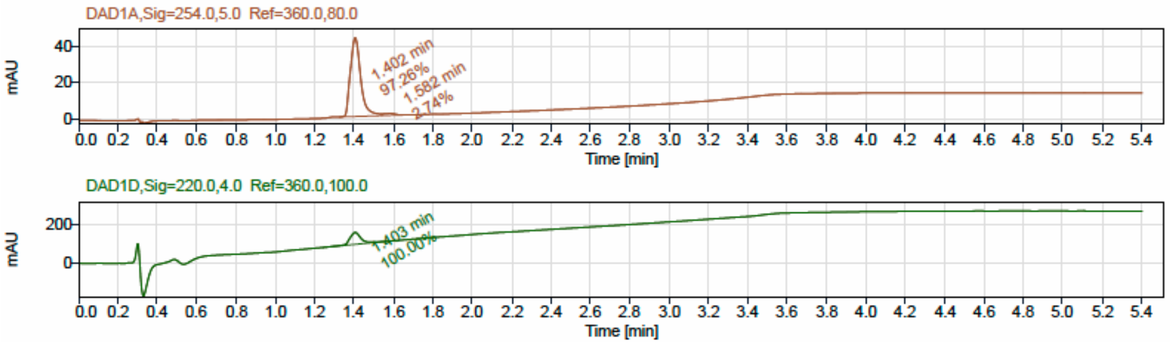


LCMS

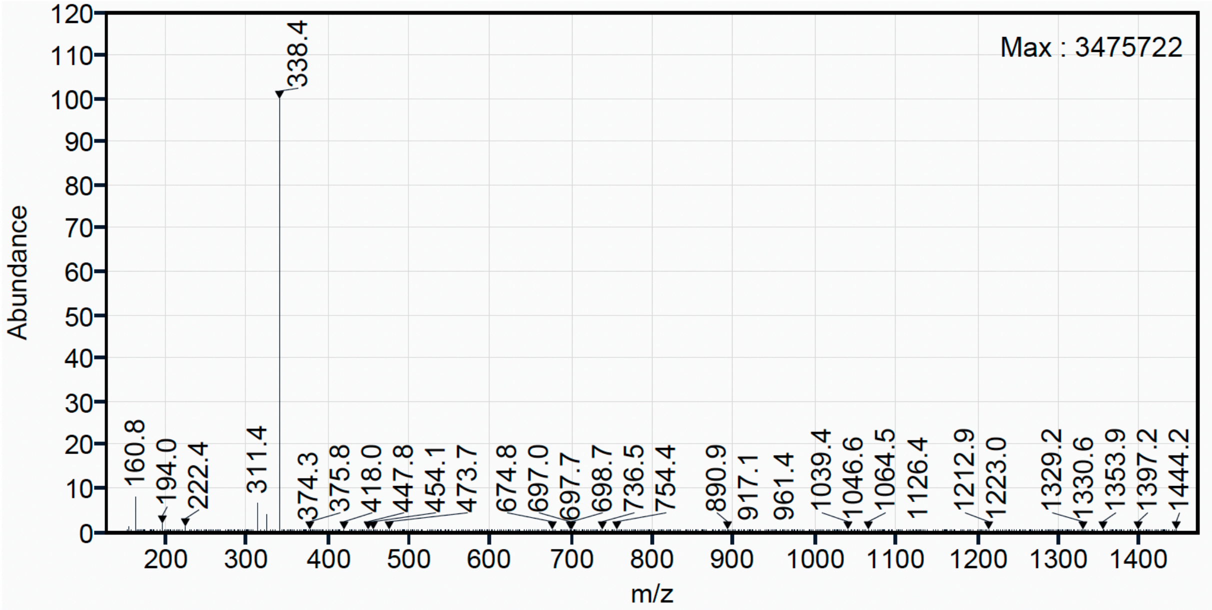

$^{1}H$ NMR (500 MHz, DMSO-$d_6$) for Compound **17**.

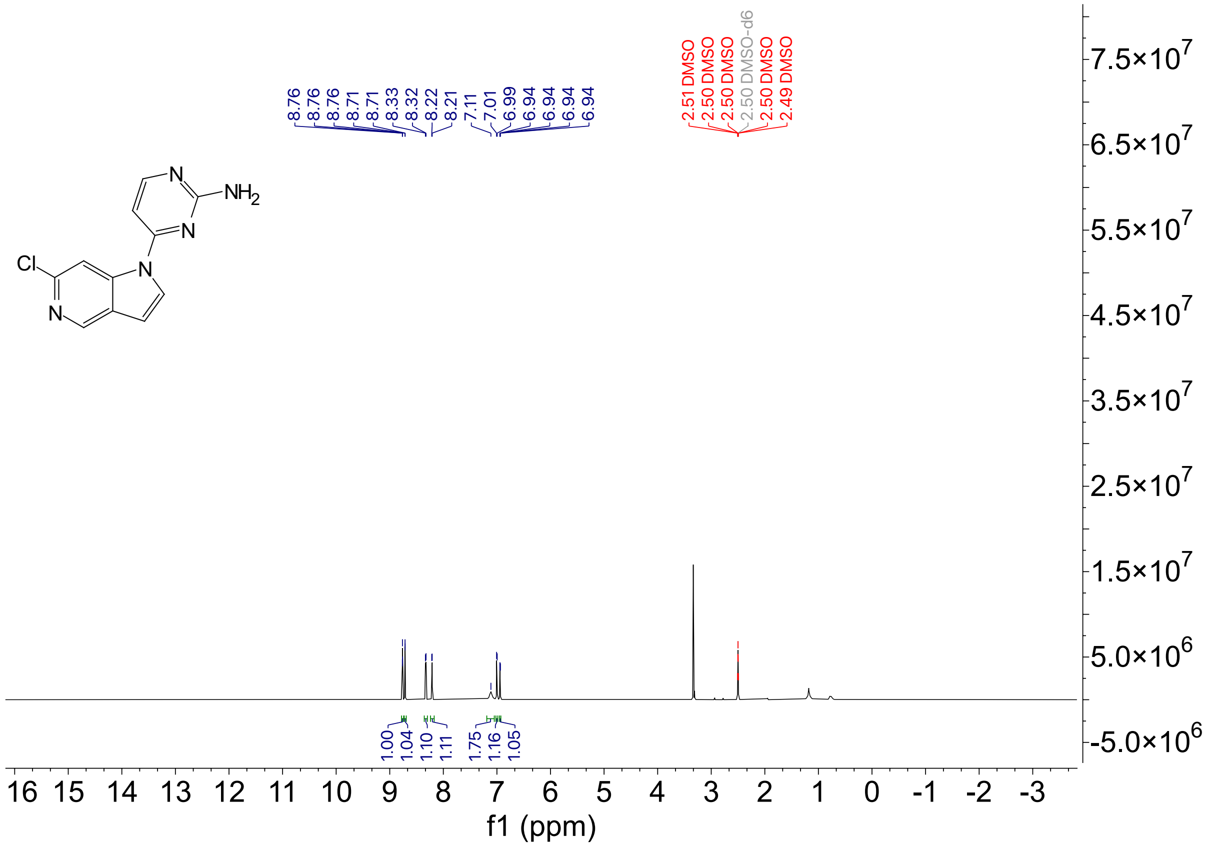

$^{13}C$ NMR (126 MHz, DMSO-$d_6$) for Compound **17.**

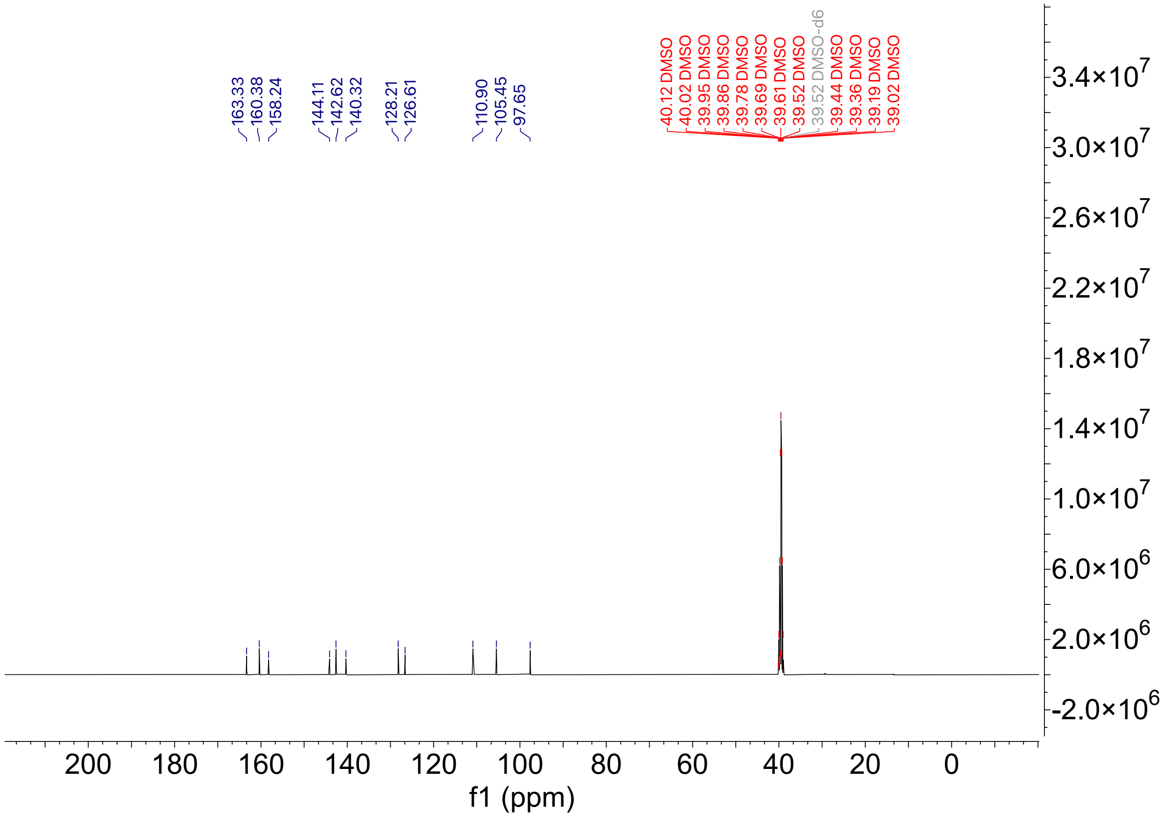

HPLC trace for compound **17**

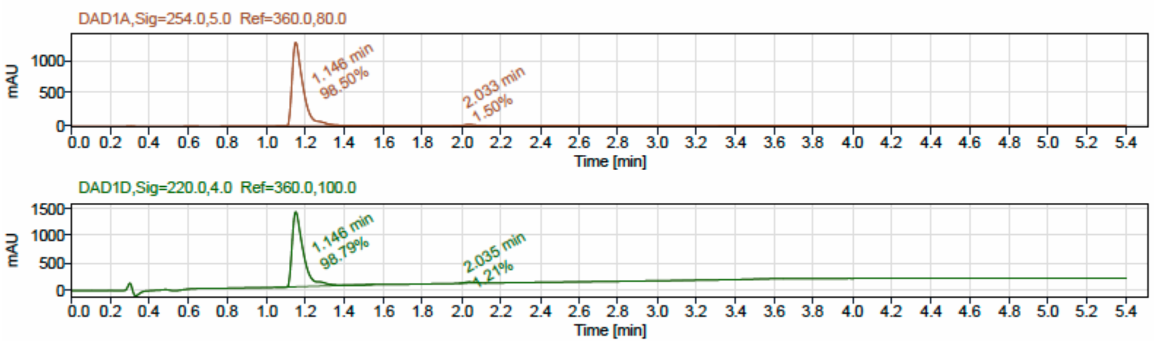


LCMS

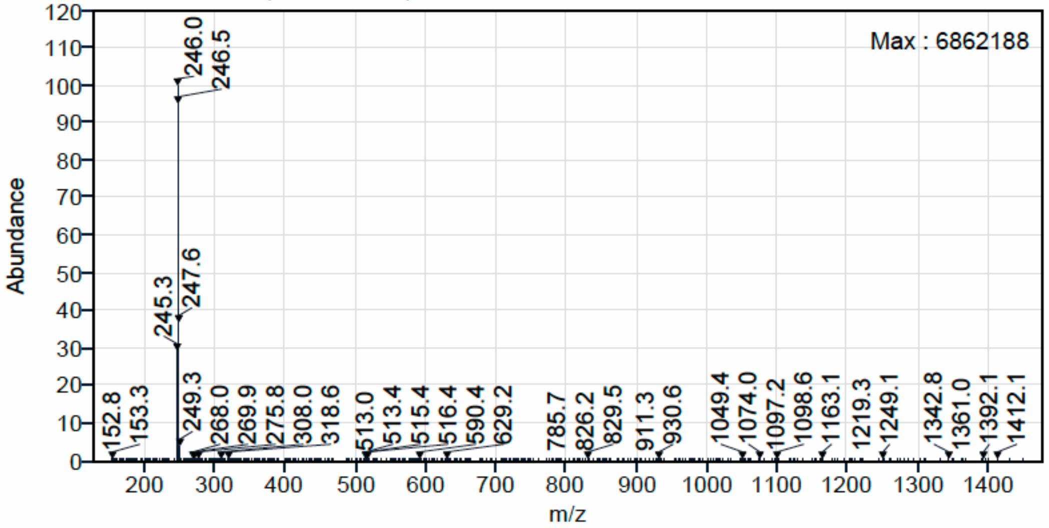

[1]H NMR (500 MHz, DMSO-$d_6$) for Compound **20**.

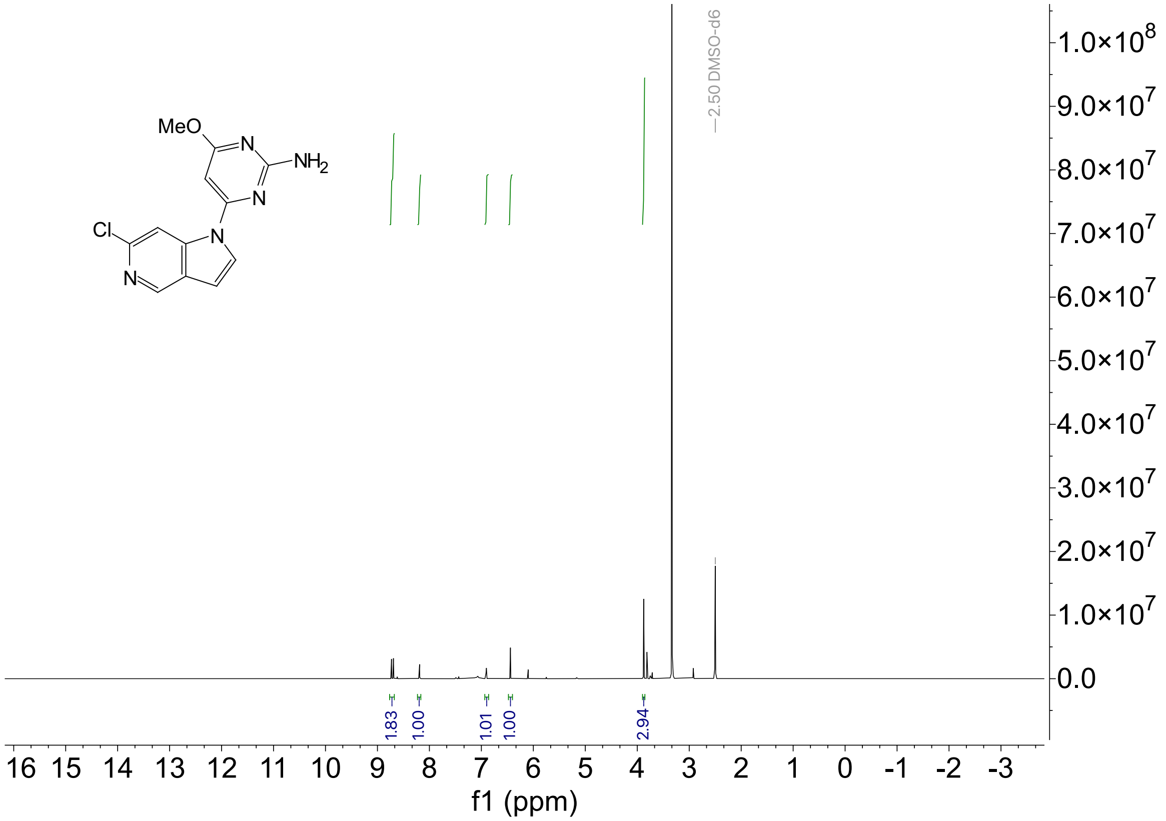

$^{13}$C NMR (126 MHz, DMSO-$d_6$) for Compound **20.**

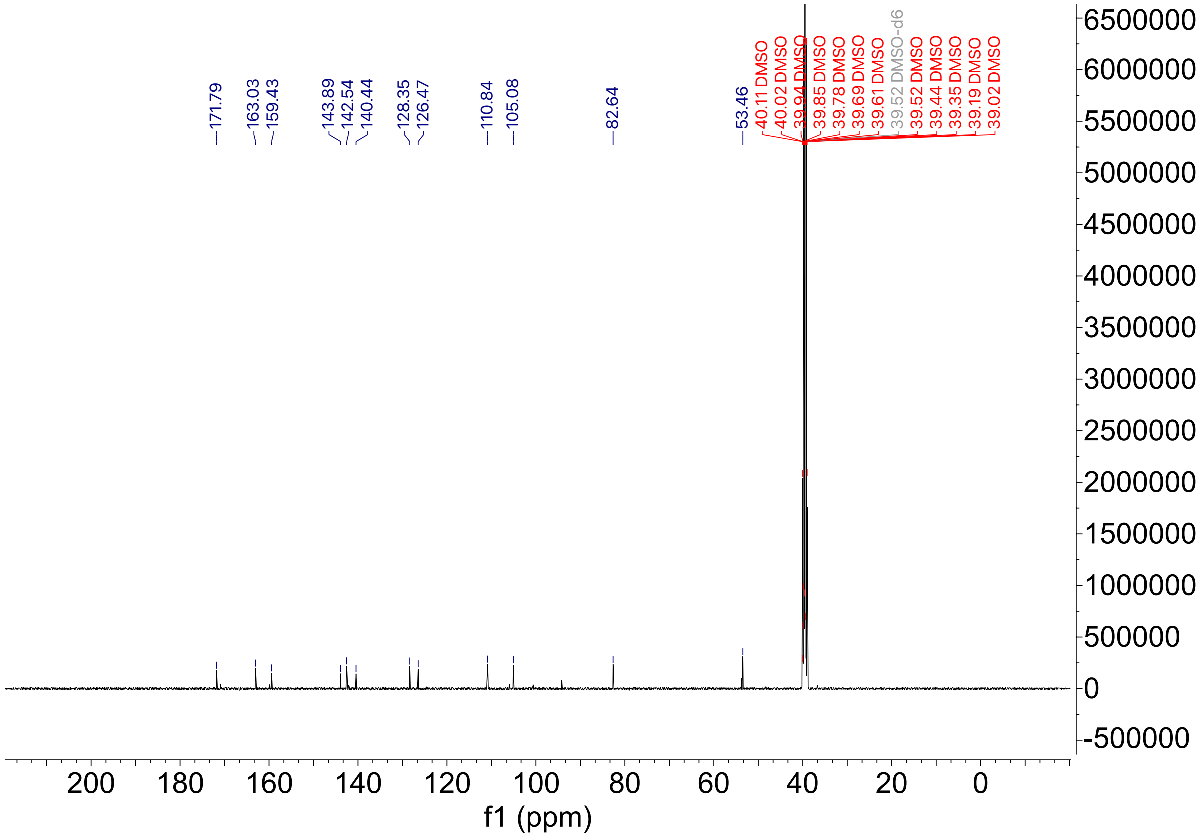

HPLC trace for compound **20**

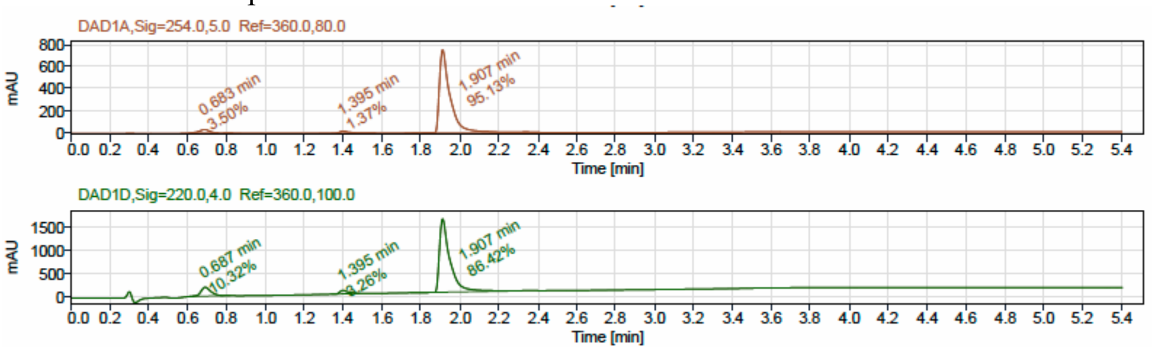


LCMS

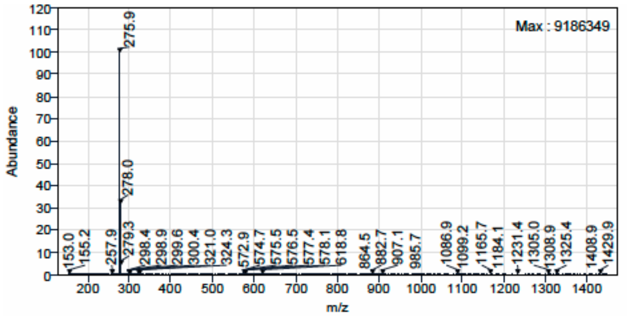

$^{1}H$ NMR (500 MHz, $CD_3OD$) for Compound **21**.

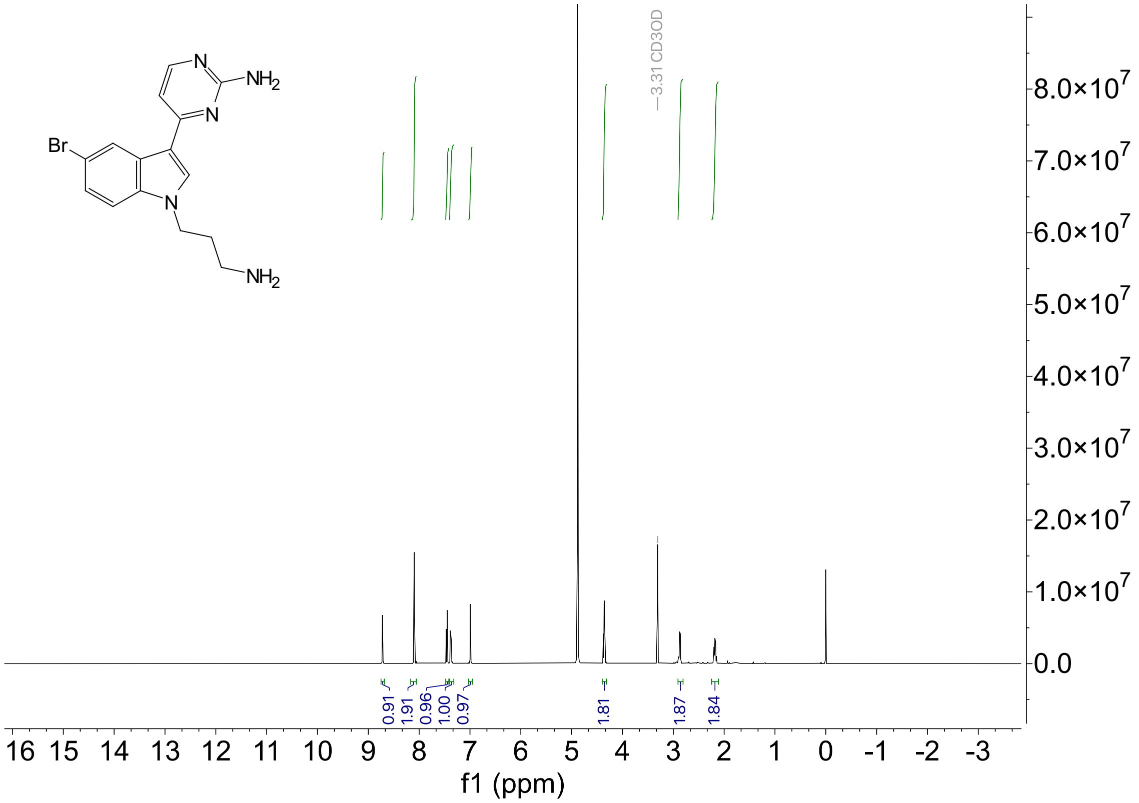

$^{13}C$ NMR (126 MHz, $CD_3OD$) for Compound **21**.

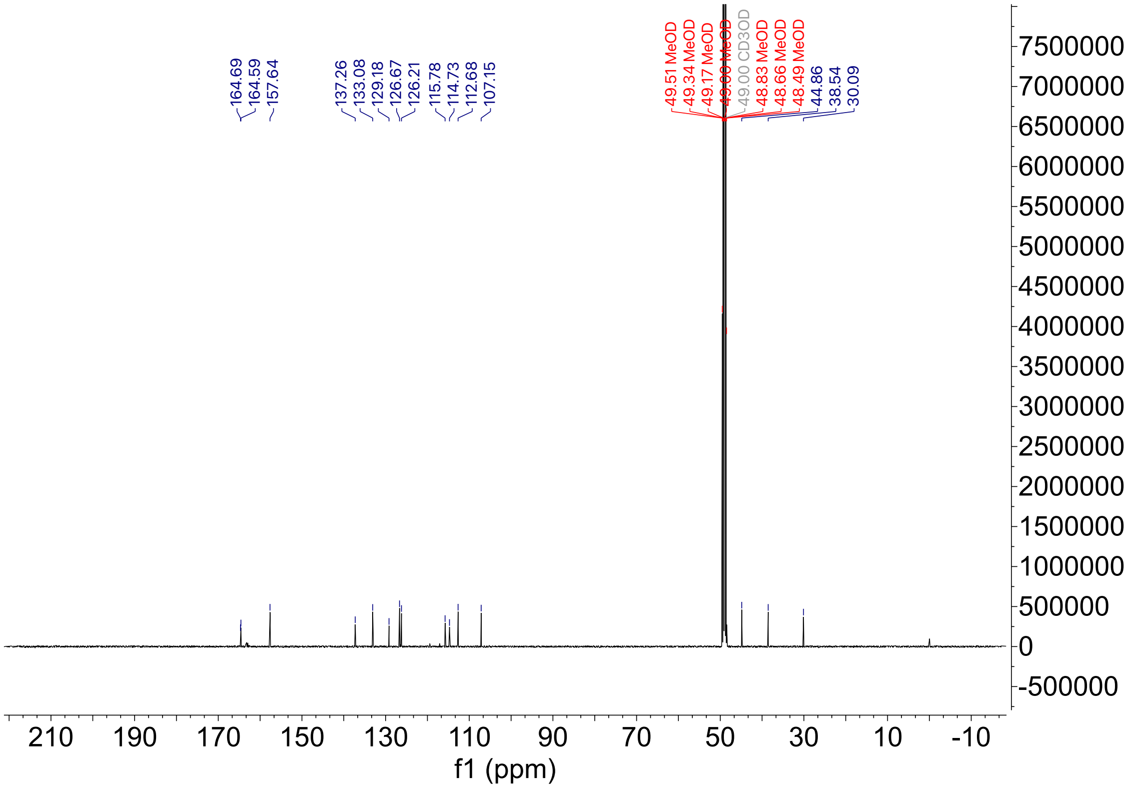

HPLC trace for compound **21**

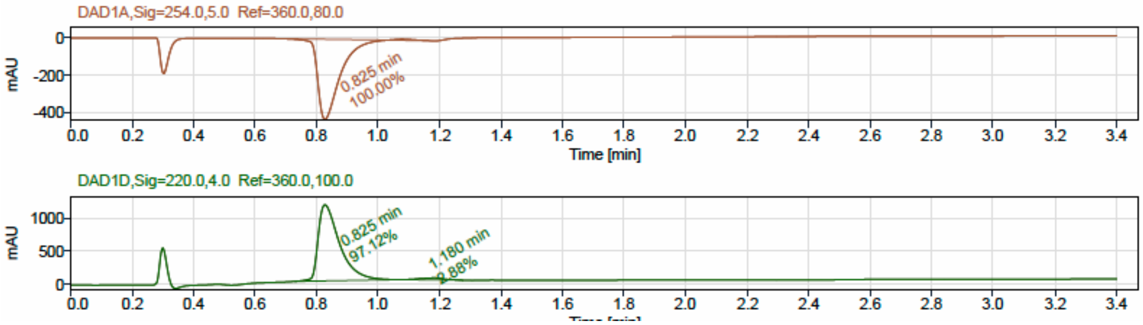


LCMS

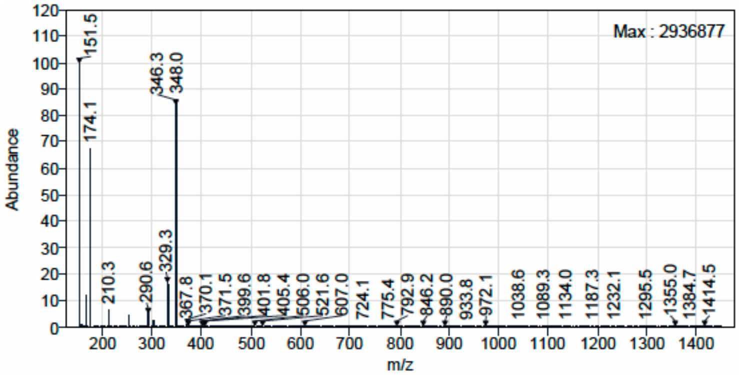

$^{1}H$ NMR (500 MHz, $CD_3OD$) for Compound **22**.

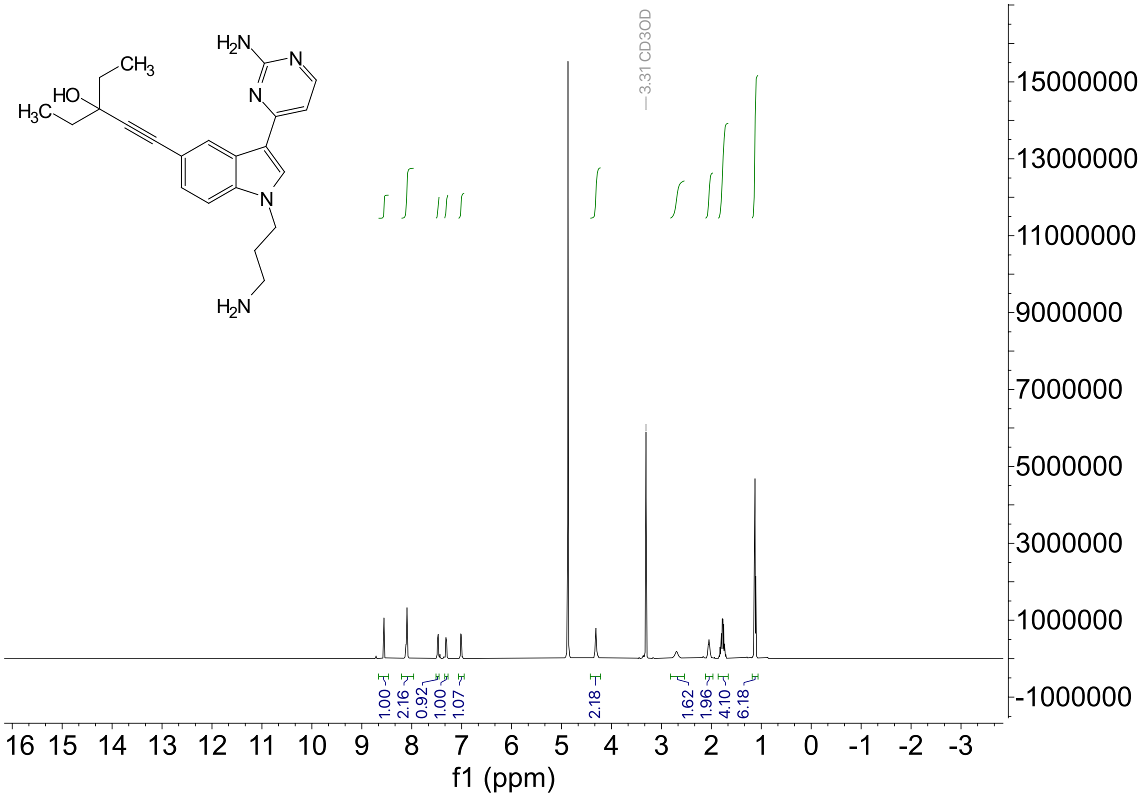

$^{13}C$ NMR (126 MHz, $CD_3OD$) for Compound **22**.

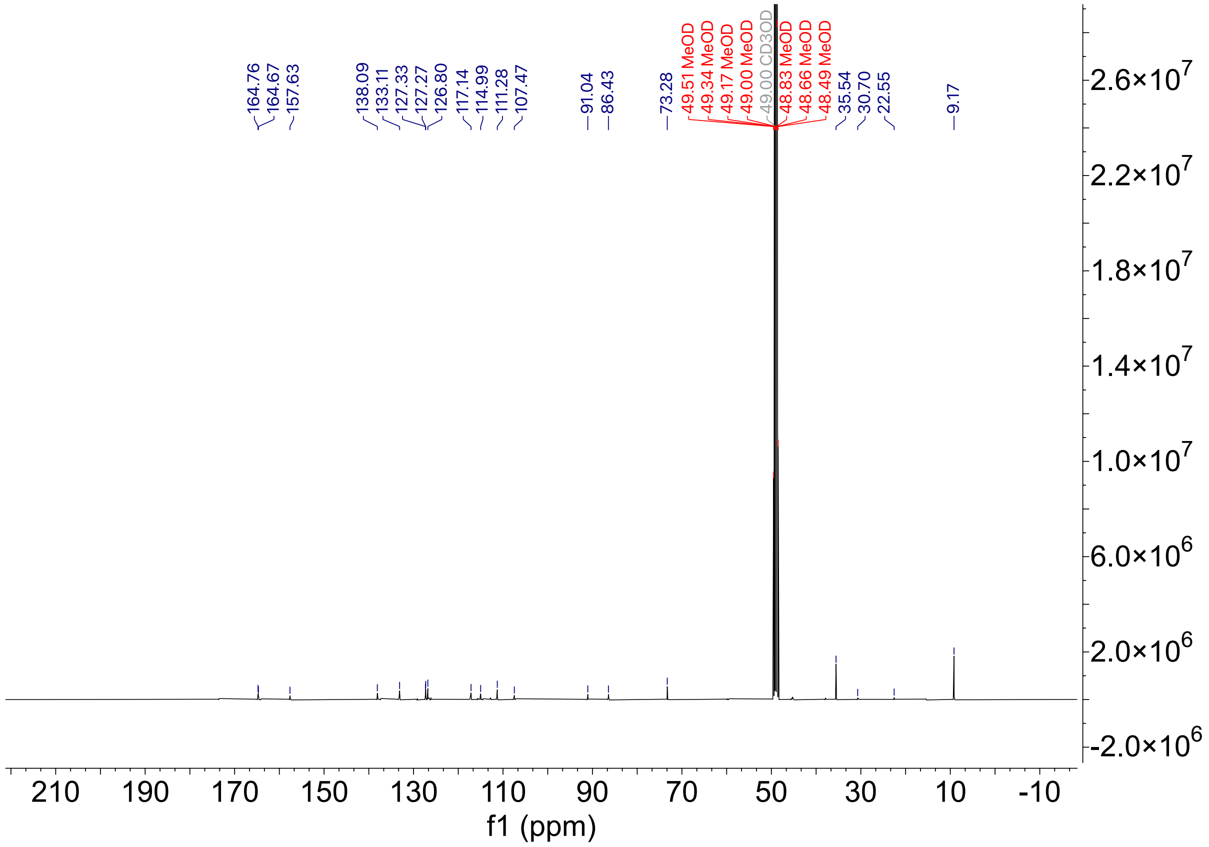

HPLC trace for compound **22**

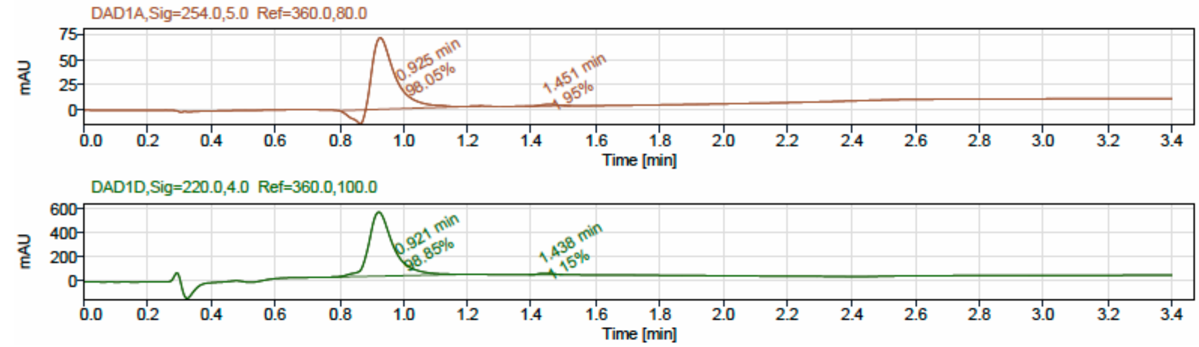


LCMS

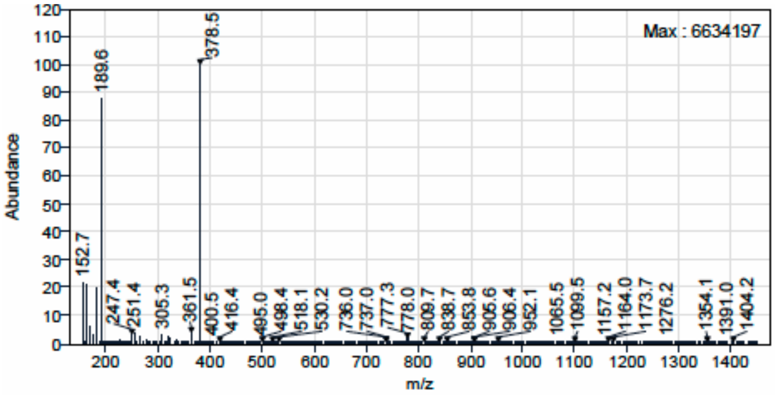

[1]H NMR (400 MHz, $CD_3OD$) for Compound **23.**

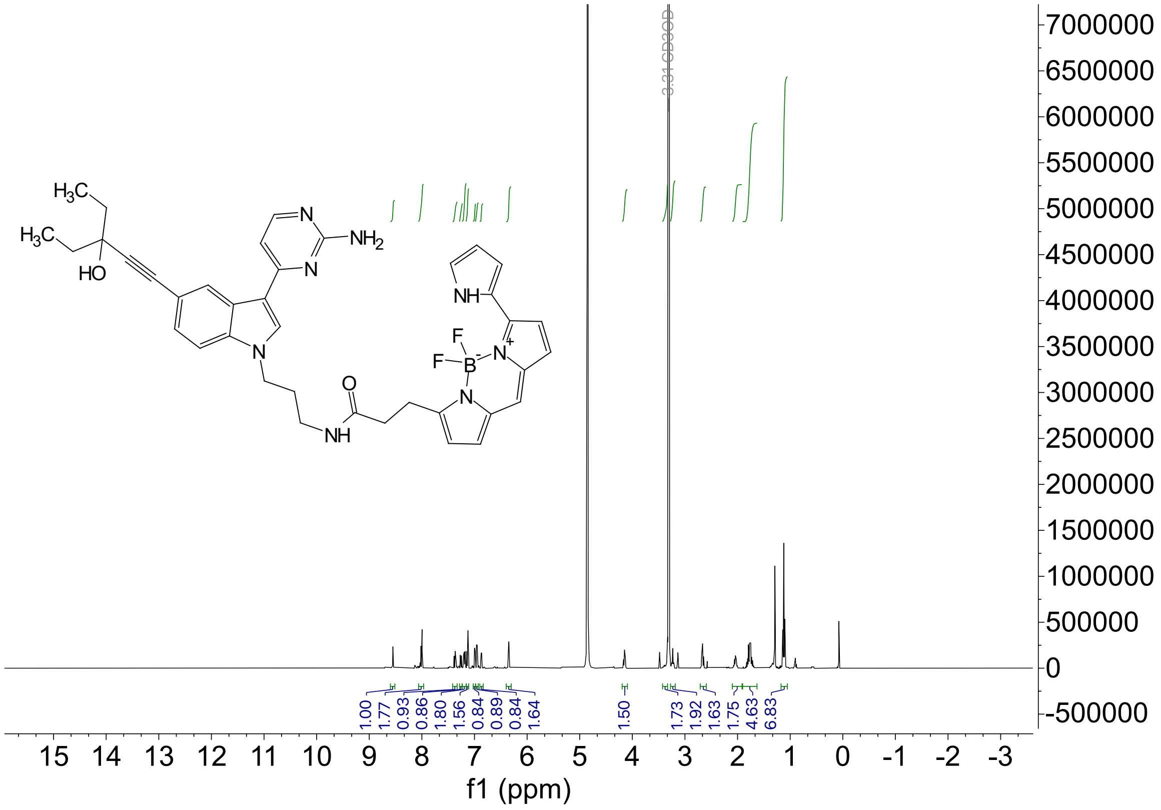

HPLC trace for compound **23**

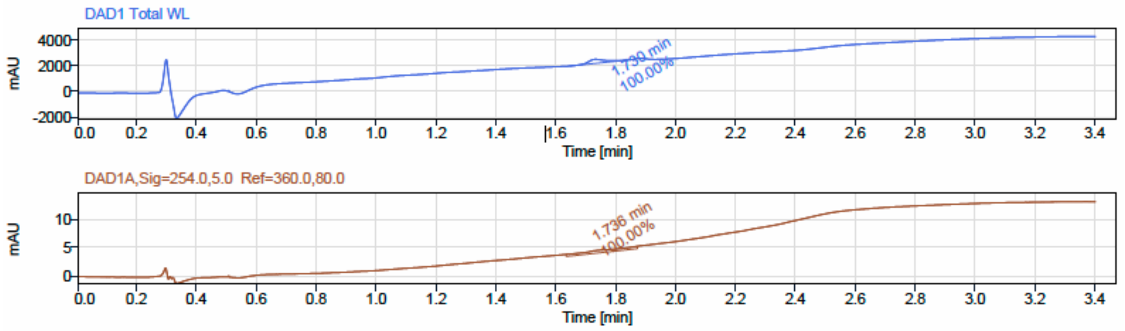


LCMS

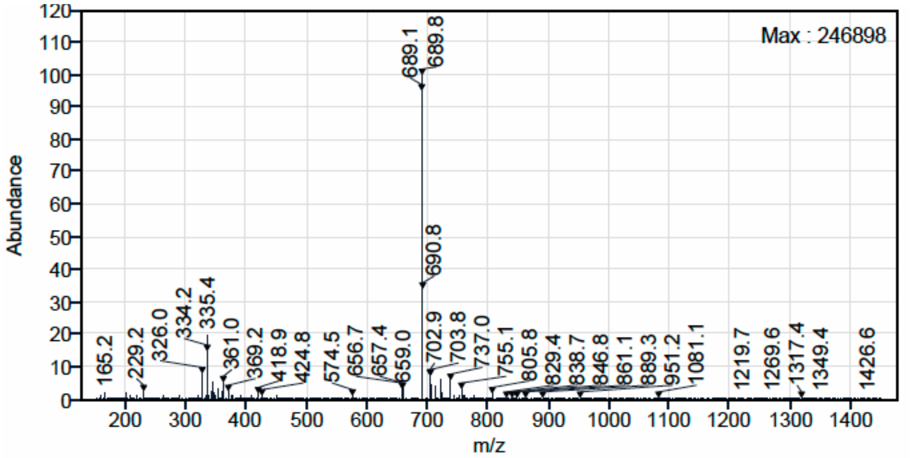

$^{1}$H NMR (400 MHz, $CD_3OD$) for Compound **24.**

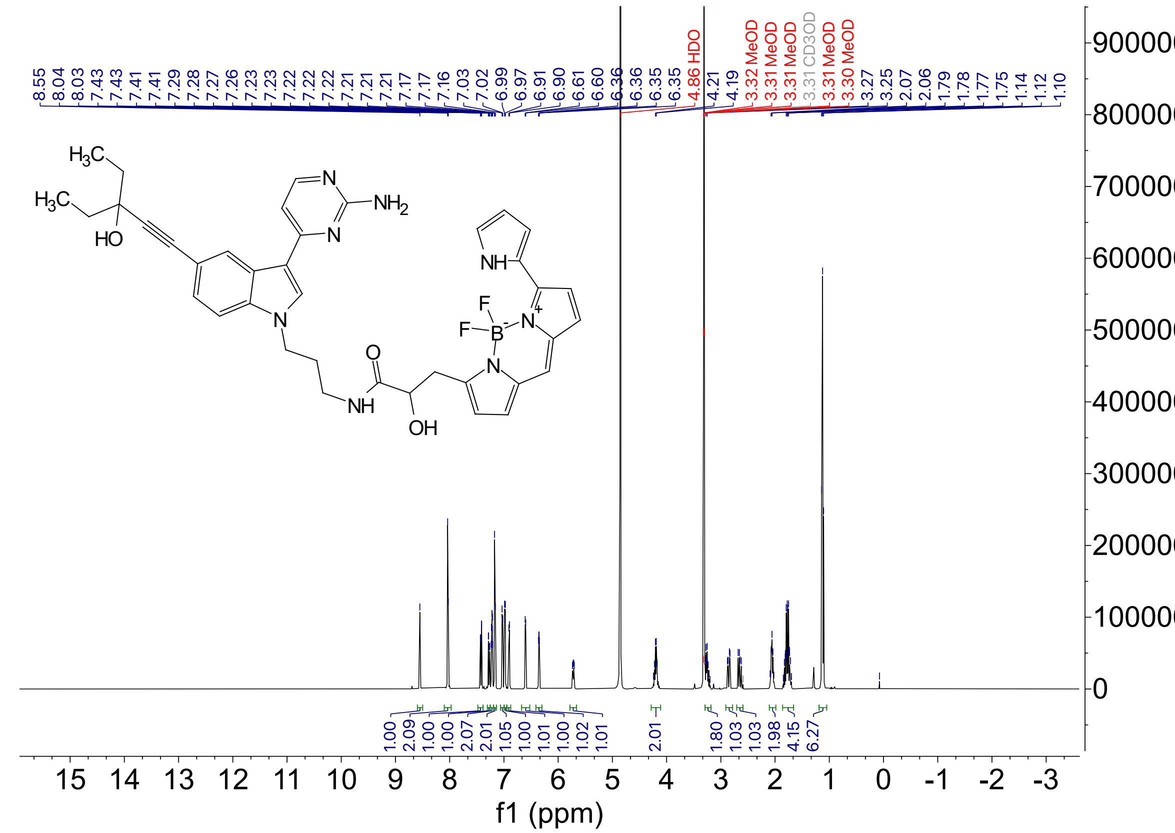

HPLC trace for compound **24**

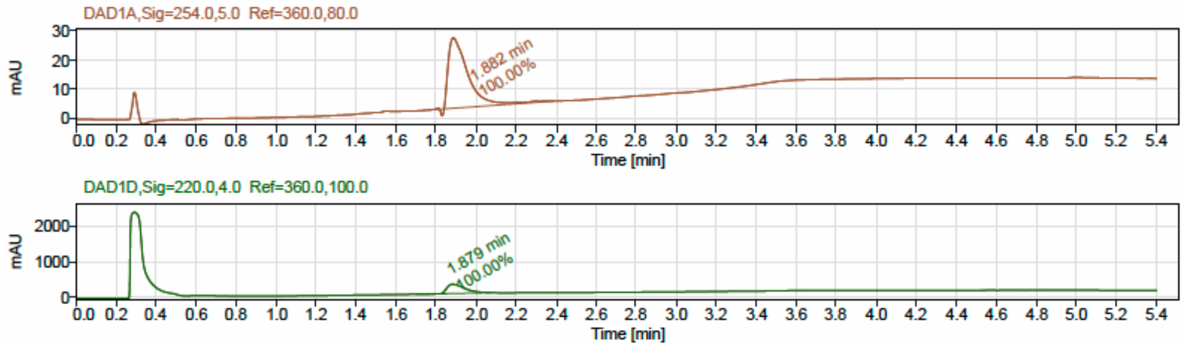


LCMS

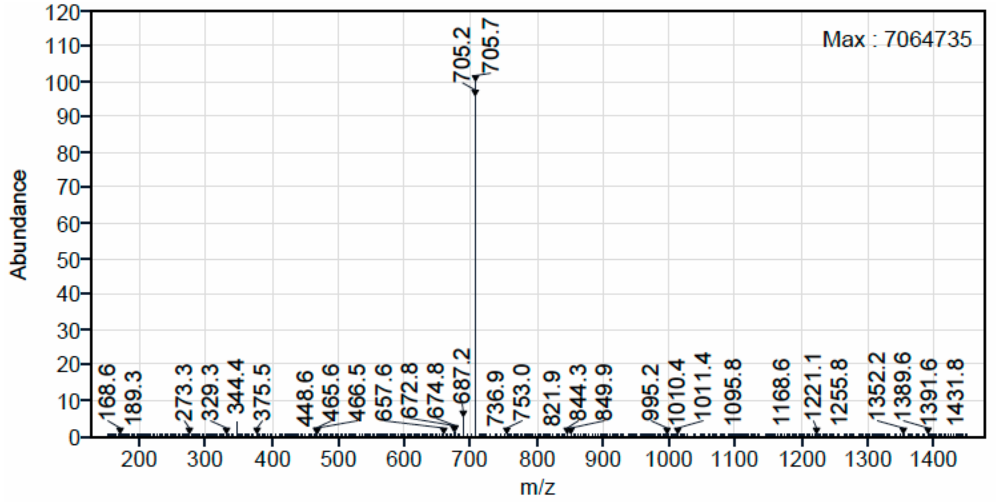

$^{1}H$ NMR (500 MHz, DMSO-$d_6$) for Compound **26**.

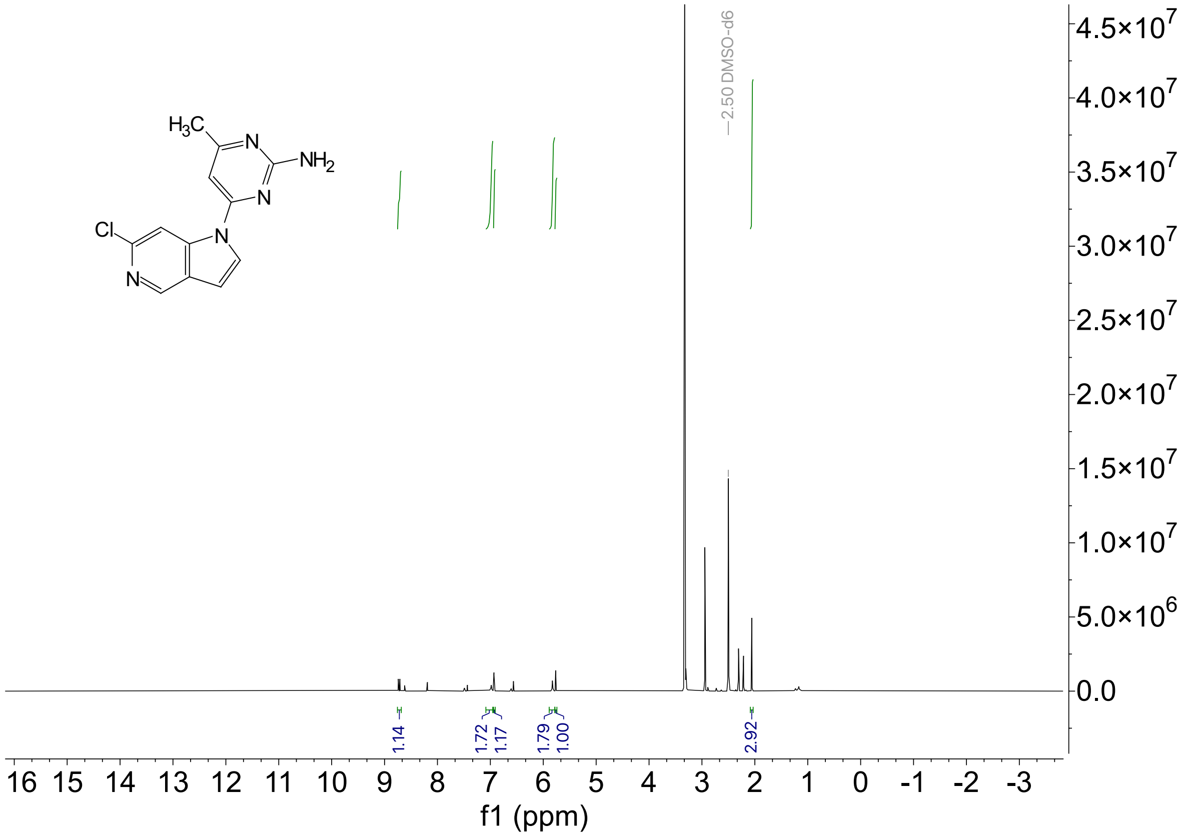

HPLC trace for compound **26**

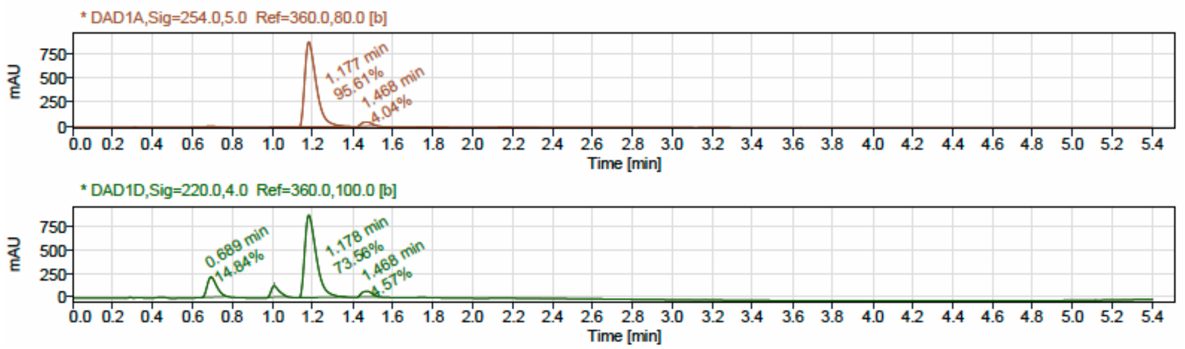


LCMS

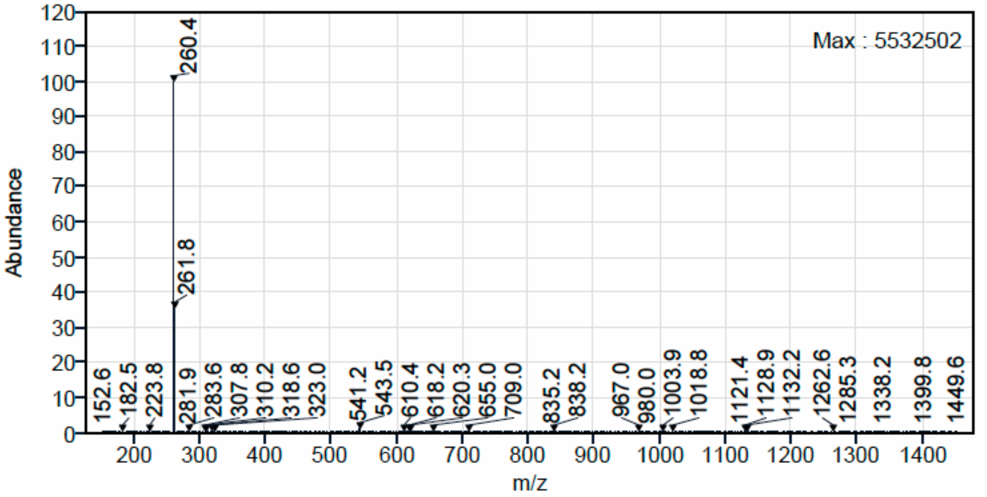
